# The ubiquitin ligase Ariadne-1 regulates NSF for neurotransmitter release

**DOI:** 10.1101/2020.01.23.916619

**Authors:** Juanma Ramírez, Miguel Morales, Nerea Osinalde, Imanol Martínez-Padrón, Ugo Mayor, Alberto Ferrús

**Affiliations:** Department of Biochemistry and Molecular Biology, Faculty of Science and Technology, UPV/EHU, Leioa 48940, Bizkaia, Spain; Cajal Institute, CSIC, Madrid 28002, Spain; Department of Biochemistry and Molecular Biology, Faculty of Pharmacy, UPV/EHU, Vitoria-Gasteiz 01006, Araba, Spain; Ikerbasque, Basque Foundation for Science, Bilbao 48013, Bizkaia, Spain

**Keywords:** Ariadne-1, *Drosophila*, E3 ubiquitin ligase, neurotransmitter release, NSF, synapse, ubiquitination

## Abstract

Ariadne-1 (Ari-1) is an essential E3 ubiquitin-ligase whose neuronal substrates are yet to be identified. We have used an *in vivo* ubiquitin biotinylation strategy coupled to quantitative proteomics to identify putative Ari-1 substrates in *Drosophila* heads. Sixteen candidates met the established criteria. Amongst those, we identified Comatose (Comt), the homologue of the N-ethylmaleimide sensitive factor (NSF). Using an *in vivo* GFP pulldown approach, we validate Comt/NSF to be an ubiquitination substrate of Ari-1 in fly neurons. The interaction results in the monoubiquitination of Comt/NSF. We also report that Ari-1 loss of function mutants display a lower rate of spontaneous neurotransmitter release due to failures at the pre-synaptic side. By contrast, evoked release in Ari-1 mutants is enhanced in a Ca^2+^ dependent manner without modifications in the number of active zones, indicating that the probability of release per synapse is increased in these mutants. The distinct Ari-1 mutant phenotypes in spontaneous versus evoked release indicate that NSF activity discriminates the two corresponding protein ensembles that mediate each mode of release. Our results, thus, provide a mechanism to regulate NSF activity in the synapse through Ari-1-dependent ubiquitination.

---

Neurotransmitter release is mediated by a set of protein-protein interactions that include the N-ethylmaleimide sensitive factor (NSF), soluble NSF attachment proteins (SNAPs) and SNAP receptors (SNAREs) (1). These proteins assemble into a tripartite complex in order to elicit synaptic vesicle fusion, which is formed by one synaptic vesicle membrane SNARE protein (v-SNARE), Synaptobrevin, and two plasma membrane SNARE proteins (t-SNAREs), Syntaxin and the 25-kDa synaptosome-associated protein (2). Following vesicle fusion, the tripartite SNARE complex disassembles by the activities of NSF and SNAPs. Free t-SNAREs from the plasma membrane can then participate in new priming reactions, while the v-SNAREs can be incorporated into recycled synaptic vesicles (3). These interactions, also routinely used for intracellular vesicle trafficking in all cell types, are conserved across species (4), including *Drosophila* (5).

Deviations on the rate of neurotransmitter release are at the origin of multiple neural diseases, including Parkinson’s disease (6). Under physiological conditions, the leucine-rich repeat serine/threonine-protein kinase 2 (LRRK2) phosphorylates NSF to enhance its ATPase activity, which facilitates the disassembly of the SNARE complex (7–9). However, the most common Parkinson’s disease mutation in LRRK2 causes an excess of kinase activity (10) that interferes with the vesicle recycling (11). Similarly, α-Synuclein, another Parkinson’s disease protein (12), alters neurotransmitter release by preventing the v-SNARE vesicle-associated membrane protein (VAMP)-2, also known as Synaptobrevin-2, from joining the SNARE complex cycle (13). Correct neural functioning, therefore, requires delicate regulation in vesicle trafficking. This regulation can be achieved by post-translational modifications, such as ubiquitination. In fact, ubiquitination of certain proteins can affect their activity or lifespan (14, 15). At the pre-synaptic side, for example, increased neurotransmitter release correlates with decreased protein ubiquitination (16). Similarly, acute pharmacological proteasomal inhibition causes rapid strengthening of neurotransmission (17).

Ariadne 1 (Ari-1) is an E3 ubiquitin-ligase, first identified in *Drosophila* (18), from a conserved gene family defined by two C_3_HC_4_ *Ring* fingers separated by a C_6_HC in-*Between*-*Rings* domain, the RBR motif (19). Ari-1 had been described to be essential for neuronal development, and its mutants reported to exhibit reduced eye rhabdomere surface and endoplasmic reticulum, as well as aberrant axonal pathfinding (18). However, despite its importance, no neuronal substrates have been reported so far. Only three Ari-1 substrates have been postulated, either in cultured cells or *in vitro* (20–22), while three Parkin substrates were reported to interact with Ari-1 in COS-1 cells (23). For this reason, with the aim to identify neuronal Ari-1 substrates, we took advantage of two methodologies developed by our lab (24). The first one, the ^bio^Ub strategy, allows the identification of hundreds of ubiquitinated proteins from neuronal tissues (25, 26). The system relies on the overexpression of a tagged ubiquitin that bears a 16 amino-acid long biotinylatable peptide (25, 27), which can be biotinylated by the *E.coli* biotin holoenzyme synthetase enzyme (BirA) in neurons *in vivo* (25, 26). Remarkably, this approach can be efficiently applied to identify neuronal E3 ligase substrates (28, 29). The second methodology, in contrast, favours the isolation of GFP-tagged proteins under denaturing conditions to further characterize their ubiquitination pattern under the presence or absence of an E3 ligase (30, 31).

Here we have combined the ^bio^Ub strategy with the overexpression of Ari-1 and identified 16 putative neuronal substrates of Ari-1. Among those, we focused on Comatose (Comt), the fly NSF orthologue (32), due to its relevance in normal and pathological function of the synapse. By the isolation of GFP-tagged Comt from *Drosophila* photoreceptor neurons overexpressing Ari-1, we confirmed Comt/NSF as an Ari-1 ubiquitin substrate, and showed that it is mostly monoubiquitinated. Furthermore, we reported that Ari-1 loss of function mutants displayed lower rate of spontaneous neurotransmitter release, but enhanced evoked release, due to failures at the pre-synaptic side. These defects in the mutants are compatible with a deregulation of Com/NSF activity. Altogether, our data show that Ari-1 regulates neurotransmitter release by controlling Comt/NSF activity through ubiquitination.

## Results

### Generation of flies for the identification of neuronal Ari-1 substrates

We generated ^*bio*^*ari* flies in order to screen, in an unbiased manner, for putative substrates of the E3 ligase Ari-1. These flies overexpress in photoreceptor neurons (using the *GMR-Gal4* driver) both an untagged version of Ari-1 protein (Fig. 1A) and the (^bio^Ub)_6_-BirA construct (Fig. 1B), which is the one that provides the biotinylatable ubiquitins and the BirA enzyme to the system. The rationale for using *GMR-Gal4* is that it has been shown to be a suitable driver for ubiquitin proteomics experiments from mature neurons, in comparison with other neuronal drivers tested (26). ^*bio*^*Ub* flies only overexpressing the (^bio^Ub)_6_-BirA construct were used as control. Overall, expression levels of biotinylated ubiquitin were similar in both genotypes, as indicated by a Western blot performed with biotin antibody using whole head extracts (see *input* fraction of Fig. 1C). Similarly, the amount of ubiquitinated material eluted from biotin pulldowns on both fly lines was also similar (see *elution* fraction of Fig. 1C).

**Figure 1.**
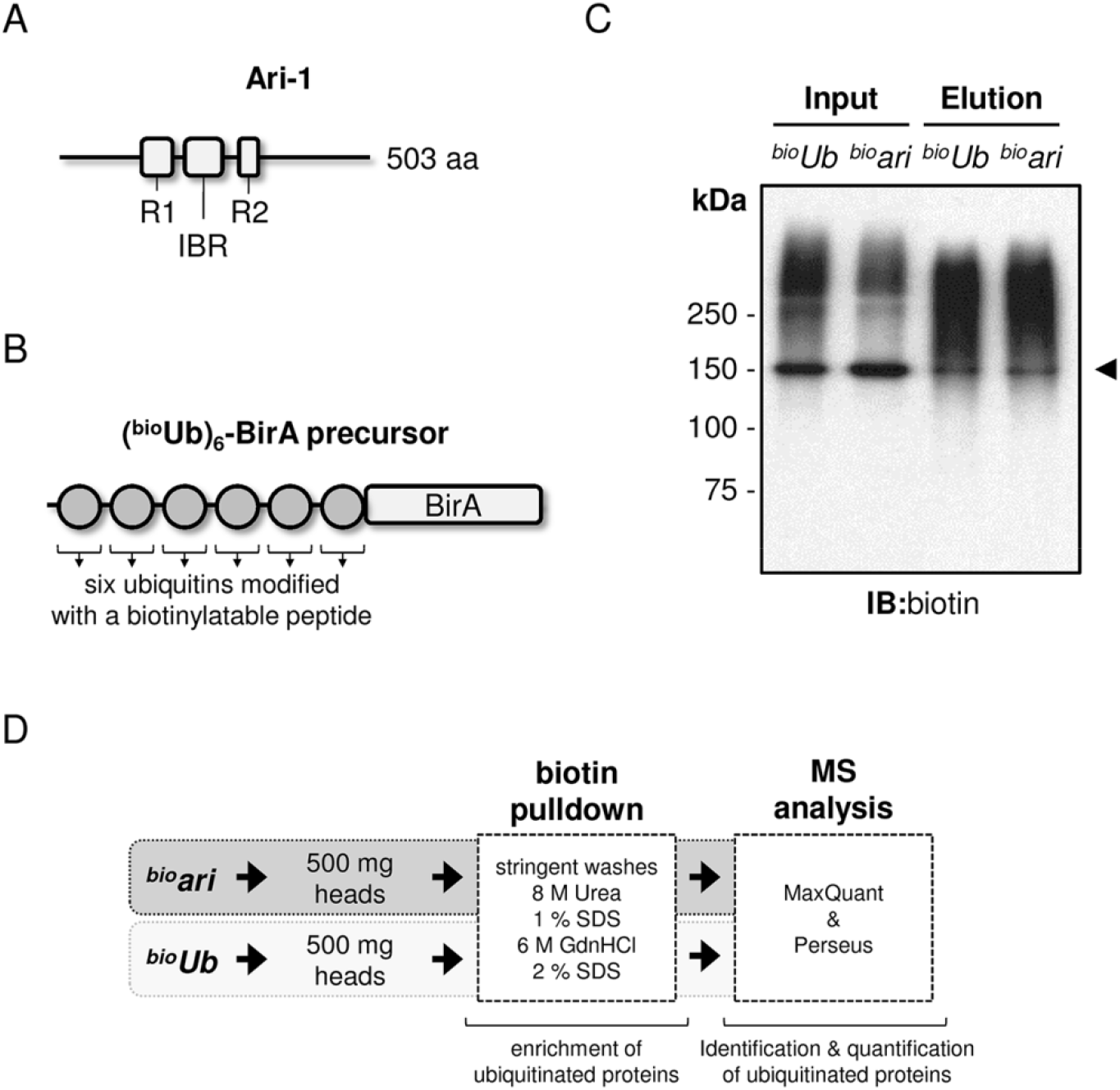
Generation of ^*bio*^*ari* flies. **A)** Schematic representation of the domain structure of *Drosophila* Ari-1 E3 ligase according to Uniprot (ID: Q94981). Ari-1 belongs to the RING between RING E3 ligase family, and as such, it is characterized by the presence of two RING domains (RING1 or R1 from residue 133 to 183 and RING2 or R2 from residue 291 to 322), which are separated by a conserved sequence called the in-between RING domain (IBR, from residue 203 to 264). ^*bio*^*ari* flies overexpress an untagged version of Ari-1 in the photoreceptor cells under the control of the *GMR-Gal4* driver. **B**) Schematic representation of the (^bio^Ub)_6_-BirA precursor (25). It is composed of six copies of ubiquitin, which had been N-terminus modified with a short biotinylatable peptide (MGLNDIFEAQKIEWHEGSGSG). The biotin holoenzyme synthetase (BirA) is found at the C-terminus. This construct is digested by endogenous deubiquitinating enzymes, allowing the biotinylation of each ubiquitin molecule by BirA *in vivo*. ^*bio*^*ari* and ^*bio*^*Ub* flies overexpress this precursor in the photoreceptor cells under the control of the *GMR-Gal4* driver. **C**) Expression of the biotinylated ubiquitin is similar between ^*bio*^*ari* and ^*bio*^*Ub* flies. Anti-biotin Western blot performed on whole lysates (inputs), as well as on the fractions coming from biotin pulldowns, where the ubiquitinated material is enriched (elutions), is shown. The abundant pyruvate carboxylase (~130 kDa), which is biotinylated endogenously, is highlighted with an arrowhead. **D**) Workflow for the identification of *Drosophila* Ari-1 substrates. 500 mg of heads from ^*bio*^*ari* and ^*bio*^*Ub* flies were subjected to biotin pulldowns and mass spectrometry analysis. MS data were analyzed by MaxQuant and Perseus softwares.

Higher levels of Ari-1 enzyme in ^*bio*^*ari* flies should result on an enhanced ubiquitination of Ari-1 substrates relative to ^*bio*^*Ub* flies. Therefore, quantitative proteomic experiments were carried out, following a previously described work-flow (28, 29), to decipher the ubiquitinated proteome from both ^*bio*^*ari* and ^*bio*^*Ub* flies, and hence, detect those proteins whose ubiquitination is regulated by Ari-1. In brief, we collected 500 mg of heads of each fly genotype and subjected triplicate samples to biotin pulldown and LC-MS/MS analysis. This allowed us to isolate, detect and quantify ubiquitinated proteins present in fly photoreceptor neurons expressing physiological (^*bio*^*Ub* samples) and high (^*bio*^*ari* samples) levels of Ari-1 (Fig. 1D). Only those proteins with at least a 2-fold enrichment determined with a p-value < 0.05 in ^*bio*^*ari* samples were considered as putative Ari-1 substrates (Fig. 2A).

**Figure 2.**
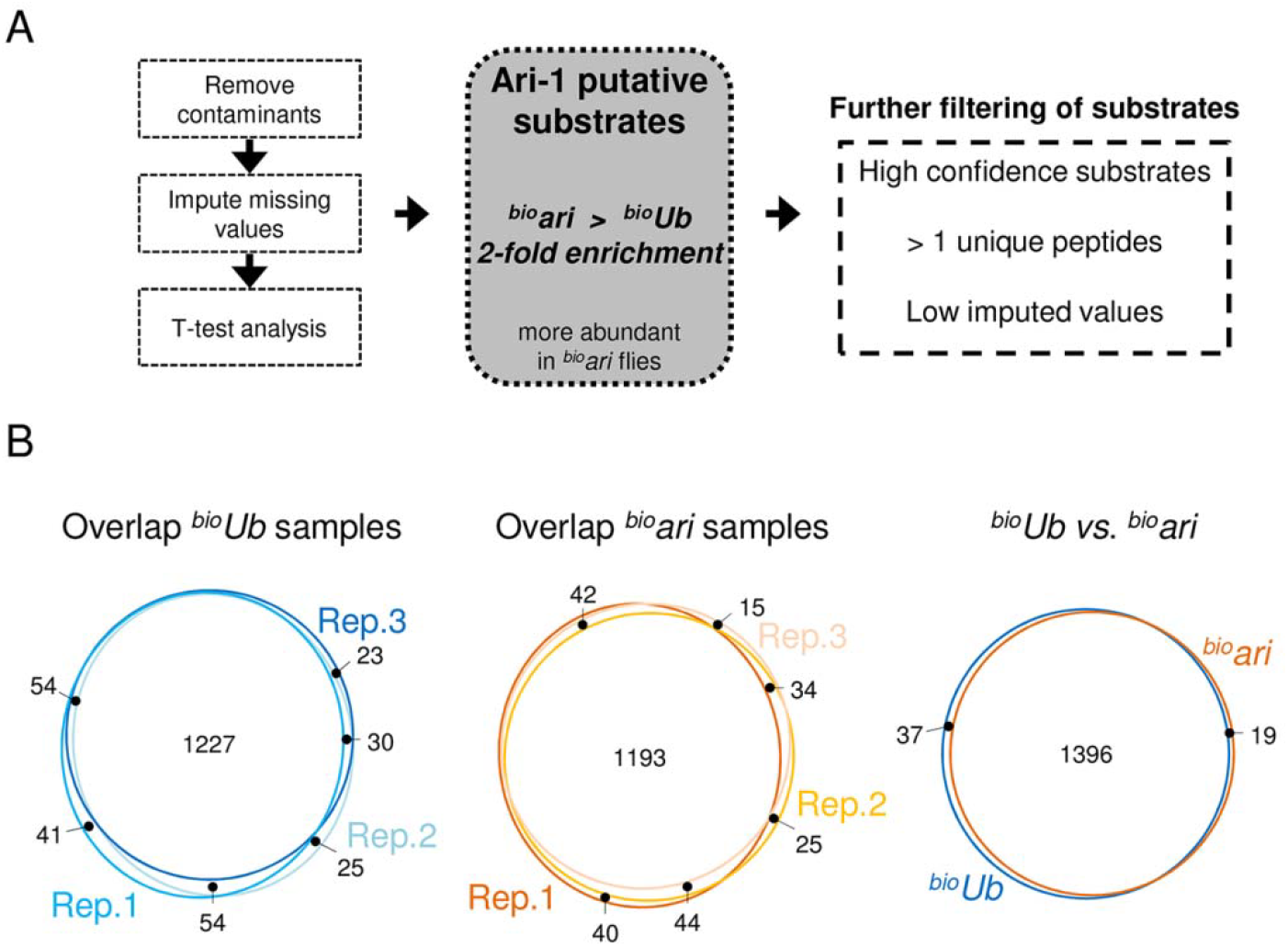
Mass spectrometry data analysis. **A**) Workflow for selection of Ari-1 putative substrates. After removing contaminants, missing LFQ values were imputed and statistical significance determined by two-tailed Student’s *t*-test. Proteins with 2-fold enrichment in ^*bio*^*ari* samples relative to ^*bio*^*Ub* were considered as putative Ari-1 substrates. These proteins were further filtered according to the number of peptides and the imputed values in order to obtain a list of high-confidence putative substrates. **B**) Overlap between the MS-identified proteins from ^*bio*^*ari* and ^*bio*^*Ub* samples. Venn diagrams displaying the overlap between the three ^*bio*^*Ub* and ^*bio*^*ari* replicas, as well as between the two conditions are shown.

### Identification of Ari-1 putative substrates by mass spectrometry

Biotin pulldowns were performed on three biological replicates for each condition. Identified ubiquitinated proteins were highly reproducible across the three replicate samples of each condition, and also between both conditions (Fig. 2B). Protein abundance was determined using label-free quantification (LFQ) intensities (33), which also displayed a high correlation across samples (Pearson correlation ≥ 0.96). For statistical analysis, missing LFQ values were imputed with low random values from a normal distribution, meant to simulate expression levels below the detection limit. It should be noted that proteins with too many imputed LFQ values were discarded from the Ari-1 putative substrate list. Only proteins with LFQ values in at least the three replicas of one of the conditions, either ^*bio*^*ari* or ^*bio*^*Ub*, or in at least two replicas of both conditions, are labelled in Fig. 3.

**Figure 3.**
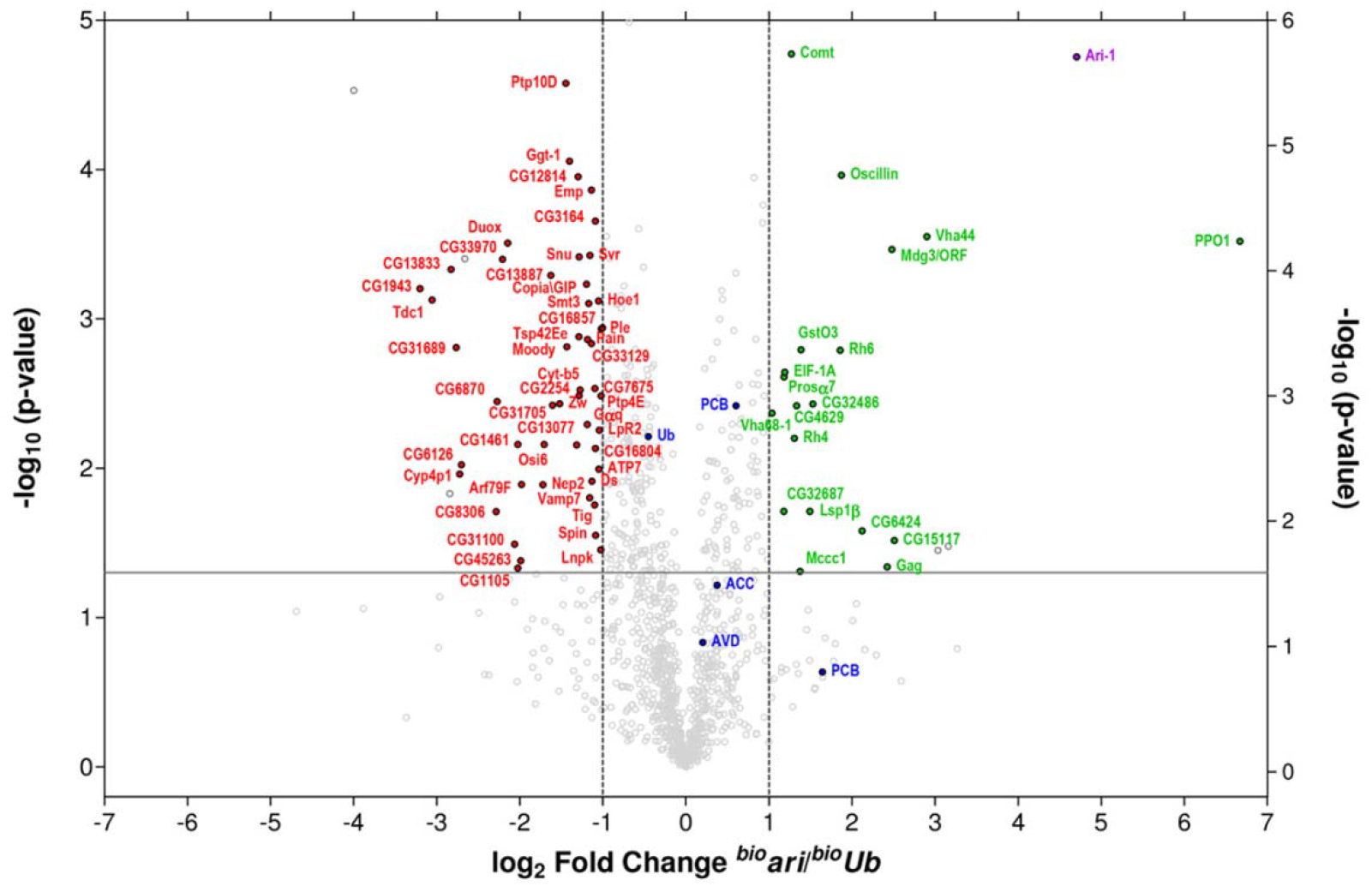
Putative Ari-1 substrates. Volcano plot showing differentially ubiquitinated proteins between ^*bio*^*ari ^bio^Ub* control flies. Abundance of each individual protein was determined by the sum of LFQ intensities of the three replicas performed. LFQ ^*bio*^*ari*/^*bio*^*Ub* ratios in the X-axis (in log_2_ scale) and the *t*-test p-values in the Y-axis (in-log_10_ scale) are displayed. The horizontal grey lane indicates the statistical significance (p-value < 0.05), while vertical grey lanes represent a 2-fold increase (log_2_ LFQ ^*bio*^*ari*/^*bio*^*Ub* = 1), or a 2-fold decrease (log_2_ LFQ ^*bio*^*ari*/^*bio*^*Ub* = −1), of the ubiquitination levels. Endogenously ubiquitinated proteins (ACC and PCB), Avidin (AVD), as well as ubiquitin (Ub) are shown in blue. Proteins whose ubiquitination increase or reduce at least 2-fold upon Ari-1 overexpression are shown in green and red, respectively. Ari-1 is shown in purple.

Ari-1 protein was found significantly enriched (p = 0.0000176) in ^*bio*^*ari* samples relative to ^*bio*^*Ub* control flies (Ari-1, ^*bio*^*ari*/^*bio*^*Ub* LFQ ratio of 26.1). By contrast, and in agreement with Western blotting results (Fig. 1C), ubiquitin levels were similar in both conditions (Ub, ^*bio*^*ari*/^*bio*^*Ub* LFQ ratio of 0.73) (Fig. 3). The relative abundance of the majority of proteins quantified did not change significantly either; this included Avidin (AVD) and proteins known to be endogenously conjugated with biotin (ACC: acetyl-CoA carboxylase, PCB: pyruvate carboxylase) (25, 34), which serve as internal controls to determine the correct processing of the biotin pulldowns (Fig. 3, marked in blue).

Out of the 1452 proteins quantified (see ^*bio*^*Ub vs ^bio^ari* Venn diagram in Fig. 2B, and **Table S1**), 22 proteins were found significantly enriched (p-value < 0.05 or –log_10_ p-value > 1.3) by at least 2-fold (log_2_ fold change > 1) in ^*bio*^*ari* flies relative to ^*bio*^*Ub*, including the Ari-1 E3 ligase itself (Fig. 3, marked in green). Two proteins, Mdg3/ORF and Gag, correspond to products encoded by genes within a transposable element, whereas another two, Calx and MoxGM95, were only identified with one unique peptide. Additionally, Mccc1 had been classified as background in previous biotin pulldown experiments (26), as it uses biotin as cofactor. After excluding these proteins, the final list of putative Ari-1 substrates contained 16 proteins (Table 1). Four of these putative Ari-1 substrates are still uncharacterized *Drosophila* proteins (CG32486, CG32687, CG4629 and CG6424), but the remaining ones are involved in different metabolic processes (CG15117, GstO3, Lsp1β, Oscillin and PPO1), translation (EIFA), phototransduction (Rh4 and Rh6), ATP-hydrolysis coupled transmembrane transport (Vha 44 and Vha 68-1) and protein degradation (Prosα7). The putative Ari-1 substrate identified with the highest significance (p = 0.0000168) was Comt, a protein known to be involved in neurotransmitter release. The human orthologue of Comt, termed NSF (32), is known to regulate the disassembly of the SNARE complexes once neurotransmitter has been released (3).

**Table 1:**
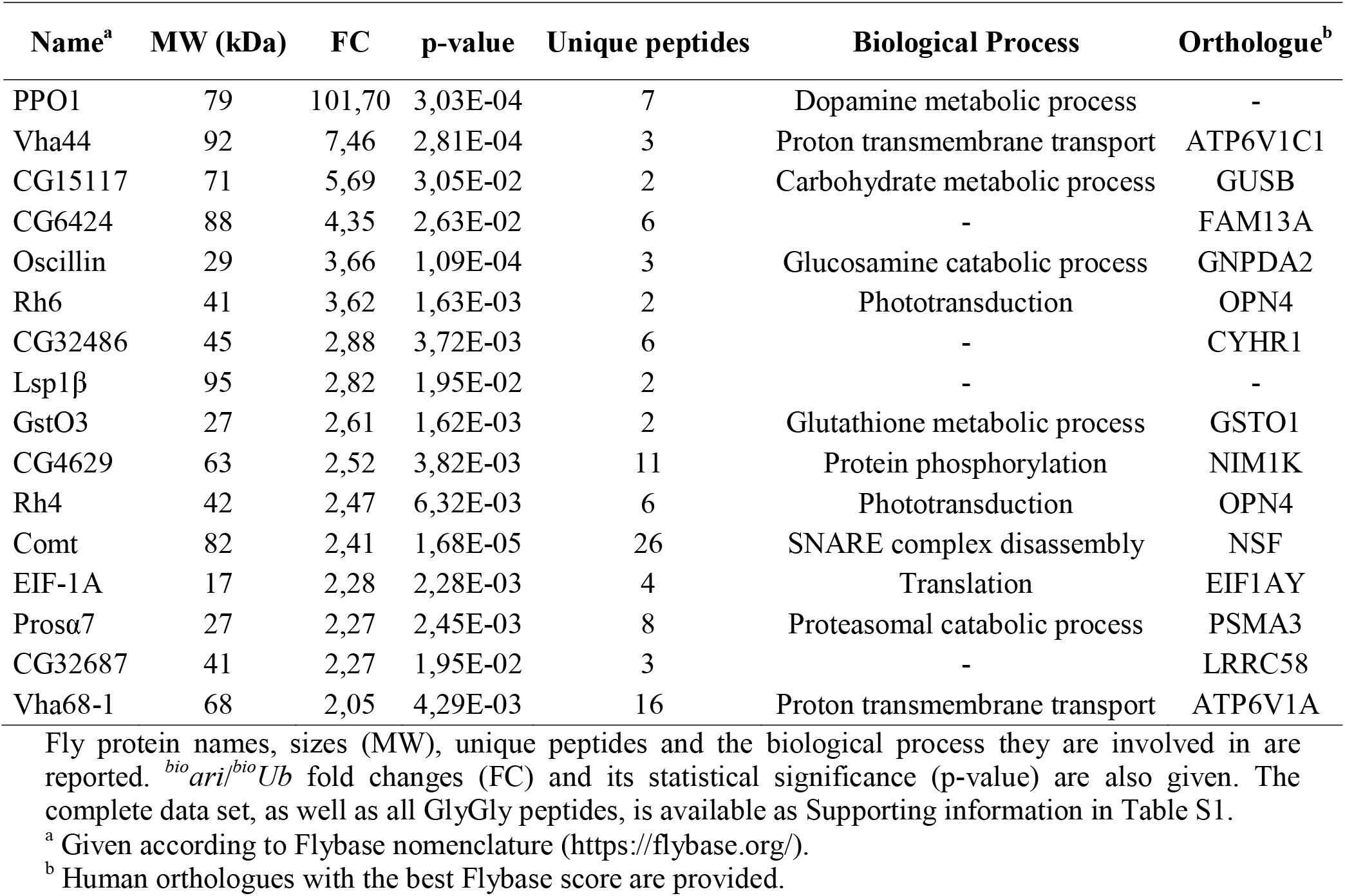
High confidence putative Ari-1 substrates.

The proteomics approach we applied does not particularly enrich for ubiquitination sites, nevertheless we did also identify 150 peptides containing the characteristic GlyGly remnant of ubiquitin. That is, the last two glycines of ubiquitins that are left covalently attached to the substrates after trypsin digestion of the proteins (35). Of those, 64 GlyGly sites had been previously reported (26, 28, 29), but to our knowledge, 86 are novel ubiquitination sites not described until now in *Drosophila* (**Table S1**). Among the sites identified, we did also detect GlyGly remnants on the seven internal lysines of ubiquitin, which indicates that all ubiquitin type chains are present in fly neurons. The abundance of all ubiquitin chain types was comparable in control and Ari-1 overexpressing flies (**Table S1**), suggesting that high levels of Ari-1 do not alter the overall landscape of the ubiquitin linkages in flies, nor directly neither indirectly by affecting other ligases.

### Comt/NSF is ubiquitinated by Ari-1 in Drosophila neurons in vivo

Having identified Comt as a putative substrate of Ari-1 by mass spectrometry analysis, we decided to validate this result biochemically. To that end, we combined flies that overexpressed a GFP-tagged version of Comt with ^*bio*^*Ub* and ^*bio*^*ari* fly lines. The first combination gave rise to flies that overexpress the (^bio^Ub)_6_-BirA construct (Fig. 1B) together with GFP-tagged Comt, termed ^*bio*^*comt* flies (Fig. 4A). The offspring of the second cross (the ^*bio*^*ari/comt* flies) express the Ari-1 E3 ligase (Fig. 1A), in addition to the GFP-tagged Comt and the (^bio^Ub)_6_-BirA construct (Fig. 4A). We then performed GFP-pulldown assays (30) using protein extracts of ^*bio*^*comt* and ^*bio*^*ari/comt* flies, in order to isolate GFP-tagged Comt and compare its ubiquitination levels in the presence of physiological (^*bio*^*comt)* or high levels of Ari-1 (^*bio*^*ari/comt*) *in vivo*. After pulldowns, which were performed in triplicate (**Fig. S1**), ubiquitinated and non-modified fractions of Comt were monitored using anti-biotin and anti-GFP antibodies, respectively. As shown in Fig. 4B and **Fig. S1**, when comparable levels of non-modified Comt were observed in both genotypes, Comt was found highly ubiquitinated in the presence of Ari-1. Altogether, these results confirmed that Comt is an *in vivo* target of Ari-1 E3 ubiquitin ligase in *Drosophila*.

**Figure 4.**
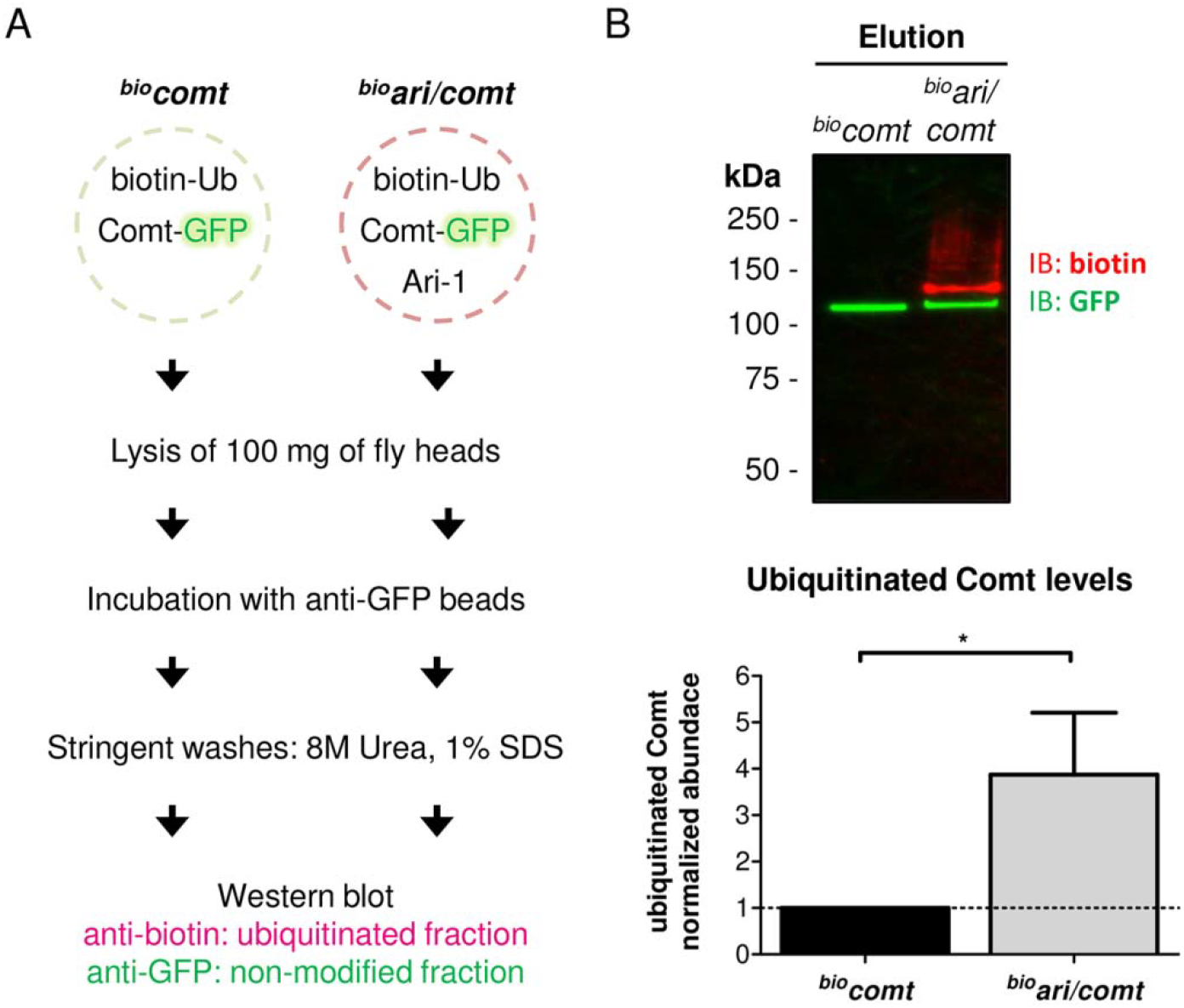
Validation of Comt/NSF as an Ari-1 substrate. **A**) Workflow for the validation of Comt/NSF as Ari-1 substrate. ^*bio*^*comt* flies express the (^bio^Ub)_6_-BirA construct (biotin-Ub; see Fig. 1B) and a GFP-tagged version of Comt (Comt-GFP) under the control of the *GMR-Gal4* driver. ^*bio*^*ari/comt* are ^*bio*^*comt* flies that express, in addition, the Ari-1 E3 ligase. 100 mg of head of each genotype were lysed and incubated with anti-GFP beads in order to isolate GFP-tagged Comt. After stringent washes with 8 M Urea and 1 % SDS, the ubiquitination levels of GFP-tagged Comt in both genotypes were determined by Western blot with anti-biotin antibody. Non-modified GFP-tagged Comt was monitored with anti-GFP antibody. **B**) GFP-Comt is ubiquitinated by Ari-1. Western blot performed on isolated GFP-tagged Comt in the presence of physiological (^*bio*^*comt*) or high levels of Ari-1 (^*bio*^*ari/comt*). Equal levels of GFP-Comt were loaded in order to have comparable ubiquitinated levels. Ubiquitinated fraction (red) was monitored with anti-biotin antibody, while anti-GFP was used to detect the non-modified fraction (green). Below, a quantification of GFP-tagged Comt from three independent pulldowns is shown. Semi-quantification of dual color Western blots was performed with Image Lab software (Bio-Rad) and statistical significance determined by two-tailed Student’s t-test. Ubiquitinated Comt levels in ^*bio*^*ari/comt* flies were normalized to the levels found in control flies (^*bio*^*comt*) for a clearer representation; in ^*bio*^*comt* flies Comt levels are, therefore, shown as 1 (dash horizontal line). * p-value = 0.0139.

Comt/NSF is a well-known component of the SNARE complex. SNARE complex formation bridges the vesicles and plasma membranes, mediating neurotransmitter release at synapses. As currently thought, NSF-SNAPs disassemble the SNARE complex by an ATP hydrolysis dependent process, allowing to recycle and to reuse SNARE components for a following round of vesicle fusion (5, 36). Consequently, we reasoned that Ari-1 might play a role in neurotransmission also.

### Ari-1 mutations alter spontaneous and evoked neurotransmitter release

We first analyzed spontaneous neurotransmitter release under two electrode voltage clamp conditions (TEVC). The frequency of spontaneous miniature events in neuromuscular junctions (NMJs) of *ari* mutant larvae was reduced over 50 % relative to controls (Fig. 5A-B) suggesting a presynaptic modification in mutant terminals. The mean of mini excitatory junctional current (mEJC) frequencies in controls were 2.19 ± 0.26 Hz in normal larvae, 2.46 ± 0.29 Hz in the male *ari^2^* mutant covered by the duplication *Dp (ari^2^; Dp)*, and 2.78 ± 0.26 Hz for the female siblings (*ari^2^/+* ♀). These genotypes constitute different forms of controls. By contrast, mEJCs in male mutant larvae were 1.02 ± 0.12 Hz for *ari^2^*, 1.20 ± 0.38 Hz for *ari^3^* and 0.92 ± 0.12 Hz for *ari^4^*. The reduced mEJC frequency of all *ari* mutant alleles indicates a relatively low rate of spontaneous synaptic vesicle fusion, which could arise from either a decreased number of release sites or a reduced probability of spontaneous vesicle fusion.

**Figure 5.**
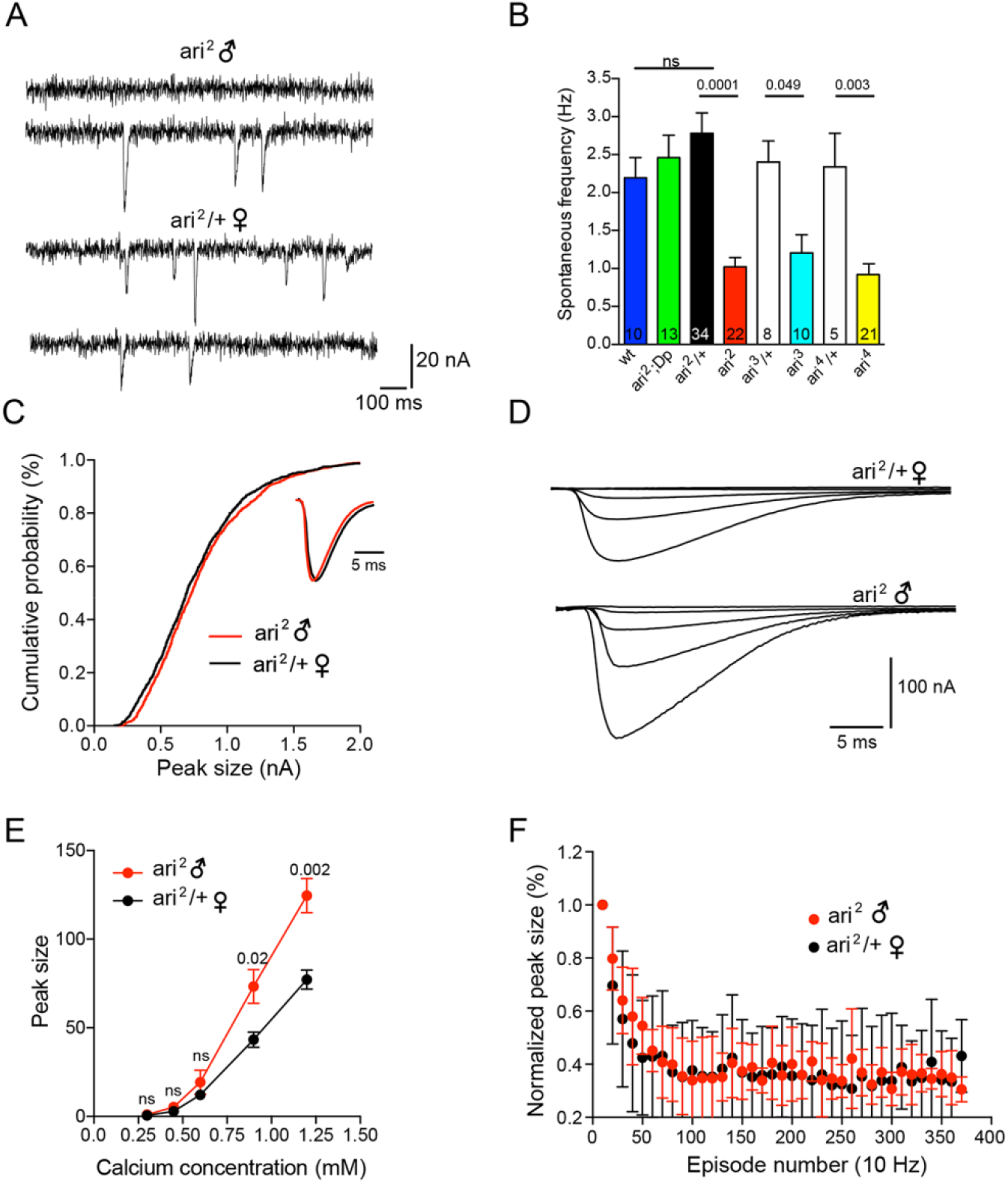
Synaptic release is affected in *ari-1* mutants. **A)** mEJC frequency in *ari-1* mutants is reduced. Representative traces from a spontaneous mEJCs recording. Events were captured under TEVC (Vh = −80 mV) from larval muscle 6 fibers in *ari^2^* mutant male (top), and in heterozygous *ari^2^/+* female siblings (bottom). **B)** Quantification of the effects over spontaneous release. mEJC frequency (mean ± SEM) was calculated from the following larvae strains; (wt), *ari^2^*♂ covered by the duplication *Dp(1;3)JC153* (*ari^2^;Dp*), heterozygous *ari^2^* female siblings (*ari^2^/+* ♀), *ari^2^* ♂ and two additional *ari* alleles (*ari^3^* and *ari^4^*), and their respective heterozygous females. Student’s *t*-test. Note the consistent reduction of mEJC frequency across all mutant genotypes with respect to the sibling female controls. **C)** Neither size nor mEJC time course is affected in *ari^2^* animals. Cumulative probability curve of mEJC amplitudes for *ari^2^/+* ♀(black) and *ari^2^* ♂ (red) larvae (n = 7 larvae). Inset: average representative mEJC traces from *ari^2^*♂ mutant superimposed to that of *ari^2^/+* ♀. Traces were scaled up to the maximum peak value. **D)** Representative traces of evoked transmitter release recorded under TEVC (Vh = −80 mV), at increasing extracellular Ca^2+^ concentrations (from 0.3 to 1.2 mM), recorded from sibling *ari^2^/+* ♀ and *ari^2^* ♂ larvae. **E)** Quantification of Ca^2+^ dependence of evoked release. Maximal EJC response was plotted versus the extracellular calcium concentration for *ari^2^* ♂ compared to *ari^2^*/+ ♀ (n = 6 for all genotypes; One way Anova post hoc: Student’s *t*-test). **F)** Quantification of EJC trains in response to a 10 Hz stimulation. Peak size was normalized to their initial response. Average of 6 different experiments from *ari^2^* ♂ and *ari^2^/+* ♀ larvae.

To further analyze spontaneous release, we compared the amplitude and time course of spontaneous mEJCs in *ari^2^* ♂ versus *ari^2^/+* ♀ larvae (Fig. 5C). The mean *ari^2^* mEJC peak amplitude (0.73 ± 0.007 nA, n = 7 cells) was not significantly different from that of *ari^2^/+* controls (0.75 ± 0.02 nA, n = 7). Cumulative frequency distribution of mEJC amplitudes (Fig. 5C) also revealed not significant differences between *ari^2^* ♂ and *ari^2^/+* ♀ larvae. Further, both genotypes showed a very similar mEJC time course (Fig. 5C, inset). The fact that mEJC amplitude and time course remain the same in *ari^2^* indicates that postsynaptic glutamate receptor properties and receptor density per active site are not altered, and further reinforce the hypothesis that the primary mutant defect is at the presynaptic site.

We also noticed a conspicuous enlargement of evoked release in *ari^2^* mutants. Postsynaptic evoked synaptic currents (EJC) in *ari^2^* males were larger than those of the heterozygous female controls (Fig. 5D). To characterize in detail these effects, EJCs amplitudes were analyzed at various extracellular Ca^2+^ concentrations (from 0.3 to 1.2 mM) by stimulating the fibers at 0.2 Hz. In all Ca^2+^ concentrations tested *ari^2^* ♂ EJC amplitudes were larger that the female controls, although differences reached statistical significance at 1.2 and 0.90 mM only (Fig. 5E). Thus, in the presence of 1.2 mM extracellular Ca^2+^, the mean peak EJC amplitude for *ari^2^* ♂ *w*as 124.61 ± 9.72 nA (n = 6) compared to 77.20 ± 5.33 nA, (n = 6) for *ari^2^*/+ females. Similar differences were found at an extracellular Ca^2+^ concentration of 0.90 mM (*ari^2^* ♂: 73.3 ± 9.5 versus 43.2 ± 4.3 nA in *ari^2^*/+ ♀) while at lower concentrations: 0.6 mM (*ari^2^*♂: 19.2 ± 6.8 versus 12.2 ± 1.6 nA in *ari^2^*/+ ♀), 0.45 mM (*ari^2^* ♂: 5.2 ± 1.7 versus 2.9 ± 0.5 nA in *ari^2^*/+ ♀) and 0.3 mM (*ari^2^* ♂: 1.0 ± 0.2 versus 0.5 ± 0.0 nA in *ari^2^*/+ ♀) differences were not statically significant.

Short-term synaptic depression observed during long high-frequency stimulation is associated with the slower replenishment of the readily-releasable pool (RRP) (37), reflecting the number of vesicles docked and ready to be released. According to this model, rapid stimulation depletes the RRP and the subsequent responses rely on the replenishment of the RRP. In order to examine the possible effects of *ari* over the whole synaptic refilling process, we employed a 10 Hz stimulation protocol to monitor synaptic vesicle replenishment (Fig. 5F). The average results of 6 different experiments did not show any difference between *ari^2^* ♂ and its sibling female controls.

A larger EJC can be explained by a relative large number of synaptic release sites. To discard this possibility, the number of synaptic contacts was analyzed and compared (Fig. 6A-B). We found that the total number of synaptic contacts onto muscle fibers 6/7 (abdominal segment 3) in *ari^2^* ♂ (82.55 ± 4.37, n = 20) was not significantly modified with respect to female controls *ari^2^*/+ ♀ (79.72 ± 3.62, n = 22). Actually, no obvious morphological difference in NMJs between mutant and control larvae could be found.

**Figure 6.**
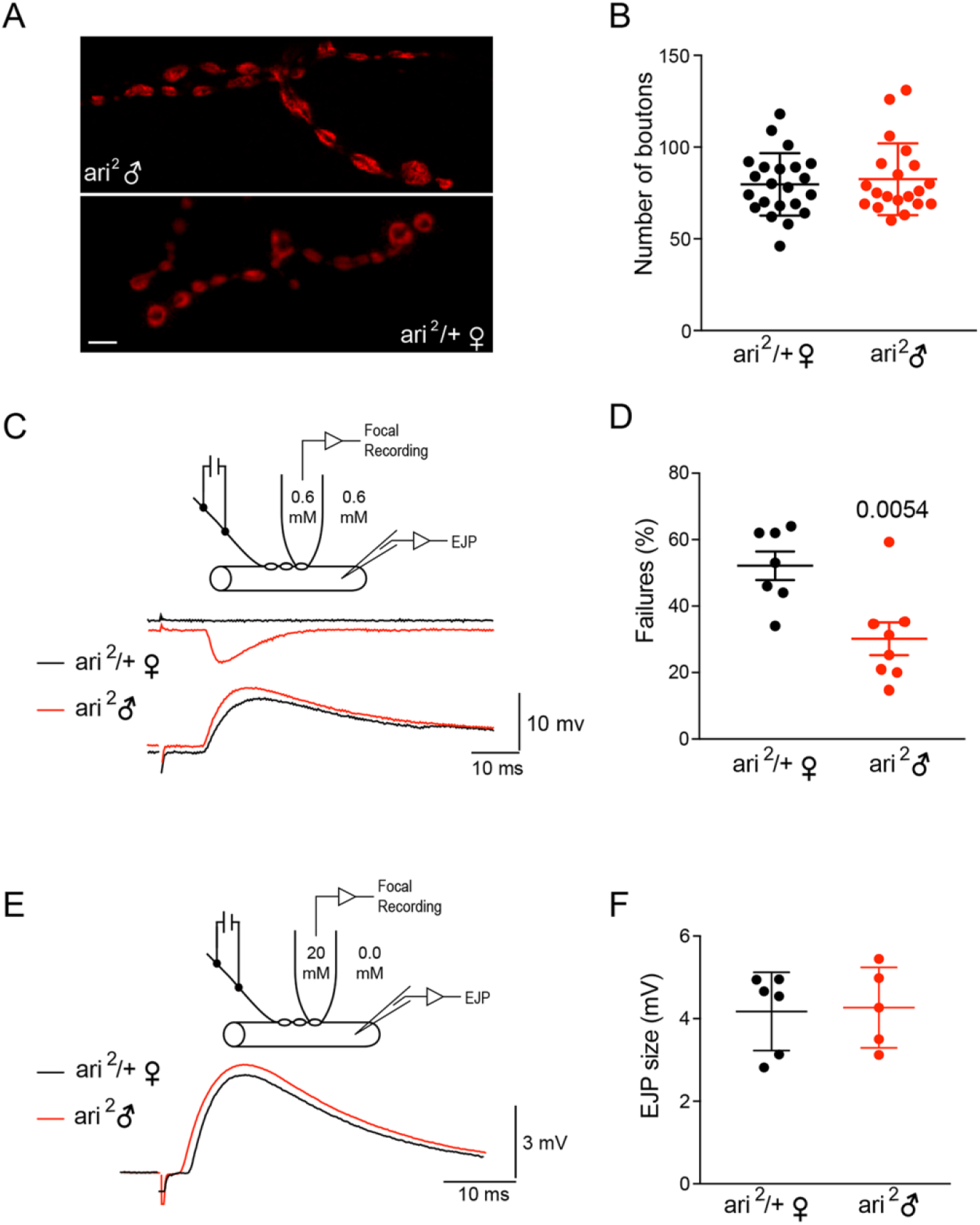
*Ari-1* mutants have low synaptic failure rate. **A)** Synaptic density is not affected by *ari^2^* mutation. Synaptic boutons were identified by inmunostainning against the presynaptic protein CSP. Representative confocal images from *ari^2^* ♂ mutant (top) and control (bottom) neuromuscular junctions (muscle 6). **B)** The average number of synaptic boutons in larval motorneurons from *ari^2^* ♂ compared with the sibling heterozygous *ari^2^/+* ♀ are not statistically different. **C)** *Ari^2^* ♂ mutants have fewer transmitter release failures. Top drawing: scheme of the recording. An extracellular focal recording of the synaptic current (type Is), was recorded by the help of a patch pipette placed over the presynaptic terminal. Calcium was kept at the same extracellular concentration (0.6 mm). The excitatory junction potential (EJP) was recorded from the muscle fiber. Bottom: Representative traces of a simultaneous focal and EJP recording. In red, an *ari^2^* ♂ recording, superimposed to a female *ari^2^/+* ♀ (black traces). Note how a single bouton failure correlates with a normal EJP at the muscle fiber level, indicating that it represents a true failure to recruit the release sites under the focal recording pipette. **D)** The mean average number of failures was smaller in the *ari^2^* ♂ mutants (n=8) when compared with the heterozygous *ari^2^/+*♀ (n=8). Student’s *t*-test. **E)** In *ari^2^* mutants, the release probability is higher in physiological calcium concentrations. Top drawing: Scheme of the recording. Extracellular calcium was kept at 0 mM nominal concentration to inhibit release, while the presynaptic bouton under the focal recording was kept at 20 mM to maximize release probability. Bottom: Representative average evoked EJPs recorded in muscle fiber 6 from *ari^2^* ♂ mutants (red) and control larvae *ari^2^/+* ♀ (black). Each trace is the average of 20 consecutive recordings at 0.1 Hz stimulation. To reduce calcium dilution inside the pipette, we employed a fresh pipette every time and proceeded with the experiment as quickly as possible. **F)** Average EJP amplitudes corresponding to data collected from *ari^2^* ♂ mutants (n = 5) and *ari^2^/+* ♀ controls (n = 6). The average size was not different when release probability was increased.

The fact that evoked synaptic release is increased in *ari*^2^ ♂ while exhibit a similar number of synaptic contacts, would argue for an upregulation of quantal release per synapse. To explore this possibility, we performed a failure test analysis (38), by means of focal recordings of single boutons (type Is) (39) while simultaneously recording the muscular depolarization (EJP: Excitatory Junctional Potential) from the corresponding muscle fiber (Fig. 6 C-D). The Ca^2+^ concentration in the bath and inside the pipette was the same (0.6 mM nominal). The rate of synaptic failures was set to yield about a 50 % failure rate in control animals by adjusting the stimulation potential. Under this condition 52.1 ± 4.2% (n = 8 recording sites from 8 different animals) of stimuli failed to elicit neurotransmitter release in *ari*^2^/+ ♀ animals whereas in *ari*^2^ ♂ mutants, transmitter release failed only in 30.1 ± 4.9% (n = 8) of the cases (Fig. 6D). This difference is statistically significant (p = 0.0054) and indicates that mutant boutons release more neurotransmitter in response to nerve stimulation.

We considered two possible mechanisms to explain this enhancement: (1) mutant boutons contain larger amounts of transmitter release machinery and therefore more release sites, and (2) the probability of vesicle fusion in response to calcium at each release site is increased in the mutant. To distinguish between these two alternatives, we examined transmitter release from single varicosities at very high calcium concentrations. The rationale behind is that, at a calcium concentration that saturates the release machinery, most release sites would be recruited during nerve stimulation, and the amplitude of the synaptic response would mainly be governed by the number of release sites.

For these experiments, the bathing solution and the focal pipette contained 0 and 20 mM calcium, respectively (Fig. 6E). The calcium-containing pipette was placed over single type Is boutons, and the EJP was recorded with an intracellular electrode (Vm in the range of −35 mV to −40 mV, Fig. 6E). Each EJP amplitude data was the average of 20 consecutive responses from the same bouton, elicited at 0.1 Hz intervals, provided that no trend towards synaptic response decrement was observed. The amplitudes of the intracellular recorded EJPs were 4.1 ± 0.3 (n = 6 recording sites from 6 different animals) and 4.2 ± 0.5 (n = 5) in control and mutant animals, respectively (Fig. 6F); the difference not being statistically significant. These results indicate that, under elevated external calcium that maximizes recruitment of release sites, mutant terminals do not release more transmitter than controls. We interpret this result in the sense that the number of release sites per bouton is not modified in the mutant, and favor the hypothesis of a regulatory mechanism that increases transmitter release probability at physiological calcium concentration.

## Discussion

We identified 16 novel putative substrates of Ari-1 in *Drosophila* photoreceptor neurons *in vivo* by means of an unbiased proteomic approach. Remarkably, despite Ari-1 has been recently shown to regulate the positioning of the cell nucleus in muscles via a direct interaction with Parkin (21), as well as to interact with some Parkin substrates (21, 23), there is no overlap between the substrates identified for Ari-1 and those previously identified for Parkin in flies (28). Taken together, the available data suggest that the 16 targets identified here are specifically regulated by Ari-1 in *Drosophila* photoreceptor neurons and that this E3 ligase has a wide functional repertoire.

We focused this study on Comt, an ATPase required for the maintenance of the neurotransmitter release (32). Ubiquitination of proteins involved in vesicle trafficking and neurotransmitter release had been previously reported (17, 25, 26, 40). Similarly, the importance of the ubiquitination machinery for the proper neuronal function has also been demonstrated (14, 16, 41, 42). The alterations produced on synaptic transmission by ubiquitination are typically attributed to an acute control of synaptic protein turnover (17, 43). However, many of these presynaptic proteins have been reported to be mainly mono- or di-ubiquitinated (25, 26), a type of ubiquitin modification that is not usually associated with protein degradation. In line with this, our results showed that Comt/NSF is mainly mono-ubiquitinated by Ari-1, suggesting that Ari-1 could be regulating Comt/NSF activity, rather than its lifespan or expression levels.

Ari-1 mutations result in abnormal synaptic function at the larval stage, a result consistent with a regulatory function of NSF. All mutant alleles examined, exhibit a reduced frequency of spontaneous synaptic release. In addition, *ari-1^2^* mutants exhibit a large calcium-dependent evoked release. Analysis of the mechanism for enhanced evoked release in *ari-1^2^* suggests that the primary defect consists in an increased probability of vesicle fusion in response to calcium entry in the presynaptic side. First, by comparing the amplitude and time course of spontaneously occurring postsynaptic events in mutant and control animals, we ruled out the possibility of a postsynaptic modification. Since no significant differences were found, we concluded that the receptor field size and kinetic properties of postsynaptic receptors are normal in the mutant.

As the mutant functional defects could result from alterations of synaptic development, we quantified the number of synaptic contacts, assuming that most release sites occur within varicosities (44). We did not observe any significant difference between mutant and control. Thus, the data point towards an upregulation of synaptic release from single terminals. The failure analysis from single varicosities represents direct evidence that, at relatively low calcium concentrations, mutant terminals release more quanta than controls in response to an action potential. We further examined whether increased quantal release could be explained on the basis of more release sites being concentrated on mutant terminals. The focal recordings using saturating calcium concentrations argue against this possibility. When mutant terminals are exposed to high calcium, in order to increase the likelihood that all active zones within the bouton will release a quantum, EJP amplitudes in the mutant are indistinguishable from that of controls. These data suggest that the number of release sites in mutant and control terminals is similar and favor the hypothesis that, at physiological calcium concentrations, the probability of vesicle fusion upon calcium entry is increased in the mutant.

We found that *ari-1* mutants have opposite effects on spontaneous and evoked release. Classically, the two modes of vesicular release have been considered to represent a single exocytotic process that functions at different rates depending on the Ca^2+^ concentration. However, recent work challenges this idea and supports the alternative model where spontaneous and evoked response might come from different vesicles pools (45). Several experimental evidences indicate that both forms of release may represent separate fusion pathways (46–49). Employing a state of the art optical imaging in larval NMJ, it has been shown that evoked and spontaneous release can be segregated across active zones. Thus, three types of active zones could be defined: those that only release vesicles in response to a rise of intracellular calcium (evoked release), a second population that only participates in spontaneous release and a third small proportion (around 4 %) that participates in both evoked and spontaneous release. This result advocates for a different molecular and spatial segregation of both modes of release (48).

Differential content or activity of regulatory SNARE binding proteins could discriminate between spontaneous and evoked release. It has been shown that the presence of the Vamp-7 isoform could participate in this differential release. Vamp-7 preferentially labels vesicles unresponsive to stimulation and it colocalizes only partially with the endogenous synaptic vesicle glycoprotein Sv2 and the vesicular glutamate transporter Vglut1, suggesting that this vesicle pool does not subserve evoked transmitter release (47). Recently, it was shown that the double knock-out mouse for Synaptobrevin genes, *syb1* and *syb2*, results in a total block of evoked release, while spontaneous release was increased in both frequency and quantal size without changes in the number of docked vesicles at the active zone (49), confirming the idea that evoked and spontaneous release are differentially regulated. Interestingly, Vamp-7 was found by MS to be less ubiquitinated when Ari-1 is overexpressed (Fig. 3), suggesting that Ari-1 mutants could be favoring evoked release through NSF, and reducing spontaneous release through Vamp7. Thus, Ari-1 could be acting as a repressor and activator of evoked and spontaneous release, respectively. All together, these results evidence a new layer of complexity over the actual fine tuning of synaptic transmission. A physiological regulatory mechanism for both types of release has been recently demonstrated for inhibitory synapses at the trapezoid body, an important brain area in auditory integration. In this nucleus, activation of metabotropic glutamate receptor mGluR1 differently modulates both spontaneous and evoked release in both GABAergic and Glycynergic synapses (50).

At this point, we can only speculate how specifically Ari-1 regulates Comt/NSF activity within the presynaptic terminal. However, a change that enhances NSF functionality, would favor the dissociation of the so called trans-SNARE complex and would build up the number of SNARE complex assembled per vesicle, thus increasing the efficiency of fusion machinery in a calcium dependent way. In this sense, it has been proved that fast release of a synaptic vesicle require at least three SNARE complexes, whereas slower release may occur with fewer complexes (51).

Interestingly, some of the additional putative substrates identified are also related to synapse physiology and neurotransmitter release. PPO1 is an enzyme with L-DOPA monooxygenase activity, hence, may be involved in the metabolism of dopamine neurotransmitter (52). Similarly, GstO3 is involved in glutathione metabolism (53), another type of neurotransmitter (54). Vha44 and Vha68-1 are components of the vacuolar proton-pump ATPase (55), whose mutations have been reported to impair neurotransmitter release (56). Vha44 has also been described as an enhancer of Tau-induced neurotoxicity (57), and CG15117, orthologue of human GUSB, has been associated with neuropathological abnormalities (58).

The data reported here may be relevant in the context of Parkinson’s disease. It should be noted that most Parkinson’s related genes encode proteins involved in vesicle recycling and neurotransmitter release at the synapse (59). Thus, the kinase LRRK2 phosphorylates NSF to enhance its ATPase activity upon the SNARE complex and facilitate its disassembly (7–9). Pathological mutations in this protein, such as G2019S, cause an excess of kinase activity (10) that interferes with vesicle recycling (11). Deregulated synaptic aggregates of α-Synuclein may target VAMP-2 hampering the formation of the SNARE complex (13, 60). Parkin is a structural relative of Ari-1, based on their common cysteine rich C_3_HC_4_ motif (19), that is also at the origin of some forms of Parkinson’s disease (61). All these genes and their corresponding mechanisms of activity sustain the scenario in which several types of Parkinson’s disease seem to result from a defective activity of the synapse. In this context, the role of Ari-1 emerges as a mechanism to regulate a key component of the SNARE complex, NSF. Conceivably, Ari-1 may become a suitable target for, either diagnosis or pharmacological treatment of Parkinson’s and related diseases.

### Experimental procedures

#### Fly strains and genetic procedures

The four *ari-1^1-4^* alleles have been previously described and sequenced (18) and the protein sequence can be found in the EMBL data bank under accession number #Q94981. The insertional duplication *Dp(1;3)JC153* (*Dp* for brevity) is routinely used to cover the lethality of *ari-1* and to demonstrate that the mutant phenotype is due to the gene *ari-1*, rather than to a putative second site mutation along the chromosome. *UAS-ari-1* flies, as well as flies overexpressing the (^bio^Ub)_6_-BirA construct in the *Drosophila* photoreceptor neurons under the control of the GMR-GAL4 driver (*GMR-Gal4, UAS-(^bio^Ub)_6_-BirA/CyO; TM2/TM6*) had been previously described (22, 26). *UAS-comt-GFP* flies, used for the overexpression of the *Drosophila* NSF1 protein, were obtained from Dr. R. Ordway (Penn State Univ. USA).

*UAS-ari-1* flies were mated to *GMR-Gal4, UAS-(^bio^Ub)_6_-BirA/CyO; TM2/TM6* (abbreviated throughout the text as ^*bio*^*Ub*) to generate *GMR-Gal4, UAS-(^bio^Ub)_6_-BirA/CyO; UAS-ari-1/TM6* flies (abbreviated throughout the text as ^*bio*^*ari*). ^*bio*^*Ub* and ^*bio*^*ari* flies were then mated to *UAS-comt-GFP* flies in order to generate *GMR-Gal4, UAS-(^bio^Ub)_6_-BirA/CyO; UAS-comt-GFP/TM6* (abbreviated throughout the text as ^*bio*^*comt*) and *GMR-Gal4, UAS-(^bio^Ub)_6_-BirA/CyO; UAS-ari-1/UAS-Comt-GFP* flies (abbreviated throughout the text as ^*bio*^*ari/comt*), respectively. Mixed-sex flies of 2-5 days old of each genotype (^*bio*^*Ub*, ^*bio*^*ari*, ^*bio*^*comt* and ^*bio*^*ari/comt*) were frozen in liquid nitrogen and their heads were collected using a pair of sieves with a nominal cut-off of 710 and 425 µm as previously described (26, 29). Fly heads were then stored at −80 ºC until required. All fly lines were grown at 25 ºC in 12 h light-dark cycles in standard *Drosophila* medium.

#### Biotin pulldown assay

Biotin pulldowns (25) from *Drosophila* heads were performed as previously described (26). In brief, about 500 mg of heads of ^*bio*^*Ub* and ^*bio*^*ari* flies were homogenized in 2.9 ml of *lysis buffer*. Lysates were centrifuged at 16000*g* at 4 ºC for 5 min and the supernatant applied to a *binding buffer*-equilibrated PD10 desalting column (GE Healthcare). Eluates, except 50 µl kept as input fraction, were incubated with 250 µl of NeutrAvidin agarose resin (Thermo Scientific) for 40 min at room temperature and further 2 h and 20 min at 4 ºC with gentle rolling. The material bound to the resin was washed twice with *washing buffer* (*WB*) 1, thrice with *WB*2, once with *WB*3, thrice with *WB*4, once again with *WB*1, once with *WB*5 and thrice with *WB*6. The material that was still bound to the resin (i.e. the ubiquitinated material) was eluted by heating the resin at 95 ºC for 5 min in 125 µl of *elution buffer*. Finally, samples were centrifuged at 16000*g* at room temperature for 2 min in a Vivaclear Mini 0.8 µm PES-micro-centrifuge filter unit (Sartorious) to discard the NeutrAvidin resin.

Buffer compositions for the biotin pulldown assays were as follows: *lysis buffer*, 8 M urea, 1 % SDS, 1x PBS, 50 mM N-ethylmaleimide (Sigma) and a protease inhibitor cocktail (Roche); *biding buffer*, 3 M urea, 1 M NaCl, 0.25 % SDS, 1x PBS and 50 mM N-ethylmaleimide; *WB1*, 8 M urea, 0.25 SDS, 1x PBS; *WB*2, 6 M guanidine-HCl, 1x PBS; *WB*3, 6.4 M urea, 1 M NaCl, 0.2 % SDS, 1x PBS; *WB*4, 4 M urea, 1 M NaCl, 10 % isopropanol, 10 % ethanol, 0.2 % SDS, 1x PBS; *WB*5, 8 M urea, 1 % SDS, 1x PBS; *WB*6, 2 % SDS, 1x PBS; *elution buffer*, 250 mM Tris-HCl, pH 7.5, 40 % glycerol, 4 % SDS, 0.2 % bromophenol blue and 100 mM DTT.

#### GFP pulldown assay

GFP pulldowns were performed as previously described (30) with slight modifications to adapt the protocol to *Drosophila* heads. Briefly, about 100 mg of mixed-sex fly heads of ^*bio*^*comt* and ^*bio*^*ari/comt* flies were homogenized in 400 µl of *lysis buffer*. Lysates were centrifuged once at 16000*g* at room temperature for 1 min, in order to get rid of most of the *Drosophila* head fragments, and then centrifuged once again at 16000*g* at 4 ºC for 10 min. 40 µl of the supernatants were kept as input, while 300 µl were diluted with 450 µl of *dilution buffer* to reduce their Triton concentration to 0.2 % for a better binding. Diluted samples (750 µl) were then incubated with 40 µl of *dilution buffer*-washed GFP-Trap-A agarose beads suspension (Chromotek GmbH) for 2 h and 30 min at room temperature with gentle rolling. GFP beads were subsequently washed once with *dilution buffer*, thrice with *washing buffer* and once with *SDS buffer*. Bound GFP-tagged Comt was eluted in 50 µl of *elution buffer* by heating at 95 ºC for 10 min. Samples were centrifuged at 16000*g* for 2 minutes and supernatant recovered, in order to get rid of the GFP-Trap-A agarose beads.

Buffer compositions for the GFP pulldown assays were as follows: *lysis buffer*, 50 mM Tris-HCl, pH 7.5, 150 mM NaCl, 1 mM EDTA, 0.5 % triton, 50 mM N-ethylmaleimide (Sigma) and 1x protease inhibitor cocktail (Roche); *dilution buffer*, 10 mM Tris-HCl, pH 7.5, 150 mM NaCl, 1 mM EDTA, 50 mM N-ethylmaleimide and 1x protease inhibitor cocktail; *washing buffer*, 8 M urea, 1% SDS, 1x PBS; *elution buffer*, 250 mM Tris-HCl, pH 7.5, 40 % glycerol, 4 % SDS, 0.2 % bromophenol blue and 100 mM DTT.

#### In-gel trypsin digestion and peptide extraction

Eluates from biotin pulldown assays were concentrated in Vivaspin 500 centrifugal filter units (Sartorius) and resolved by SDS-PAGE in 4-12 % Bolt Bis-Tris precast gels (Invitrogen). Proteins were visualized with GelCode blue stain reagent (Invitrogen) and avidin monomers (~15 kDa) and dimers (~30 kDa), still present in the samples, as well as an endogenously biotinylated protein found at ~130 kDa were excluded. For that, the gels were cut into the following slices: slice *A1* from ~15 kDa to ~25 kDa; slice *A2* from ~30 kDa to 50 kDa; slice *B* from 50 kDa to ~130 kDa and slice *C* from ~140 kDa to up to the gel. These 4 slices were then subjected to in-gel trypsin digestion as described previously (62). In brief, proteins were first reduced with DTT and then, alkylated with chloroacetamide. Afterwards, gel pieces were saturated with trypsin and incubated overnight at 37 ºC. The resulting peptides were extracted from the gel, dried down in a vacuum centrifuge and stored at −20 ºC. Peptide mixtures were resuspended in 0.1 % formic acid for LC-MS/MS analysis.

#### LC-MS/MS analysis

An EASY-nLC 1000 liquid chromatography system interfaced with a Q Exactive mass spectrometer (Thermo Scientific) via a nanospray flex ion source was employed for the mass spectrometric analyses. Peptides were loaded onto an Acclaim PepMap100 pre-column (75 µm *x* 2 cm, Thermo Scientific) connected to an Acclaim PepMap RSLC (50 µm *x* 15 cm, Thermo Scientific) analytical column. A 2 to 40 % acetonitrile in 0.1 % formic acid linear gradient, at a flow rate of 300 nl min^−1^ over 45 min, was used to elute peptides from the columns. The mass spectrometer was operated in positive ion mode. Full MS scans were acquired from *m/z* 300 to 1850 with a resolution of 70 000 at *m/z* 200. The 10 most intense ions were fragmented by higher energy C-trap dissociation with normalized collision energy of 28. MS/MS spectra were recorded with a resolution of 17 500 at *m/z* 200. The maximum ion injection time was 120 ms for both survey and MS/MS scans, whereas AGC target values of 3 *x* 10^6^ and 5 *x* 10^5^ were used for survey and MS/MS scans, respectively. Dynamic exclusion was applied for 45 s to avoid repeated sequencing of peptides. Singly charged ions or ions with unassigned charge state were also excluded from MS/MS. Data were acquired using Xcalibur software (Thermo Scientific).

#### Data processing and bioinformatics analysis

Acquired raw data files were processed with the MaxQuant (63) software (version 1.5.3.17) using the internal search engine Andromeda (64) and tested against the UniProt database filtered for *Drosophila melanogaster* entries (release 2015_11; 43712 entries). Spectra originated from the different slices corresponding to the same biological sample were combined. Carbamidomethylation (C) was set as a fixed modification, whereas methionine oxidation, protein N-terminal acetylation and lysine GlyGly (not C-term) were defined as variable modifications. Mass tolerance was set to 8 and 20 ppm at the MS and MS/MS level, respectively. Enzyme specificity was set to trypsin, allowing for a maximum of three missed cleavages. Match between runs option was enabled with 1.5 min match time window and 20 min alignment window to match identification across samples. The minimum peptide length was set to seven amino acids. The false discovery rate for peptides and proteins was set to 1%. Normalized spectral protein label-free quantification (LFQ) intensities were calculated using the MaxLFQ algorithm (33).

#### Western blot analysis

4-12% gradient Bolt Bis-Tris Plus pre-cast gels (Invitrogen) were used for SDS-PAGE. Proteins were transferred to PVDF membranes using the iBlot system (Invitrogen). Following blocking, primary and secondary antibody incubation, membranes were developed either by chemiluminescence, using the Clarity Western ECL Substrate kit (Bio-Rad), or by near-infrared fluorescence. In both cases membranes were developed using the ChemiDoc^TM^ MP Imaging system (Bio-Rad). Biotinylated proteins and, thus, ubiquitinated proteins were detected with goat anti-biotin-horseradish peroxidase (HRP)-conjugated antibody (Cell Signaling Technology, Cat# 7075) at 1:1000. A mouse monoclonal anti-GFP antibody (Roche, Cat# 11814460001) at 1:1000 was used to detect Comt-GFP. A mouse monoclonal anti-β-Tubulin (Developmental Studies Hybridoma Bank, Cat# E7-c) at 1:1000 was used for loading control. Goat anti-mouse IRDye-800CW (LI-COR Biosciences, Cat# 926-32210) at 1:4000 was used as secondary antibody.

#### Larval neuromuscular preparation

Mature third instar larvae at the wandering stage were selected for electrophysiological recordings. As previously described (65), they were pinned down onto a Sylgard-coated experimental chamber and cut open along the dorsal midline in an extracellular solution containing 100 mM NaCl, 5 mM KCl, 20 mM MgCl_2_, 5 mM HEPES and 115 mM Sucrose (pH 7.3). After pinning the cuticle flat, all internal organs were removed to expose the body-wall muscle layer, leaving only the central nervous system connected to the muscles through the segmental nerves. These were severed near the CNS and their cut-open ends sucked into a fired-polished glass suction pipette filled with extracellular saline solution. The preparations were then transferred to the microscope for electrophysiological recordings, and viewed with a 40x water immersion objective under Nomarski optics. All experiments were performed at room temperature (21-23ºC) on ventrolateral muscle fibers from segments A2-A5.

#### Electrophysiology

Spontaneous (mEJCs) and nerve-evoked (EJCs) junctional currents were recorded from muscle fibers 6 and 12, using an Axoclamp-2A amplifier in two-electrode voltage-clamp (TEVC) mode. Short-shank pipettes (8-10 MΩ resistance when filled with 1 M KCl solution) were pulled on a Flaming-Brown puller (Sutter Instruments) from thin-wall borosilicate glass (World Precision Instruments). Upon establishing TEVC, the feedback gain control of the amplifier was adjusted, while monitoring the current response to repetitive hyperpolarizing voltage commands, as to obtain a minimal clamp settling time (~ 2 ms) without introducing oscillations or excess noise in the current trace. The holding current, always less than −5 nA for a holding potential of −80 mV, was continuously monitored throughout the experiment. Current signals were low-pass filtered at 500-1000 Hz, digitized and stored in a computer for further off-line analysis using a computer program (SNAP) written in our lab. Current signals were recorded in extracellular solution to which calcium had been added as a chloride salt to attain the desired final concentration (0 - 1.2 mM) as indicated in the text. The number of mEJCs acquired in free-run mode over 3-minute periods was counted in order to determine the average frequency of spontaneous release. To compare the mEJCs amplitude distribution between genotypes, data recorded from different animals (two hundred consecutive events from each animal) of the same genotype were pooled into single data sets.

Evoked EJCs were elicited in voltage-clamped muscle fibers by delivering single pulses (100 - 200 μs) from a stimulator (Cibertec Stimulator CS 20). The stimulus intensity was adjusted slightly over the level required to recruit both motor axons innervating ventrolateral muscle fibers to produce a compound EJC. A mean EJC response for each muscle fiber sampled was obtained by averaging 25 to 50 consecutive responses elicited at 0.2 Hz. Extracellular focal synaptic signals from single boutons were recorded by placing extracellular pipettes (fired polished, 2-5 μm inner diameter) over type Is boutons, easily identified under Nomarski optics, and applying very gentle suction, which was released prior to the beginning of the experiment. Only those boutons showing no visible signs of damage after the experiment were considered in the analysis. In those experiments designed to assess single-terminal failure rates, whole-muscle EJPs were simultaneously recorded with intracellular electrodes in current-clamp mode, as to detect any possible stimulation failure susceptible of being wrongly classed as a release failure. Pooled data in the text are presented as mean ± SEM. The Student’s *t*-test with a level of significance p < 0.05 was routinely used to compare mean values between different genotypes, except when comparing mEJC amplitude distributions, where a Kolmogorov-Smirnov test was used.

#### Immunohistochemical procedures

Third instar larvae were dissected open as described and then fixed for 30 minutes in freshly made 4% paraformaldehyde in Ca^2+^-free PBS containing 5 mM EGTA. After a series of washes with PBS, the larvae were incubated in PBT (0.1 % Triton in PBS) with 2 % BSA and 5 % goat serum, added for 2 h at room temperature. Specimens were next incubated in mouse monoclonal anti-CSP antibody (mAb49) at 1:100 (66, 67), overnight at 4 ºC in PBT, thoroughly washed in PBT and then incubated in secondary anti-mouse antibody (1:1000) conjugated to Cy2 (Jackson Immuno Research, cat# 115-225-146), for 1 hour at room temperature. Preparations were mounted on PBS:glycerol (1:1) and viewed under confocal microscopy.

#### Experimental design and statistical analyses

##### Biotin pulldowns

experiments were performed in triplicate, where 500 mg of mixed-sex fly heads of each genotype (^*bio*^*Ub* and ^*bio*^*ari*) for each replica were used. The amount of heads was set to 500 mg to maximize the amount of ubiquitinated material that is purified and to minimize the amount of reagents that are used. This amount of heads ensures the isolation of enough ubiquitinated material for MS analyses, but do not saturate the PD10 columns, so only one per sample is used. Additionally, the amount of NeutrAvidin beads used can be reduced significantly. After MS analysis, MaxQuant output data from ^*bio*^*Ub* and ^*bio*^*ari* samples were analyzed with the Perseus module (version 1.5.6.0) (68) in order to determine the proteins and GlyGly peptides significantly enriched in each of the genotypes. First, contaminants, reverse hits, as well as proteins and GlyGly peptides with no intensity were removed. Later, during the analysis of proteins, those only identified by site and/or with no unique peptides were also removed. LFQ intensities were used to determine the abundance of proteins, while raw intensities, normalized by median subtraction, were used for GlyGly peptides. Missing intensity values were replaced with values from a normal distribution (width 0.3 and down shift 1.8), meant to simulate expression below the detection limit (68). Statistically significant changes in protein and GlyGly peptide abundance were assessed by two-tailed Student’s *t*-test. Proteins and GlyGly peptides displaying a fold change above 2 with a p-value below 0.05 were selected for further analysis. The selected proteins and peptides were further filtered based on the number of unique peptides and/or on the number of imputed values. For further analyses, we only considered the statistical significance of those proteins that (i) had no imputed values in any of the three replicas of at least one of the conditions, or (ii) had a maximum of one imputed value in each of the conditions.

##### GFP pulldowns

experiments were performed in triplicate, where 100 mg of mixed-sex fly heads of each genotype (^*bio*^*comt* and ^*bio*^*ari/comt*) for each replica were used. The amount of heads was determined as the minimum amount needed to saturate 40 µl of GFP-Trap_A beads. Semi-quantification of dual color Western blots was performed with Image Lab software (Bio-Rad). Ubiquitination levels of GFP-tagged Comt were normalized to the non-modified fraction. Statistical significance of the ubiquitination levels was then determined by two-tailed Student’s *t*-test using GraphPad software.

##### Electrophysiology Statistical analysis

All statistical analyses were done by the GraphPad Prism version 7.0 (GraphPad Software). Data are presented as mean ± SEM unless otherwise noted. Two-tailed Student’s *t*-test was used to assess differences between control and other groups. One way ANOVA was used for analysis of data from three or more groups followed by two-tailed Student’s *t*-test, coefficients of significance are included in the figures.

## Acknowledgements

This research was funded by grants BFU2015-65685 and PGC2018-094630-B-100 from the Spanish Ministry of Economy to A.F., and grant SAF2016-76898-P from the Spanish Ministry of Economy cofinanced with FEDER funds to U.M. JR was supported with a postdoctoral research fellowship from the University of the Basque Country (UPV/EHU). We thank the Bloomington Drosophila Stock Center (NIH P40OD018537), the Vienna Drosophila Resource Center and Prof. Richard Ordway for fly strains. The technical help of Ms. Esther Seco is most appreciated. The authors are also grateful of the technical support provided by Kerman Aloria from the UPV/EHU SGIker, which is supported with European funding (ERDF and ESF).

## Author contributions

## FOOTNOTES

Ari-1: Ariadne-1
Comt: Comatose
EJP: excitatory junctional potential
LFQ: label-free quantification
LRRK2: leucine-rich repeat serine/threonine-protein kinase 2
mEJC: mini excitatory junctional current
NMJ: neuromuscular junctions
NSF: N-ethylmaleimide sensitive factor
RRP: readily-releasable pool
SNAP: Soluble NSF attachment proteins
SNARE: SNAP receptor
v-SNARE: vesicle membrane SNARE protein
t-SNARE: plasma membrane SNARE protein
TEVC: two electrode voltage clamp conditions
VAMP: vesicle-associated membrane protein

## Supporting information

**Figure S1. Ari-1 dependent ubiquitination of GFP-Comt.** A complete data set of Comt ubiquitination by Ari-1 in fly photoreceptors. The ubiquitinated fractions were detected with anti-biotin antibody, while anti-GFP was used for the non-modified fraction. Anti-Tubulin was used as loading control in whole cell lysates (input). The original Westerns done with the pulldown samples are shown. Despite Tubulin levels were similar between genotypes, total GFP levels, as well as ubiquitin levels, were higher in ^*bio*^*Comt* flies than in ^*bio*^*ari*/^*bio*^*Comt* flies. This could be due to a GAL4-dose effect, as ^*bio*^*Comt* flies contain two UAS sequences and ^*bio*^*ari*/^*bio*^*Comt* three. In order to compare the ubiquitinated fraction of isolated Comt-GFP, therefore, comparable levels of non-modified GFP-Comt were loaded (Normalized WB).

**Table S1. Mass spectrometry analysis.** Proteins found more or less ubiquitinated in ^*bio*^*ari* flies, compared to ^*bio*^*Ub* are shown in green and red, respectively (Protein Groups sheet). All GlyGly peptides identified by Mass Spectrometry (MS) are shown in the second datasheet (GlyGly peptides). Sites already reported in previous MS analysis are also indicated. Gene symbol, CG number and protein description is provided according to Flybase.

