## Supplemental data for "The ubiquitin ligase Ariadne-1 regulates NSF for neurotransmitter release"

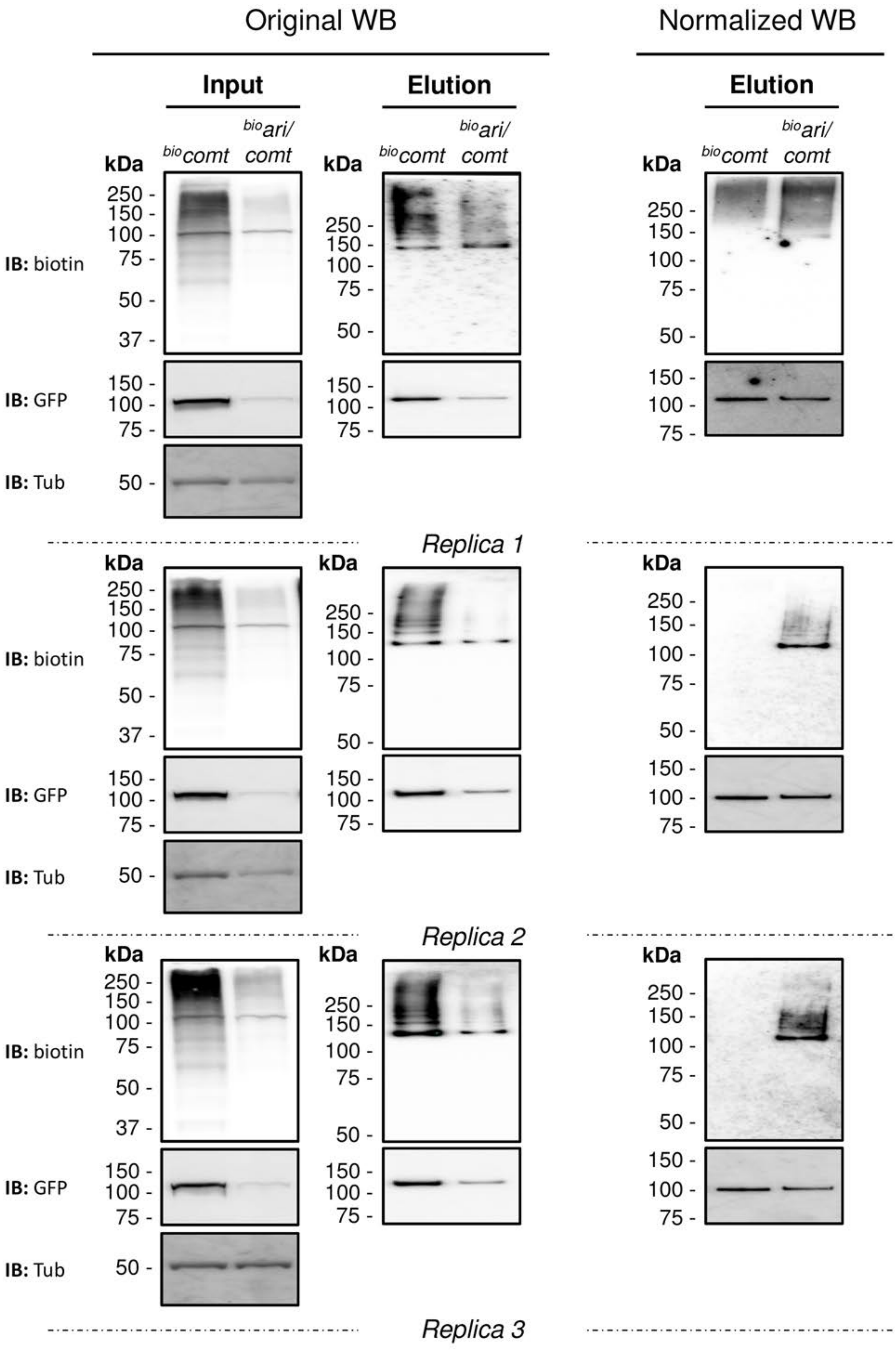

Original WB
Normalized WB
Input
Elution
Elution
bio comt
bio ari/ comt
kDa
250 -
150 -
100 -
75 -
50 -
37 -
IB: biotin
IB: GFP
IB: Tub
Replica 1
Replica 2
Replica 3

LFQ intensity ($log_2$ scale)

| $^{bio}$*Ub* -1- | $^{bio}$*Ub* -2- | $^{bio}$*Ub* -3- | $^{bio}$*ari* -1- | $^{bio}$*ari* -2- | $^{bio}$*ari* -3- | Peptides | Unique peptides |
|---|---|---|---|---|---|---|---|
| 30,52206 | 30,53213 | 30,62758 | 31,75087 | 31,86751 | 31,87291 | 39 | 26 |
| 31,63897 | 31,70141 | 31,40451 | 31,99226 | 33,5032 | 33,37205 | 25 | 25 |
| 28,1684 | 27,81885 | 28,29575 | 32,68781 | 33,06522 | 32,64528 | 16 | 16 |
| 29,54221 | 29,03436 | 29,24864 | 30,3742 | 30,44362 | 30,11706 | 24 | 16 |
| 27,33339 | 26,74548 | 27,17493 | 28,68746 | 28,28069 | 28,28635 | 11 | 11 |
| 26,33209 | 26,69405 | 26,02608 | 29,00849 | 28,83228 | 28,65448 | 9 | 9 |
| 27,59598 | 27,72845 | 27,20881 | 28,54436 | 28,80338 | 28,73292 | 8 | 8 |
| 24,31424 | 22,42708 | 22,99498 | 30,11644 | 29,76708 | 29,85717 | 16 | 7 |
| 27,62136 | 27,98174 | 27,16691 | 29,01422 | 28,74243 | 28,93258 | 6 | 6 |
| 26,48238 | 24,89758 | 24,51767 | 27,64753 | 27,43881 | 27,17703 | 6 | 6 |
| 27,15064 | 27,11347 | 27,37551 | 28,26691 | 29,01364 | 28,94344 | 6 | 6 |
| 25,78913 | 25,41216 | 25,97165 | 26,9888 | 26,82177 | 26,93731 | 4 | 4 |
| 22,02223 | 22,70115 | 22,52501 | 25,5695 | 25,12353 | 25,25465 | 3 | 3 |
| 25,66926 | 25,50513 | 25,2514 | 27,34206 | 27,31544 | 27,38228 | 3 | 3 |
| 25,98501 | 25,84518 | 25,65881 | 27,56623 | 26,91285 | 26,54887 | 3 | 3 |
| 22,62094 | 20,1984 | 22,22947 | 24,40171 | 23,90701 | 24,26842 | 2 | 2 |
| 21,11424 | 22,34051 | 22,09617 | 23,58761 | 23,15826 | 23,28757 | 2 | 2 |
| 23,57877 | 21,0264 | 21,06755 | 24,29593 | 24,20404 | 24,44411 | 2 | 2 |
| 23,63909 | 24,16676 | 23,88688 | 25,43736 | 25,72585 | 26,10043 | 2 | 2 |
| 24,48487 | 24,86722 | 24,31161 | 26,06063 | 25,97066 | 25,7891 | 2 | 2 |
| 20,55546 | 23,84642 | 21,41539 | 25,27316 | 24,90545 | 25,11474 | 10 | 1 |
| 25,51316 | 25,52576 | 22,64945 | 27,30208 | 27,90816 | 27,57967 | 27 | 1 |

| $^{bio}$*Ub* -1- | $^{bio}$*Ub* -2- | $^{bio}$*Ub* -3- | $^{bio}$*ari* -1- | $^{bio}$*ari* -2- | $^{bio}$*ari* -3- | Peptides | Unique peptides |
|---|---|---|---|---|---|---|---|
| 33,15063 | 33,19809 | 33,34916 | 31,83311 | 31,73505 | 31,79796 | 43 | 37 |
| 30,96898 | 30,75231 | 30,86541 | 28,99199 | 29,80721 | 29,22826 | 22 | 22 |
| 32,67632 | 32,58325 | 32,37938 | 31,51493 | 31,28162 | 31,24994 | 21 | 21 |
| 28,37327 | 28,60534 | 28,58459 | 27,31199 | 27,78475 | 27,40381 | 23 | 17 |
| 30,52533 | 30,53797 | 30,69511 | 29,2049 | 29,50692 | 29,19145 | 15 | 15 |
| 30,59147 | 30,31062 | 30,58603 | 29,31853 | 29,52234 | 29,58837 | 15 | 15 |
| 29,33996 | 29,76034 | 29,55611 | 28,21308 | 28,37464 | 28,51552 | 15 | 15 |
| 30,41387 | 30,28162 | 30,29909 | 29,1278 | 29,33038 | 29,128 | 14 | 14 |
| 31,1267 | 31,0236 | 30,84521 | 29,95491 | 29,87179 | 29,90067 | 15 | 14 |
| 28,68479 | 28,22936 | 28,32199 | 26,47015 | 25,96362 | 26,18052 | 12 | 12 |
| 29,62921 | 29,30245 | 29,45077 | 28,49033 | 28,56852 | 28,31363 | 10 | 10 |
| 31,25062 | 31,11514 | 30,78706 | 29,83296 | 30,15001 | 29,88277 | 10 | 10 |
| 27,78013 | 27,97295 | 27,68833 | 26,52115 | 26,54712 | 26,48567 | 8 | 8 |
| 29,63036 | 29,23414 | 28,96433 | 27,89001 | 27,9667 | 28,11664 | 8 | 8 |
| 29,47551 | 29,83945 | 29,50233 | 28,5947 | 27,83936 | 28,43667 | 8 | 8 |
| 29,80404 | 30,05582 | 29,87986 | 29,29489 | 28,60293 | 28,37044 | 7 | 7 |
| 27,01084 | 25,89825 | 26,81164 | 24,94114 | 24,42047 | 24,28623 | 7 | 7 |
| 26,89527 | 26,37202 | 26,18682 | 25,33071 | 25,30633 | 25,25857 | 14 | 6 |
| 26,55522 | 26,37431 | 27,1034 | 25,73492 | 25,27085 | 25,63481 | 6 | 6 |
| 29,76417 | 29,34172 | 29,32967 | 27,97016 | 28,28025 | 27,88739 | 7 | 5 |
| 27,5349 | 28,02332 | 27,36411 | 26,52907 | 26,43428 | 26,81702 | 5 | 5 |
| 26,76053 | 26,28477 | 26,25241 | 25,52898 | 25,43733 | 25,06632 | 5 | 5 |

| | | | | | | | |
|---|---|---|---|---|---|---|---|
| 28,62443 | 28,4338 | 28,3674 | 27,57297 | 27,4182 | 27,28445 | 5 | 5 |
| 29,92295 | 30,08626 | 29,67145 | 28,10545 | 28,23963 | 28,45909 | 5 | 5 |
| 26,36575 | 25,58941 | 25,07092 | 24,053 | 24,13962 | 23,67228 | 4 | 4 |
| 26,43965 | 26,50086 | 25,93528 | 25,16699 | 25,19404 | 25,38617 | 4 | 4 |
| 30,81398 | 31,11086 | 31,05769 | 29,91128 | 29,9167 | 29,6463 | 4 | 4 |
| 26,6245 | 26,90104 | 26,77354 | 26,2424 | 25,15449 | 25,83219 | 4 | 4 |
| 26,87244 | 26,77568 | 26,2451 | 25,41245 | 24,88586 | 24,47494 | 4 | 4 |
| 27,85507 | 27,89446 | 27,27728 | 25,17596 | 24,55098 | 24,82026 | 4 | 4 |
| 25,64804 | 25,34864 | 25,63835 | 22,32039 | 23,09885 | 22,05217 | 3 | 3 |
| 25,58423 | 25,68672 | 26,0178 | 23,64548 | 22,83469 | 22,51625 | 4 | 3 |
| 24,35991 | 24,08302 | 23,72837 | 21,66424 | 23,36639 | 21,06888 | 3 | 3 |
| 27,62909 | 27,2779 | 27,56529 | 24,53552 | 25,72268 | 25,39872 | 3 | 3 |
| 25,99062 | 25,37026 | 25,36065 | 24,28989 | 23,74006 | 23,87521 | 3 | 3 |
| 27,50309 | 27,51187 | 27,51412 | 26,42292 | 26,5952 | 26,09712 | 3 | 3 |
| 26,45013 | 26,36816 | 26,58749 | 25,18454 | 25,02682 | 24,99196 | 3 | 3 |
| 24,95899 | 24,73583 | 25,2277 | 24,25025 | 22,66275 | 22,04214 | 2 | 2 |
| 25,88707 | 25,73709 | 24,415 | 22,41354 | 23,67842 | 23,76537 | 2 | 2 |
| 27,95455 | 27,86667 | 27,43547 | 26,47197 | 26,28171 | 26,68244 | 2 | 2 |
| 24,69429 | 24,536 | 24,50227 | 22,78053 | 22,26702 | 22,24784 | 2 | 2 |
| 25,96239 | 25,82539 | 25,56069 | 25,2743 | 24,54916 | 24,26535 | 2 | 2 |
| 25,75918 | 26,31655 | 26,919 | 23,89883 | 24,9679 | 24,19227 | 2 | 2 |
| 29,02131 | 28,91869 | 28,83843 | 27,59939 | 27,90351 | 27,80759 | 22 | 2 |
| 28,31467 | 28,60661 | 28,17331 | 27,25422 | 26,91593 | 27,06157 | 2 | 2 |
| 25,10582 | 24,49857 | 24,12983 | 21,80641 | 21,7161 | 23,36185 | 2 | 2 |
| 25,59272 | 24,57508 | 23,76497 | 22,30939 | 21,50319 | 22,00921 | 2 | 2 |
| 24,61453 | 24,71968 | 25,32471 | 21,61722 | 23,29386 | 21,57574 | 2 | 2 |
| 24,47692 | 24,93988 | 25,18173 | 21,22775 | 21,65075 | 22,11623 | 2 | 2 |
| 23,07912 | 22,86695 | 22,89143 | 21,53477 | 21,61447 | 22,39385 | 2 | 2 |
| 24,31202 | 24,16852 | 24,30918 | 20,19029 | 21,49978 | 22,57826 | 21 | 1 |
| 25,22115 | 25,6329 | 25,30907 | 21,64994 | 21,35846 | 21,16324 | 28 | 1 |
| 24,14756 | 23,5341 | 24,2475 | 21,26969 | 21,50158 | 21,17441 | 12 | 1 |

| $^{bio}$*Ub* -1- | $^{bio}$*Ub* -2- | $^{bio}$*Ub* -3- | $^{bio}$*ari* -1- | $^{bio}$*ari* -2- | $^{bio}$*ari* -3- | **Peptides** | **Unique peptides** |
|---|---|---|---|---|---|---|---|
| 37,07467 | 37,52795 | 37,14386 | 37,5743 | 37,6105 | 37,68767 | 143 | 143 |
| 38,37535 | 38,23203 | 38,5571 | 38,46444 | 38,65171 | 38,66163 | 21 | 21 |
| 39,22315 | 39,37449 | 39,427 | 40,09825 | 39,90732 | 39,83116 | 111 | 2 |
| 29,4583 | 29,50728 | 29,09464 | 29,76077 | 29,89422 | 29,51616 | 17 | 1 |
| 24,11362 | 20,69188 | 24,22428 | 24,51953 | 24,93157 | 24,51364 | 110 | 1 |
| 40,31067 | 40,16648 | 40,06402 | 39,65578 | 39,70975 | 39,81664 | 40 | 21 |
| 30,23181 | 30,37015 | 30,32496 | 29,55188 | 29,739 | 29,88906 | 38 | 38 |
| 29,2169 | 29,55784 | 29,45197 | 29,15875 | 28,92912 | 28,99809 | 24 | 24 |
| 25,68921 | 25,50945 | 25,91946 | 25,04093 | 24,93899 | 25,25425 | 6 | 6 |
| 32,52542 | 32,29863 | 32,46551 | 33,34652 | 33,38291 | 33,35205 | 27 | 1 |
| 24,80174 | 24,79556 | 25,05685 | 25,20602 | 25,40695 | 25,20311 | 3 | 3 |
| 26,68979 | 26,6569 | 26,97695 | 27,01127 | 27,19385 | 27,16221 | 8 | 8 |
| 24,87183 | 24,19468 | 24,92895 | 24,16668 | 23,73469 | 23,7214 | 4 | 4 |
| 29,31657 | 28,68736 | 28,92286 | 30,06458 | 29,74386 | 29,72559 | 14 | 14 |
| 27,93393 | 27,71411 | 27,34571 | 26,77102 | 26,78357 | 26,73596 | 6 | 6 |
| 29,50179 | 29,12519 | 29,15197 | 28,63513 | 28,61919 | 28,47421 | 9 | 9 |

| | | | | | | | |
|---|---|---|---|---|---|---|---|
| 30,73723 | 30,58692 | 30,70636 | 30,12238 | 30,38628 | 30,23638 | 15 | 15 |
| 28,39938 | 28,12721 | 28,23444 | 27,62345 | 27,73447 | 27,5423 | 10 | 10 |
| 31,3639 | 31,14273 | 31,23934 | 31,07003 | 30,83017 | 30,93853 | 67 | 67 |
| 27,62303 | 27,58248 | 27,61737 | 27,25944 | 26,75888 | 26,88402 | 6 | 6 |
| 30,28295 | 30,02327 | 29,93649 | 29,42864 | 29,81415 | 29,35645 | 7 | 7 |
| 30,0624 | 29,9776 | 30,05414 | 30,22515 | 30,2366 | 30,37793 | 16 | 16 |
| 27,67209 | 27,5979 | 27,65458 | 28,15543 | 27,96444 | 27,99243 | 5 | 5 |
| 30,97869 | 30,84872 | 30,94949 | 31,25175 | 31,27475 | 31,20961 | 35 | 35 |
| 29,78673 | 29,55214 | 29,61159 | 29,24611 | 29,06361 | 29,23806 | 8 | 1 |
| 27,08375 | 27,1039 | 27,08891 | 26,36663 | 26,85698 | 26,60272 | 13 | 3 |
| 29,26669 | 29,22064 | 29,19789 | 29,57962 | 29,74839 | 29,48383 | 22 | 22 |
| 25,10077 | 25,05134 | 25,56671 | 25,94813 | 25,60071 | 26,05267 | 2 | 2 |
| 31,5163 | 31,67943 | 31,59348 | 31,9481 | 31,95342 | 32,08027 | 46 | 46 |
| 27,67916 | 27,39689 | 27,85281 | 26,88017 | 26,96884 | 26,82869 | 8 | 8 |
| 25,63945 | 26,16254 | 25,78267 | 26,47765 | 26,47073 | 26,5122 | 2 | 2 |
| 30,13602 | 30,15581 | 30,16507 | 29,57198 | 29,72164 | 29,79658 | 21 | 21 |
| 25,24905 | 25,53059 | 25,39359 | 24,83851 | 25,07961 | 24,45331 | 3 | 2 |
| 29,09939 | 28,8964 | 29,08066 | 28,61894 | 28,57582 | 28,26432 | 11 | 11 |
| 30,8455 | 30,44164 | 30,51409 | 31,57434 | 31,66729 | 31,38268 | 29 | 29 |
| 27,76585 | 27,47251 | 27,77361 | 27,21178 | 26,86938 | 27,39403 | 11 | 11 |
| 26,8867 | 27,31984 | 27,12593 | 28,24004 | 27,90776 | 27,75863 | 18 | 5 |
| 27,73467 | 27,74849 | 27,92275 | 27,4223 | 27,16116 | 27,23708 | 7 | 7 |
| 32,75149 | 32,89405 | 32,72799 | 31,97692 | 32,24607 | 32,00992 | 5 | 5 |
| 28,48001 | 28,33782 | 28,19319 | 27,618 | 27,7835 | 27,37609 | 17 | 17 |
| 29,08459 | 28,93663 | 29,02065 | 28,25696 | 28,29044 | 27,84134 | 9 | 9 |
| 33,50178 | 33,44825 | 33,03823 | 32,58982 | 32,61938 | 32,58985 | 33 | 17 |
| 31,46285 | 31,37415 | 31,10431 | 30,78714 | 30,50711 | 30,56901 | 10 | 10 |
| 29,1306 | 29,07778 | 28,80196 | 29,41657 | 29,41945 | 29,32759 | 6 | 6 |
| 32,99313 | 32,9334 | 32,82707 | 32,81153 | 32,65002 | 32,56978 | 117 | 19 |
| 28,63429 | 28,81369 | 28,83301 | 27,89678 | 27,7855 | 27,73079 | 6 | 6 |
| 25,22295 | 25,42342 | 25,63467 | 25,09993 | 24,63848 | 24,73387 | 2 | 2 |
| 30,69669 | 30,34763 | 30,65881 | 29,68569 | 30,03892 | 29,90412 | 15 | 2 |
| 24,46039 | 24,61133 | 24,3761 | 24,30341 | 24,05846 | 24,19903 | 2 | 2 |
| 26,92468 | 26,68699 | 26,9015 | 26,33508 | 25,78187 | 26,44679 | 5 | 5 |
| 28,15928 | 28,03344 | 28,26732 | 28,31034 | 28,4352 | 28,57636 | 12 | 12 |
| 28,06018 | 28,06815 | 27,46022 | 28,6599 | 28,83253 | 28,68315 | 10 | 10 |
| 26,385 | 25,61538 | 26,03997 | 25,44092 | 24,68442 | 25,10258 | 2 | 2 |
| 26,61049 | 26,65554 | 27,08952 | 25,84364 | 26,42594 | 25,74725 | 5 | 5 |
| 29,67459 | 29,54922 | 29,63523 | 29,31532 | 29,4008 | 29,21843 | 14 | 14 |
| 32,71305 | 32,80313 | 32,40819 | 31,88169 | 32,076 | 31,93139 | 21 | 21 |
| 25,29119 | 25,55488 | 25,61012 | 24,35743 | 24,89463 | 24,81787 | 2 | 2 |
| 28,33573 | 28,03475 | 28,12105 | 27,80033 | 27,51976 | 27,40414 | 6 | 6 |
| 27,80236 | 27,65928 | 27,70794 | 27,40576 | 26,85234 | 26,42488 | 9 | 3 |
| 34,31999 | 34,19484 | 33,58755 | 34,65128 | 34,67588 | 34,67651 | 31 | 25 |
| 28,78946 | 28,67795 | 28,58538 | 28,06897 | 28,39681 | 28,3132 | 22 | 22 |
| 24,85781 | 24,49479 | 24,51521 | 23,75369 | 23,69328 | 24,17013 | 3 | 3 |
| 30,83605 | 30,79576 | 30,91599 | 31,23181 | 31,28846 | 31,32764 | 47 | 47 |
| 25,69875 | 25,50709 | 25,6345 | 26,42702 | 26,14396 | 25,98089 | 4 | 4 |
| 29,75785 | 29,81636 | 29,63 | 28,91895 | 29,34336 | 29,19131 | 13 | 13 |
| 29,68482 | 29,64991 | 29,68211 | 28,98669 | 28,5037 | 28,91271 | 18 | 18 |

| | | | | | | | |
|---|---|---|---|---|---|---|---|
| 32,44663 | 32,54471 | 32,21873 | 32,14801 | 31,86548 | 31,73093 | 12 | 12 |
| 28,65513 | 28,69269 | 28,65567 | 28,83975 | 29,06296 | 28,82996 | 9 | 9 |
| 26,72078 | 26,9085 | 27,10189 | 26,25276 | 25,96206 | 25,89615 | 2 | 2 |
| 27,31544 | 27,26132 | 26,87807 | 26,50108 | 26,39873 | 26,04968 | 5 | 5 |
| 33,49679 | 33,54467 | 33,1319 | 32,85309 | 33,0708 | 32,9288 | 26 | 24 |
| 32,1498 | 32,14677 | 32,01408 | 31,55828 | 31,60076 | 31,61938 | 14 | 5 |
| 25,68429 | 25,62623 | 24,99378 | 26,08016 | 26,5209 | 26,27293 | 4 | 4 |
| 28,84242 | 28,43432 | 28,4481 | 27,98174 | 27,86944 | 27,19714 | 14 | 13 |
| 31,30791 | 31,33905 | 31,18943 | 30,93677 | 30,79815 | 30,73409 | 28 | 28 |
| 29,6657 | 29,51646 | 29,59238 | 29,99348 | 30,05634 | 30,04739 | 23 | 23 |
| 29,74431 | 29,41121 | 29,49728 | 28,99129 | 29,10575 | 29,02509 | 13 | 11 |
| 27,45928 | 27,46076 | 27,43349 | 26,8818 | 27,29421 | 27,01924 | 10 | 10 |
| 29,54337 | 29,59786 | 28,89432 | 28,50089 | 28,52865 | 28,6568 | 6 | 6 |
| 26,71542 | 27,01648 | 26,7661 | 27,56805 | 27,26078 | 27,79315 | 5 | 5 |
| 31,73622 | 31,71751 | 31,59406 | 32,3034 | 32,17593 | 31,98206 | 13 | 13 |
| 23,81274 | 24,14957 | 23,955 | 24,30487 | 24,28687 | 24,36646 | 2 | 2 |
| 24,0476 | 24,20658 | 24,15831 | 24,912 | 25,0068 | 24,95371 | 3 | 3 |
| 31,3802 | 31,18382 | 31,0768 | 31,72761 | 31,44865 | 31,56792 | 28 | 28 |
| 27,91906 | 27,90466 | 27,95532 | 28,14594 | 28,08396 | 28,05553 | 9 | 9 |
| 27,05559 | 27,17465 | 26,86455 | 26,21098 | 26,53398 | 26,08123 | 4 | 4 |
| 28,60527 | 28,72832 | 28,85831 | 28,05672 | 28,23174 | 28,10405 | 9 | 9 |
| 28,20644 | 28,4434 | 27,93016 | 28,8557 | 28,91274 | 28,50362 | 8 | 8 |
| 30,83936 | 30,76825 | 30,69345 | 30,21199 | 30,02314 | 30,23957 | 22 | 22 |
| 29,46131 | 29,40975 | 29,2871 | 28,57261 | 28,50582 | 28,7254 | 17 | 14 |
| 26,39536 | 26,38407 | 26,23063 | 27,30947 | 27,31311 | 27,15981 | 3 | 3 |
| 30,22768 | 29,96678 | 29,69166 | 30,43778 | 30,70413 | 30,53529 | 14 | 14 |
| 31,03308 | 30,60726 | 30,75883 | 30,20432 | 30,23306 | 30,33692 | 16 | 16 |
| 30,19742 | 30,40943 | 29,72436 | 29,1052 | 29,24663 | 29,53783 | 8 | 8 |
| 24,20494 | 24,41416 | 24,5764 | 24,75022 | 25,35468 | 25,36433 | 6 | 6 |
| 28,48312 | 28,4434 | 28,00715 | 28,75363 | 28,88475 | 28,77878 | 10 | 10 |
| 24,44834 | 25,06788 | 24,54445 | 23,68731 | 23,68389 | 23,70936 | 3 | 3 |
| 29,5751 | 29,42946 | 29,75159 | 30,16866 | 30,24875 | 30,26495 | 14 | 14 |
| 28,60848 | 28,23558 | 28,30639 | 28,88527 | 28,7726 | 28,87786 | 5 | 5 |
| 29,37962 | 29,80538 | 29,73192 | 29,18634 | 29,3353 | 28,96642 | 5 | 5 |
| 31,52374 | 31,47237 | 31,11582 | 30,89085 | 30,76975 | 30,87855 | 7 | 7 |
| 35,39663 | 35,34414 | 35,45586 | 35,56882 | 35,57119 | 35,70063 | 57 | 3 |
| 33,15355 | 33,04282 | 32,90158 | 33,79981 | 33,36437 | 33,42079 | 99 | 99 |
| 31,73542 | 31,81406 | 31,74979 | 31,09087 | 31,07387 | 31,08822 | 18 | 18 |
| 32,21037 | 32,20376 | 32,14394 | 32,03361 | 32,1395 | 32,04101 | 170 | 135 |
| 33,35928 | 33,25476 | 33,30157 | 33,13836 | 33,13868 | 33,16942 | 36 | 36 |
| 31,05019 | 30,82457 | 31,05265 | 30,58073 | 30,53334 | 30,48932 | 36 | 36 |
| 30,18127 | 29,92451 | 30,03656 | 29,38268 | 29,4907 | 29,24382 | 25 | 25 |
| 33,64216 | 33,60554 | 33,49226 | 33,86064 | 33,69229 | 33,83112 | 104 | 1 |
| 33,55373 | 33,39803 | 33,17639 | 33,70016 | 33,79178 | 33,70821 | 28 | 28 |
| 27,55735 | 27,43897 | 27,4128 | 27,12711 | 27,06188 | 27,00024 | 4 | 4 |
| 26,47823 | 26,603 | 26,47091 | 26,33093 | 26,02526 | 26,14585 | 6 | 6 |
| 26,09027 | 25,94682 | 26,35548 | 26,90541 | 26,99583 | 27,10009 | 8 | 8 |
| 24,76689 | 24,3654 | 24,82905 | 25,701 | 25,3979 | 25,65944 | 4 | 4 |
| 26,67223 | 26,16062 | 25,96113 | 25,65892 | 25,48778 | 25,72721 | 3 | 3 |
| 26,15368 | 26,89516 | 26,57032 | 25,58274 | 25,63243 | 25,45209 | 6 | 6 |

| | | | | | | | |
|---|---|---|---|---|---|---|---|
| 29,67845 | 29,80855 | 29,78036 | 29,40916 | 29,62587 | 29,51826 | 10 | 10 |
| 28,49171 | 28,2787 | 28,36749 | 27,7965 | 27,64588 | 28,03244 | 16 | 16 |
| 31,31154 | 31,15816 | 31,34038 | 30,7171 | 30,86356 | 30,91185 | 27 | 25 |
| 27,52567 | 28,08461 | 27,88011 | 28,78288 | 28,68248 | 28,23982 | 59 | 2 |
| 29,81135 | 29,81025 | 29,62479 | 30,42061 | 30,762 | 30,48078 | 15 | 15 |
| 26,99938 | 27,18234 | 27,40527 | 26,0341 | 26,52556 | 26,02745 | 4 | 4 |
| 29,13193 | 29,24531 | 28,88192 | 30,01531 | 30,03459 | 29,82596 | 11 | 11 |
| 24,25954 | 24,23564 | 24,4523 | 25,70023 | 24,96662 | 25,01329 | 2 | 2 |
| 30,23238 | 30,23831 | 30,28372 | 30,39836 | 30,44579 | 30,65113 | 5 | 5 |
| 29,77549 | 29,87314 | 29,55437 | 29,36353 | 29,21134 | 29,19411 | 8 | 3 |
| 27,73104 | 27,72449 | 27,58133 | 27,13995 | 27,18669 | 27,308 | 7 | 7 |
| 27,52671 | 27,45018 | 27,79334 | 27,05072 | 26,82627 | 26,76192 | 9 | 9 |
| 26,0579 | 25,83985 | 25,60368 | 24,98259 | 25,23557 | 25,22707 | 2 | 1 |
| 26,47105 | 25,72286 | 26,45772 | 25,38913 | 25,57355 | 24,9757 | 3 | 3 |
| 28,05429 | 27,91815 | 27,63312 | 28,6554 | 28,3413 | 28,33981 | 7 | 7 |
| 30,69254 | 30,47237 | 30,89973 | 29,87893 | 30,18872 | 30,07125 | 10 | 5 |
| 26,97235 | 27,01052 | 26,48378 | 26,39566 | 25,60122 | 25,74561 | 5 | 5 |
| 30,88787 | 30,78246 | 30,4446 | 31,20862 | 31,41921 | 31,32012 | 9 | 9 |
| 31,89331 | 31,74699 | 31,95404 | 31,07476 | 31,32657 | 31,0879 | 16 | 16 |
| 31,81719 | 31,81272 | 31,30905 | 31,08455 | 31,20414 | 31,02756 | 21 | 2 |
| 26,23416 | 26,87643 | 26,27195 | 27,25836 | 27,08659 | 27,77881 | 4 | 4 |
| 33,48123 | 33,56207 | 33,4184 | 34,18601 | 33,94559 | 33,82364 | 70 | 13 |
| 28,25485 | 28,35834 | 28,3416 | 28,75834 | 28,63527 | 28,68846 | 14 | 14 |
| 28,53453 | 28,36353 | 28,21341 | 28,11033 | 27,82153 | 27,62192 | 8 | 8 |
| 31,77396 | 31,59566 | 31,84689 | 31,10124 | 31,06079 | 30,8514 | 23 | 23 |
| 29,69861 | 29,87322 | 29,66885 | 30,09836 | 30,55901 | 30,37223 | 55 | 1 |
| 30,51634 | 30,5913 | 30,55326 | 30,9504 | 30,75708 | 30,97117 | 24 | 24 |
| 26,59007 | 26,79017 | 26,32364 | 27,87631 | 27,38187 | 26,9536 | 5 | 5 |
| 33,64098 | 33,3979 | 33,59924 | 34,35054 | 34,19183 | 34,01878 | 59 | 59 |
| 28,58671 | 28,51345 | 28,12194 | 27,48308 | 27,47576 | 27,31683 | 4 | 4 |
| 31,14163 | 31,11706 | 31,12022 | 31,6203 | 31,45875 | 31,52529 | 18 | 18 |
| 31,01458 | 30,96726 | 30,86386 | 31,30596 | 31,19566 | 31,18494 | 28 | 28 |
| 30,78869 | 30,79977 | 30,96148 | 30,62061 | 30,68779 | 30,7038 | 42 | 42 |
| 30,96712 | 31,09295 | 30,60576 | 31,81749 | 31,73016 | 31,59726 | 17 | 17 |
| 27,52103 | 27,52694 | 27,6513 | 26,31287 | 27,01691 | 27,08132 | 7 | 7 |
| 31,45479 | 31,54278 | 31,35332 | 32,11169 | 31,9953 | 31,96959 | 27 | 27 |
| 31,33899 | 31,59899 | 31,37088 | 31,13418 | 31,11023 | 31,13559 | 5 | 5 |
| 29,47224 | 29,39652 | 29,55945 | 28,97981 | 28,88931 | 29,11076 | 19 | 19 |
| 28,56779 | 28,31743 | 28,08385 | 27,79129 | 27,95521 | 27,91718 | 8 | 8 |
| 28,63703 | 28,16183 | 28,1043 | 29,29581 | 29,08499 | 29,10789 | 9 | 9 |
| 28,30456 | 28,07199 | 27,70872 | 28,78251 | 28,68572 | 28,87844 | 10 | 10 |
| 29,92465 | 29,9057 | 30,07521 | 30,53621 | 30,53204 | 30,2007 | 18 | 18 |
| 24,79427 | 24,65015 | 24,51671 | 23,86022 | 24,06669 | 24,02572 | 2 | 2 |
| 35,33135 | 35,20816 | 35,17604 | 34,40889 | 34,60747 | 34,46094 | 42 | 42 |
| 29,90899 | 29,95892 | 29,66078 | 30,16926 | 30,28791 | 30,09861 | 10 | 10 |
| 25,95103 | 25,92093 | 25,4947 | 25,22155 | 25,15727 | 25,13179 | 2 | 2 |
| 28,19394 | 28,08375 | 27,67788 | 27,30234 | 27,58334 | 27,14014 | 7 | 7 |
| 28,18423 | 27,75576 | 27,91946 | 28,71797 | 28,71303 | 28,84281 | 19 | 19 |
| 25,91235 | 25,66825 | 25,79097 | 25,27544 | 24,6402 | 24,69386 | 3 | 3 |
| 31,45503 | 31,37456 | 31,47141 | 30,75693 | 30,61263 | 30,46188 | 20 | 20 |

| | | | | | | | |
|---|---|---|---|---|---|---|---|
| 27,31259 | 27,12751 | 27,295 | 27,80858 | 27,94151 | 27,66064 | 6 | 6 |
| 25,37945 | 25,19107 | 24,71837 | 24,61482 | 24,23863 | 24,43924 | 2 | 2 |
| 26,65458 | 26,34853 | 26,46658 | 26,87537 | 26,9577 | 26,80156 | 5 | 5 |
| 31,31872 | 31,50824 | 31,39622 | 31,98206 | 32,00035 | 32,04576 | 23 | 1 |
| 30,13528 | 29,98101 | 30,2099 | 29,33705 | 29,4231 | 29,56258 | 27 | 27 |
| 26,66153 | 26,76522 | 25,94737 | 25,17016 | 25,64968 | 25,81965 | 5 | 5 |
| 24,84527 | 24,74304 | 25,22883 | 25,59665 | 25,72695 | 25,95491 | 2 | 2 |
| 27,65294 | 27,50142 | 27,75551 | 26,9941 | 27,13016 | 26,74945 | 5 | 5 |
| 28,59192 | 28,56344 | 28,39137 | 29,5253 | 29,30189 | 29,24434 | 4 | 4 |
| 26,23281 | 26,36796 | 26,10524 | 25,53884 | 25,43638 | 25,39953 | 5 | 5 |
| 28,62607 | 28,96521 | 28,87953 | 28,15605 | 27,48677 | 27,93629 | 3 | 3 |
| 29,88397 | 29,60906 | 29,75427 | 29,30062 | 29,43936 | 29,27115 | 18 | 3 |
| 28,27631 | 28,29141 | 28,26504 | 27,94682 | 27,86856 | 28,08836 | 10 | 10 |
| 29,19101 | 29,22037 | 29,37809 | 28,6633 | 28,84461 | 28,92751 | 20 | 20 |
| 32,36872 | 32,55595 | 31,99682 | 31,33634 | 31,4482 | 31,33026 | 16 | 16 |
| 25,3644 | 24,66074 | 24,67228 | 24,17988 | 23,76182 | 24,21571 | 2 | 2 |
| 28,36961 | 28,27209 | 27,98506 | 28,66973 | 28,7714 | 28,77869 | 11 | 11 |
| 32,64182 | 32,51239 | 32,5839 | 32,45012 | 32,40626 | 32,46117 | 34 | 31 |
| 28,11317 | 27,94218 | 27,94453 | 27,70471 | 27,60738 | 27,71784 | 5 | 5 |
| 28,86201 | 29,01675 | 28,98961 | 28,00287 | 28,42194 | 27,71332 | 8 | 8 |
| 27,16825 | 27,1947 | 27,11744 | 26,15715 | 26,60569 | 26,64952 | 4 | 4 |
| 29,51486 | 29,54195 | 29,31283 | 29,04012 | 29,05302 | 29,01337 | 7 | 7 |
| 29,01561 | 28,86606 | 29,02649 | 28,59285 | 28,71954 | 28,82211 | 15 | 15 |
| 27,4903 | 27,5326 | 27,40292 | 26,28049 | 26,74343 | 26,55801 | 7 | 7 |
| 30,18116 | 30,06149 | 30,10774 | 30,82873 | 30,65957 | 30,47528 | 18 | 2 |
| 30,28196 | 30,517 | 30,51117 | 30,85542 | 31,09572 | 31,02214 | 19 | 3 |
| 29,78071 | 29,66499 | 29,80184 | 30,24717 | 30,25214 | 30,04739 | 11 | 11 |
| 26,56075 | 26,16448 | 26,09051 | 25,54901 | 25,25598 | 25,48925 | 3 | 3 |
| 28,89423 | 28,72956 | 28,73551 | 28,07653 | 28,58007 | 28,18329 | 11 | 11 |
| 34,12054 | 33,94236 | 33,80492 | 33,73973 | 33,59379 | 33,67829 | 28 | 28 |
| 35,98958 | 35,66389 | 36,08001 | 36,39418 | 36,27152 | 36,55027 | 73 | 5 |
| 26,89261 | 26,98663 | 26,87783 | 26,6245 | 26,07069 | 26,3191 | 2 | 2 |
| 28,20974 | 28,00667 | 27,65109 | 27,47893 | 26,99927 | 27,25773 | 4 | 4 |
| 27,23663 | 26,79476 | 27,219 | 27,96312 | 27,99706 | 27,58592 | 5 | 5 |
| 28,08137 | 28,07378 | 27,45073 | 27,1776 | 27,1849 | 27,3667 | 6 | 2 |
| 25,15804 | 25,6966 | 25,46989 | 26,1865 | 26,03712 | 26,01536 | 2 | 2 |
| 32,28339 | 32,22282 | 32,21208 | 31,86426 | 31,93206 | 31,76236 | 88 | 3 |
| 31,71227 | 31,39759 | 31,29565 | 30,67228 | 30,81665 | 30,80078 | 31 | 31 |
| 27,04864 | 26,55926 | 26,8423 | 27,60894 | 27,43579 | 27,74073 | 12 | 12 |
| 30,60594 | 30,42432 | 30,29625 | 30,91171 | 30,9386 | 31,10849 | 13 | 13 |
| 28,4837 | 28,35515 | 28,28868 | 28,6441 | 28,74037 | 28,6599 | 13 | 12 |
| 28,98905 | 28,61558 | 28,87185 | 28,30669 | 28,34087 | 28,38665 | 7 | 5 |
| 31,36134 | 31,0898 | 31,31506 | 30,58101 | 30,64924 | 30,87774 | 44 | 44 |
| 25,57476 | 25,80877 | 25,63824 | 24,53005 | 25,15892 | 25,17177 | 3 | 3 |
| 27,97328 | 27,59491 | 27,59974 | 27,14375 | 27,42358 | 27,32852 | 8 | 8 |
| 29,38156 | 29,59146 | 29,64816 | 28,69959 | 28,96179 | 29,19348 | 16 | 16 |
| 27,9526 | 27,37815 | 27,97356 | 26,88297 | 26,65963 | 27,18688 | 9 | 9 |
| 27,16279 | 27,19216 | 27,36736 | 26,88262 | 26,22376 | 26,41303 | 7 | 7 |
| 30,3572 | 30,35142 | 29,94419 | 29,83623 | 29,50522 | 29,55228 | 11 | 11 |
| 28,23549 | 28,04745 | 27,9673 | 27,1916 | 27,01169 | 27,46272 | 3 | 3 |

| | | | | | | | |
|---|---|---|---|---|---|---|---|
| 29,63808 | 29,44875 | 29,32956 | 30,23957 | 30,18789 | 29,9924 | 27 | 1 |
| 27,22849 | 26,99162 | 27,04562 | 26,74945 | 26,81934 | 26,36414 | 12 | 12 |
| 25,75478 | 26,06807 | 26,14909 | 25,28164 | 24,50609 | 25,21212 | 5 | 5 |
| 27,88419 | 28,36628 | 27,86302 | 28,83328 | 28,81115 | 28,88387 | 2 | 2 |
| 30,07635 | 30,01052 | 29,95754 | 29,72436 | 29,58036 | 29,75966 | 25 | 25 |
| 29,43386 | 29,34517 | 28,88736 | 28,39579 | 28,51642 | 28,82623 | 6 | 6 |
| 29,13746 | 29,05437 | 28,58996 | 28,07801 | 28,18248 | 28,2294 | 9 | 8 |
| 30,21037 | 30,39683 | 30,35121 | 29,99739 | 29,95685 | 29,91713 | 16 | 16 |
| 26,49576 | 26,56496 | 27,12888 | 25,64719 | 26,09208 | 25,5636 | 7 | 7 |
| 30,58459 | 30,37865 | 30,34942 | 30,00718 | 30,23957 | 30,15726 | 17 | 17 |
| 27,1454 | 26,97104 | 26,53702 | 26,39642 | 26,02246 | 25,9817 | 6 | 6 |
| 23,6937 | 24,39859 | 24,48702 | 25,52373 | 25,22339 | 24,78477 | 2 | 2 |
| 31,2228 | 30,94517 | 31,11384 | 30,06907 | 30,36234 | 30,38051 | 16 | 16 |
| 30,91335 | 30,87394 | 31,03892 | 31,13498 | 31,12737 | 31,12497 | 26 | 26 |
| 28,99448 | 28,99531 | 29,00386 | 28,45376 | 28,4945 | 28,34066 | 20 | 20 |
| 28,95604 | 28,79293 | 28,88349 | 29,56215 | 29,69995 | 29,33211 | 12 | 12 |
| 29,53999 | 29,72403 | 29,70015 | 28,82199 | 28,93163 | 28,68385 | 14 | 14 |
| 30,34032 | 30,27841 | 30,18459 | 30,13112 | 29,93227 | 29,87226 | 19 | 19 |
| 26,95848 | 26,67088 | 26,57505 | 26,41529 | 26,05463 | 25,75728 | 2 | 2 |
| 30,36693 | 30,46236 | 30,46568 | 29,87247 | 30,16579 | 29,99078 | 11 | 11 |
| 25,45782 | 25,89994 | 25,71025 | 25,06489 | 25,18771 | 25,20191 | 4 | 4 |
| 28,66289 | 28,53356 | 28,16892 | 27,72058 | 27,93286 | 27,88518 | 4 | 4 |
| 32,07985 | 31,90849 | 31,96227 | 32,19325 | 32,31008 | 32,20868 | 34 | 34 |
| 28,57809 | 28,48566 | 28,5782 | 28,76797 | 28,76253 | 28,79433 | 9 | 9 |
| 32,75053 | 32,46779 | 32,62536 | 32,87295 | 32,93055 | 32,8641 | 28 | 1 |
| 28,08922 | 28,09728 | 28,10189 | 27,71417 | 27,41981 | 27,5768 | 9 | 9 |
| 26,42403 | 26,5342 | 26,21843 | 27,37567 | 26,79885 | 26,81311 | 5 | 5 |
| 27,56107 | 27,80366 | 27,87947 | 28,49511 | 28,08324 | 28,10749 | 6 | 6 |
| 25,16814 | 24,8974 | 24,60269 | 24,17219 | 24,16354 | 24,45306 | 2 | 2 |
| 29,19535 | 29,0966 | 29,05264 | 28,75786 | 28,8929 | 28,87074 | 10 | 10 |
| 28,06157 | 27,99102 | 27,50854 | 27,13212 | 27,45764 | 27,02136 | 7 | 7 |
| 30,12952 | 30,03433 | 30,2294 | 29,62361 | 29,6298 | 29,7345 | 21 | 21 |
| 26,43223 | 26,60837 | 26,57371 | 26,08687 | 25,952 | 26,30805 | 2 | 2 |
| 28,33981 | 28,15393 | 28,34422 | 28,09336 | 27,89093 | 27,64924 | 12 | 12 |
| 28,66289 | 28,50158 | 28,31648 | 27,93809 | 27,89977 | 27,81592 | 8 | 8 |
| 24,01504 | 24,52409 | 23,92904 | 24,64257 | 24,64812 | 24,87652 | 3 | 3 |
| 29,70601 | 29,4303 | 29,82675 | 28,76344 | 28,75834 | 28,76227 | 17 | 17 |
| 29,74998 | 29,50372 | 29,83892 | 28,70243 | 29,03839 | 28,91553 | 12 | 12 |
| 30,40902 | 30,28295 | 30,43002 | 29,39301 | 29,59632 | 29,38039 | 10 | 4 |
| 23,74427 | 23,62255 | 22,96556 | 22,85251 | 21,14264 | 20,93041 | 86 | 1 |
| 24,81495 | 23,72775 | 23,13781 | 23,11472 | 22,16371 | 22,48102 | 2 | 2 |
| 25,35538 | 25,80772 | 27,53697 | 25,74776 | 24,36866 | 25,01444 | 2 | 2 |
| 24,81973 | 24,75231 | 24,25386 | 24,64334 | 24,5134 | 21,39024 | 2 | 2 |
| 25,50546 | 25,36376 | 24,75918 | 25,47615 | 25,11004 | 21,41248 | 2 | 2 |
| 23,96957 | 24,28637 | 24,69594 | 22,48405 | 22,71165 | 24,35676 | 2 | 2 |
| 26,29681 | 25,43065 | 25,75313 | 25,68589 | 25,05978 | 22,68383 | 3 | 3 |
| 25,58901 | 25,85167 | 25,85531 | 25,43939 | 22,66872 | 23,0029 | 4 | 4 |
| 25,94798 | 24,49332 | 25,75154 | 24,80995 | 21,41968 | 25,3811 | 2 | 2 |
| 24,02293 | 23,49143 | 26,46756 | 22,19218 | 23,39689 | 23,27555 | 9 | 2 |
| 26,15518 | 25,93973 | 25,28344 | 25,33252 | 22,36539 | 25,72541 | 2 | 2 |

| | | | | | | | |
|---|---|---|---|---|---|---|---|
| 25,20881 | 25,91948 | 24,44916 | 24,62199 | 21,62828 | 24,39898 | 3 | 3 |
| 26,09432 | 26,70306 | 24,28602 | 24,32265 | 22,87637 | 24,34915 | 2 | 2 |
| 27,91359 | 27,58298 | 26,84482 | 23,21555 | 26,65431 | 20,82068 | 4 | 4 |
| 27,80974 | 27,4585 | 27,00496 | 27,03265 | 22,26025 | 26,88565 | 4 | 4 |
| 25,74042 | 25,78362 | 25,55789 | 24,69965 | 25,41738 | 22,86312 | 2 | 2 |
| 25,7067 | 25,96007 | 25,68493 | 25,06209 | 21,43758 | 25,31302 | 2 | 2 |
| 25,27245 | 25,58523 | 24,27611 | 24,49363 | 23,96473 | 21,1557 | 3 | 3 |
| 25,86832 | 25,09744 | 25,26327 | 24,50167 | 25,16212 | 22,61255 | 4 | 4 |
| 26,78319 | 26,87467 | 26,72936 | 26,12562 | 20,95833 | 26,17967 | 4 | 4 |
| 26,76636 | 26,11785 | 27,88198 | 26,08697 | 25,81358 | 25,35158 | 7 | 2 |
| 25,45224 | 25,45271 | 24,38331 | 21,76591 | 24,37391 | 22,4107 | 3 | 3 |
| 25,63544 | 26,14371 | 25,90786 | 24,95921 | 23,68688 | 25,25111 | 2 | 2 |
| 28,94179 | 30,4898 | 27,72501 | 21,93978 | 22,91123 | 28,24254 | 2 | 2 |
| 25,59243 | 25,57236 | 25,47587 | 24,84297 | 24,92533 | 19,60163 | 3 | 3 |
| 24,28835 | 22,98228 | 23,55262 | 22,29698 | 21,16 | 22,92796 | 2 | 2 |
| 28,28529 | 27,56114 | 27,46115 | 27,21429 | 26,76939 | 25,65253 | 8 | 8 |
| 34,19557 | 35,83404 | 27,35607 | 25,93852 | 35,68969 | 25,65736 | 2 | 2 |
| 25,87919 | 25,72497 | 25,39411 | 25,40204 | 20,21768 | 25,94523 | 3 | 3 |
| 25,8327 | 25,59275 | 25,92757 | 25,62221 | 25,72018 | 22,35675 | 2 | 2 |
| 25,84826 | 25,00423 | 25,62034 | 25,18162 | 22,20823 | 21,60293 | 3 | 3 |
| 25,25803 | 24,22575 | 25,33644 | 24,25494 | 21,87254 | 24,83229 | 2 | 2 |
| 28,30704 | 28,16404 | 28,05698 | 27,57694 | 27,50036 | 26,25303 | 12 | 12 |
| 27,46162 | 27,09376 | 28,02538 | 24,63334 | 27,45576 | 21,56042 | 3 | 3 |
| 29,4308 | 29,73127 | 29,48882 | 28,84395 | 27,31579 | 26,11718 | 2 | 2 |
| 24,40801 | 24,19182 | 24,09877 | 23,74448 | 24,14437 | 21,55126 | 2 | 2 |
| 25,65138 | 25,85015 | 23,4853 | 20,13144 | 23,25681 | 22,70428 | 3 | 3 |
| 25,10362 | 25,40406 | 24,97823 | 25,28747 | 21,93358 | 25,22181 | 3 | 3 |
| 24,88106 | 25,86313 | 26,20747 | 25,13864 | 21,3417 | 25,13794 | 4 | 4 |
| 25,60744 | 25,50996 | 24,83186 | 25,21982 | 21,44338 | 25,89169 | 2 | 2 |
| 27,56224 | 26,17488 | 26,82043 | 24,92084 | 25,12376 | 26,11353 | 12 | 12 |
| 24,7397 | 24,68148 | 23,57173 | 23,45224 | 22,66249 | 23,52325 | 2 | 2 |
| 26,51817 | 26,31098 | 26,00693 | 25,11617 | 25,62213 | 23,00183 | 3 | 3 |
| 26,56634 | 27,10959 | 25,00988 | 25,57323 | 24,35461 | 24,06019 | 2 | 2 |
| 26,0536 | 25,9819 | 26,31318 | 26,2637 | 25,87058 | 22,86771 | 6 | 6 |
| 27,38969 | 27,87033 | 28,43357 | 26,80858 | 27,37136 | 24,98941 | 8 | 8 |
| 25,64631 | 25,08148 | 23,02713 | 23,6231 | 21,71678 | 22,69342 | 2 | 2 |
| 23,77476 | 21,92635 | 23,85148 | 25,04055 | 24,50767 | 24,4188 | 4 | 4 |
| 23,28842 | 27,06301 | 27,21178 | 27,09204 | 27,90461 | 27,2141 | 4 | 4 |
| 21,42491 | 22,50776 | 24,625 | 24,37643 | 24,04494 | 24,60303 | 69 | 1 |
| 25,08712 | 21,66337 | 24,97369 | 25,81331 | 25,22784 | 26,02386 | 2 | 2 |
| 25,95296 | 24,76744 | 23,54663 | 26,25057 | 26,93528 | 25,0748 | 3 | 3 |
| 23,31354 | 23,57138 | 22,99552 | 26,83762 | 25,36626 | 23,83374 | 21 | 1 |
| 24,40795 | 24,50616 | 22,01313 | 24,75588 | 24,78806 | 24,82934 | 3 | 3 |
| 22,17411 | 25,62308 | 25,39891 | 24,24227 | 25,92515 | 26,88507 | 2 | 2 |
| 21,592 | 22,95976 | 28,09406 | 26,4885 | 26,4697 | 27,46669 | 2 | 2 |
| 21,65918 | 21,4707 | 23,73552 | 22,75904 | 23,51677 | 23,62399 | 2 | 2 |
| 22,46478 | 27,01009 | 23,19669 | 23,11938 | 28,7165 | 23,84173 | 2 | 2 |
| 24,50142 | 21,32339 | 22,27564 | 24,44278 | 25,08485 | 24,59889 | 3 | 3 |
| 25,85916 | 23,20245 | 19,62515 | 24,95123 | 26,81066 | 26,71908 | 3 | 3 |
| 22,87296 | 22,14205 | 24,63871 | 24,72448 | 24,53617 | 23,84833 | 5 | 4 |

| | | | | | | | |
|---|---|---|---|---|---|---|---|
| 24,47482 | 21,80369 | 20,94347 | 23,20619 | 25,05059 | 25,42294 | 2 | 2 |
| 21,83136 | 23,30299 | 21,63542 | 24,74873 | 21,46585 | 25,21727 | 2 | 2 |
| 23,27573 | 25,64158 | 25,7001 | 25,87713 | 25,7751 | 26,58606 | 3 | 3 |
| 21,23869 | 24,2717 | 23,88418 | 24,17425 | 24,33967 | 23,9785 | 2 | 2 |
| 21,21642 | 22,30708 | 22,67984 | 22,29788 | 22,6256 | 24,4513 | 13 | 1 |
| 24,4771 | 24,58612 | 21,55445 | 24,9693 | 25,51557 | 25,52459 | 3 | 3 |
| 22,04871 | 25,60459 | 25,63334 | 26,64801 | 25,89481 | 25,68067 | 2 | 2 |
| 21,88305 | 24,13985 | 24,71733 | 24,7752 | 25,32828 | 25,66392 | 2 | 2 |
| 22,23558 | 23,17455 | 20,80511 | 23,87305 | 23,7144 | 22,99281 | 2 | 2 |
| 21,59519 | 21,83592 | 19,66652 | 23,73376 | 21,99331 | 21,43369 | 2 | 2 |
| 21,75831 | 26,46208 | 23,23261 | 25,66096 | 26,3838 | 26,28759 | 2 | 2 |
| 26,47678 | 26,0167 | 25,33231 | 24,88003 | 25,2737 | 24,82866 | 3 | 3 |
| 26,23487 | 26,10041 | 25,88274 | 26,084 | 26,49835 | 26,29966 | 8 | 5 |
| 27,28613 | 27,21669 | 27,4178 | 27,0592 | 27,20695 | 27,52917 | 7 | 7 |
| 23,76405 | 23,98598 | 24,1404 | 24,90651 | 24,22192 | 21,87243 | 2 | 2 |
| 27,03612 | 26,87185 | 27,00228 | 26,85852 | 26,48501 | 27,25863 | 8 | 8 |
| 31,28444 | 31,35857 | 30,74142 | 31,58042 | 31,38684 | 31,35642 | 17 | 17 |
| 27,12701 | 27,14385 | 26,63644 | 26,95172 | 27,26266 | 27,12888 | 7 | 7 |
| 27,32337 | 28,10059 | 27,82347 | 27,73085 | 27,56674 | 27,5721 | 5 | 5 |
| 24,13891 | 23,58474 | 24,15491 | 24,32526 | 22,24345 | 24,3278 | 2 | 2 |
| 27,49396 | 27,64849 | 27,53022 | 27,44561 | 27,49953 | 27,04124 | 6 | 6 |
| 26,98196 | 27,01999 | 26,42817 | 27,0876 | 26,854 | 26,47673 | 5 | 5 |
| 29,30776 | 29,31404 | 28,87654 | 28,35464 | 29,30101 | 28,93058 | 7 | 7 |
| 26,16793 | 25,95821 | 25,79892 | 26,10713 | 25,90012 | 25,98253 | 4 | 4 |
| 24,46283 | 24,90228 | 24,65971 | 24,38146 | 24,55321 | 25,0721 | 2 | 2 |
| 25,39365 | 25,71707 | 25,68298 | 25,35209 | 25,97502 | 25,95009 | 4 | 4 |
| 25,74686 | 25,58317 | 25,91356 | 26,01801 | 25,58305 | 24,58337 | 2 | 2 |
| 29,04017 | 24,21052 | 29,31471 | 27,23013 | 26,91228 | 28,57788 | 2 | 2 |
| 27,35817 | 28,75318 | 28,53081 | 27,85258 | 28,13153 | 29,15424 | 9 | 9 |
| 26,46899 | 27,58628 | 27,05092 | 27,89574 | 27,49052 | 28,04651 | 8 | 8 |
| 25,26635 | 25,76063 | 24,68014 | 25,29632 | 23,89745 | 23,89551 | 2 | 2 |
| 27,26222 | 27,76148 | 27,18952 | 27,08111 | 27,308 | 27,30443 | 4 | 4 |
| 27,25836 | 26,97257 | 27,00174 | 26,87009 | 26,68392 | 27,09949 | 5 | 5 |
| 26,24543 | 25,73534 | 25,66131 | 25,54981 | 26,33552 | 25,94409 | 5 | 5 |
| 24,28912 | 25,11656 | 24,24598 | 24,56976 | 24,41738 | 23,80065 | 4 | 4 |
| 22,07267 | 22,47835 | 23,06189 | 22,6558 | 22,31866 | 23,09233 | 3 | 3 |
| 27,56049 | 27,24409 | 27,71017 | 27,41691 | 27,16442 | 27,5018 | 8 | 8 |
| 23,73159 | 24,84623 | 23,92452 | 24,97014 | 24,36359 | 21,70915 | 2 | 2 |
| 23,635 | 22,80906 | 23,06549 | 23,35192 | 23,89781 | 22,97518 | 3 | 3 |
| 27,68552 | 27,55764 | 27,45967 | 27,78967 | 27,88075 | 27,63187 | 11 | 11 |
| 27,91194 | 27,68258 | 27,69365 | 28,54156 | 27,84751 | 28,02099 | 7 | 7 |
| 27,06969 | 26,71908 | 26,74932 | 26,80637 | 26,73854 | 27,41078 | 8 | 8 |
| 28,26316 | 28,24004 | 27,81121 | 28,55512 | 28,21257 | 28,18315 | 10 | 10 |
| 25,6312 | 25,89977 | 25,9968 | 25,57303 | 25,84766 | 25,25137 | 4 | 2 |
| 32,15289 | 32,02759 | 31,69789 | 31,76161 | 32,04124 | 31,85556 | 12 | 10 |
| 26,47198 | 25,80538 | 25,89816 | 25,90157 | 25,7685 | 25,72559 | 4 | 4 |
| 22,27034 | 21,85243 | 22,07835 | 22,03295 | 21,60209 | 21,75979 | 2 | 2 |
| 28,10214 | 27,9856 | 27,52694 | 27,85436 | 27,68372 | 26,51572 | 7 | 7 |
| 28,23073 | 28,01009 | 28,24427 | 27,82408 | 27,41683 | 26,82857 | 9 | 9 |
| 25,97001 | 27,08456 | 26,85246 | 26,7107 | 26,31638 | 26,18499 | 9 | 9 |

| | | | | | | | |
|---|---|---|---|---|---|---|---|
| 25,36353 | 25,59511 | 22,0771 | 23,94798 | 25,86729 | 23,64405 | 7 | 7 |
| 26,74868 | 26,49052 | 26,58764 | 26,48622 | 26,43971 | 26,4222 | 4 | 4 |
| 28,50737 | 27,73937 | 28,3736 | 28,67108 | 29,71658 | 28,41635 | 3 | 3 |
| 25,01095 | 23,78835 | 23,69051 | 23,51701 | 23,30841 | 23,34449 | 4 | 4 |
| 30,39785 | 30,13638 | 30,37555 | 30,12176 | 30,35269 | 30,25935 | 17 | 17 |
| 28,52638 | 27,20023 | 28,43062 | 27,28074 | 27,93601 | 28,01919 | 6 | 6 |
| 24,95536 | 24,77595 | 25,18435 | 25,1512 | 24,47035 | 25,99608 | 4 | 4 |
| 24,38212 | 24,0878 | 24,72114 | 24,21089 | 23,98676 | 25,77062 | 4 | 4 |
| 25,98222 | 25,81675 | 25,5583 | 25,71295 | 25,84117 | 26,28553 | 4 | 4 |
| 28,63072 | 28,44055 | 28,81794 | 28,59149 | 29,20751 | 29,06193 | 5 | 5 |
| 25,43476 | 25,10949 | 24,58251 | 24,40594 | 24,30709 | 24,35448 | 3 | 3 |
| 23,15724 | 25,09071 | 25,16599 | 25,01822 | 24,16162 | 24,05556 | 3 | 3 |
| 25,56813 | 23,02051 | 23,8658 | 24,41931 | 23,99473 | 26,86821 | 3 | 3 |
| 30,17009 | 30,22331 | 29,87685 | 29,88469 | 29,60883 | 29,27267 | 12 | 12 |
| 31,93343 | 32,13816 | 32,00978 | 32,14136 | 31,91046 | 31,97555 | 47 | 47 |
| 34,37166 | 34,37477 | 34,18201 | 34,37484 | 34,0027 | 33,99951 | 8 | 6 |
| 31,16447 | 30,94014 | 30,46022 | 30,43946 | 30,81826 | 30,72297 | 14 | 14 |
| 21,41586 | 22,66169 | 23,79937 | 22,7633 | 23,26728 | 23,2548 | 2 | 2 |
| 26,6679 | 26,34159 | 26,9812 | 27,35574 | 26,82189 | 26,7375 | 8 | 8 |
| 26,82141 | 27,77593 | 27,18253 | 27,27355 | 27,38105 | 27,40438 | 6 | 6 |
| 21,28648 | 24,83016 | 24,64988 | 24,6084 | 21,24994 | 24,69275 | 3 | 3 |
| 29,73705 | 29,67212 | 29,49576 | 29,41355 | 29,30572 | 29,68161 | 13 | 3 |
| 32,27122 | 32,36346 | 31,87199 | 32,16135 | 31,65762 | 31,82373 | 7 | 7 |
| 29,25858 | 28,97265 | 29,08203 | 29,17653 | 28,86558 | 29,00582 | 15 | 15 |
| 26,41146 | 25,68942 | 25,9769 | 25,39679 | 25,10797 | 25,54901 | 2 | 2 |
| 29,91698 | 30,15774 | 30,03708 | 29,91085 | 30,1575 | 30,15025 | 18 | 18 |
| 28,12923 | 28,05734 | 27,83545 | 28,01404 | 28,65976 | 28,44892 | 4 | 4 |
| 29,59237 | 29,58287 | 29,21484 | 29,7128 | 29,72639 | 29,84734 | 16 | 16 |
| 25,04693 | 25,39257 | 24,52283 | 25,98111 | 24,99002 | 21,93858 | 3 | 3 |
| 24,42137 | 25,11494 | 24,93674 | 23,89069 | 24,25609 | 24,78332 | 5 | 5 |
| 31,73828 | 32,0364 | 30,80847 | 31,92405 | 31,86308 | 31,4068 | 10 | 10 |
| 23,18912 | 22,60851 | 24,11219 | 22,99222 | 22,77604 | 22,63969 | 2 | 2 |
| 27,69724 | 27,33833 | 27,32569 | 27,62708 | 27,84799 | 27,64258 | 6 | 6 |
| 26,41835 | 26,34554 | 26,59092 | 26,74227 | 26,43159 | 26,37353 | 4 | 4 |
| 22,81658 | 22,93023 | 22,8507 | 22,67098 | 22,45538 | 21,86645 | 11 | 1 |
| 25,57652 | 25,08217 | 25,54675 | 25,12569 | 25,18223 | 24,18117 | 5 | 5 |
| 29,22297 | 28,86747 | 27,97323 | 28,69495 | 29,65779 | 29,3878 | 12 | 12 |
| 29,04085 | 29,17995 | 29,20074 | 29,02986 | 29,11791 | 29,33371 | 18 | 18 |
| 23,59719 | 23,55169 | 23,66438 | 24,0535 | 22,90012 | 24,4188 | 2 | 2 |
| 26,48045 | 26,38334 | 26,65376 | 26,34315 | 25,29506 | 26,12037 | 2 | 2 |
| 26,31022 | 26,37592 | 26,37721 | 26,45419 | 26,31986 | 25,853 | 5 | 2 |
| 20,75316 | 24,59161 | 25,09148 | 23,16719 | 24,88012 | 24,9461 | 17 | 1 |
| 29,96719 | 30,02049 | 30,92451 | 30,00865 | 29,66851 | 30,85348 | 21 | 21 |
| 28,28754 | 28,80796 | 28,34448 | 28,67071 | 28,56194 | 28,41389 | 10 | 10 |
| 25,17976 | 26,19169 | 25,62414 | 26,28738 | 26,08383 | 26,44092 | 2 | 2 |
| 26,27654 | 27,01297 | 26,22735 | 26,82262 | 26,49111 | 26,46855 | 7 | 7 |
| 28,58534 | 28,50309 | 27,57932 | 28,26754 | 28,04593 | 28,31488 | 6 | 6 |
| 24,25832 | 24,54975 | 24,44493 | 22,43216 | 24,4688 | 24,39355 | 2 | 2 |
| 28,97861 | 28,68983 | 28,57174 | 29,09162 | 28,85641 | 28,43289 | 9 | 9 |
| 21,59336 | 22,63823 | 23,08755 | 22,42183 | 21,68855 | 20,9399 | 2 | 2 |

| | | | | | | | |
|---|---|---|---|---|---|---|---|
| 28,94849 | 28,93309 | 28,98921 | 28,6731 | 28,93103 | 28,66027 | 12 | 12 |
| 25,48597 | 25,54731 | 26,19924 | 26,09192 | 26,15646 | 26,23511 | 4 | 4 |
| 27,19188 | 27,16202 | 26,78481 | 26,51961 | 27,15393 | 26,04241 | 4 | 4 |
| 26,58835 | 26,90219 | 26,25335 | 26,61203 | 26,49225 | 26,85103 | 5 | 5 |
| 25,60781 | 25,98096 | 26,35572 | 25,91214 | 25,78187 | 25,96084 | 6 | 6 |
| 26,19088 | 25,94559 | 25,86712 | 26,07763 | 25,63517 | 25,16603 | 4 | 4 |
| 27,39877 | 27,56027 | 27,18698 | 27,19357 | 27,31242 | 26,46011 | 5 | 5 |
| 24,51503 | 24,05805 | 24,31734 | 24,06201 | 23,93822 | 24,01265 | 3 | 3 |
| 26,5242 | 27,00174 | 26,94089 | 27,779 | 26,96917 | 26,97169 | 8 | 8 |
| 28,47646 | 28,29649 | 27,88489 | 28,25714 | 28,48608 | 28,88658 | 3 | 3 |
| 24,06669 | 24,5044 | 24,07798 | 24,32958 | 24,33163 | 24,24779 | 3 | 3 |
| 25,16928 | 24,94561 | 25,43619 | 25,31631 | 24,98958 | 25,33981 | 2 | 1 |
| 25,88008 | 25,84771 | 25,84795 | 25,85481 | 25,69962 | 25,956 | 3 | 3 |
| 25,20929 | 24,83152 | 23,39008 | 25,71453 | 25,35421 | 24,31119 | 7 | 7 |
| 25,16534 | 25,38245 | 24,9664 | 25,33616 | 25,10318 | 25,07606 | 2 | 2 |
| 25,40077 | 25,44878 | 25,7583 | 25,51141 | 25,58689 | 25,24818 | 4 | 4 |
| 26,23308 | 26,32869 | 26,09485 | 26,43951 | 26,79971 | 26,47795 | 5 | 5 |
| 26,34494 | 26,24784 | 26,59605 | 26,43793 | 26,47291 | 26,34652 | 9 | 9 |
| 24,14429 | 23,92778 | 23,18971 | 23,51629 | 23,58233 | 23,0034 | 3 | 3 |
| 23,44127 | 22,2727 | 24,00196 | 23,78247 | 22,6231 | 24,21556 | 2 | 2 |
| 26,81103 | 26,22927 | 26,63312 | 26,39447 | 26,31603 | 25,98609 | 5 | 5 |
| 24,50094 | 25,11088 | 25,93238 | 25,14507 | 26,34089 | 24,85667 | 2 | 2 |
| 27,14336 | 26,25472 | 26,47816 | 26,83147 | 26,22008 | 26,81934 | 8 | 8 |
| 23,42008 | 22,74043 | 21,69908 | 22,80581 | 22,21745 | 22,57725 | 11 | 1 |
| 25,16921 | 25,40876 | 25,21349 | 25,00457 | 25,99481 | 24,76795 | 2 | 2 |
| 26,76028 | 25,74193 | 25,09973 | 26,14577 | 26,44985 | 25,98592 | 2 | 2 |
| 25,79566 | 25,76134 | 25,37554 | 25,58018 | 26,59719 | 26,42824 | 4 | 4 |
| 24,65414 | 25,13046 | 25,11438 | 24,89296 | 25,2793 | 25,24322 | 3 | 3 |
| 28,09607 | 27,52477 | 28,04999 | 27,78463 | 28,1211 | 27,65956 | 3 | 3 |
| 26,88972 | 26,82055 | 27,18026 | 27,05341 | 26,60102 | 26,79922 | 4 | 4 |
| 26,07071 | 25,25807 | 25,9666 | 26,34399 | 26,2719 | 26,0879 | 4 | 4 |
| 31,06001 | 30,66949 | 30,88707 | 30,53612 | 30,66399 | 30,50266 | 24 | 16 |
| 28,58946 | 28,85171 | 28,811 | 28,46673 | 28,73473 | 28,56903 | 13 | 1 |
| 28,79182 | 28,55149 | 28,39468 | 28,67912 | 28,44853 | 28,6412 | 10 | 10 |
| 25,96763 | 25,03859 | 25,6373 | 25,95894 | 25,67188 | 25,26792 | 3 | 3 |
| 28,35149 | 28,13109 | 27,53341 | 28,38566 | 28,15166 | 27,99523 | 8 | 6 |
| 25,41445 | 24,9193 | 25,57199 | 25,99677 | 26,23004 | 25,16982 | 13 | 4 |
| 23,53908 | 23,97281 | 23,79818 | 22,77901 | 23,92171 | 24,00067 | 2 | 2 |
| 27,82019 | 27,79619 | 27,32852 | 27,37609 | 27,32886 | 27,29754 | 8 | 8 |
| 28,61807 | 28,37588 | 28,45442 | 29,04195 | 28,9028 | 28,59053 | 17 | 17 |
| 26,81849 | 26,41732 | 26,20301 | 26,32017 | 26,38217 | 26,48508 | 3 | 3 |
| 25,25548 | 24,41635 | 24,59616 | 24,97189 | 23,46089 | 24,17333 | 6 | 6 |
| 24,39225 | 24,296 | 24,10701 | 23,80627 | 24,67912 | 24,70033 | 16 | 1 |
| 26,78057 | 27,41893 | 26,89203 | 27,17217 | 26,51932 | 27,10719 | 6 | 6 |
| 27,03118 | 27,23763 | 27,26392 | 27,28657 | 27,42575 | 27,24799 | 7 | 7 |
| 32,2144 | 32,3556 | 32,17126 | 31,98118 | 32,1767 | 31,75871 | 12 | 12 |
| 23,04833 | 22,61495 | 21,36196 | 21,29019 | 19,50297 | 23,56999 | 2 | 2 |
| 25,86825 | 26,35968 | 26,03299 | 26,01263 | 25,91454 | 25,7077 | 2 | 2 |
| 25,44695 | 25,3803 | 25,64334 | 25,63425 | 25,5357 | 25,71759 | 3 | 3 |

| | | | | | | | |
|---|---|---|---|---|---|---|---|
| 32,51559 | 32,52843 | 32,44287 | 32,57997 | 32,60399 | 32,47259 | 113 | 65 |
| 25,89141 | 25,97474 | 25,88758 | 25,80257 | 25,78579 | 26,14872 | 10 | 10 |
| 25,69509 | 25,41232 | 25,42124 | 25,24282 | 24,91953 | 25,47797 | 8 | 8 |
| 24,81416 | 25,56069 | 25,68143 | 26,69856 | 25,66251 | 26,11245 | 8 | 8 |
| 30,22527 | 30,17284 | 29,67003 | 30,61193 | 30,53519 | 30,40882 | 17 | 17 |
| 25,84984 | 25,33906 | 25,35864 | 25,20337 | 26,14106 | 25,64268 | 2 | 2 |
| 29,30138 | 29,22267 | 28,86154 | 29,2469 | 29,12401 | 29,01606 | 11 | 11 |
| 27,9198 | 27,56841 | 27,67054 | 27,91285 | 27,30783 | 27,93988 | 13 | 2 |
| 27,07439 | 27,0475 | 26,65239 | 27,0626 | 26,82941 | 27,21132 | 8 | 2 |
| 26,56342 | 25,74386 | 26,67506 | 27,27409 | 26,77191 | 26,84566 | 6 | 6 |
| 25,29691 | 26,30598 | 26,59634 | 25,16917 | 24,89176 | 25,91022 | 4 | 4 |
| 28,81659 | 28,84867 | 28,90673 | 28,57982 | 28,933 | 28,82545 | 13 | 4 |
| 24,19558 | 23,76152 | 25,01606 | 24,50978 | 24,81279 | 24,00607 | 3 | 3 |
| 25,30281 | 25,46469 | 25,09977 | 25,07826 | 25,24496 | 25,17725 | 4 | 4 |
| 25,44234 | 24,84096 | 25,2792 | 25,52552 | 25,30295 | 24,90775 | 3 | 3 |
| 30,91006 | 30,58755 | 30,59646 | 30,2457 | 30,55883 | 30,15967 | 15 | 15 |
| 25,61667 | 24,63594 | 24,9345 | 25,2488 | 26,07938 | 25,34787 | 6 | 6 |
| 28,07643 | 28,3385 | 28,57546 | 28,36457 | 28,39284 | 28,27954 | 8 | 3 |
| 30,32249 | 30,65241 | 29,92606 | 30,64538 | 30,53492 | 30,55746 | 11 | 11 |
| 25,65322 | 25,81123 | 25,39754 | 26,55141 | 25,11454 | 25,81414 | 2 | 2 |
| 24,78905 | 25,13798 | 25,04206 | 25,1712 | 25,31513 | 22,51862 | 2 | 2 |
| 29,28516 | 29,10559 | 28,96268 | 29,04913 | 28,54182 | 28,11639 | 6 | 6 |
| 26,76875 | 25,93692 | 25,93818 | 26,95116 | 26,46242 | 27,17741 | 6 | 6 |
| 25,23114 | 24,61941 | 24,1988 | 22,48636 | 25,06118 | 26,04514 | 2 | 2 |
| 30,18825 | 29,90383 | 29,80664 | 30,03486 | 29,91399 | 29,86211 | 15 | 15 |
| 25,9574 | 25,51431 | 25,52674 | 25,11858 | 25,20471 | 25,40067 | 2 | 2 |
| 24,77269 | 24,32876 | 24,11514 | 24,55882 | 24,01299 | 24,03372 | 2 | 2 |
| 30,06433 | 29,93495 | 30,10699 | 30,03092 | 30,19917 | 30,00531 | 20 | 20 |
| 31,69627 | 32,17902 | 31,83624 | 31,46684 | 31,41468 | 31,6571 | 62 | 62 |
| 27,41555 | 26,84948 | 27,45482 | 26,97378 | 27,18338 | 27,35347 | 3 | 3 |
| 27,25178 | 27,22582 | 26,8411 | 26,84613 | 26,50859 | 26,34873 | 3 | 3 |
| 27,43428 | 27,52552 | 27,2615 | 27,49244 | 27,11555 | 26,86597 | 4 | 4 |
| 29,81572 | 29,63822 | 29,23171 | 29,79535 | 30,04114 | 29,71807 | 13 | 13 |
| 27,34189 | 27,4334 | 27,64993 | 27,13516 | 26,96653 | 27,73176 | 6 | 6 |
| 24,85776 | 25,01419 | 24,47265 | 24,84891 | 24,96671 | 25,18696 | 2 | 2 |
| 24,24627 | 24,62282 | 24,38824 | 24,30835 | 24,09748 | 24,19535 | 2 | 2 |
| 27,92162 | 27,61302 | 27,86609 | 27,65403 | 27,73841 | 27,84032 | 6 | 6 |
| 26,5302 | 26,66899 | 26,49852 | 26,5773 | 26,05588 | 26,16246 | 5 | 5 |
| 27,23023 | 27,27062 | 27,04833 | 27,84368 | 27,41675 | 27,28118 | 8 | 8 |
| 29,27573 | 29,15228 | 28,77354 | 29,14603 | 29,42484 | 29,40574 | 7 | 7 |
| 25,73024 | 25,91461 | 23,55667 | 25,99583 | 25,36966 | 21,73591 | 6 | 1 |
| 23,84192 | 24,58187 | 22,13064 | 24,58055 | 23,54898 | 22,99372 | 2 | 2 |
| 28,18726 | 27,98951 | 27,57564 | 27,34748 | 27,58133 | 27,28762 | 7 | 7 |
| 35,66792 | 35,46815 | 35,07507 | 34,9801 | 35,16448 | 34,97852 | 18 | 18 |
| 28,07265 | 28,17231 | 28,34181 | 28,18504 | 28,46451 | 28,06579 | 4 | 4 |
| 36,34517 | 36,19735 | 36,06712 | 36,21766 | 36,27708 | 36,25318 | 56 | 47 |
| 24,02284 | 24,4562 | 24,45613 | 24,26155 | 23,17776 | 22,91183 | 2 | 2 |
| 28,46209 | 28,68939 | 28,38722 | 28,57499 | 28,47112 | 28,28745 | 13 | 7 |
| 28,09818 | 28,00603 | 28,15253 | 28,19662 | 28,1856 | 28,3986 | 8 | 8 |
| 30,99598 | 30,83425 | 30,37078 | 30,50323 | 30,40355 | 30,45944 | 23 | 23 |

| | | | | | | | |
|---|---|---|---|---|---|---|---|
| 24,51959 | 24,26234 | 24,00222 | 24,38765 | 23,98372 | 24,51208 | 2 | 2 |
| 26,76648 | 26,37127 | 26,13208 | 26,44542 | 26,6764 | 26,12501 | 5 | 5 |
| 26,34391 | 26,86455 | 26,37433 | 26,77065 | 26,71293 | 26,23996 | 3 | 3 |
| 27,37625 | 26,768 | 26,67506 | 26,89308 | 26,54934 | 26,54719 | 6 | 6 |
| 25,63741 | 25,51806 | 25,74929 | 25,31648 | 24,61913 | 25,51776 | 3 | 3 |
| 23,38535 | 23,85795 | 23,47605 | 23,87014 | 23,46351 | 23,54534 | 2 | 2 |
| 28,25273 | 27,74529 | 27,96047 | 27,06209 | 27,53022 | 27,62101 | 9 | 9 |
| 29,51305 | 29,19974 | 28,91199 | 28,65421 | 29,01308 | 28,796 | 9 | 9 |
| 26,85674 | 26,86432 | 25,72427 | 26,43605 | 26,27131 | 26,1278 | 5 | 5 |
| 30,72558 | 30,32667 | 30,0927 | 30,71792 | 30,69644 | 30,60346 | 9 | 4 |
| 26,09073 | 26,57255 | 26,05788 | 26,37619 | 26,08762 | 25,74553 | 5 | 5 |
| 29,43187 | 29,41784 | 29,10842 | 29,06584 | 29,20658 | 29,12635 | 8 | 8 |
| 25,47957 | 25,52666 | 22,9186 | 25,31413 | 24,96596 | 25,59383 | 2 | 2 |
| 28,33641 | 28,47781 | 27,91809 | 28,02464 | 28,08284 | 28,09053 | 5 | 5 |
| 28,53586 | 28,82542 | 28,38825 | 28,51631 | 28,07806 | 28,36628 | 8 | 8 |
| 26,26612 | 25,84881 | 26,43649 | 26,8839 | 25,93986 | 23,44847 | 3 | 3 |
| 28,06769 | 28,40129 | 28,08121 | 28,32877 | 28,14667 | 27,95316 | 11 | 2 |
| 24,10909 | 25,78154 | 24,86283 | 26,08241 | 25,65567 | 25,5823 | 3 | 3 |
| 24,93692 | 25,25285 | 24,05474 | 25,80048 | 24,95234 | 21,49412 | 3 | 3 |
| 27,01977 | 26,5041 | 26,35881 | 26,56792 | 26,11301 | 26,1207 | 2 | 2 |
| 28,04192 | 27,52649 | 27,98397 | 27,95682 | 28,08071 | 28,12549 | 10 | 10 |
| 23,98659 | 22,71157 | 21,60058 | 23,3002 | 23,1798 | 23,31976 | 2 | 2 |
| 29,62286 | 29,62178 | 29,35483 | 29,37805 | 29,14766 | 29,16504 | 5 | 4 |
| 28,66268 | 28,32423 | 28,42894 | 28,40141 | 28,19938 | 28,47591 | 12 | 2 |
| 26,28139 | 25,91265 | 26,21443 | 26,01272 | 26,02437 | 25,81069 | 4 | 4 |
| 25,47643 | 25,52373 | 25,75251 | 25,50443 | 25,02221 | 25,5581 | 4 | 4 |
| 28,28065 | 27,0703 | 28,29413 | 29,01316 | 28,91016 | 28,58302 | 2 | 2 |
| 28,24109 | 28,16332 | 28,31752 | 27,99459 | 28,08851 | 28,23137 | 10 | 1 |
| 25,14802 | 25,07611 | 26,24102 | 25,28641 | 24,37013 | 25,6091 | 3 | 3 |
| 25,98839 | 25,79 | 26,40472 | 26,6238 | 26,10528 | 26,51694 | 4 | 4 |
| 22,75873 | 22,34611 | 23,31741 | 21,52536 | 22,99943 | 23,39506 | 2 | 2 |
| 27,85436 | 28,51107 | 28,30926 | 27,88075 | 27,93966 | 27,3157 | 2 | 2 |
| 29,11808 | 29,36799 | 28,84613 | 28,9405 | 29,05359 | 28,77662 | 14 | 14 |
| 24,7291 | 25,81477 | 24,21386 | 24,22435 | 23,9722 | 23,59457 | 3 | 1 |
| 28,98688 | 29,02261 | 29,24043 | 28,85611 | 28,73728 | 28,97221 | 20 | 20 |
| 27,9961 | 28,21354 | 27,64382 | 28,35712 | 28,44217 | 27,80206 | 7 | 2 |
| 22,54182 | 22,8395 | 22,83225 | 23,08618 | 22,93225 | 23,29025 | 7 | 2 |
| 25,6384 | 26,5491 | 25,63896 | 25,40724 | 26,10977 | 26,08612 | 2 | 2 |
| 27,85644 | 27,65581 | 27,85389 | 28,01728 | 27,79891 | 28,11763 | 15 | 15 |
| 28,95222 | 29,29572 | 29,40131 | 29,08512 | 29,15349 | 29,22138 | 13 | 13 |
| 32,35253 | 32,51606 | 32,48513 | 32,28389 | 32,39051 | 32,39397 | 36 | 36 |
| 32,06359 | 32,11759 | 32,05012 | 31,97326 | 32,0753 | 32,0783 | 29 | 29 |
| 30,67287 | 29,7808 | 30,95387 | 30,88744 | 30,93965 | 30,56593 | 6 | 6 |
| 29,15161 | 28,07913 | 28,3333 | 29,27697 | 29,26542 | 29,9621 | 12 | 12 |
| 30,06522 | 29,94447 | 30,34043 | 30,53297 | 30,17248 | 29,89322 | 4 | 4 |
| 28,7662 | 28,85549 | 28,62725 | 28,57283 | 27,97635 | 28,68201 | 5 | 5 |
| 21,79448 | 23,35797 | 23,43797 | 23,87221 | 23,86467 | 23,62176 | 2 | 2 |
| 28,21169 | 27,83744 | 28,05502 | 28,12514 | 28,1052 | 27,93072 | 10 | 10 |
| 31,66941 | 31,81757 | 31,81176 | 31,78425 | 31,81639 | 31,84176 | 30 | 1 |
| 22,26227 | 23,20453 | 23,17782 | 23,26871 | 22,95987 | 24,03767 | 30 | 1 |

| | | | | | | | |
|---|---|---|---|---|---|---|---|
| 30,82138 | 30,53733 | 30,79118 | 31,06798 | 30,8282 | 31,02828 | 25 | 25 |
| 28,71696 | 28,63104 | 28,51555 | 28,58967 | 28,8546 | 28,64904 | 14 | 14 |
| 27,32766 | 27,45246 | 26,99755 | 27,12879 | 27,49831 | 27,26042 | 5 | 5 |
| 26,9788 | 25,94537 | 25,63492 | 25,01712 | 25,21456 | 25,66701 | 6 | 6 |
| 30,15158 | 29,99321 | 29,57384 | 28,73725 | 29,71547 | 29,33811 | 4 | 4 |
| 27,35699 | 27,5847 | 27,4649 | 27,37169 | 26,52187 | 25,88418 | 12 | 8 |
| 26,03269 | 26,51488 | 25,87124 | 25,46033 | 25,83067 | 25,3691 | 8 | 3 |
| 26,23277 | 26,40329 | 26,16421 | 26,17146 | 26,42544 | 26,08543 | 3 | 3 |
| 29,43131 | 29,34773 | 28,87182 | 28,9656 | 28,92102 | 28,8788 | 11 | 11 |
| 25,07879 | 24,61088 | 24,88455 | 25,4108 | 24,92561 | 24,84551 | 11 | 1 |
| 23,73635 | 24,37338 | 23,94262 | 24,69546 | 24,26399 | 24,46401 | 2 | 2 |
| 24,86618 | 23,39807 | 23,68955 | 24,68201 | 24,43517 | 24,06159 | 3 | 3 |
| 25,74558 | 25,78162 | 25,97237 | 25,95271 | 26,03641 | 26,22929 | 6 | 6 |
| 23,03246 | 23,32841 | 23,20611 | 22,45686 | 22,43797 | 23,39205 | 2 | 2 |
| 27,31553 | 26,9173 | 27,01914 | 27,31164 | 27,90828 | 26,43756 | 8 | 8 |
| 24,98733 | 25,26177 | 25,25158 | 25,29337 | 25,08966 | 24,81748 | 4 | 3 |
| 27,69319 | 27,26526 | 26,98739 | 26,76989 | 26,83966 | 26,58907 | 5 | 5 |
| 28,60784 | 28,11491 | 27,49762 | 27,33824 | 28,06866 | 27,78625 | 7 | 7 |
| 26,19927 | 25,47475 | 25,84206 | 25,71062 | 25,44 | 25,81892 | 4 | 4 |
| 26,1433 | 26,3936 | 26,69936 | 26,31136 | 26,62715 | 25,63159 | 3 | 3 |
| 28,17522 | 27,97618 | 28,30125 | 28,27111 | 27,57456 | 28,01993 | 14 | 14 |
| 24,92976 | 24,83089 | 24,63539 | 24,6207 | 24,51791 | 24,43644 | 2 | 2 |
| 28,42858 | 28,13075 | 28,68535 | 28,23197 | 28,14778 | 27,85043 | 9 | 9 |
| 25,80442 | 25,61099 | 25,52037 | 25,76188 | 26,24927 | 25,80132 | 4 | 4 |
| 30,07674 | 30,07202 | 29,48178 | 29,68001 | 29,99091 | 29,91869 | 11 | 11 |
| 28,30391 | 27,71398 | 27,88402 | 27,96653 | 27,82353 | 27,98387 | 7 | 7 |
| 25,29869 | 23,30091 | 25,75496 | 25,09019 | 24,91479 | 22,00207 | 38 | 3 |
| 25,8959 | 25,8158 | 25,33125 | 25,89266 | 25,58397 | 25,9616 | 5 | 5 |
| 27,08689 | 27,19723 | 25,92893 | 27,14268 | 27,06147 | 27,22048 | 4 | 4 |
| 26,23352 | 27,65123 | 27,03507 | 26,42182 | 26,75073 | 26,49797 | 5 | 5 |
| 26,41843 | 26,37309 | 26,62909 | 26,14163 | 26,79971 | 26,49479 | 3 | 3 |
| 26,41164 | 25,12798 | 25,22762 | 26,13929 | 25,93686 | 26,19764 | 4 | 4 |
| 27,66438 | 27,61435 | 27,80532 | 28,0704 | 26,71856 | 27,93106 | 8 | 8 |
| 28,83879 | 29,3678 | 29,0713 | 29,07301 | 28,74298 | 29,1938 | 12 | 12 |
| 25,52984 | 25,87065 | 25,45579 | 25,32488 | 25,33565 | 25,41438 | 4 | 4 |
| 27,45293 | 27,29255 | 27,24146 | 27,22057 | 27,18281 | 27,28277 | 7 | 7 |
| 30,87291 | 30,89461 | 31,22682 | 30,69511 | 30,79715 | 30,91706 | 26 | 26 |
| 23,99688 | 23,66557 | 23,90573 | 23,32251 | 23,76243 | 23,06017 | 2 | 2 |
| 27,84924 | 27,65082 | 27,54392 | 27,4567 | 27,77562 | 27,90426 | 10 | 10 |
| 28,89675 | 29,18577 | 28,67496 | 28,96199 | 28,73857 | 28,2611 | 3 | 3 |
| 28,55344 | 28,28965 | 28,253 | 28,65857 | 28,54315 | 28,33151 | 9 | 9 |
| 27,8642 | 27,63796 | 27,43881 | 27,58154 | 27,70002 | 27,70207 | 8 | 2 |
| 25,20579 | 24,99261 | 25,34374 | 25,3946 | 25,05316 | 25,14915 | 4 | 4 |
| 25,72606 | 25,53715 | 26,05736 | 26,15184 | 26,04589 | 25,5786 | 4 | 4 |
| 29,58384 | 29,87379 | 29,84939 | 29,59673 | 29,70464 | 29,77651 | 24 | 24 |
| 25,47954 | 25,05163 | 22,38175 | 22,62169 | 25,84525 | 25,21564 | 2 | 2 |
| 29,71767 | 29,29007 | 29,34634 | 29,35825 | 29,00161 | 29,10442 | 14 | 14 |
| 31,94517 | 31,82514 | 32,22026 | 31,89833 | 32,00025 | 32,38154 | 89 | 89 |
| 25,32598 | 25,64111 | 25,53303 | 26,15362 | 25,82167 | 25,66273 | 2 | 2 |
| 27,47568 | 26,86302 | 26,74407 | 27,4303 | 27,33083 | 27,01829 | 8 | 8 |

| | | | | | | | |
|---|---|---|---|---|---|---|---|
| 34,48412 | 34,42711 | 34,46927 | 34,32133 | 34,45544 | 34,34229 | 142 | 142 |
| 26,53396 | 26,6147 | 26,63229 | 25,80843 | 26,45678 | 26,16538 | 7 | 5 |
| 24,83918 | 24,68046 | 24,94078 | 25,44357 | 24,69349 | 24,7145 | 3 | 3 |
| 34,1284 | 33,77313 | 33,85731 | 33,81993 | 33,96206 | 33,9728 | 48 | 48 |
| 26,1813 | 25,88143 | 24,98407 | 26,44255 | 25,83531 | 25,70221 | 8 | 8 |
| 26,42762 | 25,66229 | 25,89571 | 25,37352 | 25,70506 | 25,12321 | 3 | 3 |
| 26,17805 | 26,46663 | 26,93157 | 24,66855 | 24,70993 | 27,48654 | 2 | 2 |
| 26,53228 | 26,78007 | 26,3226 | 26,70279 | 26,44482 | 27,21401 | 5 | 5 |
| 31,25715 | 31,05323 | 30,89266 | 30,66535 | 30,73788 | 30,86644 | 10 | 10 |
| 26,408 | 26,0701 | 25,93921 | 26,34937 | 25,80233 | 25,8584 | 5 | 5 |
| 26,20391 | 26,47598 | 26,54506 | 26,35213 | 26,58764 | 26,05122 | 8 | 8 |
| 26,09569 | 25,76979 | 26,23424 | 25,98702 | 26,25607 | 26,2903 | 5 | 5 |
| 27,67465 | 27,27799 | 27,09668 | 27,75385 | 27,41191 | 26,55289 | 4 | 4 |
| 26,74343 | 27,1211 | 26,41743 | 26,28701 | 26,62087 | 26,43898 | 2 | 2 |
| 28,0462 | 27,72832 | 27,55281 | 27,41893 | 27,50559 | 27,33577 | 6 | 6 |
| 26,9955 | 26,92536 | 26,54062 | 27,3827 | 26,9391 | 26,95082 | 3 | 3 |
| 25,49628 | 25,22059 | 24,98451 | 25,18223 | 25,16465 | 24,8892 | 3 | 3 |
| 25,51758 | 25,3979 | 23,98885 | 23,53231 | 23,70271 | 25,52295 | 3 | 3 |
| 28,83735 | 28,58714 | 28,25903 | 28,23444 | 28,45073 | 28,31687 | 8 | 8 |
| 27,62687 | 27,80894 | 27,20732 | 27,43054 | 27,54215 | 27,65007 | 13 | 13 |
| 26,9149 | 26,24773 | 27,0663 | 26,92739 | 27,18442 | 27,25187 | 11 | 11 |
| 26,41775 | 26,08472 | 27,00539 | 25,46354 | 26,6327 | 25,74878 | 2 | 2 |
| 28,98245 | 28,60424 | 29,31376 | 29,35609 | 29,34951 | 29,36133 | 8 | 8 |
| 31,56184 | 30,90182 | 30,33266 | 31,7531 | 31,35531 | 31,20967 | 7 | 1 |
| 26,45937 | 25,58081 | 27,32217 | 25,95414 | 26,50547 | 26,53687 | 7 | 7 |
| 28,73321 | 28,66316 | 28,91134 | 28,45423 | 28,49857 | 28,84933 | 7 | 7 |
| 29,08626 | 29,04265 | 28,91314 | 28,86859 | 28,97432 | 29,12115 | 10 | 10 |
| 28,14778 | 27,72123 | 27,82857 | 27,87812 | 28,12893 | 27,71024 | 8 | 8 |
| 34,12078 | 33,88831 | 33,96619 | 33,80569 | 33,80559 | 33,90311 | 116 | 4 |
| 28,7337 | 28,54885 | 28,26226 | 28,4271 | 28,61439 | 28,15962 | 11 | 11 |
| 26,32548 | 26,70306 | 26,64375 | 26,48886 | 26,2163 | 26,49753 | 4 | 4 |
| 26,44579 | 26,00661 | 25,45234 | 26,603 | 25,69633 | 25,60394 | 2 | 2 |
| 24,35104 | 23,30925 | 24,12731 | 23,59991 | 24,60286 | 25,36403 | 3 | 3 |
| 33,16286 | 32,58931 | 32,96222 | 32,57425 | 33,16722 | 32,74547 | 10 | 10 |
| 27,66628 | 27,61583 | 27,19366 | 27,69995 | 27,5852 | 26,84482 | 8 | 8 |
| 24,37073 | 25,73663 | 24,19167 | 24,99183 | 25,95753 | 26,21753 | 4 | 4 |
| 20,30419 | 21,26052 | 22,50913 | 21,89682 | 21,05232 | 19,72444 | 2 | 2 |
| 29,82467 | 29,35756 | 29,50173 | 29,45315 | 28,81955 | 29,14063 | 12 | 12 |
| 27,52589 | 27,16749 | 27,07521 | 27,27746 | 27,55098 | 27,65854 | 5 | 5 |
| 28,31069 | 28,17741 | 26,98707 | 27,3441 | 26,56526 | 27,14248 | 4 | 4 |
| 25,51638 | 25,34746 | 26,37444 | 25,66611 | 25,57894 | 26,3199 | 6 | 6 |
| 28,25201 | 28,10724 | 27,47576 | 27,58534 | 27,2129 | 27,1045 | 7 | 7 |
| 24,23564 | 24,39401 | 24,85024 | 24,51779 | 24,31036 | 24,01801 | 3 | 3 |
| 25,85863 | 26,05194 | 26,09188 | 25,59135 | 26,08517 | 25,85329 | 5 | 5 |
| 27,80224 | 26,94704 | 27,38039 | 27,01839 | 26,84876 | 26,9486 | 4 | 4 |
| 23,90802 | 24,2322 | 24,14965 | 22,54538 | 24,47976 | 24,18056 | 2 | 2 |
| 30,02393 | 30,53399 | 30,00892 | 29,87135 | 29,67665 | 30,10137 | 9 | 9 |
| 30,9698 | 30,95151 | 30,29527 | 30,08499 | 30,70849 | 30,6513 | 9 | 8 |
| 28,47676 | 28,62377 | 28,01164 | 28,28176 | 28,6526 | 28,90147 | 9 | 9 |
| 26,11527 | 26,72481 | 26,31983 | 26,75009 | 25,39963 | 26,01546 | 5 | 4 |

| | | | | | | | |
|---|---|---|---|---|---|---|---|
| 23,77193 | 23,5657 | 23,64328 | 24,99084 | 23,56639 | 23,88054 | 3 | 3 |
| 27,77228 | 27,91678 | 27,27088 | 27,98408 | 28,01201 | 27,86455 | 11 | 10 |
| 24,62355 | 24,80277 | 25,47277 | 24,73908 | 24,69019 | 24,67428 | 3 | 3 |
| 29,19373 | 29,00945 | 29,41474 | 29,21503 | 29,4833 | 29,24136 | 16 | 16 |
| 29,47756 | 29,20053 | 28,51912 | 29,2891 | 29,8746 | 29,23398 | 12 | 12 |
| 26,04783 | 25,4929 | 25,92073 | 25,10242 | 26,21666 | 25,54831 | 4 | 4 |
| 30,80793 | 30,76351 | 30,25293 | 30,29625 | 30,37803 | 30,50436 | 17 | 17 |
| 29,93972 | 29,93354 | 29,86952 | 29,82554 | 29,85938 | 30,00504 | 15 | 15 |
| 28,99664 | 28,75325 | 28,62641 | 28,88093 | 28,96441 | 28,91593 | 13 | 13 |
| 28,95391 | 28,49106 | 28,87771 | 29,31129 | 28,67805 | 28,78971 | 14 | 14 |
| 24,47296 | 24,74217 | 23,53231 | 24,0053 | 23,47085 | 23,49167 | 2 | 2 |
| 25,52522 | 25,79196 | 26,00753 | 25,76215 | 25,60834 | 25,14752 | 3 | 3 |
| 24,55028 | 24,51905 | 24,12195 | 24,47035 | 24,13469 | 25,35252 | 2 | 2 |
| 26,33829 | 26,73712 | 25,89768 | 26,58477 | 25,94318 | 24,51473 | 3 | 3 |
| 28,26723 | 28,35468 | 27,94553 | 27,98027 | 28,41038 | 27,74541 | 5 | 5 |
| 23,91059 | 23,48358 | 23,56918 | 24,71576 | 21,82663 | 23,58566 | 8 | 2 |
| 26,54113 | 26,60865 | 25,96455 | 26,63256 | 26,26367 | 26,14961 | 2 | 2 |
| 29,01773 | 28,97345 | 29,15661 | 28,63917 | 28,94946 | 28,70586 | 13 | 13 |
| 24,39702 | 24,2183 | 24,49692 | 24,24023 | 24,36259 | 24,29754 | 3 | 3 |
| 25,67885 | 26,10071 | 26,1752 | 25,5528 | 26,31608 | 25,88862 | 4 | 4 |
| 27,73821 | 27,71371 | 27,32105 | 27,48108 | 27,44403 | 26,83485 | 5 | 5 |
| 24,49161 | 22,32094 | 24,34618 | 24,79293 | 24,38469 | 22,66166 | 4 | 4 |
| 25,99835 | 25,05503 | 25,22916 | 24,92868 | 24,9354 | 24,93939 | 3 | 3 |
| 27,59953 | 27,84272 | 28,02496 | 27,67929 | 27,48547 | 27,47406 | 3 | 3 |
| 24,86032 | 24,61257 | 24,51833 | 24,48308 | 24,13938 | 24,50422 | 3 | 3 |
| 27,30365 | 27,20872 | 27,36803 | 27,30887 | 27,71109 | 27,37343 | 6 | 6 |
| 24,94253 | 25,71691 | 24,88488 | 25,67689 | 25,95409 | 25,71411 | 3 | 3 |
| 33,07575 | 32,73904 | 32,687 | 32,65421 | 32,70718 | 32,6304 | 23 | 23 |
| 31,27598 | 31,21489 | 31,34377 | 31,45518 | 31,06053 | 30,89656 | 7 | 7 |
| 30,26383 | 30,62401 | 30,12139 | 30,29307 | 30,46704 | 30,37192 | 6 | 6 |
| 31,30247 | 31,15406 | 30,89988 | 30,80355 | 30,8795 | 30,83161 | 15 | 15 |
| 22,27997 | 24,69429 | 24,58486 | 26,35085 | 24,66123 | 22,72375 | 2 | 2 |
| 24,84995 | 25,07321 | 24,58818 | 24,3379 | 24,80794 | 24,1937 | 2 | 2 |
| 33,33684 | 33,43986 | 34,88908 | 33,71957 | 33,79585 | 33,68833 | 24 | 5 |
| 27,97257 | 27,75334 | 27,62874 | 27,84476 | 27,88902 | 27,56972 | 6 | 4 |
| 25,82257 | 25,22615 | 25,59309 | 25,79479 | 25,91504 | 24,31507 | 6 | 6 |
| 27,00496 | 27,1757 | 27,32637 | 27,31536 | 27,41127 | 27,03265 | 5 | 5 |
| 31,69316 | 31,67316 | 31,66848 | 31,77518 | 31,9244 | 31,634 | 18 | 18 |
| 27,30348 | 27,23178 | 26,9165 | 26,69657 | 26,95903 | 27,0071 | 4 | 3 |
| 28,57856 | 28,35708 | 28,52914 | 28,65857 | 28,46311 | 28,87675 | 19 | 19 |
| 27,64038 | 27,45183 | 27,42238 | 27,79005 | 27,23553 | 27,26607 | 6 | 6 |
| 24,14686 | 23,89032 | 23,20334 | 21,94008 | 23,7601 | 23,72078 | 2 | 2 |
| 27,77235 | 27,56049 | 27,37327 | 27,87238 | 27,39926 | 27,67364 | 11 | 11 |
| 31,99453 | 31,83228 | 31,91823 | 31,57943 | 31,80973 | 31,76042 | 28 | 28 |
| 28,86423 | 29,46939 | 29,09421 | 29,59046 | 28,95197 | 28,65625 | 21 | 1 |
| 33,09462 | 32,91085 | 32,37358 | 32,29587 | 32,56556 | 32,35371 | 22 | 2 |
| 27,32027 | 27,54392 | 26,75646 | 27,14657 | 27,75156 | 27,88466 | 4 | 4 |
| 30,71653 | 30,51794 | 30,96478 | 30,55536 | 30,16459 | 30,62758 | 21 | 21 |
| 31,94217 | 31,6708 | 31,39938 | 31,85274 | 32,31265 | 32,06603 | 17 | 9 |

| | | | | | | | |
|---|---|---|---|---|---|---|---|
| 30,4006 | 30,5351 | 29,78821 | 30,54232 | 30,74455 | 30,82472 | 9 | 9 |
| 27,74426 | 27,89643 | 28,50233 | 27,82129 | 27,80366 | 27,60675 | 9 | 9 |
| 27,65697 | 27,7692 | 27,11555 | 27,95249 | 27,73538 | 28,00603 | 9 | 8 |
| 28,07327 | 27,93101 | 27,70147 | 28,14545 | 27,93663 | 28,52977 | 9 | 9 |
| 26,98153 | 26,52367 | 26,52153 | 26,50106 | 26,66275 | 26,09581 | 6 | 6 |
| 26,02357 | 26,25796 | 26,51164 | 26,903 | 26,50346 | 26,44847 | 3 | 3 |
| 21,8826 | 23,33074 | 22,29364 | 23,84336 | 23,28136 | 21,20855 | 2 | 2 |
| 24,34313 | 24,67659 | 25,50373 | 25,4256 | 26,32287 | 24,65453 | 4 | 4 |
| 27,48631 | 27,46217 | 27,45505 | 25,895 | 27,18291 | 26,89574 | 3 | 3 |
| 28,53753 | 27,93517 | 28,36465 | 27,90684 | 27,75945 | 28,23224 | 5 | 5 |
| 25,45676 | 25,61759 | 26,4049 | 26,23971 | 26,229 | 26,32917 | 2 | 2 |
| 25,90265 | 25,53694 | 25,03565 | 25,20221 | 25,02242 | 24,94405 | 3 | 3 |
| 25,84988 | 26,06983 | 26,14921 | 26,52589 | 26,30795 | 26,2074 | 5 | 5 |
| 25,00372 | 24,71041 | 24,65807 | 24,74052 | 25,42159 | 24,4872 | 4 | 1 |
| 26,32665 | 26,26973 | 25,98366 | 25,63259 | 25,54872 | 26,22714 | 2 | 2 |
| 24,91178 | 25,83277 | 23,39506 | 25,89634 | 24,57479 | 23,64284 | 4 | 4 |
| 25,65412 | 26,1116 | 25,10017 | 25,84014 | 25,65657 | 25,24158 | 4 | 3 |
| 26,97564 | 25,51926 | 27,85816 | 27,84775 | 28,23471 | 25,67223 | 2 | 2 |
| 30,22803 | 30,49648 | 30,13136 | 30,26461 | 30,45778 | 29,87146 | 5 | 5 |
| 26,22773 | 25,80622 | 25,89034 | 25,92662 | 26,41001 | 26,16821 | 3 | 3 |
| 25,54854 | 26,44528 | 25,31469 | 24,88455 | 25,48999 | 26,07979 | 4 | 4 |
| 27,01127 | 26,74701 | 27,36586 | 26,54464 | 27,67721 | 28,27249 | 2 | 2 |
| 29,17719 | 29,32376 | 27,34266 | 29,00158 | 28,35997 | 28,10814 | 7 | 5 |
| 28,11714 | 28,23836 | 28,66112 | 28,64282 | 28,17374 | 28,28145 | 115 | 3 |
| 30,25384 | 29,99806 | 29,95809 | 30,21327 | 29,9632 | 29,67504 | 16 | 16 |
| 28,6285 | 28,36699 | 28,54576 | 28,81632 | 28,57622 | 28,33981 | 11 | 10 |
| 28,61782 | 28,52058 | 28,65345 | 28,79387 | 28,66275 | 28,82493 | 12 | 12 |
| 27,07653 | 26,84566 | 27,01329 | 26,53832 | 26,8484 | 26,76737 | 6 | 6 |
| 21,16848 | 24,0426 | 23,60863 | 22,53893 | 23,38995 | 23,28362 | 3 | 3 |
| 27,23123 | 27,00988 | 26,44613 | 27,04291 | 26,64595 | 26,81201 | 3 | 3 |
| 27,96119 | 28,12819 | 27,68819 | 27,92281 | 27,66851 | 27,9551 | 5 | 5 |
| 25,4338 | 25,74214 | 25,63303 | 25,53677 | 25,37955 | 26,07568 | 3 | 3 |
| 27,12554 | 26,73725 | 26,55802 | 27,09093 | 27,0134 | 26,65103 | 6 | 6 |
| 26,36421 | 26,48786 | 26,13414 | 26,48866 | 25,46678 | 25,43377 | 3 | 3 |
| 27,79284 | 27,53934 | 27,68271 | 27,28921 | 27,59562 | 27,12613 | 10 | 10 |
| 26,43248 | 27,40568 | 26,73982 | 26,15977 | 25,57462 | 26,26345 | 7 | 7 |
| 23,30508 | 24,52725 | 24,52277 | 24,31665 | 24,88152 | 24,86008 | 2 | 2 |
| 25,48148 | 24,92393 | 25,38225 | 25,50443 | 24,42937 | 25,27835 | 4 | 4 |
| 30,54075 | 30,59246 | 29,82438 | 30,39069 | 30,75366 | 30,65113 | 10 | 10 |
| 28,74204 | 28,52996 | 28,21716 | 27,97027 | 28,63471 | 28,21951 | 8 | 8 |
| 24,64493 | 25,09156 | 24,3682 | 22,06855 | 24,93661 | 25,27291 | 4 | 4 |
| 27,71816 | 27,73989 | 27,29334 | 27,4243 | 27,48938 | 27,15103 | 6 | 6 |
| 25,61026 | 25,74509 | 26,62756 | 26,06766 | 25,51415 | 26,28198 | 7 | 7 |
| 28,52268 | 28,32487 | 28,49922 | 28,21984 | 28,46018 | 28,34613 | 11 | 11 |
| 26,93708 | 27,68284 | 27,34223 | 27,18518 | 27,09154 | 27,18471 | 5 | 5 |
| 25,07904 | 26,43663 | 25,96585 | 26,30372 | 26,76028 | 25,94213 | 3 | 3 |
| 28,17997 | 27,73731 | 28,03459 | 28,97197 | 28,72289 | 28,17493 | 9 | 9 |
| 27,50491 | 27,4684 | 27,55976 | 27,67209 | 26,97804 | 27,66777 | 10 | 10 |
| 25,16672 | 25,07997 | 25,03699 | 25,10574 | 25,04323 | 24,67309 | 2 | 2 |
| 25,05441 | 25,12282 | 24,73959 | 24,98529 | 25,06007 | 24,94673 | 3 | 3 |

| | | | | | | | |
|---|---|---|---|---|---|---|---|
| 26,72702 | 25,62751 | 26,8236 | 24,8561 | 26,61821 | 25,19028 | 5 | 5 |
| 27,03213 | 26,68003 | 26,99335 | 26,49962 | 26,95382 | 26,21751 | 4 | 4 |
| 28,12362 | 28,36678 | 27,8521 | 27,99723 | 28,18702 | 28,31043 | 6 | 6 |
| 32,36494 | 32,59308 | 32,19824 | 32,26352 | 32,34824 | 32,39928 | 10 | 10 |
| 30,19566 | 30,37648 | 30,4342 | 30,53408 | 30,50654 | 30,52337 | 25 | 25 |
| 25,79749 | 26,87408 | 26,97016 | 27,28189 | 24,75913 | 25,96673 | 2 | 2 |
| 23,81802 | 24,37524 | 24,95367 | 24,32766 | 24,66552 | 24,53919 | 2 | 2 |
| 27,96152 | 28,16485 | 28,29557 | 27,78481 | 28,17902 | 28,22159 | 9 | 9 |
| 26,641 | 27,16576 | 26,02542 | 25,80504 | 25,81919 | 26,85792 | 2 | 2 |
| 24,7415 | 24,69912 | 24,96274 | 24,6609 | 24,26335 | 24,81294 | 29 | 2 |
| 26,21345 | 26,77618 | 25,62171 | 26,37552 | 25,36289 | 24,86036 | 30 | 3 |
| 26,72495 | 27,37493 | 26,53739 | 27,76269 | 27,33356 | 27,43508 | 3 | 3 |
| 25,49048 | 26,22 | 25,88169 | 25,95626 | 25,94905 | 25,82396 | 5 | 5 |
| 25,47494 | 24,79828 | 24,89708 | 24,47482 | 24,71633 | 24,63976 | 2 | 2 |
| 25,73531 | 25,92103 | 26,19078 | 25,7341 | 25,8719 | 26,12424 | 4 | 4 |
| 31,23638 | 31,92 | 31,1343 | 31,78842 | 31,75529 | 31,86009 | 10 | 2 |
| 25,59215 | 25,92477 | 24,87202 | 25,10717 | 25,32783 | 25,48582 | 5 | 5 |
| 28,03779 | 27,79538 | 28,15451 | 28,19756 | 28,29491 | 28,29758 | 10 | 10 |
| 24,53594 | 24,19708 | 23,85158 | 24,22656 | 24,04668 | 23,91789 | 2 | 2 |
| 25,50176 | 25,68675 | 24,88488 | 25,2505 | 25,45905 | 24,27938 | 3 | 3 |
| 28,25948 | 27,78525 | 27,02971 | 28,18834 | 28,4273 | 28,54344 | 6 | 6 |
| 31,26411 | 31,32866 | 31,3628 | 31,24161 | 31,24632 | 31,41084 | 19 | 1 |
| 27,55786 | 27,46256 | 27,56827 | 27,09979 | 27,67909 | 27,70121 | 8 | 8 |
| 26,74548 | 26,56834 | 26,4356 | 26,0842 | 26,67801 | 26,45555 | 3 | 3 |
| 25,26259 | 25,07022 | 24,62282 | 24,78352 | 25,17665 | 25,01644 | 2 | 2 |
| 25,18733 | 24,78831 | 25,30466 | 25,13907 | 24,59679 | 24,86646 | 4 | 4 |
| 25,62762 | 25,10121 | 25,46286 | 25,27412 | 25,03955 | 25,18279 | 4 | 4 |
| 33,20615 | 33,3028 | 33,27489 | 33,41665 | 33,31757 | 33,47024 | 70 | 70 |
| 25,92626 | 25,98561 | 25,50294 | 25,88779 | 25,74237 | 25,94156 | 3 | 3 |
| 26,22008 | 26,94 | 27,37832 | 26,56152 | 26,66357 | 25,1902 | 5 | 5 |
| 28,03941 | 27,73982 | 27,45967 | 27,89574 | 27,74746 | 27,90828 | 8 | 8 |
| 26,50167 | 26,15172 | 26,49132 | 26,82675 | 26,42856 | 26,65499 | 6 | 6 |
| 26,78669 | 26,77392 | 26,14435 | 26,53995 | 25,9572 | 26,49069 | 2 | 2 |
| 25,48821 | 25,07957 | 23,51581 | 25,64405 | 25,54144 | 24,61189 | 2 | 2 |
| 24,85339 | 25,11406 | 25,04226 | 25,35609 | 25,10013 | 25,23422 | 3 | 3 |
| 26,68992 | 26,79699 | 26,81495 | 26,36418 | 27,12416 | 26,6513 | 2 | 2 |
| 27,75239 | 27,18801 | 26,41387 | 27,07327 | 26,98848 | 26,83774 | 9 | 9 |
| 26,48968 | 26,78194 | 26,45436 | 26,61274 | 26,41928 | 26,39699 | 4 | 4 |
| 30,61316 | 30,95109 | 30,29669 | 31,01691 | 31,02393 | 30,78207 | 10 | 10 |
| 24,92583 | 24,92456 | 24,92266 | 25,18726 | 24,82236 | 24,72697 | 4 | 4 |
| 21,32921 | 21,54182 | 22,41723 | 19,99907 | 21,84556 | 22,35842 | 2 | 2 |
| 24,97836 | 24,46787 | 23,99516 | 23,83045 | 24,02513 | 23,85005 | 2 | 2 |
| 25,35269 | 26,16331 | 24,76901 | 25,81115 | 25,53133 | 24,98107 | 2 | 2 |
| 28,05745 | 28,06142 | 27,61905 | 27,6476 | 27,91553 | 27,13575 | 7 | 7 |
| 28,4299 | 28,4847 | 28,22209 | 28,20373 | 28,04713 | 28,26392 | 11 | 1 |
| 27,16576 | 26,81238 | 27,48232 | 27,52709 | 27,42085 | 27,87168 | 8 | 8 |
| 27,58999 | 27,63402 | 27,37493 | 27,34883 | 27,67707 | 27,30722 | 8 | 8 |
| 28,57456 | 28,81885 | 28,45227 | 28,82829 | 28,67711 | 28,70909 | 15 | 15 |
| 25,71358 | 25,53913 | 25,25407 | 25,71694 | 25,55441 | 25,59369 | 3 | 3 |
| 32,20557 | 30,58826 | 30,66102 | 31,73788 | 32,33261 | 31,06516 | 10 | 10 |

| | | | | | | | |
|---|---|---|---|---|---|---|---|
| 26,7604 | 26,35178 | 26,52977 | 26,31622 | 25,95757 | 25,75496 | 6 | 6 |
| 26,25296 | 26,82723 | 25,83651 | 26,96246 | 26,16594 | 25,81736 | 97 | 1 |
| 28,06907 | 27,81543 | 28,16883 | 27,96113 | 27,74746 | 27,9635 | 11 | 11 |
| 24,9604 | 25,42832 | 24,81719 | 24,71785 | 24,9822 | 25,41693 | 2 | 2 |
| 25,6445 | 25,78159 | 25,80366 | 27,14005 | 25,6502 | 26,23621 | 4 | 4 |
| 28,19531 | 28,04874 | 28,07465 | 28,13629 | 27,74849 | 27,89406 | 11 | 11 |
| 26,4016 | 26,99281 | 26,68779 | 26,61034 | 26,50937 | 26,17037 | 3 | 3 |
| 32,15065 | 31,94688 | 32,08023 | 32,04257 | 32,17754 | 32,08322 | 33 | 33 |
| 30,12436 | 29,83123 | 30,05271 | 29,89836 | 29,96292 | 29,94838 | 22 | 22 |
| 25,21171 | 24,86084 | 25,25703 | 25,12617 | 24,73836 | 24,33231 | 3 | 3 |
| 27,42358 | 27,63519 | 26,99776 | 27,12347 | 27,36961 | 26,90885 | 6 | 6 |
| 28,4717 | 28,60134 | 28,48331 | 28,56678 | 27,55516 | 28,07403 | 9 | 9 |
| 29,88371 | 29,52002 | 29,75344 | 29,69752 | 29,85554 | 29,91912 | 17 | 1 |
| 27,51443 | 27,08477 | 27,60414 | 27,36461 | 27,53927 | 27,46428 | 7 | 7 |
| 24,86344 | 25,42256 | 25,02732 | 25,55124 | 25,45601 | 25,56069 | 5 | 5 |
| 24,23739 | 24,14942 | 23,48987 | 24,6602 | 25,16932 | 24,51376 | 2 | 2 |
| 25,02952 | 25,61411 | 25,88846 | 24,92103 | 25,41754 | 25,34564 | 3 | 3 |
| 26,0112 | 25,8593 | 25,98574 | 26,03938 | 26,15607 | 26,2917 | 4 | 4 |
| 26,01171 | 25,71557 | 26,02403 | 26,19768 | 25,18873 | 25,76426 | 6 | 6 |
| 31,86116 | 31,53036 | 31,8445 | 31,58491 | 31,65745 | 31,6388 | 33 | 33 |
| 27,86284 | 27,64389 | 28,79132 | 27,51698 | 27,59199 | 28,44526 | 3 | 3 |
| 26,96102 | 27,10799 | 27,14802 | 26,24256 | 26,49222 | 26,89087 | 6 | 6 |
| 25,49204 | 24,12825 | 22,55259 | 23,51232 | 23,84747 | 21,86738 | 17 | 1 |
| 30,32646 | 30,17462 | 30,50966 | 30,20163 | 30,10262 | 30,3226 | 20 | 20 |
| 27,24336 | 27,29299 | 26,75136 | 26,78519 | 26,71594 | 26,6803 | 4 | 4 |
| 29,97951 | 29,80264 | 29,42786 | 29,35195 | 29,60848 | 29,30149 | 5 | 5 |
| 30,3952 | 30,35448 | 30,37493 | 30,49943 | 30,38577 | 30,58172 | 13 | 13 |
| 27,9015 | 27,49747 | 28,15706 | 27,49777 | 27,95532 | 27,76231 | 9 | 9 |
| 23,31327 | 22,26213 | 23,22219 | 23,41326 | 22,78479 | 24,32375 | 2 | 2 |
| 26,79934 | 26,73673 | 26,15341 | 26,96367 | 27,13673 | 27,16461 | 6 | 6 |
| 24,5412 | 24,70972 | 25,19468 | 24,50803 | 24,42265 | 24,50785 | 18 | 1 |
| 25,17348 | 25,29793 | 25,32978 | 24,80971 | 25,10805 | 25,34604 | 3 | 3 |
| 27,61723 | 27,65191 | 27,19517 | 27,67788 | 27,82499 | 27,29789 | 10 | 10 |
| 26,23608 | 26,03465 | 25,81558 | 25,98917 | 25,1979 | 25,8501 | 2 | 2 |
| 27,39836 | 27,34223 | 27,21928 | 27,10199 | 27,2203 | 27,41393 | 8 | 8 |
| 25,37829 | 24,71905 | 24,47976 | 24,67293 | 24,98081 | 24,73702 | 3 | 3 |
| 25,89025 | 26,59662 | 26,07366 | 24,35616 | 25,85816 | 25,47676 | 6 | 6 |
| 29,24577 | 29,33134 | 28,81231 | 29,40787 | 29,39562 | 29,64959 | 9 | 9 |
| 23,91579 | 22,79755 | 23,19436 | 24,60676 | 22,83936 | 22,86208 | 3 | 3 |
| 26,62729 | 26,05877 | 25,99166 | 25,94156 | 24,50361 | 25,59912 | 4 | 4 |
| 26,94659 | 27,00592 | 26,73879 | 26,52585 | 26,59833 | 25,93148 | 6 | 6 |
| 25,94815 | 25,88409 | 25,2913 | 25,45262 | 25,24978 | 23,99411 | 2 | 2 |
| 25,41674 | 24,32237 | 24,48339 | 25,22483 | 24,73195 | 25,54954 | 3 | 3 |
| 29,08241 | 28,64379 | 28,75344 | 28,77096 | 28,69495 | 29,35641 | 11 | 11 |
| 24,64686 | 24,00924 | 23,93238 | 24,19925 | 22,93577 | 24,04427 | 4 | 4 |
| 25,81651 | 25,97018 | 21,40283 | 25,54937 | 24,95327 | 25,23418 | 4 | 4 |
| 34,35021 | 34,56666 | 34,32268 | 34,35508 | 34,25673 | 34,22367 | 18 | 18 |
| 26,32925 | 26,39547 | 26,05012 | 26,55605 | 26,33006 | 25,90389 | 5 | 5 |
| 21,10791 | 24,29334 | 24,69376 | 24,10988 | 23,98216 | 24,2617 | 2 | 2 |
| 30,94705 | 30,55206 | 31,01585 | 30,4685 | 30,13761 | 30,60761 | 18 | 1 |

| | | | | | | | |
|---|---|---|---|---|---|---|---|
| 28,4451 | 27,9462 | 27,72813 | 28,1055 | 28,03123 | 27,83635 | 6 | 6 |
| 28,68027 | 28,44439 | 27,96422 | 27,96224 | 27,76591 | 27,66126 | 3 | 3 |
| 26,76926 | 26,80058 | 26,55541 | 26,87572 | 27,12859 | 26,89885 | 9 | 1 |
| 28,28352 | 28,19046 | 27,96785 | 27,96659 | 27,94815 | 27,7775 | 5 | 5 |
| 26,60258 | 26,89006 | 25,43275 | 26,0425 | 26,05041 | 26,13718 | 6 | 6 |
| 31,89969 | 31,81597 | 31,96513 | 31,72886 | 31,75561 | 31,81803 | 40 | 40 |
| 27,5974 | 27,40194 | 27,99868 | 27,24391 | 26,90369 | 28,11252 | 2 | 1 |
| 25,15815 | 25,20195 | 25,6115 | 25,77911 | 25,26474 | 25,17634 | 5 | 5 |
| 24,25018 | 24,68019 | 24,94664 | 24,33245 | 24,503 | 24,4309 | 3 | 2 |
| 27,25755 | 27,50506 | 27,05352 | 26,64774 | 26,79625 | 27,09537 | 11 | 6 |
| 25,48142 | 26,52953 | 25,67441 | 26,37764 | 26,12662 | 26,61063 | 5 | 5 |
| 27,95166 | 27,84212 | 28,05889 | 27,8584 | 27,71293 | 27,90483 | 9 | 9 |
| 25,95496 | 25,807 | 25,60473 | 25,68579 | 25,07953 | 25,88085 | 2 | 2 |
| 28,3362 | 28,13951 | 28,22582 | 28,1155 | 27,96025 | 28,10869 | 9 | 9 |
| 26,52106 | 26,47748 | 27,07989 | 26,9266 | 27,21632 | 26,64113 | 2 | 2 |
| 23,59537 | 23,83248 | 23,6049 | 23,6513 | 24,0385 | 23,81078 | 2 | 2 |
| 24,61986 | 25,22938 | 24,51057 | 23,87961 | 24,54162 | 24,18684 | 2 | 2 |
| 27,68892 | 27,74387 | 27,57824 | 27,77549 | 27,68659 | 27,56049 | 8 | 8 |
| 26,95626 | 27,24654 | 26,68926 | 27,16519 | 27,17331 | 27,68499 | 5 | 5 |
| 29,91257 | 29,87012 | 29,56692 | 30,08854 | 29,85091 | 29,8235 | 11 | 11 |
| 23,77505 | 23,90876 | 23,70007 | 22,42653 | 23,6634 | 24,81783 | 17 | 1 |
| 26,99873 | 27,22435 | 26,41487 | 26,40934 | 27,31112 | 27,23936 | 5 | 5 |
| 30,34805 | 29,41597 | 31,04616 | 30,53955 | 32,03901 | 29,88129 | 2 | 2 |
| 25,35877 | 24,64785 | 24,12943 | 24,30966 | 24,04519 | 24,57439 | 2 | 2 |
| 27,41256 | 28,5968 | 28,54008 | 28,25723 | 27,37393 | 28,18943 | 4 | 4 |
| 26,17703 | 26,66804 | 25,75318 | 25,62087 | 25,49415 | 25,10673 | 4 | 4 |
| 28,72159 | 28,35288 | 27,27124 | 28,57376 | 28,57308 | 27,67169 | 13 | 1 |
| 28,10275 | 28,51058 | 28,38356 | 28,18782 | 28,31437 | 28,35229 | 20 | 20 |
| 22,52174 | 22,39799 | 24,21963 | 22,74236 | 23,61941 | 23,08622 | 3 | 3 |
| 27,51051 | 26,84386 | 26,90759 | 26,69975 | 26,98218 | 26,71529 | 15 | 1 |
| 27,31709 | 27,80261 | 27,1046 | 27,82863 | 27,19995 | 27,47088 | 6 | 6 |
| 24,75771 | 25,57415 | 25,2285 | 25,55789 | 25,24202 | 25,45472 | 3 | 3 |
| 28,59046 | 28,86432 | 28,53063 | 28,36611 | 28,61618 | 28,53738 | 3 | 3 |
| 26,7433 | 26,71908 | 26,4971 | 26,76648 | 26,56745 | 26,38765 | 7 | 7 |
| 25,77658 | 26,79042 | 27,23945 | 26,48267 | 26,50676 | 26,98783 | 5 | 5 |
| 28,18575 | 27,79185 | 28,28162 | 27,77982 | 27,57131 | 28,22619 | 12 | 12 |
| 29,65803 | 29,49295 | 29,16209 | 29,3663 | 29,01422 | 29,02308 | 10 | 10 |
| 26,36465 | 26,13434 | 26,5027 | 27,03559 | 26,39284 | 26,66397 | 50 | 2 |
| 29,94224 | 29,71673 | 29,59336 | 29,40377 | 29,49932 | 29,52518 | 15 | 15 |
| 27,92954 | 27,50165 | 27,8929 | 27,92993 | 28,00651 | 28,1038 | 10 | 10 |
| 28,15682 | 28,12401 | 27,79023 | 28,22002 | 28,35246 | 28,11436 | 7 | 7 |
| 25,4641 | 24,87061 | 25,07014 | 26,48173 | 25,08902 | 26,0507 | 4 | 4 |
| 28,70458 | 28,52563 | 27,67405 | 27,5087 | 27,35321 | 27,51646 | 7 | 7 |
| 27,02654 | 26,475 | 27,35405 | 27,78457 | 26,83062 | 27,09225 | 5 | 5 |
| 29,32496 | 28,85156 | 29,00512 | 28,66919 | 29,07074 | 28,14511 | 5 | 5 |
| 26,05082 | 25,86257 | 25,19156 | 26,05329 | 25,23907 | 24,6324 | 7 | 7 |
| 27,77543 | 27,38878 | 27,78238 | 27,81415 | 27,63851 | 27,91929 | 13 | 1 |
| 24,97666 | 25,32364 | 25,15132 | 24,93894 | 25,42742 | 25,10897 | 3 | 3 |
| 27,05569 | 27,29316 | 27,02623 | 27,21956 | 27,33091 | 27,26222 | 4 | 4 |
| 26,20749 | 26,43285 | 26,33865 | 26,6064 | 26,33579 | 26,20247 | 4 | 4 |

| | | | | | | | |
|---|---|---|---|---|---|---|---|
| 27,20937 | 27,14462 | 26,86915 | 26,81054 | 27,1812 | 27,00796 | 3 | 3 |
| 28,16265 | 28,20802 | 27,77894 | 28,16734 | 27,99599 | 28,08071 | 6 | 6 |
| 26,06651 | 26,38877 | 25,71759 | 25,45105 | 25,74114 | 25,17489 | 6 | 6 |
| 28,90455 | 28,82451 | 28,36666 | 28,30256 | 28,42766 | 28,71221 | 4 | 4 |
| 32,22472 | 31,93044 | 31,81711 | 31,78058 | 32,02934 | 31,89475 | 23 | 23 |
| 28,38944 | 28,75022 | 28,21697 | 27,70154 | 28,17897 | 28,30208 | 5 | 5 |
| 26,52809 | 26,03913 | 26,64141 | 26,27805 | 26,69299 | 25,6803 | 5 | 5 |
| 26,81775 | 26,54234 | 26,96213 | 26,63686 | 26,43817 | 26,6224 | 6 | 6 |
| 26,06126 | 26,76433 | 25,99317 | 26,38829 | 25,69755 | 25,3221 | 7 | 7 |
| 25,38952 | 25,19272 | 24,97701 | 25,09438 | 24,96076 | 24,72349 | 3 | 3 |
| 26,83425 | 27,0357 | 27,38047 | 27,41062 | 27,23142 | 27,33833 | 8 | 8 |
| 25,96133 | 25,32711 | 24,98598 | 22,38437 | 26,53559 | 25,17862 | 2 | 2 |
| 24,60722 | 25,14554 | 25,53228 | 24,84537 | 24,84666 | 25,05945 | 4 | 4 |
| 25,96766 | 26,34533 | 26,00494 | 26,25425 | 26,25278 | 26,41368 | 5 | 5 |
| 24,60925 | 24,49631 | 25,05205 | 24,35669 | 24,60546 | 24,79576 | 3 | 3 |
| 31,81019 | 31,60585 | 31,57849 | 31,70997 | 31,69793 | 31,5472 | 39 | 39 |
| 29,8941 | 29,62716 | 29,71506 | 29,60291 | 29,47837 | 29,5341 | 12 | 12 |
| 25,1208 | 23,66036 | 24,11012 | 25,11831 | 25,27459 | 25,31921 | 2 | 2 |
| 27,52111 | 27,66037 | 27,18971 | 27,07255 | 27,1574 | 27,26464 | 7 | 7 |
| 24,93292 | 25,53638 | 24,36833 | 24,68169 | 24,24285 | 25,41248 | 3 | 3 |
| 27,38204 | 27,91656 | 27,20574 | 27,69199 | 27,14793 | 27,34189 | 4 | 4 |
| 24,51256 | 24,65606 | 25,1617 | 25,27185 | 24,2214 | 25,37186 | 2 | 2 |
| 27,22858 | 27,22628 | 26,7265 | 27,03107 | 27,15374 | 27,15277 | 4 | 4 |
| 26,82214 | 26,43349 | 26,84038 | 25,97773 | 26,32423 | 25,14239 | 2 | 2 |
| 26,25909 | 26,31057 | 25,86389 | 25,97029 | 26,17919 | 26,53838 | 4 | 4 |
| 25,39816 | 25,45271 | 25,77731 | 25,40354 | 26,1809 | 25,61156 | 3 | 1 |
| 26,09702 | 26,03735 | 25,76271 | 25,49118 | 25,86495 | 25,87586 | 4 | 4 |
| 25,95385 | 26,23063 | 25,50963 | 26,33103 | 26,46616 | 26,49733 | 3 | 3 |
| 29,93593 | 29,55083 | 29,6268 | 29,47717 | 29,13631 | 29,37706 | 15 | 15 |
| 23,78027 | 24,5415 | 23,70567 | 23,73097 | 24,0703 | 24,70446 | 2 | 2 |
| 24,61257 | 25,03065 | 21,8951 | 24,32711 | 24,45651 | 24,86797 | 2 | 2 |
| 23,33715 | 23,42905 | 23,181 | 23,045 | 23,05416 | 23,24879 | 2 | 2 |
| 26,36328 | 26,10797 | 26,17238 | 26,08898 | 25,83502 | 25,64243 | 5 | 5 |
| 28,00341 | 27,60527 | 27,29378 | 27,66316 | 27,73124 | 27,59427 | 7 | 7 |
| 30,64693 | 30,45268 | 30,1332 | 30,42972 | 30,14321 | 30,42642 | 11 | 11 |
| 28,17455 | 28,78372 | 28,68003 | 28,0748 | 29,00021 | 29,01039 | 6 | 6 |
| 26,91718 | 26,82833 | 26,80943 | 26,75494 | 26,70846 | 26,54637 | 4 | 4 |
| 23,97858 | 22,49885 | 21,0454 | 22,97008 | 22,81306 | 22,86597 | 2 | 2 |
| 25,98074 | 25,19494 | 25,79444 | 25,68683 | 25,50176 | 25,91568 | 3 | 3 |
| 28,84915 | 28,51202 | 28,64107 | 28,49133 | 28,54797 | 28,73854 | 11 | 11 |
| 25,46326 | 25,93267 | 25,96596 | 25,57695 | 25,61515 | 25,0537 | 2 | 2 |
| 23,70176 | 23,7166 | 25,19749 | 24,81201 | 24,82474 | 24,58187 | 3 | 3 |
| 27,31346 | 27,06353 | 25,81512 | 26,38657 | 26,81201 | 25,98603 | 5 | 5 |
| 30,23683 | 30,19437 | 30,08601 | 30,12964 | 29,94977 | 30,15834 | 12 | 12 |
| 27,90133 | 28,04328 | 27,9081 | 27,51337 | 27,89816 | 29,00961 | 6 | 6 |
| 29,38078 | 29,26033 | 28,91835 | 29,54359 | 28,86662 | 29,2252 | 12 | 12 |
| 29,53699 | 29,22057 | 29,27689 | 29,06199 | 29,04458 | 29,10996 | 8 | 8 |
| 28,49987 | 28,5333 | 28,57333 | 28,7326 | 28,79433 | 28,50843 | 11 | 11 |
| 28,81925 | 29,06175 | 28,57261 | 28,86564 | 29,04025 | 28,82908 | 9 | 9 |
| 28,52174 | 27,99642 | 28,51717 | 28,49362 | 28,12223 | 28,05227 | 13 | 13 |

| | | | | | | | |
|---|---|---|---|---|---|---|---|
| 23,74991 | 23,23421 | 23,08161 | 23,5254 | 23,2397 | 21,20729 | 3 | 3 |
| 24,93197 | 23,95553 | 24,17912 | 24,11514 | 23,79174 | 24,21393 | 3 | 3 |
| 27,90351 | 27,75583 | 27,50408 | 27,95532 | 27,86638 | 27,59156 | 8 | 8 |
| 25,3931 | 25,34682 | 25,14985 | 25,90871 | 25,04381 | 24,91178 | 3 | 3 |
| 24,61223 | 24,16615 | 23,94566 | 23,87558 | 24,09184 | 23,72682 | 4 | 4 |
| 28,01829 | 27,87947 | 27,51141 | 27,75646 | 27,82675 | 27,21067 | 5 | 5 |
| 28,76623 | 28,69369 | 28,37509 | 28,57777 | 28,31428 | 28,08735 | 6 | 6 |
| 25,95256 | 25,83441 | 26,286 | 25,95613 | 25,54766 | 25,61541 | 5 | 5 |
| 28,06157 | 27,86644 | 27,14899 | 26,86361 | 27,29693 | 28,03941 | 12 | 12 |
| 29,0833 | 28,97766 | 28,54613 | 29,09989 | 28,94924 | 28,89467 | 14 | 14 |
| 24,98107 | 24,41429 | 24,48308 | 24,59542 | 23,20305 | 23,40119 | 2 | 2 |
| 26,26878 | 26,92705 | 25,98085 | 26,04059 | 26,68793 | 26,65663 | 2 | 2 |
| 27,85858 | 28,11426 | 28,31251 | 27,84625 | 27,86136 | 28,37638 | 9 | 9 |
| 26,05399 | 25,73337 | 25,76149 | 25,76562 | 26,38388 | 26,15111 | 3 | 3 |
| 25,96882 | 25,94219 | 25,65946 | 25,11287 | 25,6184 | 25,428 | 4 | 4 |
| 27,52238 | 27,69073 | 27,47839 | 27,52358 | 27,31726 | 26,98174 | 4 | 4 |
| 23,83441 | 24,04618 | 23,76577 | 23,84086 | 23,89504 | 23,85709 | 2 | 2 |
| 28,05941 | 27,10799 | 26,64155 | 27,80397 | 26,51265 | 27,01041 | 3 | 3 |
| 24,13273 | 25,02301 | 23,65458 | 24,40503 | 24,03171 | 24,56674 | 3 | 3 |
| 26,92218 | 26,94078 | 27,00099 | 27,00539 | 26,74111 | 27,25584 | 7 | 7 |
| 24,55245 | 24,74088 | 24,96014 | 25,06275 | 25,70179 | 24,50312 | 3 | 3 |
| 29,76721 | 29,52416 | 29,40377 | 29,4008 | 29,11299 | 29,44851 | 11 | 11 |
| 29,59561 | 30,00919 | 29,82224 | 29,90842 | 29,59731 | 30,01132 | 6 | 6 |
| 25,56895 | 25,57528 | 24,98298 | 25,47698 | 25,14612 | 25,38245 | 3 | 3 |
| 28,10345 | 27,82772 | 28,0055 | 27,32225 | 28,01105 | 27,53727 | 10 | 10 |
| 28,79002 | 28,52073 | 28,59188 | 28,57355 | 28,6059 | 28,7785 | 8 | 8 |
| 28,31406 | 27,66472 | 27,62902 | 27,3238 | 27,55837 | 27,91336 | 9 | 9 |
| 25,58092 | 25,57816 | 25,27452 | 26,86349 | 26,05115 | 25,33623 | 4 | 4 |
| 27,83485 | 27,73389 | 27,81048 | 27,74554 | 27,99868 | 28,00999 | 10 | 10 |
| 26,69564 | 26,26132 | 26,83123 | 26,12841 | 26,84757 | 25,71683 | 2 | 2 |
| 27,53289 | 27,30191 | 27,34951 | 27,99658 | 27,21937 | 27,48999 | 2 | 2 |
| 27,29754 | 27,54929 | 26,96862 | 27,87156 | 27,39697 | 27,45967 | 9 | 2 |
| 26,52998 | 26,63907 | 26,27565 | 26,47761 | 26,33685 | 26,55416 | 4 | 4 |
| 27,00174 | 26,52323 | 26,93112 | 26,7465 | 26,24889 | 26,50038 | 2 | 2 |
| 27,20462 | 28,40722 | 28,42514 | 28,96041 | 28,52966 | 28,50207 | 4 | 4 |
| 28,46552 | 28,14914 | 28,35569 | 28,54639 | 28,47792 | 28,49781 | 6 | 6 |
| 26,96334 | 26,81861 | 26,91718 | 26,77077 | 26,55169 | 26,86302 | 5 | 5 |
| 31,61404 | 31,41498 | 31,51164 | 31,31127 | 31,60761 | 31,4404 | 15 | 15 |
| 28,63204 | 28,38101 | 29,18348 | 28,90747 | 29,13783 | 29,39832 | 9 | 9 |
| 26,37292 | 26,62895 | 27,00024 | 27,11506 | 26,63367 | 26,66289 | 5 | 5 |
| 25,88008 | 26,55767 | 25,83827 | 25,52286 | 25,71196 | 24,97246 | 4 | 4 |
| 25,4849 | 25,86493 | 24,94838 | 24,6621 | 26,45435 | 25,01899 | 3 | 3 |
| 27,14112 | 27,09376 | 26,54511 | 27,13085 | 27,59356 | 27,28339 | 6 | 6 |
| 24,47593 | 25,24858 | 24,8122 | 24,75261 | 24,74473 | 25,79635 | 2 | 2 |
| 29,63597 | 29,51452 | 28,51612 | 28,21873 | 29,0874 | 28,25273 | 5 | 5 |
| 27,8811 | 27,85144 | 28,20317 | 27,83159 | 28,21354 | 27,82281 | 11 | 11 |
| 30,34435 | 30,3662 | 30,44894 | 30,28162 | 30,12485 | 30,28273 | 22 | 22 |
| 28,74464 | 28,3758 | 28,61358 | 28,80045 | 27,84912 | 28,22871 | 16 | 16 |
| 28,98563 | 29,15137 | 29,30456 | 28,91886 | 28,70118 | 29,0723 | 15 | 15 |

| | | | | | | | |
|---|---|---|---|---|---|---|---|
| 28,9377 | 28,81287 | 28,13785 | 28,41224 | 28,65031 | 28,54893 | 10 | 10 |
| 29,29467 | 29,12655 | 29,07505 | 29,16835 | 29,25023 | 29,16914 | 16 | 16 |
| 25,12562 | 25,38611 | 25,7507 | 25,29169 | 25,13371 | 25,90336 | 3 | 3 |
| 28,42029 | 28,11872 | 28,11053 | 28,59477 | 28,39738 | 28,26405 | 8 | 8 |
| 25,09043 | 25,71994 | 25,40458 | 25,49598 | 24,95762 | 25,1029 | 3 | 3 |
| 29,42972 | 29,36394 | 28,96832 | 29,50106 | 29,87912 | 29,75527 | 12 | 12 |
| 25,37753 | 25,35461 | 25,03834 | 23,23747 | 25,23319 | 24,98203 | 4 | 4 |
| 26,10258 | 27,17855 | 26,94414 | 27,82165 | 27,21827 | 26,98283 | 2 | 2 |
| 29,17926 | 28,69189 | 28,4642 | 29,36601 | 28,95912 | 29,00239 | 8 | 8 |
| 21,80601 | 22,12676 | 21,28777 | 20,87455 | 23,05763 | 22,6027 | 2 | 2 |
| 24,55526 | 24,3648 | 25,71819 | 25,41047 | 25,5627 | 24,62377 | 3 | 3 |
| 28,29938 | 27,98087 | 27,39583 | 27,41538 | 26,94011 | 26,79328 | 2 | 2 |
| 30,84109 | 29,36942 | 30,19378 | 30,49933 | 30,90239 | 30,17902 | 5 | 5 |
| 27,04521 | 27,08891 | 27,128 | 27,68752 | 27,21327 | 26,99367 | 5 | 5 |
| 24,97531 | 25,17192 | 25,65884 | 24,61172 | 25,07565 | 25,06312 | 3 | 3 |
| 25,25944 | 24,67098 | 26,13408 | 25,00123 | 24,72879 | 24,57467 | 5 | 5 |
| 28,40755 | 28,0391 | 27,84362 | 28,0607 | 27,87015 | 27,66702 | 8 | 8 |
| 29,4187 | 29,50926 | 29,54427 | 29,26752 | 29,66748 | 29,74029 | 19 | 19 |
| 28,4186 | 28,43261 | 28,40852 | 27,01489 | 27,22177 | 28,20658 | 8 | 8 |
| 25,28023 | 25,72705 | 25,47584 | 25,67357 | 25,21679 | 24,89611 | 3 | 3 |
| 30,83748 | 30,59183 | 30,66518 | 30,45346 | 30,64693 | 30,50427 | 22 | 22 |
| 29,90067 | 29,66711 | 29,45906 | 29,64151 | 29,24059 | 29,17962 | 7 | 7 |
| 29,34568 | 28,78538 | 28,9384 | 28,93593 | 29,28767 | 29,10047 | 5 | 5 |
| 26,78481 | 27,15837 | 26,79934 | 26,90931 | 26,564 | 25,73787 | 4 | 4 |
| 24,83379 | 24,42092 | 24,45864 | 24,50306 | 24,37132 | 24,59895 | 2 | 2 |
| 27,98044 | 27,96119 | 28,33799 | 28,52526 | 28,30835 | 28,29933 | 7 | 7 |
| 26,27957 | 26,18771 | 25,60676 | 25,89238 | 26,21245 | 26,40617 | 7 | 7 |
| 26,19197 | 26,23794 | 25,95684 | 25,90974 | 26,18522 | 26,29037 | 3 | 3 |
| 29,1655 | 29,15509 | 29,01404 | 28,9755 | 28,67512 | 28,5244 | 13 | 13 |
| 31,14364 | 30,89627 | 31,10724 | 30,84311 | 30,55837 | 30,82805 | 21 | 7 |
| 29,20799 | 28,55142 | 28,08806 | 29,32882 | 29,67361 | 28,6442 | 6 | 6 |
| 27,70938 | 27,70734 | 28,14749 | 27,67573 | 27,50673 | 27,85055 | 6 | 6 |
| 30,49962 | 30,33884 | 30,27531 | 30,2518 | 30,33639 | 30,11309 | 17 | 17 |
| 35,18486 | 35,17262 | 35,06892 | 35,0975 | 34,93473 | 34,95246 | 78 | 78 |
| 30,37958 | 30,54499 | 29,82929 | 30,30062 | 30,24558 | 30,17926 | 13 | 13 |
| 27,61541 | 27,61491 | 27,71594 | 27,32208 | 27,66648 | 27,23498 | 11 | 11 |
| 27,10069 | 27,03065 | 26,79811 | 26,84158 | 27,49221 | 27,26517 | 4 | 4 |
| 28,00287 | 28,03538 | 27,80913 | 28,06415 | 27,97738 | 28,05703 | 14 | 14 |
| 25,88451 | 26,34166 | 26,40648 | 25,87368 | 26,17013 | 26,02878 | 4 | 2 |
| 25,96508 | 25,60215 | 25,71028 | 25,75972 | 26,12794 | 25,5602 | 6 | 6 |
| 26,74689 | 26,02081 | 25,28263 | 26,07149 | 25,0739 | 25,32526 | 4 | 4 |
| 24,96504 | 25,02416 | 25,10973 | 25,18741 | 25,03658 | 25,22107 | 2 | 2 |
| 26,9556 | 26,80255 | 26,10711 | 27,11396 | 27,18064 | 27,23498 | 7 | 7 |
| 29,81508 | 29,69385 | 29,41764 | 29,57423 | 29,80761 | 29,60895 | 14 | 14 |
| 24,32045 | 26,90323 | 26,58319 | 25,82479 | 25,5168 | 25,89291 | 7 | 7 |
| 25,92461 | 25,77034 | 25,33909 | 25,65428 | 25,28185 | 25,70205 | 4 | 4 |
| 28,68759 | 28,57924 | 28,15485 | 28,27515 | 28,41885 | 28,14953 | 12 | 12 |
| 24,54663 | 24,86325 | 23,30605 | 25,35205 | 25,10829 | 24,87084 | 2 | 2 |
| 27,49098 | 27,19779 | 28,28586 | 28,4654 | 27,98685 | 28,92167 | 13 | 13 |

| | | | | | | | |
|---|---|---|---|---|---|---|---|
| 28,57694 | 28,57073 | 28,50714 | 28,42926 | 28,75656 | 27,67512 | 11 | 3 |
| 23,77595 | 24,58852 | 24,27711 | 24,38495 | 24,26771 | 24,42297 | 17 | 2 |
| 22,72315 | 23,44063 | 25,55549 | 23,51797 | 24,30285 | 23,63389 | 3 | 3 |

| | | Student's *t* -t |
|---|---|---|
| **Sequence coverage [%]** | **Intensity (natural scale)** | **p-value (-log$_{10}$ scale)** |
| 525 | 1,14E+10 | 4,774100131 |
| 381 | 2,63E+10 | 1,309304225 |
| 308 | 1,67E+10 | 4,754444436 |
| 495 | 4,17E+09 | 2,367237855 |
| 156 | 1,05E+09 | 2,417759161 |
| 63 | 1,20E+09 | 3,465208624 |
| 344 | 1,60E+09 | 2,610867252 |
| 309 | 2,04E+09 | 3,519086027 |
| 151 | 1,78E+09 | 2,199139963 |
| 93 | 4,88E+08 | 1,580078148 |
| 146 | 1,32E+09 | 2,42935673 |
| 311 | 3,98E+08 | 2,642606905 |
| 32 | 9,35E+07 | 3,551277598 |
| 117 | 5,52E+08 | 3,962028154 |
| 8 | 4,62E+08 | 1,710565654 |
| 32 | 3,87E+07 | 1,515420454 |
| 28 | 2,12E+07 | 1,70983972 |
| 46 | 6,51E+07 | 1,338753669 |
| 54 | 1,77E+08 | 2,788508715 |
| 71 | 2,36E+08 | 2,79168912 |
| 353 | 7,15E+07 | 1,477324674 |
| 251 | 4,73E+08 | 1,449187049 |

| | | Student's *t* -t |
|---|---|---|
| **Sequence coverage [%]** | **Intensity (natural scale)** | **p-value (-log$_{10}$ scale)** |
| 322 | 3,14E+10 | 4,578156979 |
| 534 | 6,15E+09 | 2,430975953 |
| 352 | 2,44E+10 | 3,231123588 |
| 162 | 1,29E+09 | 2,483191555 |
| 241 | 5,31E+09 | 3,414644862 |
| 278 | 5,08E+09 | 2,93351579 |
| 185 | 2,57E+09 | 2,861517493 |
| 227 | 5,05E+09 | 3,862203037 |
| 234 | 7,47E+09 | 3,655238155 |
| 166 | 1,11E+09 | 3,39899754 |
| 251 | 2,50E+09 | 2,93939179 |
| 467 | 9,54E+09 | 2,533099932 |
| 176 | 9,44E+08 | 3,951819116 |
| 284 | 2,76E+09 | 2,484508427 |
| 26 | 3,15E+09 | 2,155043965 |
| 307 | 3,92E+09 | 1,801449309 |
| 208 | 3,18E+08 | 2,157618344 |
| 357 | 3,66E+08 | 2,291580028 |
| 22 | 3,64E+08 | 1,911334721 |
| 175 | 2,41E+09 | 2,812911654 |
| 44 | 7,14E+08 | 1,992557546 |
| 149 | 3,96E+08 | 2,13199412 |

| | | |
|---|---|---|
| 82 | 1,29E+09 | 3,119680666 |
| 167 | 3,51E+09 | 3,290826774 |
| 87 | 1,68E+08 | 1,888474501 |
| 57 | 2,84E+08 | 2,253662115 |
| 356 | 8,45E+09 | 3,102192954 |
| 119 | 4,71E+08 | 1,452962412 |
| 196 | 3,89E+08 | 2,158363384 |
| 193 | 7,38E+08 | 3,331416161 |
| 66 | 1,22E+08 | 3,126231675 |
| 8 | 1,42E+08 | 2,807179936 |
| 85 | 5,60E+07 | 1,329821995 |
| 255 | 5,78E+08 | 2,445977152 |
| 95 | 1,53E+08 | 2,419824958 |
| 143 | 8,02E+08 | 2,834230061 |
| 52 | 3,81E+08 | 4,056476813 |
| 49 | 9,21E+07 | 1,380096146 |
| 18 | 1,40E+08 | 1,490978449 |
| 187 | 8,73E+08 | 2,523861088 |
| 17 | 5,77E+07 | 3,507811313 |
| 45 | 2,24E+08 | 1,550942335 |
| 104 | 2,62E+08 | 1,890667295 |
| 179 | 1,79E+09 | 3,425240578 |
| 96 | 1,33E+09 | 2,87919035 |
| 45 | 8,63E+07 | 1,708627788 |
| 36 | 8,81E+07 | 2,021995705 |
| 43 | 7,35E+07 | 1,959985222 |
| 39 | 9,87E+07 | 3,201645343 |
| 11 | 1,91E+07 | 1,753163369 |
| 173 | 4,77E+07 | 1,829762887 |
| 327 | 1,02E+08 | 4,530656343 |
| 127 | 3,99E+07 | 3,401668764 |

| | | **Student's *t*-t** |
|---|---|---|
| **Sequence coverage [%]** | **Intensity (natural scale)** | **p-value (-$\log_{10}$ scale)** |
| 606 | 8,78E+11 | 1,217116466 |
| 852 | 3,45E+12 | 0,832747562 |
| 784 | 4,37E+12 | 2,418004469 |
| 519 | 6,43E+09 | 1,017264019 |
| 776 | 7,97E+07 | 0,63509358 |
| 915 | 6,63E+12 | 2,210645616 |
| 225 | 5,04E+09 | 2,276123474 |
| 101 | 2,93E+09 | 1,452783792 |
| 45 | 2,03E+08 | 1,851998015 |
| 25 | 3,65E+10 | 3,762491172 |
| 74 | 1,58E+08 | 1,620000128 |
| 28 | 5,67E+08 | 1,39585642 |
| 138 | 9,80E+07 | 1,333590678 |
| 237 | 3,20E+09 | 1,814193955 |
| 121 | 9,60E+08 | 2,195779496 |
| 171 | 2,36E+09 | 2,183249785 |

| | | |
|---|---|---|
| 32 | 6,95E+09 | 2,06584944 |
| 156 | 1,19E+09 | 2,516291083 |
| 311 | 1,03E+10 | 1,484532227 |
| 114 | 7,50E+08 | 1,877868342 |
| 152 | 4,51E+09 | 1,445472976 |
| 352 | 5,10E+09 | 1,943414074 |
| 91 | 1,09E+09 | 2,469187507 |
| 354 | 1,03E+10 | 2,729541821 |
| 249 | 3,50E+09 | 2,147060226 |
| 204 | 5,41E+08 | 1,568689728 |
| 196 | 3,09E+09 | 2,030444502 |
| 38 | 2,19E+08 | 1,372039911 |
| 394 | 1,70E+10 | 2,468147779 |
| 141 | 8,26E+08 | 2,244109218 |
| 8 | 4,16E+08 | 1,789876291 |
| 243 | 4,55E+09 | 2,62224066 |
| 81 | 1,97E+08 | 1,401940406 |
| 244 | 2,21E+09 | 1,854633359 |
| 292 | 1,08E+10 | 2,482939101 |
| 72 | 8,20E+08 | 1,311468436 |
| 229 | 8,52E+08 | 1,97427778 |
| 108 | 8,93E+08 | 2,236554173 |
| 366 | 3,29E+10 | 2,700634793 |
| 264 | 1,20E+09 | 2,172804554 |
| 248 | 2,15E+09 | 2,3760458 |
| 49 | 4,61E+10 | 2,117136452 |
| 583 | 1,22E+10 | 2,139875011 |
| 211 | 3,12E+09 | 1,649746934 |
| 158 | 3,61E+10 | 1,309880369 |
| 129 | 1,57E+09 | 3,552960794 |
| 37 | 1,68E+08 | 1,513318742 |
| 366 | 7,74E+09 | 1,991227521 |
| 71 | 1,04E+08 | 1,394675958 |
| 76 | 4,55E+08 | 1,384180173 |
| 192 | 1,40E+09 | 1,3147615 |
| 336 | 1,87E+09 | 1,841133648 |
| 79 | 3,01E+08 | 1,400207754 |
| 10 | 4,22E+08 | 1,391558806 |
| 241 | 3,26E+09 | 2,058689661 |
| 458 | 2,75E+10 | 2,156823968 |
| 49 | 1,65E+08 | 1,824659527 |
| 96 | 1,13E+09 | 1,787277134 |
| 211 | 7,76E+08 | 1,350069248 |
| 663 | 1,19E+11 | 1,311469957 |
| 53 | 1,70E+09 | 1,682031941 |
| 41 | 9,24E+07 | 1,769058396 |
| 17 | 9,63E+09 | 3,18822918 |
| 58 | 2,78E+08 | 1,79897276 |
| 277 | 3,50E+09 | 1,897078955 |
| 267 | 3,02E+09 | 2,35292096 |

| | | |
|---|---|---|
| 688 | 2,48E+10 | 1,451891321 |
| 135 | 1,96E+09 | 1,463113821 |
| 164 | 4,92E+08 | 2,308821151 |
| 195 | 6,85E+08 | 1,899963109 |
| 672 | 5,69E+10 | 1,41282732 |
| 32 | 2,07E+10 | 3,345828929 |
| 392 | 3,37E+08 | 1,546251149 |
| 277 | 1,45E+09 | 1,481244774 |
| 40 | 9,95E+09 | 2,426982339 |
| 254 | 4,02E+09 | 3,130636089 |
| 354 | 3,54E+09 | 2,075332762 |
| 308 | 7,21E+08 | 1,472862275 |
| 141 | 3,09E+09 | 1,560009351 |
| 112 | 6,79E+08 | 1,766652801 |
| 748 | 2,10E+10 | 1,982836941 |
| 29 | 7,90E+07 | 1,58429404 |
| 33 | 1,08E+08 | 3,945199699 |
| 392 | 1,28E+10 | 1,425157227 |
| 195 | 1,18E+09 | 2,275411228 |
| 85 | 5,07E+08 | 2,019837157 |
| 212 | 1,75E+09 | 2,582519658 |
| 291 | 2,01E+09 | 1,3465564 |
| 189 | 6,77E+09 | 2,79569954 |
| 329 | 2,46E+09 | 3,155748441 |
| 137 | 5,44E+08 | 3,642292382 |
| 517 | 6,92E+09 | 1,582839168 |
| 35 | 7,23E+09 | 1,840858263 |
| 248 | 5,18E+09 | 1,565074842 |
| 56 | 1,25E+08 | 1,522976456 |
| 262 | 1,94E+09 | 1,456243804 |
| 15 | 9,17E+07 | 2,172794817 |
| 20 | 4,56E+09 | 2,555614841 |
| 187 | 1,95E+09 | 1,736866711 |
| 309 | 3,61E+09 | 1,315565407 |
| 372 | 1,37E+10 | 1,76136095 |
| 517 | 2,27E+11 | 1,778083841 |
| 314 | 4,63E+10 | 1,482386901 |
| 362 | 1,30E+10 | 4,986952365 |
| 165 | 2,03E+10 | 1,336287275 |
| 524 | 4,73E+10 | 2,093748906 |
| 301 | 8,28E+09 | 2,276322715 |
| 399 | 4,24E+09 | 2,550865202 |
| 47 | 6,10E+10 | 1,450227714 |
| 734 | 6,26E+10 | 1,463236896 |
| 229 | 8,09E+08 | 2,672583787 |
| 38 | 3,86E+08 | 1,622275003 |
| 103 | 4,31E+08 | 2,557551876 |
| 54 | 1,62E+08 | 2,236821812 |
| 89 | 3,91E+08 | 1,339975255 |
| 61 | 3,33E+08 | 1,948213132 |

| | | |
|---|---|---|
| 255 | 4,51E+09 | 1,489736139 |
| 164 | 1,35E+09 | 1,904773212 |
| 243 | 1,03E+10 | 2,243384311 |
| 526 | 2,13E+09 | 1,467281581 |
| 564 | 6,52E+09 | 2,56303407 |
| 127 | 5,13E+08 | 2,106884852 |
| 365 | 4,41E+09 | 2,637057191 |
| 161 | 1,60E+08 | 1,677351885 |
| 139 | 6,81E+09 | 1,447888954 |
| 349 | 4,43E+09 | 1,932971123 |
| 181 | 9,60E+08 | 2,583508928 |
| 173 | 7,28E+08 | 2,19381727 |
| 113 | 2,44E+08 | 1,937783802 |
| 62 | 3,40E+08 | 1,388311822 |
| 261 | 1,65E+09 | 1,623549856 |
| 206 | 6,26E+09 | 1,862838921 |
| 605 | 4,56E+08 | 1,421237164 |
| 209 | 1,27E+10 | 1,849333385 |
| 411 | 1,70E+10 | 2,625563002 |
| 726 | 1,62E+10 | 1,426205511 |
| 57 | 5,94E+08 | 1,443118107 |
| 484 | 6,58E+10 | 1,917251567 |
| 93 | 1,62E+09 | 2,844274849 |
| 151 | 1,30E+09 | 1,426087315 |
| 258 | 1,25E+10 | 2,619031482 |
| 513 | 4,93E+09 | 1,80109072 |
| 212 | 7,67E+09 | 2,045149211 |
| 208 | 7,42E+08 | 1,310146132 |
| 401 | 7,40E+10 | 2,206022214 |
| 201 | 1,48E+09 | 2,50712962 |
| 337 | 1,33E+10 | 2,997664105 |
| 309 | 9,74E+09 | 2,048389943 |
| 293 | 8,05E+09 | 1,364209116 |
| 459 | 1,53E+10 | 2,180928527 |
| 393 | 8,17E+08 | 1,421545322 |
| 261 | 1,62E+10 | 2,921279837 |
| 242 | 1,37E+10 | 1,704971796 |
| 206 | 2,81E+09 | 2,426608914 |
| 224 | 1,53E+09 | 1,370630736 |
| 113 | 1,91E+09 | 2,046356201 |
| 125 | 1,55E+09 | 1,842278973 |
| 378 | 5,63E+09 | 1,674561801 |
| 67 | 1,16E+08 | 2,554666228 |
| 706 | 1,57E+11 | 3,219415253 |
| 293 | 5,49E+09 | 1,476854007 |
| 43 | 2,10E+08 | 1,839305944 |
| 55 | 1,06E+09 | 1,467043298 |
| 276 | 1,51E+09 | 2,438156414 |
| 106 | 2,62E+08 | 1,890223444 |
| 419 | 9,95E+09 | 3,095447823 |

| | | |
|---|---|---|
| 159 | 8,28E+08 | 2,293001989 |
| 66 | 1,70E+08 | 1,382144351 |
| 103 | 4,57E+08 | 1,751506434 |
| 234 | 1,65E+10 | 3,306792882 |
| 205 | 4,35E+09 | 2,68103216 |
| 153 | 3,28E+08 | 1,32377721 |
| 36 | 1,93E+08 | 1,975098057 |
| 183 | 8,12E+08 | 2,149840835 |
| 211 | 2,62E+09 | 2,863384766 |
| 128 | 2,70E+08 | 3,069785974 |
| 147 | 1,78E+09 | 1,914240273 |
| 366 | 3,95E+09 | 1,913382908 |
| 9 | 1,31E+09 | 2,056990521 |
| 195 | 2,44E+09 | 2,011821014 |
| 571 | 2,52E+10 | 2,286117708 |
| 66 | 1,47E+08 | 1,434436008 |
| 82 | 1,61E+09 | 1,932525643 |
| 582 | 2,78E+10 | 1,571386285 |
| 87 | 1,04E+09 | 2,083310456 |
| 45 | 2,08E+09 | 1,901498652 |
| 205 | 6,68E+08 | 1,910663352 |
| 538 | 3,13E+09 | 2,343496426 |
| 273 | 2,10E+09 | 1,42664651 |
| 189 | 8,28E+08 | 2,607118091 |
| 195 | 6,23E+09 | 2,121196629 |
| 186 | 7,70E+09 | 2,2083869 |
| 118 | 4,51E+09 | 2,253811438 |
| 51 | 2,95E+08 | 2,097601675 |
| 218 | 1,74E+09 | 1,448870558 |
| 487 | 8,06E+10 | 1,32647213 |
| 678 | 3,68E+11 | 1,522395522 |
| 175 | 5,12E+08 | 1,622647284 |
| 88 | 1,05E+09 | 1,531059652 |
| 93 | 8,15E+08 | 1,761117546 |
| 254 | 1,17E+09 | 1,342553177 |
| 21 | 4,51E+08 | 1,742933746 |
| 437 | 2,04E+10 | 2,69303976 |
| 268 | 1,09E+10 | 2,211048472 |
| 105 | 6,82E+08 | 2,017822186 |
| 245 | 9,07E+09 | 2,12386012 |
| 40 | 2,11E+09 | 2,042510692 |
| 165 | 1,93E+09 | 1,886533076 |
| 293 | 9,76E+09 | 1,965748243 |
| 46 | 1,97E+08 | 1,495132505 |
| 168 | 8,47E+08 | 1,324581714 |
| 275 | 2,92E+09 | 1,63820714 |
| 123 | 8,03E+08 | 1,589575096 |
| 12 | 5,60E+08 | 1,629128035 |
| 424 | 6,00E+09 | 1,572514976 |
| 33 | 1,08E+09 | 2,307109902 |

| | | |
|---|---|---|
| 247 | 4,06E+09 | 2,329168836 |
| 46 | 5,52E+08 | 1,311670708 |
| 125 | 2,28E+08 | 1,641946646 |
| 85 | 2,00E+09 | 2,080931308 |
| 223 | 4,29E+09 | 2,14012446 |
| 314 | 2,61E+09 | 1,409627283 |
| 505 | 2,35E+09 | 1,910927983 |
| 212 | 5,13E+09 | 2,402427177 |
| 147 | 3,84E+08 | 1,686745087 |
| 243 | 6,11E+09 | 1,404229059 |
| 67 | 4,32E+08 | 1,545691616 |
| 31 | 1,20E+08 | 1,390212298 |
| 286 | 8,04E+09 | 2,506472662 |
| 237 | 9,77E+09 | 1,701334548 |
| 149 | 2,04E+09 | 3,603480132 |
| 193 | 2,94E+09 | 2,29579452 |
| 198 | 2,86E+09 | 3,100915312 |
| 285 | 5,90E+09 | 1,482820177 |
| 39 | 4,13E+08 | 1,382815526 |
| 262 | 5,62E+09 | 2,008379983 |
| 83 | 1,99E+08 | 1,78398083 |
| 106 | 1,63E+09 | 1,709839546 |
| 537 | 2,01E+10 | 1,814848395 |
| 163 | 2,01E+09 | 2,667290449 |
| 382 | 3,24E+10 | 1,50490527 |
| 8 | 1,11E+09 | 2,457552481 |
| 218 | 5,48E+08 | 1,336637135 |
| 464 | 1,61E+09 | 1,365470855 |
| 231 | 1,27E+08 | 1,530152161 |
| 343 | 2,53E+09 | 2,004703353 |
| 271 | 1,18E+09 | 1,393162694 |
| 428 | 4,41E+09 | 2,66139249 |
| 9 | 4,09E+08 | 1,647813961 |
| 205 | 1,29E+09 | 1,315951532 |
| 177 | 1,80E+09 | 2,336564347 |
| 69 | 1,20E+08 | 1,319010315 |
| 263 | 3,11E+09 | 2,795956321 |
| 289 | 3,36E+09 | 2,35390325 |
| 206 | 5,04E+09 | 3,405069788 |
| 426 | 2,80E+07 | 1,290462967 |
| 83 | 7,19E+07 | 1,087260311 |
| 71 | 3,66E+08 | 0,699974054 |
| 21 | 9,79E+07 | 0,434062464 |
| 4 | 1,48E+08 | 0,386660335 |
| 18 | 6,88E+07 | 0,839588666 |
| 61 | 2,04E+08 | 0,642628963 |
| 74 | 1,76E+08 | 1,10460776 |
| 47 | 1,66E+08 | 0,507424967 |
| 196 | 1,46E+08 | 0,793097295 |
| 8 | 2,64E+08 | 0,531605501 |

| | | |
|---|---|---|
| 91 | 1,31E+08 | 0,713247162 |
| 141 | 2,09E+08 | 0,990054422 |
| 115 | 5,91E+08 | 1,059685448 |
| 69 | 5,86E+08 | 0,570320167 |
| 164 | 1,97E+08 | 0,830035949 |
| 46 | 2,39E+08 | 0,667026696 |
| 181 | 1,54E+08 | 0,763772993 |
| 96 | 1,53E+08 | 0,758165212 |
| 69 | 3,58E+08 | 0,615689145 |
| 132 | 4,93E+08 | 0,983862218 |
| 75 | 1,31E+08 | 1,224750258 |
| 114 | 2,28E+08 | 1,182701208 |
| 11 | 2,10E+09 | 1,040446168 |
| 44 | 1,58E+08 | 0,61825734 |
| 14 | 4,15E+07 | 1,085583796 |
| 276 | 7,87E+08 | 1,08103488 |
| 21 | 1,09E+11 | 0,33025487 |
| 39 | 2,11E+08 | 0,421829793 |
| 21 | 2,08E+08 | 0,476235135 |
| 39 | 1,46E+08 | 1,032266296 |
| 97 | 1,16E+08 | 0,590176559 |
| 251 | 1,02E+09 | 1,151845217 |
| 21 | 6,93E+08 | 0,797985308 |
| 135 | 2,76E+09 | 1,255352842 |
| 81 | 9,07E+07 | 0,599254115 |
| 47 | 1,25E+08 | 1,138077778 |
| 163 | 1,69E+08 | 0,382809421 |
| 55 | 1,80E+08 | 0,599938443 |
| 103 | 2,15E+08 | 0,331603128 |
| 73 | 4,07E+08 | 1,263984107 |
| 51 | 6,96E+07 | 1,120007845 |
| 68 | 2,60E+08 | 0,975268145 |
| 57 | 3,27E+08 | 0,937227815 |
| 71 | 2,70E+08 | 0,44504201 |
| 177 | 8,33E+08 | 0,902576899 |
| 4 | 7,76E+07 | 0,920761353 |
| 37 | 7,53E+07 | 1,049721489 |
| 53 | 8,62E+08 | 0,520057583 |
| 712 | 6,18E+07 | 0,713394232 |
| 23 | 1,69E+08 | 0,708111959 |
| 61 | 3,13E+08 | 0,686813677 |
| 505 | 1,84E+08 | 1,09278555 |
| 143 | 1,18E+08 | 0,635071011 |
| 4 | 2,36E+08 | 0,401510518 |
| 58 | 4,53E+08 | 0,575402644 |
| 2 | 3,56E+07 | 0,581964444 |
| 6 | 3,95E+08 | 0,167823977 |
| 36 | 7,26E+07 | 0,979299476 |
| 33 | 2,12E+08 | 0,791311873 |
| 95 | 7,43E+07 | 0,662753361 |

| | | |
|---|---|---|
| 38 | 8,36E+07 | 0,785118878 |
| 29 | 6,26E+07 | 0,529540281 |
| 63 | 2,19E+08 | 0,651036942 |
| 48 | 1,23E+08 | 0,465890187 |
| 111 | 4,24E+07 | 0,589465106 |
| 208 | 1,45E+08 | 0,824804522 |
| 51 | 2,75E+08 | 0,601117679 |
| 32 | 1,15E+08 | 0,863327675 |
| 34 | 3,47E+07 | 0,918025396 |
| 18 | 1,78E+07 | 0,624893468 |
| 82 | 2,65E+08 | 0,748189937 |
| 73 | 2,24E+08 | 1,233186133 |
| 508 | 4,16E+08 | 0,632784759 |
| 88 | 7,14E+08 | 0,099367765 |
| 44 | 7,38E+07 | 0,116540526 |
| 61 | 6,87E+08 | 0,169460322 |
| 436 | 1,53E+10 | 0,689085503 |
| 23 | 7,53E+08 | 0,314001983 |
| 56 | 9,14E+08 | 0,208566977 |
| 42 | 6,89E+07 | 0,172237789 |
| 168 | 8,65E+08 | 0,684742814 |
| 118 | 5,12E+08 | 0,004924978 |
| 105 | 2,81E+09 | 0,415961691 |
| 62 | 3,67E+08 | 0,061026737 |
| 18 | 1,14E+08 | 0,008123723 |
| 49 | 2,30E+08 | 0,285117813 |
| 71 | 2,95E+08 | 0,334588562 |
| 37 | 1,42E+09 | 0,009804175 |
| 247 | 1,60E+09 | 0,101341262 |
| 5 | 8,31E+08 | 1,002726869 |
| 58 | 1,60E+08 | 0,709709624 |
| 295 | 9,34E+08 | 0,37251256 |
| 122 | 5,75E+08 | 0,569952222 |
| 228 | 3,48E+08 | 0,075190301 |
| 61 | 9,78E+07 | 0,3207374 |
| 24 | 2,84E+07 | 0,15546291 |
| 49 | 8,79E+08 | 0,349894561 |
| 37 | 8,53E+07 | 0,174090447 |
| 22 | 4,47E+07 | 0,26244758 |
| 64 | 9,47E+08 | 0,956659723 |
| 197 | 1,45E+09 | 0,778250916 |
| 129 | 5,51E+08 | 0,225500822 |
| 213 | 1,82E+09 | 0,487791789 |
| 67 | 2,55E+08 | 0,62941064 |
| 492 | 2,40E+10 | 0,175258396 |
| 47 | 2,87E+08 | 0,532561681 |
| 1 | 1,80E+07 | 0,702869532 |
| 121 | 9,42E+08 | 0,498409626 |
| 235 | 1,05E+09 | 1,264658073 |
| 36 | 4,40E+08 | 0,244394493 |

| | | |
|---|---|---|
| 47 | 1,18E+08 | 0,035881171 |
| 58 | 4,18E+08 | 0,962062868 |
| 35 | 2,16E+09 | 0,718921915 |
| 33 | 7,22E+07 | 0,834905056 |
| 287 | 6,32E+09 | 0,212206852 |
| 231 | 1,31E+09 | 0,250035207 |
| 49 | 1,60E+08 | 0,196888759 |
| 12 | 1,14E+08 | 0,165381146 |
| 74 | 2,67E+08 | 0,307973315 |
| 388 | 2,13E+09 | 0,683647042 |
| 72 | 1,53E+08 | 1,287581417 |
| 16 | 1,08E+08 | 0,027572316 |
| 36 | 1,71E+08 | 0,332785405 |
| 35 | 4,93E+09 | 1,138117806 |
| 379 | 1,92E+10 | 0,06908789 |
| 461 | 1,01E+11 | 0,586178801 |
| 377 | 1,02E+10 | 0,339926636 |
| 41 | 3,54E+07 | 0,264779505 |
| 155 | 5,09E+08 | 0,503747051 |
| 72 | 7,26E+08 | 0,12072348 |
| 139 | 1,09E+08 | 0,014708816 |
| 207 | 3,47E+09 | 0,560601704 |
| 347 | 2,24E+10 | 0,611101242 |
| 191 | 2,45E+09 | 0,292049523 |
| 178 | 2,78E+08 | 1,283317926 |
| 387 | 4,98E+09 | 0,121743836 |
| 296 | 1,59E+09 | 0,809597901 |
| 442 | 4,42E+09 | 1,068508952 |
| 59 | 2,00E+08 | 0,213850261 |
| 2 | 1,11E+08 | 0,706613383 |
| 314 | 1,81E+10 | 0,193178706 |
| 22 | 4,16E+07 | 0,48416644 |
| 254 | 9,86E+08 | 0,82845363 |
| 208 | 4,56E+08 | 0,180010274 |
| 389 | 2,52E+07 | 1,034675469 |
| 48 | 1,71E+08 | 0,723209641 |
| 432 | 3,04E+09 | 0,523359008 |
| 235 | 2,59E+09 | 0,067507803 |
| 1 | 8,85E+07 | 0,15158181 |
| 153 | 5,73E+08 | 0,827416937 |
| 316 | 4,04E+08 | 0,325652251 |
| 148 | 8,25E+07 | 0,223553765 |
| 244 | 6,04E+09 | 0,096941264 |
| 473 | 2,15E+09 | 0,141024668 |
| 73 | 3,58E+08 | 0,910110478 |
| 192 | 5,45E+08 | 0,115347836 |
| 162 | 1,77E+09 | 0,013034201 |
| 52 | 1,05E+08 | 0,412962479 |
| 292 | 2,29E+09 | 0,072275998 |
| 18 | 2,02E+07 | 0,54278614 |

| | | |
|---|---|---|
| 141 | 2,13E+09 | 1,058073383 |
| 63 | 2,81E+08 | 0,833831826 |
| 96 | 6,71E+08 | 0,612377746 |
| 124 | 4,68E+08 | 0,11947454 |
| 47 | 2,77E+08 | 0,16328011 |
| 85 | 3,34E+08 | 0,597635488 |
| 169 | 8,47E+08 | 0,613943291 |
| 67 | 8,19E+07 | 1,000671904 |
| 127 | 6,39E+08 | 0,606958095 |
| 159 | 1,80E+09 | 0,56692698 |
| 21 | 8,98E+07 | 0,231735602 |
| 58 | 2,17E+08 | 0,060290653 |
| 228 | 3,20E+08 | 0,104051184 |
| 223 | 1,59E+08 | 0,394291614 |
| 75 | 1,83E+08 | 0,000903045 |
| 3 | 2,12E+08 | 0,223938882 |
| 10 | 3,70E+08 | 1,249258713 |
| 65 | 3,79E+08 | 0,072551176 |
| 44 | 5,94E+07 | 0,49279748 |
| 19 | 5,33E+07 | 0,162859072 |
| 77 | 4,01E+08 | 0,697449446 |
| 113 | 1,95E+08 | 0,162670752 |
| 10 | 4,69E+08 | 0,001741409 |
| 161 | 2,35E+07 | 0,056393768 |
| 66 | 2,20E+08 | 0,006894076 |
| 49 | 3,25E+08 | 0,258822652 |
| 173 | 3,58E+08 | 0,748223895 |
| 25 | 1,48E+08 | 0,36094561 |
| 71 | 1,09E+09 | 0,052826671 |
| 5 | 5,35E+08 | 0,353856831 |
| 98 | 3,53E+08 | 0,815701469 |
| 487 | 8,42E+09 | 1,162521222 |
| 291 | 1,99E+09 | 0,642245871 |
| 241 | 1,72E+09 | 0,025375526 |
| 52 | 2,64E+08 | 0,089609991 |
| 203 | 1,58E+09 | 0,253724644 |
| 305 | 3,35E+08 | 0,58863299 |
| 27 | 5,86E+07 | 0,187085571 |
| 205 | 1,01E+09 | 0,906844253 |
| 208 | 1,84E+09 | 1,125849657 |
| 54 | 3,80E+08 | 0,169563126 |
| 133 | 1,68E+08 | 0,475358654 |
| 305 | 9,02E+07 | 0,15945826 |
| 62 | 5,80E+08 | 0,124853936 |
| 54 | 6,95E+08 | 0,713518396 |
| 217 | 2,42E+10 | 0,968406856 |
| 31 | 2,41E+07 | 0,278532584 |
| 106 | 3,23E+08 | 0,541913529 |
| 177 | 2,70E+08 | 0,66411741 |

| | | |
|---|---|---|
| 32 | 2,69E+10 | 0,512257255 |
| 106 | 2,83E+08 | 0,015104359 |
| 173 | 1,90E+08 | 0,726497629 |
| 39 | 2,58E+08 | 0,931411786 |
| 355 | 7,28E+09 | 1,247574891 |
| 92 | 2,80E+08 | 0,174476325 |
| 18 | 2,54E+09 | 0,000990296 |
| 127 | 9,81E+08 | 0,000851687 |
| 62 | 5,62E+08 | 0,246558361 |
| 198 | 4,81E+08 | 0,891736052 |
| 29 | 2,64E+08 | 0,678328304 |
| 62 | 2,19E+09 | 0,292561432 |
| 55 | 1,02E+08 | 0,097167546 |
| 98 | 1,82E+08 | 0,453672313 |
| 25 | 1,72E+08 | 0,08024622 |
| 367 | 7,93E+09 | 1,099190348 |
| 87 | 1,88E+08 | 0,563695225 |
| 109 | 1,52E+09 | 0,035475158 |
| 408 | 7,68E+09 | 0,585265613 |
| 54 | 3,11E+08 | 0,181581135 |
| 49 | 1,47E+08 | 0,289157631 |
| 277 | 2,84E+09 | 0,896914109 |
| 119 | 4,70E+08 | 0,867044484 |
| 18 | 1,56E+08 | 0,047277188 |
| 261 | 4,51E+09 | 0,082487859 |
| 123 | 2,50E+08 | 1,189543506 |
| 16 | 9,32E+07 | 0,316434762 |
| 174 | 5,13E+09 | 0,208762669 |
| 321 | 1,58E+10 | 1,140505211 |
| 19 | 7,31E+08 | 0,11273549 |
| 61 | 5,38E+08 | 1,276749764 |
| 79 | 9,07E+08 | 0,558335254 |
| 358 | 4,70E+09 | 0,66129805 |
| 276 | 9,03E+08 | 0,32484354 |
| 44 | 1,32E+08 | 0,508196494 |
| 51 | 9,32E+07 | 0,80550309 |
| 98 | 1,00E+09 | 0,197139584 |
| 114 | 4,41E+08 | 0,832813211 |
| 216 | 9,21E+08 | 0,840073928 |
| 183 | 3,36E+09 | 0,666241151 |
| 289 | 2,13E+08 | 0,173431463 |
| 52 | 6,69E+07 | 0,077446363 |
| 141 | 1,25E+09 | 1,195848958 |
| 529 | 2,36E+11 | 0,91670768 |
| 48 | 1,63E+09 | 0,109191928 |
| 674 | 3,86E+11 | 0,218467673 |
| 136 | 7,75E+07 | 0,919985244 |
| 164 | 1,68E+09 | 0,214639755 |
| 16 | 1,31E+09 | 1,007299985 |
| 626 | 8,63E+09 | 0,665465838 |

| | | |
|---|---|---|
| 61 | 9,96E+07 | 0,052112799 |
| 113 | 3,86E+08 | 0,010337746 |
| 108 | 4,91E+08 | 0,068826855 |
| 11 | 5,15E+08 | 0,484867236 |
| 5 | 2,01E+08 | 0,79732636 |
| 41 | 5,89E+07 | 0,10011912 |
| 176 | 1,03E+09 | 1,20344717 |
| 216 | 2,83E+09 | 0,891139081 |
| 152 | 5,04E+08 | 0,201501493 |
| 357 | 8,81E+09 | 0,706112451 |
| 137 | 4,20E+08 | 0,277884936 |
| 261 | 2,85E+09 | 0,757604972 |
| 214 | 2,00E+08 | 0,299679282 |
| 296 | 1,69E+09 | 0,453193288 |
| 306 | 1,95E+09 | 0,654777996 |
| 7 | 3,16E+08 | 0,296398857 |
| 248 | 1,32E+09 | 0,094139119 |
| 93 | 2,46E+08 | 0,776084587 |
| 62 | 1,46E+08 | 0,186279098 |
| 236 | 4,70E+08 | 0,649916838 |
| 77 | 1,17E+09 | 0,524535716 |
| 49 | 4,29E+07 | 0,293338647 |
| 213 | 3,41E+09 | 1,227503914 |
| 153 | 1,60E+09 | 0,363767705 |
| 8 | 3,03E+08 | 0,633590957 |
| 137 | 2,57E+08 | 0,512202547 |
| 559 | 1,86E+09 | 1,052078842 |
| 237 | 1,35E+09 | 0,762202212 |
| 41 | 2,00E+08 | 0,308249673 |
| 28 | 3,74E+08 | 0,668463239 |
| 22 | 2,28E+07 | 0,094244302 |
| 108 | 1,25E+09 | 0,857578019 |
| 341 | 3,26E+09 | 0,475321748 |
| 86 | 1,32E+08 | 0,912253866 |
| 115 | 2,34E+09 | 1,026851845 |
| 545 | 1,38E+09 | 0,405954142 |
| 532 | 3,81E+07 | 1,202324171 |
| 145 | 3,46E+08 | 0,068235076 |
| 6 | 1,09E+09 | 0,755037674 |
| 75 | 2,72E+09 | 0,168720555 |
| 33 | 2,65E+10 | 0,701783316 |
| 282 | 2,03E+10 | 0,361059756 |
| 49 | 8,46E+09 | 0,368989401 |
| 287 | 2,56E+09 | 1,160489511 |
| 199 | 5,88E+09 | 0,139912844 |
| 25 | 1,97E+09 | 0,671972921 |
| 35 | 5,01E+07 | 0,786626235 |
| 113 | 1,20E+09 | 0,052285504 |
| 256 | 1,70E+10 | 0,394902814 |
| 256 | 3,75E+07 | 0,534760899 |

| | | |
|---|---|---|
| 238 | 8,52E+09 | 1,039171761 |
| 182 | 1,94E+09 | 0,315790118 |
| 126 | 9,31E+08 | 0,07407396 |
| 148 | 2,87E+08 | 0,921687981 |
| 216 | 4,86E+09 | 0,900351004 |
| 216 | 6,49E+08 | 0,940318092 |
| 147 | 2,79E+08 | 1,151725214 |
| 19 | 3,91E+08 | 0,114780357 |
| 427 | 3,18E+09 | 0,772321245 |
| 112 | 1,43E+08 | 0,382419936 |
| 105 | 8,56E+07 | 0,949259547 |
| 28 | 8,74E+07 | 0,350624767 |
| 95 | 2,66E+08 | 1,042482811 |
| 28 | 3,78E+07 | 0,582999049 |
| 84 | 6,75E+08 | 0,110296072 |
| 155 | 1,86E+08 | 0,239607674 |
| 28 | 6,32E+08 | 1,251642267 |
| 165 | 1,24E+09 | 0,372365769 |
| 52 | 2,54E+08 | 0,313418779 |
| 106 | 3,66E+08 | 0,264795959 |
| 154 | 1,27E+09 | 0,363781005 |
| 31 | 1,15E+08 | 1,263930063 |
| 257 | 1,69E+09 | 0,789717349 |
| 98 | 2,76E+08 | 0,7576275 |
| 498 | 5,54E+09 | 0,020780591 |
| 12 | 1,13E+09 | 0,082659951 |
| 63 | 1,22E+08 | 0,24700387 |
| 67 | 2,48E+08 | 0,246811468 |
| 45 | 5,73E+08 | 0,421919651 |
| 329 | 7,32E+08 | 0,420337775 |
| 66 | 4,17E+08 | 0,008262878 |
| 65 | 2,99E+08 | 0,526292861 |
| 108 | 9,57E+08 | 0,100605472 |
| 133 | 2,48E+09 | 0,165034931 |
| 82 | 2,05E+08 | 0,930578063 |
| 97 | 6,96E+08 | 0,646569074 |
| 10 | 9,15E+09 | 0,674565724 |
| 49 | 4,39E+07 | 0,977285291 |
| 107 | 9,47E+08 | 0,0671682 |
| 285 | 2,46E+09 | 0,449105783 |
| 162 | 1,55E+09 | 0,468509422 |
| 134 | 9,47E+08 | 0,037325685 |
| 29 | 1,70E+08 | 0,043206658 |
| 21 | 2,63E+08 | 0,260116574 |
| 143 | 4,00E+09 | 0,289828904 |
| 39 | 1,42E+08 | 0,064567896 |
| 209 | 2,94E+09 | 0,801949023 |
| 27 | 1,94E+10 | 0,19753373 |
| 53 | 2,83E+08 | 1,037916077 |
| 114 | 6,57E+08 | 0,377085739 |

| | | |
|---|---|---|
| 558 | 1,04E+11 | 0,90328015 |
| 33 | 3,84E+08 | 1,114618438 |
| 43 | 1,32E+08 | 0,19392395 |
| 543 | 7,24E+10 | 0,003741156 |
| 2 | 2,77E+08 | 0,296288067 |
| 89 | 3,25E+08 | 0,987584822 |
| 115 | 4,33E+08 | 0,399126439 |
| 153 | 5,95E+08 | 0,39009241 |
| 229 | 1,01E+10 | 1,210787534 |
| 89 | 3,12E+08 | 0,239968127 |
| 33 | 3,75E+08 | 0,156175942 |
| 53 | 3,03E+08 | 0,359049302 |
| 114 | 7,63E+08 | 0,099904198 |
| 79 | 5,70E+08 | 0,622889379 |
| 128 | 9,94E+08 | 1,096750997 |
| 109 | 6,69E+08 | 0,594962109 |
| 59 | 1,68E+08 | 0,36932637 |
| 22 | 1,33E+08 | 0,372699413 |
| 194 | 1,83E+09 | 0,563748284 |
| 306 | 1,03E+09 | 0,011859808 |
| 82 | 6,04E+08 | 0,630580316 |
| 64 | 3,69E+08 | 0,554234126 |
| 21 | 2,79E+09 | 0,883620953 |
| 147 | 1,44E+10 | 0,578737169 |
| 105 | 4,14E+08 | 0,080130148 |
| 154 | 1,97E+09 | 0,508296278 |
| 181 | 2,36E+09 | 0,1042457 |
| 202 | 1,12E+09 | 0,012283685 |
| 51 | 7,15E+10 | 0,950401515 |
| 236 | 1,66E+09 | 0,23683602 |
| 108 | 5,27E+08 | 0,451753676 |
| 192 | 3,49E+08 | 0,000368789 |
| 43 | 9,22E+07 | 0,4206552 |
| 437 | 3,89E+10 | 0,112910019 |
| 154 | 9,13E+08 | 0,138824794 |
| 161 | 2,38E+08 | 0,711057709 |
| 9 | 7,75E+06 | 0,200078805 |
| 181 | 3,11E+09 | 0,859159978 |
| 113 | 7,73E+08 | 0,601697348 |
| 119 | 1,04E+09 | 0,773446059 |
| 74 | 2,57E+08 | 0,098930749 |
| 241 | 1,24E+09 | 1,084220099 |
| 23 | 1,07E+08 | 0,378317898 |
| 15 | 3,55E+08 | 0,420250083 |
| 98 | 6,91E+08 | 0,804423691 |
| 72 | 8,64E+07 | 0,232980231 |
| 585 | 6,54E+09 | 0,653230099 |
| 517 | 9,58E+09 | 0,359819443 |
| 408 | 2,00E+09 | 0,39535828 |
| 319 | 4,32E+08 | 0,315906303 |

| | | |
|---|---|---|
| 42 | 7,19E+07 | 0,484109831 |
| 373 | 1,19E+09 | 0,679192626 |
| 38 | 1,29E+08 | 0,438829546 |
| 209 | 2,81E+09 | 0,30063978 |
| 355 | 3,90E+09 | 0,497936594 |
| 83 | 2,68E+08 | 0,210426369 |
| 434 | 8,63E+09 | 0,499974824 |
| 255 | 4,45E+09 | 0,106843716 |
| 286 | 2,47E+09 | 0,504326902 |
| 168 | 2,17E+09 | 0,248814122 |
| 43 | 7,81E+07 | 0,661478811 |
| 7 | 2,35E+08 | 0,508818748 |
| 101 | 1,29E+08 | 0,262219578 |
| 54 | 3,28E+08 | 0,416197541 |
| 195 | 1,57E+09 | 0,246090088 |
| 99 | 1,41E+08 | 0,119277486 |
| 48 | 4,29E+08 | 0,030629371 |
| 243 | 2,29E+09 | 1,223014501 |
| 28 | 9,27E+07 | 0,326902344 |
| 92 | 3,62E+08 | 0,08655835 |
| 113 | 7,88E+08 | 0,60716861 |
| 63 | 7,58E+07 | 0,083857997 |
| 66 | 1,73E+08 | 0,784774554 |
| 137 | 1,09E+09 | 0,920884951 |
| 101 | 1,42E+08 | 0,857900839 |
| 153 | 9,20E+08 | 0,57187015 |
| 87 | 2,39E+08 | 0,998525881 |
| 382 | 3,40E+10 | 0,616079433 |
| 447 | 1,33E+10 | 0,343040101 |
| 303 | 7,26E+09 | 0,092524458 |
| 704 | 1,25E+10 | 1,102957664 |
| 46 | 1,68E+08 | 0,215149835 |
| 136 | 1,44E+08 | 0,774198081 |
| 553 | 8,32E+10 | 0,111048874 |
| 175 | 1,28E+09 | 0,040960903 |
| 53 | 2,07E+08 | 0,140178811 |
| 91 | 6,64E+08 | 0,223782775 |
| 312 | 1,60E+10 | 0,519555973 |
| 116 | 6,94E+08 | 0,793342156 |
| 237 | 1,72E+09 | 0,578323691 |
| 158 | 9,32E+08 | 0,143214282 |
| 10 | 5,53E+07 | 0,385033736 |
| 75 | 8,93E+08 | 0,167902388 |
| 525 | 1,75E+10 | 1,107874531 |
| 688 | 3,10E+09 | 0,082544252 |
| 688 | 3,74E+10 | 0,772539158 |
| 253 | 9,76E+08 | 0,521660245 |
| 237 | 7,45E+09 | 0,665641602 |
| 482 | 2,00E+10 | 0,924135741 |

| | | |
|---|---|---|
| 303 | 7,79E+09 | 0,880297668 |
| 188 | 1,40E+09 | 0,557655644 |
| 637 | 1,28E+09 | 0,815160065 |
| 139 | 1,31E+09 | 0,669346661 |
| 13 | 4,38E+08 | 0,489123869 |
| 10 | 5,54E+08 | 0,81537156 |
| 116 | 3,28E+07 | 0,109488923 |
| 47 | 1,77E+08 | 0,45572009 |
| 236 | 8,39E+08 | 0,971835638 |
| 68 | 1,40E+09 | 0,619302922 |
| 66 | 3,90E+08 | 0,677667956 |
| 85 | 2,69E+08 | 0,762876355 |
| 45 | 3,38E+08 | 1,173916807 |
| 363 | 1,66E+08 | 0,111931333 |
| 42 | 3,66E+08 | 0,752465774 |
| 244 | 1,76E+08 | 0,002892496 |
| 109 | 2,89E+08 | 0,042397618 |
| 164 | 8,95E+08 | 0,168018568 |
| 246 | 7,22E+09 | 0,160624327 |
| 93 | 3,72E+08 | 0,436776864 |
| 69 | 2,73E+08 | 0,228683609 |
| 241 | 7,86E+08 | 0,353232391 |
| 427 | 2,15E+09 | 0,062704965 |
| 508 | 1,53E+09 | 0,042512074 |
| 33 | 4,95E+09 | 0,263423359 |
| 216 | 1,67E+09 | 0,150595229 |
| 227 | 1,90E+09 | 1,205915756 |
| 105 | 5,25E+08 | 1,058681056 |
| 56 | 3,49E+07 | 0,048050277 |
| 93 | 5,88E+08 | 0,084472606 |
| 286 | 1,34E+09 | 0,187515566 |
| 17 | 2,32E+08 | 0,09509153 |
| 163 | 5,33E+08 | 0,199214489 |
| 104 | 3,92E+08 | 0,667994527 |
| 8 | 8,47E+08 | 1,006721408 |
| 56 | 4,48E+08 | 1,128419053 |
| 29 | 1,00E+08 | 0,564445709 |
| 111 | 1,70E+08 | 0,199906209 |
| 36 | 8,29E+09 | 0,443736998 |
| 36 | 2,00E+09 | 0,377072345 |
| 162 | 1,32E+08 | 0,229808637 |
| 105 | 8,08E+08 | 0,570113305 |
| 98 | 2,85E+08 | 0,034189657 |
| 163 | 1,52E+09 | 0,500291725 |
| 41 | 8,65E+08 | 0,313204545 |
| 64 | 3,24E+08 | 0,476170259 |
| 164 | 1,53E+09 | 1,117417451 |
| 107 | 8,14E+08 | 0,111942968 |
| 125 | 1,64E+08 | 0,475840911 |
| 56 | 1,45E+08 | 0,071682446 |

| | | |
|---|---|---|
| 127 | 3,79E+08 | 0,561040419 |
| 69 | 5,05E+08 | 0,64428776 |
| 19 | 1,50E+09 | 0,104829669 |
| 459 | 2,69E+10 | 0,148888795 |
| 277 | 6,28E+09 | 1,208245984 |
| 21 | 4,49E+08 | 0,265389479 |
| 7 | 1,01E+08 | 0,138766375 |
| 85 | 1,25E+09 | 0,176339519 |
| 136 | 4,26E+08 | 0,396508141 |
| 254 | 1,20E+08 | 0,534697046 |
| 257 | 2,96E+08 | 0,531053481 |
| 13 | 8,02E+08 | 1,040946186 |
| 66 | 2,67E+08 | 0,0746148 |
| 4 | 1,34E+08 | 0,937336544 |
| 37 | 2,86E+08 | 0,078585547 |
| 304 | 1,82E+10 | 0,678141533 |
| 74 | 1,90E+08 | 0,180109755 |
| 144 | 1,27E+09 | 1,13556129 |
| 42 | 1,07E+08 | 0,238067769 |
| 43 | 1,71E+08 | 0,342445312 |
| 84 | 1,34E+09 | 0,866587796 |
| 24 | 1,14E+10 | 0,109118914 |
| 55 | 8,23E+08 | 0,062985604 |
| 55 | 4,06E+08 | 0,381909971 |
| 121 | 1,94E+08 | 0,010413066 |
| 41 | 1,43E+08 | 0,438067197 |
| 63 | 1,83E+08 | 0,612557869 |
| 319 | 4,90E+10 | 1,238804858 |
| 124 | 2,92E+08 | 0,116538017 |
| 73 | 4,63E+08 | 0,535494867 |
| 96 | 1,00E+09 | 0,233582885 |
| 82 | 4,17E+08 | 0,716439746 |
| 35 | 4,84E+08 | 0,351634122 |
| 68 | 1,47E+08 | 0,345705172 |
| 14 | 2,06E+08 | 0,991896595 |
| 105 | 5,52E+08 | 0,085201302 |
| 222 | 8,12E+08 | 0,142651779 |
| 181 | 5,37E+08 | 0,327321863 |
| 383 | 1,08E+10 | 0,714928955 |
| 117 | 1,73E+08 | 0,029141231 |
| 6 | 2,22E+07 | 0,173400592 |
| 106 | 1,19E+08 | 0,930766324 |
| 107 | 2,42E+08 | 0,008954848 |
| 157 | 1,00E+09 | 0,5661269 |
| 424 | 1,68E+09 | 0,943135669 |
| 155 | 7,54E+08 | 0,892912337 |
| 147 | 8,45E+08 | 0,247366721 |
| 491 | 2,16E+09 | 0,45189337 |
| 58 | 2,74E+08 | 0,347311347 |
| 61 | 1,64E+10 | 0,364418178 |

| | | |
|---|---|---|
| 6 | 3,74E+08 | 1,248028046 |
| 221 | 4,03E+08 | 0,007158894 |
| 186 | 1,13E+09 | 0,426535439 |
| 41 | 1,60E+08 | 0,036529624 |
| 102 | 3,89E+08 | 0,616768174 |
| 282 | 1,48E+09 | 0,670469739 |
| 105 | 5,48E+08 | 0,538371653 |
| 431 | 2,12E+10 | 0,227929217 |
| 33 | 4,61E+09 | 0,297246757 |
| 49 | 1,45E+08 | 0,654124693 |
| 119 | 8,28E+08 | 0,401977071 |
| 245 | 1,50E+09 | 0,701037222 |
| 246 | 4,19E+09 | 0,348110038 |
| 191 | 9,22E+08 | 0,119109647 |
| 12 | 1,82E+08 | 1,161735726 |
| 104 | 1,02E+08 | 1,251533586 |
| 68 | 2,83E+08 | 0,403345858 |
| 46 | 2,97E+08 | 1,140378081 |
| 12 | 3,27E+08 | 0,25768074 |
| 479 | 1,52E+10 | 0,466151525 |
| 23 | 1,87E+09 | 0,208257178 |
| 328 | 6,64E+08 | 1,262538056 |
| 818 | 9,13E+07 | 0,39650579 |
| 286 | 6,15E+09 | 0,479631242 |
| 53 | 5,66E+08 | 0,984500113 |
| 204 | 5,03E+09 | 0,772904181 |
| 118 | 6,39E+09 | 0,918677808 |
| 194 | 1,04E+09 | 0,185740096 |
| 48 | 4,76E+07 | 0,441190807 |
| 27 | 5,19E+08 | 1,148396115 |
| 235 | 1,27E+08 | 0,782200592 |
| 37 | 1,64E+08 | 0,479262995 |
| 443 | 1,07E+09 | 0,20105333 |
| 35 | 2,75E+08 | 0,570875637 |
| 179 | 6,98E+08 | 0,286016407 |
| 97 | 1,64E+08 | 0,076814285 |
| 92 | 2,69E+08 | 0,895669813 |
| 47 | 3,45E+09 | 0,916352193 |
| 79 | 5,51E+07 | 0,069596907 |
| 57 | 2,71E+08 | 0,85284753 |
| 127 | 4,67E+08 | 1,133379848 |
| 46 | 2,01E+08 | 0,739469611 |
| 59 | 1,54E+08 | 0,4418734 |
| 305 | 2,23E+09 | 0,175314227 |
| 67 | 7,72E+07 | 0,440282873 |
| 93 | 1,77E+08 | 0,219460875 |
| 593 | 1,09E+11 | 0,709767625 |
| 159 | 4,56E+08 | 0,007598174 |
| 54 | 7,60E+07 | 0,264271114 |
| 47 | 7,57E+09 | 1,013902874 |

| | | |
|---|---|---|
| 252 | 1,53E+09 | 0,075583697 |
| 159 | 1,72E+09 | 1,166256713 |
| 30 | 6,46E+08 | 1,092733092 |
| 292 | 1,32E+09 | 1,054604143 |
| 78 | 3,64E+08 | 0,199810497 |
| 487 | 1,80E+10 | 1,171528274 |
| 72 | 9,99E+08 | 0,242375822 |
| 23 | 1,90E+08 | 0,128297571 |
| 23 | 1,09E+08 | 0,41466965 |
| 26 | 7,39E+08 | 1,079127113 |
| 104 | 3,40E+08 | 0,608672667 |
| 127 | 1,07E+09 | 0,668191073 |
| 225 | 4,01E+08 | 0,386423901 |
| 15 | 1,33E+09 | 1,063099736 |
| 218 | 5,78E+08 | 0,388214101 |
| 23 | 5,98E+07 | 0,49898377 |
| 71 | 1,48E+08 | 0,927314504 |
| 288 | 1,17E+09 | 0,016110608 |
| 271 | 8,40E+08 | 0,733972193 |
| 473 | 5,44E+09 | 0,428006385 |
| 472 | 7,70E+07 | 0,080857228 |
| 314 | 6,93E+08 | 0,102414182 |
| 59 | 1,20E+10 | 0,278397182 |
| 34 | 1,16E+08 | 0,446045552 |
| 71 | 1,36E+09 | 0,194788251 |
| 103 | 3,55E+08 | 1,215401321 |
| 374 | 1,61E+09 | 0,107595328 |
| 174 | 1,45E+09 | 0,13421437 |
| 72 | 4,01E+07 | 0,055413835 |
| 25 | 5,75E+08 | 0,55169034 |
| 24 | 1,01E+09 | 0,12160395 |
| 48 | 2,35E+08 | 0,382832743 |
| 253 | 2,23E+09 | 0,542117112 |
| 197 | 5,79E+08 | 0,231117511 |
| 77 | 4,71E+08 | 0,041894623 |
| 226 | 1,22E+09 | 0,39241087 |
| 242 | 3,63E+09 | 0,74726977 |
| 339 | 4,22E+08 | 0,779399873 |
| 192 | 3,60E+09 | 1,188936328 |
| 295 | 1,12E+09 | 0,751778882 |
| 209 | 1,65E+09 | 0,687449152 |
| 27 | 2,33E+08 | 0,760120645 |
| 77 | 1,18E+09 | 1,22704568 |
| 88 | 6,38E+08 | 0,301247147 |
| 149 | 2,60E+09 | 0,646364777 |
| 92 | 2,14E+08 | 0,332783712 |
| 8 | 9,63E+08 | 0,38884399 |
| 129 | 2,09E+08 | 0,014983737 |
| 127 | 8,53E+08 | 0,739145536 |
| 138 | 4,57E+08 | 0,151999348 |

| | | |
|---|---|---|
| 132 | 7,15E+08 | 0,190806661 |
| 20 | 1,46E+09 | 0,07647085 |
| 24 | 3,20E+08 | 1,116443171 |
| 157 | 2,27E+09 | 0,453772545 |
| 78 | 1,99E+10 | 0,250468418 |
| 298 | 1,66E+09 | 0,744360679 |
| 10 | 3,70E+08 | 0,20692986 |
| 149 | 4,74E+08 | 0,682437172 |
| 159 | 3,15E+08 | 0,519114568 |
| 61 | 1,50E+08 | 0,74097069 |
| 65 | 6,61E+08 | 0,656469121 |
| 128 | 2,58E+08 | 0,225989966 |
| 34 | 1,47E+08 | 0,254558726 |
| 129 | 4,25E+08 | 0,696351088 |
| 33 | 1,19E+08 | 0,248947094 |
| 428 | 1,48E+10 | 0,050183564 |
| 191 | 3,77E+09 | 1,126509232 |
| 166 | 1,47E+08 | 1,01180499 |
| 141 | 7,33E+08 | 0,907652058 |
| 86 | 1,43E+08 | 0,127566088 |
| 161 | 1,03E+09 | 0,150416132 |
| 149 | 1,69E+08 | 0,160372303 |
| 211 | 7,49E+08 | 0,109571524 |
| 73 | 4,39E+08 | 1,108307153 |
| 111 | 4,13E+08 | 0,144522532 |
| 89 | 3,05E+08 | 0,294038847 |
| 246 | 3,42E+08 | 0,609883279 |
| 73 | 4,07E+08 | 1,160793171 |
| 191 | 3,44E+09 | 1,134016616 |
| 169 | 8,55E+07 | 0,152318165 |
| 36 | 9,77E+07 | 0,28518013 |
| 49 | 5,27E+07 | 0,951211914 |
| 132 | 3,84E+08 | 1,123309588 |
| 211 | 1,17E+09 | 0,047046317 |
| 375 | 7,28E+09 | 0,165296424 |
| 299 | 1,94E+09 | 0,153487576 |
| 87 | 5,02E+08 | 1,196119998 |
| 16 | 3,93E+07 | 0,166943029 |
| 55 | 2,37E+08 | 0,058259077 |
| 261 | 1,90E+09 | 0,238650836 |
| 71 | 2,58E+08 | 0,697182914 |
| 111 | 1,20E+08 | 0,459045114 |
| 85 | 5,10E+08 | 0,256144386 |
| 238 | 5,13E+09 | 0,515865037 |
| 135 | 1,30E+09 | 0,157420233 |
| 383 | 3,38E+09 | 0,035762933 |
| 205 | 2,79E+09 | 1,286143483 |
| 128 | 1,74E+09 | 0,732524234 |
| 254 | 2,51E+09 | 0,237341386 |
| 101 | 1,45E+09 | 0,214470581 |

| | | |
|---|---|---|
| 27 | 4,17E+07 | 0,388322746 |
| 84 | 8,99E+07 | 0,417298695 |
| 304 | 1,25E+09 | 0,200604865 |
| 83 | 2,37E+08 | 0,008687259 |
| 112 | 9,92E+07 | 0,702556887 |
| 256 | 1,25E+09 | 0,344628611 |
| 333 | 1,84E+09 | 0,700021289 |
| 227 | 3,61E+08 | 0,793239658 |
| 289 | 1,12E+09 | 0,264414501 |
| 404 | 2,78E+09 | 0,25412113 |
| 77 | 1,25E+08 | 0,884949002 |
| 75 | 4,89E+08 | 0,069302394 |
| 65 | 1,23E+09 | 0,111390005 |
| 93 | 3,27E+08 | 0,532418082 |
| 88 | 2,35E+08 | 1,243944867 |
| 246 | 8,86E+08 | 0,782906571 |
| 56 | 8,45E+07 | 0,072607966 |
| 323 | 8,42E+08 | 0,102884908 |
| 78 | 1,22E+08 | 0,051336298 |
| 104 | 5,62E+08 | 0,110969055 |
| 22 | 1,38E+08 | 0,389700959 |
| 157 | 3,26E+09 | 0,749479108 |
| 355 | 5,00E+09 | 0,060270559 |
| 82 | 2,35E+08 | 0,064238609 |
| 283 | 1,05E+09 | 0,74521114 |
| 181 | 1,81E+09 | 0,062392274 |
| 177 | 9,88E+08 | 0,40919231 |
| 63 | 2,57E+08 | 0,59876076 |
| 18 | 1,03E+09 | 0,612985004 |
| 99 | 4,49E+08 | 0,417452211 |
| 72 | 8,83E+08 | 0,295657108 |
| 321 | 1,06E+09 | 0,607480065 |
| 38 | 4,07E+08 | 0,070978946 |
| 2 | 4,62E+08 | 0,705490543 |
| 136 | 2,02E+09 | 0,689435608 |
| 115 | 1,62E+09 | 0,903828281 |
| 233 | 6,23E+08 | 0,775708708 |
| 302 | 1,47E+10 | 0,229407919 |
| 157 | 2,24E+09 | 0,684602064 |
| 10 | 4,94E+08 | 0,222241958 |
| 65 | 2,58E+08 | 1,005387513 |
| 107 | 2,68E+08 | 0,029991159 |
| 427 | 7,38E+08 | 0,806237374 |
| 55 | 1,59E+08 | 0,239759048 |
| 243 | 2,97E+09 | 0,705469761 |
| 63 | 1,13E+09 | 0,045118998 |
| 146 | 5,80E+09 | 1,200788301 |
| 31 | 1,80E+09 | 0,407864132 |
| 135 | 2,39E+09 | 0,816075645 |

| | | |
|---|---|---|
| 279 | 2,10E+09 | 0,131702708 |
| 206 | 2,66E+09 | 0,159592492 |
| 219 | 2,69E+08 | 0,024964747 |
| 213 | 1,54E+09 | 0,653115664 |
| 58 | 1,83E+08 | 0,37999398 |
| 358 | 3,70E+09 | 1,182254486 |
| 116 | 1,91E+08 | 0,534698266 |
| 128 | 8,10E+08 | 0,659862249 |
| 139 | 2,61E+09 | 0,598590373 |
| 29 | 1,37E+07 | 0,244287754 |
| 52 | 1,58E+08 | 0,246145129 |
| 79 | 1,10E+09 | 1,219976364 |
| 329 | 7,49E+09 | 0,341995972 |
| 55 | 6,70E+08 | 0,438492479 |
| 165 | 2,20E+08 | 0,622782623 |
| 67 | 1,70E+08 | 0,591741705 |
| 179 | 1,16E+09 | 0,502959011 |
| 213 | 3,31E+09 | 0,168466783 |
| 479 | 1,40E+09 | 1,200064005 |
| 77 | 2,49E+08 | 0,374386863 |
| 295 | 7,50E+09 | 0,812219847 |
| 205 | 4,16E+09 | 0,767233622 |
| 335 | 2,69E+09 | 0,163148956 |
| 17 | 6,40E+08 | 0,623240129 |
| 66 | 1,48E+08 | 0,210363488 |
| 53 | 1,39E+09 | 0,929324699 |
| 46 | 3,10E+08 | 0,219550226 |
| 46 | 3,94E+08 | 0,001082896 |
| 23 | 2,23E+09 | 1,282807561 |
| 313 | 9,20E+09 | 1,193885952 |
| 223 | 2,95E+09 | 0,605410275 |
| 104 | 1,06E+09 | 0,42784211 |
| 235 | 6,07E+09 | 0,668965244 |
| 413 | 1,68E+11 | 1,093164486 |
| 406 | 6,72E+09 | 0,014297105 |
| 262 | 8,52E+08 | 0,820756682 |
| 181 | 7,75E+08 | 0,454979502 |
| 448 | 1,51E+09 | 0,478995288 |
| 79 | 3,35E+08 | 0,431172295 |
| 19 | 2,52E+08 | 0,103152454 |
| 149 | 3,72E+08 | 0,435414632 |
| 52 | 2,02E+08 | 0,750898463 |
| 257 | 6,98E+08 | 0,987371493 |
| 219 | 3,65E+09 | 0,053328138 |
| 8 | 3,33E+08 | 0,082151243 |
| 147 | 2,93E+08 | 0,236004862 |
| 373 | 2,04E+09 | 0,461989088 |
| 58 | 1,53E+08 | 0,815148484 |
| 15 | 1,25E+09 | 0,881766842 |

| | | |
|---|---|---|
| 191 | 1,65E+09 | 0,341036334 |
| 691 | 9,54E+07 | 0,235811607 |
| 4 | 8,55E+07 | 0,033676081 |

| est ($^{bio}$*ari* /$^{bio}$*Ub*) | | | |
|---|---|---|---|
| **Fold Change ($log_2$ scale)** | **Protein IDs** | **Majority protein IDs** | **Gene names** |
| 1,269837697 | M9PH10;P46461; | M9PH10;P46461 | comt |
| 1,37420845 | Q9V9T5;Q8I0S9 | Q9V9T5;Q8I0S9 | Mccc1 |
| 4,705107371 | A4V4Q7;Q94981; | A4V4Q7;Q94981 | ari-1 |
| 1,036558787 | V9GZU6;M9PD1 | V9GZU6;M9PD18;P48602 | Vha68-1 |
| 1,333567301 | Q9VPW3;Q8MS0 | Q9VPW3;Q8MS09 | CG4629 |
| 2,481012344 | Q94885 | Q94885 | mdg3/ORF |
| 1,182474772 | Q9V5C6 | Q9V5C6 | Prosalpha7 |
| 6,668132146 | Q7K2W6;Q27598 | Q7K2W6;Q27598 | PPO1 |
| 1,306407928 | B5RJL1;P08255;( | B5RJL1;P08255;Q9TX52 | Rh4 |
| 2,121911367 | Q9U4H1;Q0E938 | Q9U4H1;Q0E938;T2GGE1 | CG6424 |
| 1,528125763 | Q9VZV5 | Q9VZV5 | CG32486 |
| 1,191648483 | Q9VEA1;Q9NJB | Q9VEA1;Q9NJB4 | eIF1A |
| 2,89976565 | B7YZI0;Q9V7N5 | B7YZI0;Q9V7N5-2;Q9V7N | Vha44 |
| 1,871335347 | M9NCR7;M9ND3 | M9NCR7;M9ND31;M9NE | Oscillin |
| 1,179653804 | Q9W2U2;F6J9D1 | Q9W2U2;F6J9D1;F6J9C7 | CG32687 |
| 2,509445826 | A0A0B4LFU6;A0 | A0A0B4LFU6;A0A0B4LG | CG15117 |
| 1,49417305 | X2J8F5;P11996 | X2J8F5;P11996 | Lsp1beta |
| 2,423790614 | Q967T6 | Q967T6 | Gag |
| 1,856972377 | O01668 | O01668 | Rh6 |
| 1,385567347 | Q9VSL2 | Q9VSL2 | GstO3 |
| 3,158693949 | Q9GV29;E2RWQ | Q9GV29 | MoxGM95 |
| 3,033847809 | A0A0B4LHD7 | A0A0B4LHD7 | Calx |

| est ($^{bio}$*ari* /$^{bio}$*Ub*) | | | |
|---|---|---|---|
| **Fold Change ($log_2$ scale)** | **Protein IDs** | **Majority protein IDs** | **Gene names** |
| -1,443918864 | P35992-2;P35992- | P35992-2;P35992-1;P35992 | Ptp10D |
| -1,519742966 | P12646;P12646-2; | P12646;P12646-2;B7FNK0 | Zw |
| -1,197481155 | P04146-2;P04146; | P04146-2;P04146 | copia\GIP |
| -1,020881653 | Q8IRS0;E1JJE4;X | Q8IRS0;E1JJE4;X2JCK2;Q | Ptp4E |
| -1,285050074 | A0A0B4JD13;A0. | A0A0B4JD13;A0A0B4KH | snu |
| -1,019625982 | B5X0J4;P18459-2 | B5X0J4;P18459-2;P18459 | ple |
| -1,184392293 | Q9W0Y6;A0A0B | Q9W0Y6;A0A0B4KEZ8;A | pain |
| -1,136137644 | Q24336;Q7KVF1; | Q24336;Q7KVF1;Q9W0X( | emp |
| -1,089377721 | Q9VPJ9;Q8SZF5; | Q9VPJ9;Q8SZF5;Q86P66;( | CG3164 |
| -2,207279205 | Q86P18;Q9VBF7; | Q86P18;Q9VBF7 | CG33970 |
| -1,003317515 | C0PTX2;Q9U4G1 | C0PTX2;Q9U4G1;Q95U86 | bdl |
| -1,095693588 | Q8T0J5;Q9VE80; | Q8T0J5;Q9VE80;D0IQ98 | CG7675 |
| -1,295824051 | A0A0B4KFP1;Q0 | A0A0B4KFP1;Q0KI92;Q9 | CG12814 |
| -1,285161336 | Q9W3K9;Q8SWZ | Q9W3K9;Q8SWZ9 | Ldsdh1 |
| -1,315522512 | Q9VIT5;M9PG89 | Q9VIT5;M9PG89 | CG13077 |
| -1,157152176 | Q7JYX5 | Q7JYX5 | Vamp7 |
| -2,024297714 | Q9VY42 | Q9VY42 | CG1461 |
| -1,186169942 | P23625-2 | P23625-2 | Galphaq |
| -1,130781174 | X2J8E9;Q24292;( | X2J8E9;Q24292;Q29QI1 | ds |
| -1,432588577 | M9MS14;Q9W53 | M9MS14 | moody |
| -1,047320048 | Q9VYT4;Q6IDF6 | Q9VYT4;Q6IDF6 | ATP7 |
| -1,088359197 | Q7K2V9 | Q7K2V9 | CG33672 |

| | | | |
|---|---|---|---|
| -1,050004323 | Q9Y163;Q8I930;( | Q9Y163;Q8I930;Q9VR47;( | hoe1 |
| -1,625497818 | Q9W0M4 | Q9W0M4 | CG13887 |
| -1,720391591 | Q9XZ01;A0A0B4 | Q9XZ01;A0A0B4K692;D3 | Nep2 |
| -1,042865117 | Q86B77;A0A0B4 | Q86B77;A0A0B4KGZ9;Q7 | LpR2 |
| -1,169418335 | O97102 | O97102 | smt3 |
| -1,023333232 | Q7K2N0 | Q7K2N0 | Lnpk |
| -1,706658681 | Q7KN81;Q9XZ1 | Q7KN81;Q9XZ15 | Osi6 |
| -2,826539358 | Q9VCS2 | Q9VCS2 | CG13833 |
| -3,05454127 | Q8MZ32;A1Z6N | Q8MZ32;A1Z6N2 | Tdc1 |
| -2,764109294 | Q9VQF1;Q8T998 | Q9VQF1;Q8T998;M9PBX | CG31689 |
| -2,023932139 | Q9VI53 | Q9VI53 | CG1105 |
| -2,271789551 | Q9VJC0 | Q9VJC0 | CG6870 |
| -1,605457306 | Q9VKG4 | Q9VKG4 | CG31705 |
| -1,137947718 | Q9VKM8;Q9VK | Q9VKM8;Q9VKM7 | CG33129 |
| -1,400824865 | Q9VWT3 | Q9VWT3 | Ggt-1 |
| -1,989123027 | Q9VHQ8;A0A0B | Q9VHQ8;A0A0B4K642;A( | CG45263 |
| -2,060609182 | Q9VHI9;A0A0B4 | Q9VHI9;A0A0B4KGW3 | CG31100 |
| -1,273523966 | E9P245;A8VEL0; | E9P245;A8VEL0;Q9V4N3 | Cyt-b5 |
| -2,145721436 | D0Z752;M9PCF1 | D0Z752;M9PCF1;Q9VQH | Duox |
| -1,086554209 | R4NR58;K4M4D | R4NR58;K4M4D5;Q9GQQ | spin |
| -1,978580475 | M9PG22;P61209; | M9PG22;P61209;L0MNA8 | Arf79F |
| -1,155976613 | X2J9Z1;P42787-5 | X2J9Z1;P42787-5;P42787; | svr |
| -1,287623088 | Q8SY17;Q7KJ73 | Q8SY17;Q7KJ73 | Tsp42Ee |
| -2,283281326 | Q960W6;A0A0B | Q960W6;A0A0B4LFD0;A1 | CG8306 |
| -2,703658422 | Q961R9 | Q961R9 | CG6126 |
| -2,72403717 | Q9V558;Q9V559 | Q9V558;Q9V559 | Cyp4p1 |
| -3,201266607 | Q9VI56 | Q9VI56 | CG1943 |
| -1,098136266 | X2JDD7;Q9VMD | X2JDD7;Q9VMD9 | Tig |
| -2,840459824 | P42787-4;P42787 | P42787-4;P42787-3 | svr |
| -3,997159958 | Q8IQR5;B7Z067; | Q8IQR5;B7Z067;B7Z068;N | Nedd4 |
| -2,661157608 | Q94880 | Q94880 | pyd |

**est ($^{bio}$*ari* /$^{bio}$*Ub* )**

| Fold Change ($log_2$ scale) | Protein IDs | Majority protein IDs | Gene names |
|---|---|---|---|
| 0,375330607 | A1Z784;Q7JV23; | A1Z784;Q7JV23;A8DY67 | ACC |
| 0,204429626 | P02701 | P02701 | AVD |
| 0,604033152 | Q7KN97;Q6NKL | Q7KN97 | PCB |
| 0,370307922 | Q8IGR0;P15357 | Q8IGR0;P15357 | RpS27A |
| 1,644987742 | Q0E9E2;Q86NV1 | Q0E9E2;Q86NV1 | PCB |
| -0,452999115 | P00002 | P00002 | bioUb |
| -0,58232371 | Q9GQV2;A0A0B | Q9GQV2;A0A0B4JCY6;Q | cher |
| -0,38025411 | A0A0B4LGR2;A( | A0A0B4LGR2;A0A0B4JD | ctrip |
| -0,627981186 | B7Z0T2;A0A0B4 | B7Z0T2;A0A0B4K6R6;B7 | smash |
| 0,930639903 | A0A0B4K6A6;A( | A0A0B4K6A6;A0A0B4K6 | Calx |
| 0,387308757 | A0A0B4KGY9;D | A0A0B4KGY9;D0Z767;A( | msi |
| 0,347897847 | A0A0B4K6W2;A | A0A0B4K6W2;A0A0B4K7 | faf |
| -0,79089419 | A0A0B4K713;Q9 | A0A0B4K713;Q9Y091-2;Q | fl(2)d |
| 0,869082133 | Q9W1Y2;Q8T0K | Q9W1Y2;Q8T0K5;Q9W1Y | PIP5K59B |
| -0,901065191 | A0A0B4KEH8;A( | A0A0B4KEH8;A0A0B4KE | CG17883 |
| -0,683474223 | E3CTS3;Q8IGH3; | E3CTS3;Q8IGH3;A1Z6Z3; | Aldh-III |

| | | | |
|---|---|---|---|
| -0,428490957 | Q9VHX9;Q8SYR | Q9VHX9;Q8SYR7;A0A0B | CD98hc |
| -0,620267868 | Q86NS4;A1Z7H3 | Q86NS4;A1Z7H3;Q8T3L1; | Acsl |
| -0,302412669 | A0A0B4KFE9;A1 | A0A0B4KFE9;A1Z8P9 | Mtor |
| -0,640182495 | Q24468;Q24229; | Q24468;Q24229;A0A0B4K | put |
| -0,547822316 | A0A0B4KGG6;Q | A0A0B4KGG6;Q9VN12;Q | CG1090 |
| 0,248516719 | Q9VDI1;A0A0B4 | Q9VDI1;A0A0B4KH28 | Synd |
| 0,39590772 | A0A0B4KHE7;Q | A0A0B4KHE7;Q9VAA2 | AdoR |
| 0,319738388 | F0JAL7;A0A0B4 | F0JAL7;A0A0B4KHG5;A0 | chp |
| -0,467562358 | A0A0B4KHQ8;Q | A0A0B4KHQ8;Q9VB13;A | tau |
| -0,483411789 | A0A0B4KHT5;Q | A0A0B4KHT5;Q95TW4;A | Syp |
| 0,375541687 | A0A0B4KHY8;Q | A0A0B4KHY8;Q9XZ32 | CG10254 |
| 0,627560298 | A0A0B4LF49;Q8 | A0A0B4LF49;Q8T5S9;B7 | metro |
| 0,397527059 | A0A0B4LGC0;A | A0A0B4LGC0;A0A0B4LF | TppII |
| -0,750387828 | A8DYJ3;A0A0B4 | A8DYJ3;A0A0B4LG26;Q1 | CalpA |
| 0,625303268 | A0A0B4LGC6;Q | A0A0B4LGC6;Q8MLS7;A | l(2)k09913 |
| -0,455568314 | E4NKH1;A0A0B | E4NKH1;A0A0B4LGB0;Q | Gp150 |
| -0,600602468 | A0A0B4LGI1;P2 | A0A0B4LGI1;P20354-2;P2 | Galphas |
| -0,53911972 | A0A0B4LGL5;Q | A0A0B4LGL5;Q8MSU3 | CG8399 |
| 0,941026688 | A0A0B4LGM0;B | A0A0B4LGM0;B7YZQ3;A | prom |
| -0,512256622 | N0D584;A0A0H4 | N0D584;A0A0H4Y1G5;A0 | hyd |
| 0,85798645 | A0A0B4LH53;P5 | A0A0B4LH53;P54351 | Nsf2 |
| -0,528455734 | A0A0B4LHB3;P1 | A0A0B4LHB3;P12428;P12 | bw |
| -0,713542302 | A0A0B4LJ12;A0 | A0A0B4LJ12;A0A0B4LH | awd |
| -0,744475047 | A0A0B7P9G0;Q6 | A0A0B7P9G0 | uex |
| -0,884375254 | A1Z746-2;A1Z74 | A1Z746-2;A1Z746 | cn |
| -0,7297376 | A1Z7S3;A1Z7S2; | A1Z7S3;A1Z7S2 | Rab32 |
| -0,692685445 | Q7K2P3;A1Z7Z4 | Q7K2P3;A1Z7Z4;Q8IGD2; | CG1648 |
| 0,384422302 | A1Z8B5;D3DMQ | A1Z8B5;D3DMQ0;Q8T97 | CG7220 |
| -0,240754445 | A1Z9J3 | A1Z9J3 | shot |
| -0,955977758 | A1Z9M5;Q95TJ2 | A1Z9M5;Q95TJ2 | CG30069 |
| -0,602920532 | E1JH83;A1ZAI5 | E1JH83;A1ZAI5 | CG5065 |
| -0,691467921 | E0R982;A1ZBL0; | E0R982;A1ZBL0;Q24560 | betaTub56D |
| -0,295642853 | Q9V948;E1JGN7; | Q9V948;E1JGN7;Q7KA80 | HnRNP-K |
| -0,649810791 | A2RVF0;Q9VQY | A2RVF0;Q9VQY4;Q9UAF | Atet |
| 0,287287394 | A4V1B2;Q9NB04 | A4V1B2;Q9NB04 | Patj |
| 0,86233902 | A4V2B8;P08181 | A4V2B8;P08181 | CkIIalpha |
| -0,937478383 | A4V2K7;P22058 | A4V2K7;P22058 | D1 |
| -0,779572805 | A4V441;Q8SY66 | A4V441;Q8SY66;Q24524 | sn |
| -0,308163325 | A4V448;Q9W3K | A4V448;Q9W3K5 | Gclc |
| -0,678427378 | A4V488;Q07152; | A4V488;Q07152;Q07152-2 | ras |
| -0,795420329 | Q9VPQ7;A7KX2 | Q9VPQ7;A7KX20;A7KX1 | Spp |
| -0,589102427 | D5SHT8;Q5U1C | D5SHT8;Q5U1C5;B7Z029; | Nhe3 |
| -0,82886823 | A8DZ14;X2J6N7; | A8DZ14;X2J6N7;A2RVH3 | Fas3 |
| 0,63376236 | A8E775;P19107; | A8E775;P19107;Q5BIJ0 | Arr2 |
| -0,424602509 | A8JTM7;Q95SN5 | A8JTM7 | mgl |
| -0,750235875 | A8JUY0;Q9W4D | A8JUY0;Q9W4D2 | Rnp4F |
| 0,433369954 | B3DN78;Q00174 | B3DN78;Q00174 | LanA |
| 0,570513407 | B5RIN0;M9PB50 | B5RIN0;M9PB50;Q9VMD | retm |
| -0,583531062 | B5X0J6;Q9V9J3; | B5X0J6;Q9V9J3;A1Z6I9 | Src42A |
| -0,871247609 | B7YZL6;Q7YTZ | B7YZL6;Q7YTZ4;B7YZL | CG10737 |

| | | | |
|---|---|---|---|
| -0,488550186 | M9NEQ9;D4G7H | M9NEQ9;D4G7H1;C0H6Z | RpS10b |
| 0,24306043 | Q8MSJ2;C4XVJ6 | Q8MSJ2;C4XVJ6;Q24265; | boss |
| -0,873401642 | Q0E8T6;H0RNG5 | Q0E8T6;H0RNG5;C6SV06 | CG34125 |
| -0,835114161 | Q8T9J9;C6TP50;( | Q8T9J9;C6TP50;Q9W401;( | kdn |
| -0,440219879 | D3DMY5;C6TP7( | D3DMY5;C6TP70;P92177- | 14-3-3epsilon |
| -0,510741552 | C6TP87;P08736;( | C6TP87;P08736;Q8T3U3 | eEF1alpha1 |
| 0,856564204 | Q8IQX3;C7LA72 | Q8IQX3;C7LA72;Q4V5C5 | CG32544 |
| -0,89217186 | E1JJ68;C7LAE4;I | E1JJ68;C7LAE4;P19109-3; | Rm62 |
| -0,455792745 | M9NG50;M9PHG | M9NG50;M9PHG2;C7LAH | Moe |
| 0,440887451 | Q9W0E4;C8VUZ | Q9W0E4;C8VUZ1;Q8IRH( | Psa |
| -0,510222117 | C8VV33;D3DMH | C8VV33;D3DMH8;C9QP4 | eIF4A |
| -0,386089325 | Q7K4I5;C8VV52 | Q7K4I5;C8VV52 | CG8892 |
| -0,783068975 | X2JJG8;Q494K4; | X2JJG8;Q494K4;X2JDA5;( | Gs2 |
| 0,707992554 | M9MRI9;C9QPB9 | M9MRI9;C9QPB9;Q59DZ9 | qtc |
| 0,471200307 | Q86BQ0;D0IQJ1; | Q86BQ0;D0IQJ1;Q9W0C2 | nSyb |
| 0,346959432 | Q76NQ9;Q961Q7 | Q76NQ9;Q961Q7;Q9VY76 | AMPdeam |
| 0,82001241 | D2NUF0;Q24273; | D2NUF0;Q24273;Q0E8B8; | nrm |
| 0,3677845 | D3DMF9;Q24008 | D3DMF9;Q24008;D3DMI7 | inaD |
| 0,168793996 | Q9V3N7;Q2XST3 | Q9V3N7;Q2XST3;D3YE81 | CRMP |
| -0,75619634 | D4G7C9;M9NFB6 | D4G7C9;M9NFB6;E1JHM | spir |
| -0,599796931 | M9PFU6;D6W4W | M9PFU6;D6W4W6;M9PC2 | CG18507 |
| 0,564021428 | E1JGZ9;Q9XZJ4 | E1JGZ9;Q9XZJ4 | Prosalpha1 |
| -0,608789444 | Q8T4M0;E1JHB8 | Q8T4M0;E1JHB8;M9MRE | Pvr |
| -0,784776688 | Q9VKU1;E1JHE4 | Q9VKU1;E1JHE4 | Fatp1 |
| 0,924114863 | Q8T0S9;Q9U9K1 | Q8T0S9;Q9U9K1;Q9VPS6 | Eaat2 |
| 0,597025553 | E1JHR5;P15007-2 | E1JHR5;P15007-2;P15007 | Eno |
| -0,541624705 | Q9VMV9;E1JHT( | Q9VMV9;E1JHT6;Q6NP2( | Rtnl1 |
| -0,813851674 | E1JJF3;Q9V427 | E1JJF3;Q9V427 | Inx2 |
| 0,757907232 | E1JJK8;E1JJK9;E | E1JJK8;E1JJK9;E1JJK7;E1 | rdgB |
| 0,494493484 | Q9V405;E1UIA5; | Q9V405;E1UIA5 | Rpt3 |
| -0,993372599 | Q5BHW4;E2QD6 | Q5BHW4;E2QD66;Q0KH( | CG34120 |
| 0,64207077 | Q8IQA9;E6PBV8 | Q8IQA9;E6PBV8;Q8IQA8; | Culd |
| 0,461761475 | Q95R82;Q7KGU6 | Q95R82;Q7KGU6;F2FB78 | CG14767 |
| -0,47628657 | F6J6B9;X2JF59;( | F6J6B9;X2JF59;Q9V3P0 | Jafrac1 |
| -0,524266561 | H1UUB1;P18489- | H1UUB1;P18489-4;P18489 | Syb |
| 0,214670817 | H9ZJM5;P13607- | H9ZJM5;P13607-4;P13607 | Atpalpha |
| 0,495672862 | L0MPS3;Q9V496 | L0MPS3;Q9V496 | apolpp |
| -0,682102203 | M9MQH9;P36179 | M9MQH9;P36179;M9NDI( | Pp2A-29B |
| -0,114650726 | M9MRD1;M9PC8 | M9MRD1;M9PC84;M9PCI | Msp300 |
| -0,156387329 | M9MS15;P23654; | M9MS15;P23654 | Nrt |
| -0,441336314 | M9MSM5;Q7YZ/ | M9MSM5;Q7YZA4;Q8IRY | DAAM |
| -0,6750501 | M9NE89;P08928 | M9NE89;P08928 | Lam |
| 0,214694977 | M9NEP1;M9ND9 | M9NEP1;M9ND95;E1JHJ4 | Mhc |
| 0,357332865 | M9NES0;P15372; | M9NES0;P15372;X2J8V5 | Arr1 |
| -0,406628291 | M9NEW1;P54357 | M9NEW1;P54357 | Mlc-c |
| -0,350038528 | M9NFA7;Q9VX9 | M9NFA7;Q9VX91 | Ubr1 |
| 0,869586309 | M9PB59;Q8IPI3;I | M9PB59;Q8IPI3;M9PC92;( | Ndae1 |
| 0,932332993 | Q8MQR4;Q9VIV( | Q8MQR4;Q9VIV0;M9PBE | CG10188 |
| -0,640023549 | O76268;Q9VQI8; | O76268;Q9VQI8;M9PBV2 | NTPase |
| -0,983971278 | M9PCA7;M9PI37 | M9PCA7;M9PI37;Q9VTW | Ncc69 |

| | | | |
|---|---|---|---|
| -0,238021215 | M9PCE0;Q01604 | M9PCE0;Q01604 | Pgk |
| -0,554360072 | M9PCF8;Q9VUC | M9PCF8;Q9VUC6-2;Q9VU | Frl |
| -0,439188004 | M9PCI6;Q9VQW | M9PCI6;Q9VQW7;Q9BN1 | ed |
| 0,738259633 | M9PCU0;P10676- | M9PCU0;P10676-2 | ninaC |
| 0,805667241 | M9PD75;Q94920 | M9PD75;Q94920 | porin |
| -0,99995931 | Q7K3K5;M9PEG | Q7K3K5;M9PEG6 | ZnT63C |
| 0,872236252 | M9PEL3;Q9VS54 | M9PEL3;Q9VS54;O61539; | qm |
| 0,910884857 | M9PF40;Q9VTE5 | M9PF40;Q9VTE5 | Pfdn2 |
| 0,2469546 | M9PIG8;P55035 | M9PIG8;P55035 | Rpn10 |
| -0,478009542 | M9PJN8;P07487; | M9PJN8;P07487 | Gapdh2 |
| -0,467407227 | O02373;Q95SJ7 | O02373 | sgl |
| -0,710442861 | Q8T410;Q0KHZ9 | Q8T410;Q0KHZ9;Q9VAL7 | Cnx99A |
| -0,685401917 | O15971 | O15971 | Rab10 |
| -0,904417674 | O62619-2;O6261 | O62619-2;O62619 | PyK |
| 0,57698377 | O76863 | O76863 | eIF2Bbeta |
| -0,641915639 | O97125;Q95NM1 | O97125 | Hsp68 |
| -0,908052444 | O97428;D5AEL7; | O97428;D5AEL7;Q8IRS7 | cib |
| 0,611006419 | P06002;S5M1F0; | P06002 | ninaE |
| -0,701702754 | P06605;P06603;K | P06605;P06603;K7X561;K | alphaTub84D |
| -0,540903091 | P06754-2;P06754- | P06754-2;P06754-5 | Tm1 |
| 0,913740158 | P08510-8;P08510- | P08510-8;P08510-7;P0851 | Sh |
| 0,497844696 | P10676 | P10676 | ninaC |
| 0,375759761 | P11046 | P11046 | LanB1 |
| -0,51922671 | X2JE30;X2JAU0; | X2JE30;X2JAU0;P11584 | mys |
| -0,734357834 | P13060 | P13060 | eEF2 |
| 0,596307755 | P13607-5;E1JIR4; | P13607-5;E1JIR4;B5RIT8; | Atpalpha |
| 0,339251836 | X2JAW6;P15215 | X2JAW6;P15215 | LanB2 |
| 0,835968653 | P18053;Q9VA12 | P18053 | Prosalpha3 |
| 0,641012828 | U3PXB8;P19334; | U3PXB8;P19334;F0JAI0;Q | trp |
| -0,982139587 | P20432 | P20432 | GstD1 |
| 0,40847524 | X2J4C1;P21521-2 | X2J4C1;P21521-2;P21521; | Syt1 |
| 0,280286789 | P22700-2;A0A0B- | P22700-2;A0A0B4LGB7;E | SERCA |
| -0,179249446 | X2JC31;P29742 | X2JC31;P29742 | Chc |
| 0,826359431 | P32392 | P32392 | Arp3 |
| -0,762725194 | P48611 | P48611 | pr |
| 0,575235367 | P48994;B5T1W9; | P48994 | trpl |
| -0,309620539 | P55828 | P55828 | RpS20 |
| -0,482777913 | P91926;E8NH12; | P91926;E8NH12;P91926-2 | AP-2alpha |
| -0,435131073 | P91938-4;P91938- | P91938-4;P91938-2;P9193 | Trxr-1 |
| 0,861848195 | Q06003-4;Q4QPC | Q06003-4;Q4QPQ6;Q0600 | gol |
| 0,753797531 | Q09103-3;A0A02 | Q09103-3;A0A023GPM5;M | rdgA |
| 0,454465866 | Q0KI96;A0A023U | Q0KI96;A0A023UKB8;Q9 | ps |
| -0,669497808 | Q9U4Y0;Q9U4Y1 | Q9U4Y0;Q9U4Y1;Q0KI98 | TrpRS |
| -0,746087392 | Q59E30;Q24036; | Q59E30;Q24036;Q95RI5;M | fax |
| 0,342360814 | Q86DS7;Q53ZT0; | Q86DS7;Q53ZT0;Q24133 | DnaJ-1 |
| -0,618682226 | Q24201;Q9VIB5; | Q24201;Q9VIB5;Q8IH63 | alpha-Est7 |
| -0,643250783 | Q24298 | Q24298 | shg |
| 0,804782867 | Q24311;D3DMX2 | Q24311;D3DMX2 | Cul1 |
| -0,920689265 | Q24324;X2JAZ3 | Q24324;X2JAZ3 | Dsor1 |
| -0,823188782 | Q26459;Q9VRM5 | Q26459;Q9VRM5;Q8IGD4 | Msr-110 |

| | | | |
|---|---|---|---|
| 0,558543523 | Q9VG62;Q29R43 | Q9VG62;Q29R43 | trus |
| -0,665400823 | Q95T75;Q9TWZ1 | Q95T75;Q9TWZ1;Q3YMU | ERp60 |
| 0,388317744 | Q961T2;Q494G8 | Q961T2;Q494G8 | Fbl6 |
| 0,601664225 | Q7KTU2;Q58CL2 | Q7KTU2;Q58CL2;C1C5A( | SPoCk |
| -0,667819977 | Q5U133;Q9VAF5 | Q5U133;Q9VAF5;Q5D716 | Cad99C |
| -0,911539714 | Q5U130;P05812 | Q5U130;P05812 | Hsp67Ba |
| 0,820453644 | Q8MSY7;Q5U16 | Q8MSY7;Q5U168;Q8IM95 | CG11155 |
| -0,678721746 | Q7JQU9;Q5U191 | Q7JQU9;Q5U191 | puml |
| 0,841597875 | Q7JYY8 | Q7JYY8 | Tsp42En |
| -0,777085622 | Q7K0E6;Q9XYM | Q7K0E6;Q9XYM1 | AspRS |
| -0,963895162 | Q9U3V4;Q7K1I4 | Q9U3V4;Q7K1I4 | Tsp42Ea |
| -0,412054698 | Q8IGB6;Q7K4M9 | Q8IGB6;Q7K4M9;A1ZBH | CG15118 |
| -0,309672674 | Q960V6;Q7K9H6 | Q960V6;Q7K9H6 | Sara |
| -0,451351802 | Q7KN75;Q9U982 | Q7KN75;Q9U982;Q7KN84 | Dp1 |
| -0,935563405 | Q7KNR7;Q9VV4 | Q7KNR7;Q9VV42;B7Z061 | Pdh |
| -0,846667608 | Q7KS11;Q961C4 | Q7KS11;Q961C4 | Ude |
| 0,531020482 | Q7KUK9;Q0E8E( | Q7KUK9 | CG17839 |
| -0,140183767 | Q9W1W0;Q7KVJ | Q9W1W0;Q7KVJ6;Q8SXT | Fatp2 |
| -0,323315938 | Q8T0H3;Q9NHP1 | Q8T0H3;Q9NHP1;Q7PLI0; | p120ctn |
| -0,910081228 | Q9VZQ8;Q8IRD4 | Q9VZQ8;Q8IRD4;Q8IRD3 | PHGPx |
| -0,689341227 | Q9VG23;Q86PF4 | Q9VG23;Q86PF4 | CG10126 |
| -0,421040217 | Q9VGM2;Q8IGA | Q9VGM2;Q8IGA2;Q8INK | fabp |
| -0,257887522 | Q8IGY6;Q8SWR | Q8IGY6;Q8SWR2 | CG32137 |
| -0,947961807 | Q9I7M3;Q8INW5 | Q9I7M3;Q8INW5;Q9VIW | CG10237 |
| 0,537730535 | Q8IQE6;A8JNR2; | Q8IQE6;A8JNR2;A8JNR3 | IRSp53 |
| 0,554385503 | Q8MLQ5;A0A0B | Q8MLQ5;A0A0B4LHE0;Q | kcc |
| 0,433055878 | Q8MQQ8;X2JDJ3 | Q8MQQ8;X2JDJ3;Q9VLS | PAPLA1 |
| -0,840496699 | Q9VIB3;Q8MRB | Q9VIB3;Q8MRB9;Q24202 | alpha-Est8 |
| -0,506467819 | Q8MRM6;Q9W3I | Q8MRM6;Q9W3D6 | CG12065 |
| -0,285336812 | Q8MS59 | Q8MS59 | wat |
| 0,494164785 | Q8T0L3 | Q8T0L3 | Uba1 |
| -0,580928802 | Q8T0T9 | Q8T0T9 | CG13920 |
| -0,71052742 | Q95U72;Q9W266 | Q95U72;Q9W266;Q8IGN2 | wdp |
| 0,765235265 | Q961U9;Q9VIK2; | Q961U9;Q9VIK2;D6W4U | CarT |
| -0,625562032 | Q962I2 | Q962I2 | Rala |
| 0,638150533 | Q967S6;Q9BMP4 | Q967S6;Q9BMP4 | HMS-Beagle\pol |
| -0,386535009 | Q99323-2;Q9932 | Q99323-2;Q99323-4;Q9932 | zip |
| -0,70526886 | Q9GP66;Q9W1X | Q9GP66;Q9W1X5;A0A0B | nahoda |
| 0,778416951 | Q9NHE5-7;Q9NH | Q9NHE5-7;Q9NHE5-4;Q9I | Cadps |
| 0,544094721 | Q9U6P7;Q8SYD5 | Q9U6P7;Q8SYD5 | Syt4 |
| 0,305612564 | Q9V3V6;Q9XZC | Q9V3V6;Q9XZC3 | Rpt5 |
| -0,480754217 | Q9V3W1;Q8IP18 | Q9V3W1;Q8IP18;M9PDJ9 | Tpr2 |
| -0,552735011 | Q9V8R9;A0A0B4 | Q9V8R9;A0A0B4LFX4;A( | cora |
| -0,720342 | Q9V9R2 | Q9V9R2 | Cul2 |
| -0,42403094 | Q9VAA9 | Q9VAA9 | CG7946 |
| -0,588776271 | Q9VAE7;Q7K1D | Q9VAE7;Q7K1D9 | CG31038 |
| -0,858279546 | Q9VBW3 | Q9VBW3 | Cad96Ca |
| -0,734301249 | Q9VC18;Q95RT1 | Q9VC18 | CG11089 |
| -0,586362203 | Q9VCR9 | Q9VCR9 | CG17121 |
| -0,861410141 | Q9VCW2;Q6GK | Q9VCW2 | cd |

| | | | |
|---|---|---|---|
| 0,667823156 | Q9VDG5;O18367 | Q9VDG5;O18367 | Calx |
| -0,444265366 | Q9VDW6-1;Q9VI | Q9VDW6-1;Q9VDW6;Q9V | Dys |
| -0,990696589 | Q9VE49;E1NZD7 | Q9VE49;E1NZD7 | CG7702 |
| 0,804938634 | Q9VEJ4 | Q9VEJ4 | CG5823 |
| -0,326679866 | Q9VF03;A0A0B4 | Q9VF03;A0A0B4JDA0;Q7 | mor |
| -0,642653147 | S5PU30;Q9VGS2 | S5PU30;Q9VGS2 | Tctp |
| -0,763963699 | Q9VH76 | Q9VH76 | Snap24 |
| -0,362349828 | Q9VH90 | Q9VH90 | trbd |
| -0,962242126 | Q9VH93;Q8SX48 | Q9VH93;Q8SX48 | CG9444 |
| -0,302886327 | Q9VHR8-2;Q9VH | Q9VHR8-2;Q9VHR8;Q6B( | DppIII |
| -0,750958125 | Q9VIE8 | Q9VIE8 | mAcon1 |
| 0,98419253 | Q9VJ39;R4GRV7 | Q9VJ39;R4GRV7 | CG10602 |
| -0,823295593 | Q9VJA9;F0JAM7 | Q9VJA9;F0JAM7;X2J6K6 | rdo |
| 0,187039693 | Q9VK44;Q8T3I1 | Q9VK44;Q8T3I1 | CG9934 |
| -0,56824557 | Q9VKK1 | Q9VKK1 | Ge-1 |
| 0,653910319 | Q9VKW5;Q961N | Q9VKW5;Q961N7 | CG5355 |
| -0,842237473 | Q9VNA6;Q95S15 | Q9VNA6;Q95S15 | Cerk |
| -0,289223989 | Q9VND3;Q8STF7 | Q9VND3;Q8STF7;Q8I0B7; | Pi4KIIalpha |
| -0,659069061 | Q9VQ34 | Q9VQ34 | CG17660 |
| -0,421977997 | Q9VRD9 | Q9VRD9 | Cbs |
| -0,537834803 | Q9VSS2 | Q9VSS2 | Srp68 |
| -0,608914693 | Q9VU17;Q8IQH6 | Q9VU17;Q8IQH6 | CG10960 |
| 0,25380071 | Q9VUC1;Q9XZT | Q9VUC1;Q9XZT5;M9MSI | Hsc70Cb |
| 0,227624893 | Q9VUQ7;E5KZJ( | Q9VUQ7;E5KZJ0;E5KZJ7 | CG7739 |
| 0,274636587 | Q9VVI3-4;Q9VV | Q9VVI3-4;Q9VVI3-3 | Nedd4 |
| -0,525870641 | Q9VWE0 | Q9VWE0 | dome |
| 0,603656133 | Q9VWS1 | Q9VWS1 | CG6617 |
| 0,480548223 | X2JFX0;Q9VX25 | X2JFX0;Q9VX25 | CG8188 |
| -0,62648201 | X2JFG6;Q9VXC1 | X2JFG6;Q9VXC1 | CG34325 |
| -0,2743632 | Q9VXF9 | Q9VXF9 | rngo |
| -0,650004705 | Q9VXG4-3;Q9VX | Q9VXG4-3;Q9VXG4-2;Q9 | AnxB11 |
| -0,468446096 | Q9VXQ5 | Q9VXQ5 | CCT6 |
| -0,422460556 | Q9VZ71 | Q9VZ71 | CG15211 |
| -0,401477814 | Q9W0H9 | Q9W0H9 | Rabex-5 |
| -0,609052658 | Q9W404;B3DNM | Q9W404;B3DNM5 | CG3842 |
| 0,566348394 | Q9W461 | Q9W461 | CG16721 |
| -0,892999649 | X2JFT2;Q9XZ14 | X2JFT2;Q9XZ14 | goe |
| -0,812088013 | Q9Y165;Q86NX0 | Q9Y165;Q86NX0 | morgue |
| -0,917427063 | X2JAT7;P15278 | X2JAT7;P15278 | Fas3 |
| -1,802269618 | Q59E58;A0A0B4 | Q59E58;A0A0B4JD57;Q59 | zip |
| -1,307018916 | A0A0B4K807;D3 | A0A0B4K807;D3DN03;A0 | Mlf |
| -1,189736684 | A0A0B4K812;Q7 | A0A0B4K812;Q7KNS3;Q7 | Lis-1 |
| -1,092972438 | Q9U6X3;Q9VNF | Q9U6X3;Q9VNF7;B7Z0T5 | MTA1-like |
| -1,209909439 | A0A0B4KGQ0;Q | A0A0B4KGQ0;Q9VCI7 | RanBP3 |
| -1,133141836 | A0A0B4KI19;Q9 | A0A0B4KI19;Q9VBP3 | Tnks |
| -1,350357691 | Q7K3C2;A0A0B4 | Q7K3C2;A0A0B4LFF6 | CG8180 |
| -2,061658859 | A0A0B4LFH2;Q7 | A0A0B4LFH2;Q7JUP3;F0 | casp |
| -1,527371724 | F3YDG2;A1Z792 | F3YDG2;A1Z792;Q8SZH3 | CG12769 |
| -1,705768585 | A0A0B4LGH1;P0 | A0A0B4LGH1;P08841;Q7 | betaTub60D |
| -1,318346659 | Q95SA6;Q8SYQ2 | Q95SA6;Q8SYQ2;A0AQH | Prosbeta1 |

| | | | |
|---|---|---|---|
| -1,642735163 | Q5BI38;A1Z9K4; | Q5BI38;A1Z9K4;Q7KQC4 | RN-tre |
| -1,845076879 | A4UZI6;P48148 | A4UZI6;P48148 | Rho1 |
| -3,88361613 | B3DMP7;Q9W1G | B3DMP7;Q9W1G0 | Taldo |
| -2,031548818 | B5RIN5;Q7KML | B5RIN5;Q7KML2 | CG5009 |
| -1,367257436 | Q8IQX5;Q9VWQ | Q8IQX5;Q9VWQ7;Q8MQ | CG6891 |
| -1,846337001 | Q59DZ2;Q9VJR4 | Q59DZ2;Q9VJR4;B6CLF8 | CG15279 |
| -1,839913686 | X2JC80;C6SV50; | X2JC80;C6SV50;P38979-2 | sta |
| -1,317566554 | Q8IPJ7;D3DMD4 | Q8IPJ7;D3DMD4;M9PCQ | Cpr |
| -2,37453715 | M9PDV9;M9PDI | M9PDV9;M9PDI9;D3DML | sky |
| -1,171352386 | F3YDH5;Q8IH88 | F3YDH5;Q8IH88;P29843; | Hsc70-1 |
| -2,24591128 | Q7PLL1;Q8SYL3 | Q7PLL1;Q8SYL3;L7EEF1; | eIF4B |
| -1,26327006 | M9MRT6;P08645 | M9MRT6;P08645 | Rap1 |
| -4,687680562 | M9MRU0;Q9XZ | M9MRU0;Q9XZC9;Q8IP5 | wb |
| -2,423575719 | M9MS42;Q8IRM | M9MS42;Q8IRM7;Q9W32 | Ptpmeg2 |
| -1,479436239 | Q8IQ69;M9NE38 | Q8IQ69;M9NE38 | jv |
| -1,223794301 | M9NFR5;Q95RE | M9NFR5;Q95RE4;O46173 | yps |
| -3,366704941 | M9PBT2;Q960M | M9PBT2;Q960M7;Q9VS2 | corn |
| -1,811104457 | Q9VZI9;Q9NGX | Q9VZI9;Q9NGX8;Q95V83 | wit |
| -1,217960358 | M9PEB1;Q95R81 | M9PEB1;Q95R81;Q9VRV | fmt |
| -2,493350347 | Q6SAX0;Q6SAF | Q6SAX0;Q6SAF6;Q6SAK | Egfr |
| -1,286816279 | Q86BZ5;Q86BZ4 | Q86BZ5;Q86BZ4;Q86BZ3; | Hsf |
| -1,065912247 | Q03427;A0A0B4 | Q03427;A0A0B4KFY9 | LamC |
| -2,97708257 | Q0KHR5;Q9VX9 | Q0KHR5;Q9VX92;D0Z73 | RhoGAP15B |
| -2,124657313 | Q8T015;Q0KI06 | Q8T015;Q0KI06 | CG5938 |
| -1,086164474 | Q7K5M6 | Q7K5M6 | CG10939 |
| -2,964766184 | Q8WQ84;Q961Q | Q8WQ84;Q961Q5;Q9GTZ | kkv |
| -1,014348984 | Q9TW62;Q9VBI | Q9TW62;Q9VBI2;Q29QY | Dak1 |
| -1,777795792 | Q9VAY2;Q1RKX | Q9VAY2;Q1RKX0 | Gp93 |
| -1,131458282 | Q9VBU6 | Q9VBU6 | CG11857 |
| -1,466471354 | Q9VGG5 | Q9VGG5 | Cad87A |
| -1,118305206 | Q9VIF2 | Q9VIF2 | Pld3 |
| -1,698652903 | Q9VQ91 | Q9VQ91 | papi |
| -1,565923691 | Q9VR44 | Q9VR44 | CG3036 |
| -1,11555926 | Q9VWI2 | Q9VWI2 | Naa15-16 |
| -1,508078893 | Q9W330;Q9GNH | Q9W330;Q9GNH8;B4YW | Hex-A |
| -1,90720431 | Q9Y128;M9PEI8 | Q9Y128;M9PEI8 | cert |
| 1,471474965 | A0A0B4LEY5;A1 | A0A0B4LEY5;A1Z7T2;A0 | Pkn |
| 1,549180349 | Q9W2J2;A0A0B4 | Q9W2J2;A0A0B4KF17;Q8 | ASPP |
| 1,488913218 | A0A0B4KF46;O4 | A0A0B4KF46;O46111 | Uba1 |
| 1,780279795 | A0A0B4KG14;A | A0A0B4KG14;A0A0B4KF | aux |
| 1,331207275 | Q8SWW8;A0A0 | Q8SWW8;A0A0B4KFY0 | dnr1 |
| 2,05239296 | A0A0B4LH50;P1 | A0A0B4LH50;P10981;B6I | Act87E |
| 1,148681005 | A1ZBM2;Q7K1S | A1ZBM2;Q7K1S6 | TBCB |
| 1,285463969 | A8JQT5 | A8JQT5 | Cdep |
| 2,593026479 | B3DMK6;M9MS | B3DMK6;M9MSI1;Q9VK | w-cup |
| 1,011467616 | B5RJS0;X2JAX8; | B5RJS0;X2JAX8;Q7K115; | Nost |
| 1,00201416 | M9PFL9;M9PD8 | M9PFL9;M9PD87;M9PCX | cmet |
| 2,008688609 | O46094;Q95R90 | O46094 | Ocrl |
| 3,26473554 | Q9W1M7;P91635 | Q9W1M7;P91635 | Pi3K59F |
| 1,151755015 | Q24113;Q7JPC7; | Q24113;Q7JPC7 | nonA-l |

| | | | |
|---|---|---|---|
| 2,152582804 | Q6NP06;Q960R8; | Q6NP06;Q960R8;Q9VB20; | dsd |
| 1,554024378 | Q8IPZ5;Q9VQJ6 | Q8IPZ5;Q9VQJ6 | CG9641 |
| 1,206963857 | Q9VIE7;Q8WSN4 | Q9VIE7;Q8WSN4;C4IXY5 | bur |
| 1,032612483 | X2JE14;Q9V7D2; | X2JE14;Q9V7D2;Q9NEF6 | Vha36-3 |
| 1,057146072 | Q9VHK1;Q8MQ( | Q9VHK1;Q8MQQ3;Q8INC | pyd |
| 1,797264735 | Q9VJZ6 | Q9VJZ6 | CG6523 |
| 1,645618439 | Q9VMX3 | Q9VMX3 | CG3756 |
| 1,675722758 | Q9VPA9 | Q9VPA9 | CG3618 |
| 1,455001831 | Q9VRK9 | Q9VRK9 | Klp64D |
| 1,354380925 | Q9VTC1 | Q9VTC1 | CG6418 |
| 2,293118159 | Q9VVJ5 | Q9VVJ5 | Ndfip |
| -0,947799047 | Q95R28;Q8IP47;( | Q95R28;Q8IP47;Q9VJP8;C | yuri |
| 0,2213281 | A0A077HCY9;A( | A0A077HCY9;A0A077HC | Adh |
| -0,041765849 | A0A0B4JCS1;A0. | A0A0B4JCS1;A0A0B4JCQ | Lpin |
| -0,296525319 | Q9XYM5;C1C3E | Q9XYM5;C1C3E9;Q5BIG7 | Su(var)2-10 |
| -0,10269928 | Q9XZL7;A0A0B4 | Q9XZL7;A0A0B4JCV4;A0 | tacc |
| 0,313086192 | A0A0B4JCZ4;Q2 | A0A0B4JCZ4;Q24547;K7Z | Syx1A |
| 0,145320892 | A0A0B4JD11;Q9 | A0A0B4JD11;Q9NJH0 | eEF1gamma |
| -0,125912984 | A0A0B4JD16;B7 | A0A0B4JD16;B7YZF9;B7 | ATP8A |
| -0,327354431 | A0A0B4KGP6;A( | A0A0B4KGP6;A0A0B4KF | Fmr1 |
| -0,22876358 | Q8IHB0;Q9VEX2 | Q8IHB0;Q9VEX2;Q8INB6 | gish |
| -0,003927867 | A0A0B4K6I5;Q9 | A0A0B4K6I5;Q9VN45 | spartin |
| -0,304035187 | A0A0B4LHZ6;A( | A0A0B4LHZ6;A0A0B4K6 | CG6051 |
| 0,021572749 | A0A0B4K6N4;O4 | A0A0B4K6N4;O46231;E1J | tmod |
| -0,006015142 | Q1EBY4;A0A0B4 | Q1EBY4;A0A0B4KEF4;Q( | Dscam1 |
| 0,161166509 | A0A0B4K6T1;A0 | A0A0B4K6T1;A0A0B4K7 | lig |
| -0,353052139 | A0A0B4K6U6;Q( | A0A0B4K6U6;Q08473-3;Q | sqd |
| 0,051629384 | Q0E9B2;A1Z904; | Q0E9B2;A1Z904;E8NH80; | Dyb |
| 0,165397008 | A0A0B4K730;E2 | A0A0B4K730;E2QC90;E0 | ced-6 |
| 0,775526683 | A0A0B4KF91;A0 | A0A0B4KF91;A0A0B4K7( | unc-104 |
| -0,872617722 | A0A0C4DHD5;Q | A0A0C4DHD5;Q7JMZ7;Q | sm |
| -0,173225403 | A1ZB78;A1ZB77 | A1ZB78;A1ZB77;Q7JRI6;( | CG5174 |
| -0,19305865 | A0A0B4K7G4;P1 | A0A0B4K7G4;P13469;Q8I | mod |
| 0,062444687 | B7YZQ7;A0A0B4 | B7YZQ7;A0A0B4K7G9;B9 | Nurf-38 |
| -0,287956874 | A0A0B4K7H0;Q7 | A0A0B4K7H0;Q7KN74 | Ttc7 |
| 0,151290258 | D3DMF2;Q9VBU | D3DMF2;Q9VBU7;A0A0E | Nup358 |
| -0,1438694 | Q25B55;A1Z7V1; | Q25B55;A1Z7V1;A8DY79 | brp |
| -0,486487071 | H5V888;C8VV04 | H5V888;C8VV04;C7LA80 | Pfk |
| 0,238450368 | A1Z8W9;A0A0B4 | A1Z8W9;A0A0B4K7N0;A | garz |
| 0,199821472 | A0A0B4KHS1;A( | A0A0B4KHS1;A0A0B4K7 | crb |
| 0,373964945 | A0A0B4K7Z5;Q7 | A0A0B4K7Z5;Q7KLV9;Q7 | Rpn6 |
| 0,139200211 | A0A0B4K851;C4 | A0A0B4K851;C4IY22;Q4Z | RanBPM |
| 0,212141673 | A0A0B4K885;Q9 | A0A0B4K885;Q9GYV5;B6 | key |
| -0,28523763 | A1Z6Q1;A0A0B4 | A1Z6Q1;A0A0B4KEE4;Q8 | koi |
| -0,073319753 | A0A0B4KEH0;P2 | A0A0B4KEH0;P29310;P29 | 14-3-3zeta |
| -0,259952545 | A0A0B4KEI5;Q9 | A0A0B4KEI5;Q9V9K7;E8 | Ars2 |
| -0,268763224 | A1Z8M2;A0A0B4 | A1Z8M2;A0A0B4KFE3;A( | tou |
| -0,520296733 | A0A0B4KEQ1;I0 | A0A0B4KEQ1;I0B8M2;B6 | Hdc |
| -0,805203756 | A0A0B4KEW6;Q | A0A0B4KEW6;Q7KLE5;A | Amph |
| -0,231653214 | A0A0B4KEX6;Q9 | A0A0B4KEX6;Q9W205;A | jbug |

| | | | |
|---|---|---|---|
| 0,14119339 | A0A0B4KF84;A0 | A0A0B4KF84;A0A0B4KG | Strn-Mlck |
| -0,159568151 | Q2XY38;Q7JQL5 | Q2XY38;Q7JQL5;A0A0B4 | Atg9 |
| 0,727888107 | A8DY98;Q8T0M | A8DY98;Q8T0M2;Q6AW | CG30015 |
| -0,773300171 | A0A0B4KFB8;Q | A0A0B4KFB8;Q95U01;C0 | CG42724 |
| -0,058661779 | C1C3H4;B7Z0U7 | C1C3H4;B7Z0U7;A0A0B4 | lap |
| -0,307093302 | A0A0B4KGE6;A | A0A0B4KGE6;A0A0B4K | Rpn8 |
| 0,233994166 | A0A0B4KFN1;Q | A0A0B4KFN1;Q6NQX0;A | Asx |
| 0,259070714 | A0A0B4KH36;A | A0A0B4KH36;A0A0B4KF | Bruce |
| 0,160792033 | D3DMK5;A0A0B | D3DMK5;A0A0B4KFT3;A | Pkc53E |
| 0,323907216 | A0A0B4KFZ9;P8 | A0A0B4KFZ9;P84040 | His4r |
| -0,686414083 | Q8SY21;A0A0B4 | Q8SY21;A0A0B4KG94;Q | CG6908 |
| -0,059514364 | A0A0B4KHB1;Q | A0A0B4KHB1;Q7KSE8;A | osa |
| 0,94260025 | A0A0B4KH12;A | A0A0B4KH12;A0A0B4K | bon |
| -0,501359304 | A0A0B4KGG9;A | A0A0B4KGG9;A0A0B4K | CtBP |
| -0,018002192 | A0A0B4KGX1;Q | A0A0B4KGX1;Q01989-2; | jar |
| -0,183801015 | A0A0B4KH25;P0 | A0A0B4KH25;P08985 | His2Av |
| -0,194710414 | A0A0B4KH34;P2 | A0A0B4KH34;P22464-3;P | AnxB9 |
| 0,46948878 | Q9VB04;Q7KRX | Q9VB04;Q7KRX5;C8VV4 | btz |
| 0,308148702 | B7Z0L1;B7Z0L0; | B7Z0L1;B7Z0L0;Q9VES3; | Fas1 |
| 0,093039195 | A0A0B4KHE9;A | A0A0B4KHE9;A0A0B4KI | Rpn2 |
| -0,071809769 | Q8T405;Q7KS06; | Q8T405;Q7KS06;A0A0B4 | Nmnat |
| -0,168020884 | A0A0B4KHH8;A | A0A0B4KHH8;A0A0B4K | Syp |
| -0,287991842 | A0A0B4KHJ1;P2 | A0A0B4KHJ1;P25867 | eff |
| -0,088445028 | G4LU37;A0A0B4 | G4LU37;A0A0B4KHL2;Q | gammaCOP |
| -0,674669266 | A0A0B4KHQ5;C | A0A0B4KHQ5;C6TP30;B7 | koko |
| 0,035598755 | Q9VDZ1;A0A0B | Q9VDZ1;A0A0B4KHR2 | CG5555 |
| 0,366898855 | A0A0B4KI24;Q2 | A0A0B4KI24;Q27415-2;Q | Nlp |
| 0,298816681 | Q9V9Y9;A0A0B | Q9V9Y9;A0A0B4KI68 | spn-F |
| -0,684208552 | A0A0B4LF00;A1 | A0A0B4LF00;A1Z833;A0 | CG1407 |
| -0,514316559 | A8DY82;E2QCN | A8DY82;E2QCN4;A8DY8 | Not1 |
| 0,203592936 | A0A0B4LF26;Q7 | A0A0B4LF26;Q7K550 | eIF3j |
| -0,500627518 | C7LA61;A0A0B4 | C7LA61;A0A0B4LFF9;A0 | Pcf11 |
| 0,25213178 | Q8T085;A1Z8S4; | Q8T085;A1Z8S4;A0A0B4 | Mppe |
| 0,064196269 | A0A0B4LF57;C6 | A0A0B4LF57;C6SUZ2;P6 | Cam |
| -0,53490003 | A0A0B4LFA6;Q7 | A0A0B4LFA6;Q7K2G1-2 | Rpn13 |
| -0,572117488 | A0A0B4LFI7;G2 | A0A0B4LFI7;G2J629;A8D | Nox |
| 0,55895551 | A0A0B4LFM0;P4 | A0A0B4LFM0;P41044 | Cbp53E |
| 0,019978205 | A0A0B4LFU5;Q | A0A0B4LFU5;Q8IGJ0-1;Q | stmA |
| 0,186386108 | A0A0B4LG05;P2 | A0A0B4LG05;P25823 | tud |
| -0,586328506 | A0A0B4LG36;Q0 | A0A0B4LG36;Q06943 | HmgZ |
| -0,145432154 | A0A0B4LG79;B7 | A0A0B4LG79;B7YZM9;B | Sdc |
| 0,852387746 | A0A0B4LGD3 | A0A0B4LGD3 | kcc |
| -0,127180099 | Q9VNC3;A0A0B | Q9VNC3;A0A0B4LGM5;A | Hcs |
| 0,068856557 | A0A0B4LGS4;P5 | A0A0B4LGS4;P54611 | Vha26 |
| 0,605513891 | F3YDF6;A0A0B4 | F3YDF6;A0A0B4LH71;Q9 | Sap47 |
| 0,088474909 | A0A0B4LHF9;R4 | A0A0B4LHF9;R4HZM7;Q | Sirt2 |
| -0,01313146 | A0A0B4LHT3;Q | A0A0B4LHT3;Q9VAS7 | Inx3 |
| -0,652827581 | A0A0B4LHV8;Q | A0A0B4LHV8;Q9VC57 | atl |
| 0,046912511 | A0A0B4LIJ0;P48 | A0A0B4LIJ0;P48601 | Rpt2 |
| -0,756285985 | C0PTW9;Q9W1T | C0PTW9;Q9W1T6;Q6NR5 | side-V |

| | | | |
|---|---|---|---|
| -0,202126821 | A0A0C4DHE6;Q9 | A0A0C4DHE6;Q9VFS5;Q9 | flfl |
| 0,416992188 | Q9GT67;Q8WR09 | Q9GT67;Q8WR09;A1ZA89 | Dg |
| -0,474254608 | A0A0C4DHG5;P2 | A0A0C4DHG5;P26686-3;P | B52 |
| 0,070472717 | Q9V3D8;Q9NHV | Q9V3D8;Q9NHV8;Q8IH82 | grsm |
| -0,09654363 | M9PJQ3;A0A0C5 | M9PJQ3;A0A0C5KMR0;M | RhoGAPp190 |
| -0,374919256 | M9PIS3;A0A0F67 | M9PIS3;A0A0F6T1J9;P415 | Pgd |
| -0,393304825 | Q7KSM5;Q7KJV | Q7KSM5;Q7KJV5;A0ANV | Aos1 |
| -0,292511622 | Q9U5W4;A0ANZ | Q9U5W4;A0ANZ0;A0ANY | nop5 |
| 0,417674383 | A0T1Z5;A0T1Z4; | A0T1Z5;A0T1Z4;Q9VFU8 | yrt |
| 0,323987961 | A1A701 | A1A701 | CG34138 |
| 0,086640676 | Q9GPI0;A1Z6H7; | Q9GPI0;A1Z6H7;Q6NP18 | Gp210 |
| 0,031536738 | Q960C7;A1Z729; | Q960C7;A1Z729;C1C590;C | CG2064 |
| -0,021767934 | Q8MSH0;A1Z765 | Q8MSH0;A1Z765;D3DN02 | CG12159 |
| 0,649676641 | Q8IGW5;A1Z7Q8 | Q8IGW5;A1Z7Q8;Q95SG9 | alc |
| 0,000403722 | A1Z8K7 | A1Z8K7 | Tsp47F |
| -0,087120056 | A1Z8S6;Q95U25 | A1Z8S6;Q95U25 | pds5 |
| 0,353517532 | A1Z8U4;C8VV32 | A1Z8U4;C8VV32;Q7KKI0 | CCT5 |
| 0,022843043 | A1Z968;Q5KTT4 | A1Z968;Q5KTT4;Q86NR8 | NAT1 |
| -0,386585236 | Q95TH2;A1Z971; | Q95TH2;A1Z971;Q8T0T5 | CG17019 |
| 0,301736832 | A1ZAB5;Q8MQR | A1ZAB5;Q8MQR3;A0A0B | clu |
| -0,325607936 | Q960R6;L0CRT5; | Q960R6;L0CRT5;L0CRT0; | Psi |
| 0,266143163 | Q8T9F0;A1ZAM0 | Q8T9F0;A1ZAM0 | CG6665 |
| -0,001785278 | C7LAA5;A8E6M | C7LAA5;A8E6M1;A1ZBB | CG30122 |
| -0,086358388 | A1ZBL7;E1JGM9 | A1ZBL7;E1JGM9;Q963E6 | par-1 |
| -0,008041382 | A2RVG4;E2QCY | A2RVG4;E2QCY5;A4VCL | CG34342 |
| 0,326533635 | Q8IQZ1;A3FP35; | Q8IQZ1;A3FP35;Q86B44;X | Gss1 |
| 0,557689031 | A4UZL5;P41043; | A4UZL5;P41043;A0A0B4K | GstS1 |
| 0,17216746 | Q8MLP1;Q7YZB | Q8MLP1;Q7YZB2;Q59E61 | Slik |
| -0,035184224 | Q8SZG1;A4UZX7 | Q8SZG1;A4UZX7;Q27367 | crq |
| -0,145624797 | A4UZZ4;Q9XTL9 | A4UZZ4;Q9XTL9 | GlyP |
| 0,469472885 | M9MRD7;D1Z388 | M9MRD7;D1Z388;A4V0D | Slob |
| -0,304604212 | A4V0N4;Q27331; | A4V0N4;Q27331 | Vha68-2 |
| -0,160561879 | A4V134;Q00168- | A4V134;Q00168-3;Q00168 | CaMKII |
| 0,010286331 | A4V303;P48605 | A4V303;P48605 | CCT3 |
| 0,085074743 | A4V391;P12613 | A4V391;P12613 | CCT1 |
| 0,172186534 | C8VV67;A4V3J6 | C8VV67;A4V3J6;A4V3J5; | Hrb98DE |
| 0,496967951 | A4V3Q6;P05303 | A4V3Q6;P05303 | eEF1α2 |
| -0,202894847 | A4V4A1;Q960Z0 | A4V4A1;Q960Z0 | Klp10A |
| -0,314136505 | A4V4F2;M9PHT0 | A4V4F2;M9PHT0;F2FB46 | Flo2 |
| 0,362300873 | A4V4J0;M9PJN9; | A4V4J0;M9PJN9;E1JJA5;A | shi |
| -0,083797455 | E8NH77;E6PBX0 | E8NH77;E6PBX0;X2JLF7; | sw |
| -0,5539608 | Q0GA05;Q0GA02 | Q0GA05;Q0GA02;A5YVH | gypsy\gag |
| 0,130152384 | A8DYI9;A1A6W0 | A8DYI9 | CG15118 |
| -0,097614288 | Q8IG91;L0MLL1 | Q8IG91;L0MLL1;Q8IMA6 | anne |
| 0,142523448 | O61380;A8DZ29; | O61380;A8DZ29;Q7YU50 | eIF4G1 |
| -0,274887085 | D6QLX6;A8E1U1 | D6QLX6;A8E1U1;P10090; | w |
| -0,887365341 | Q8IMC5;L0MLN | Q8IMC5;L0MLN1;L0MLK | Asator |
| -0,20868365 | A8E774;P25007;F | A8E774;P25007;F6J7B3 | Cyp1 |
| 0,138982773 | Q8T046;O18336;A | Q8T046;O18336;A8E777;O | Rab14 |

| | | | |
|---|---|---|---|
| 0,056551615 | Q7KV70;M9PBL( | Q7KV70;M9PBL6;A8JNJ6 | kst |
| -0,00554657 | M9PBN6;M9PE9. | M9PBN6;M9PE93;M9PHB | ens |
| -0,296110789 | M9PBP3;M9PE52 | M9PBP3;M9PE52;M9NE1( | Cip4 |
| 0,805746714 | Q7KU95;A8JNM4 | Q7KU95;A8JNM4;X2JGD. | Ank2 |
| 0,49593544 | A8JNP2;P48610-3 | A8JNP2;P48610-3;P48610- | Argk |
| 0,146524429 | B7Z0G0;A8JNQ0 | B7Z0G0;A8JNQ0;A8JNQ1 | nudE |
| 0,000458399 | A8JQV2;A0A0B4 | A8JQV2;A0A0B4KFF3;Q8 | CG17816 |
| 0,000603994 | A8JQV8;A0A0B4 | A8JQV8;A0A0B4K6C7;Q7 | pyd |
| 0,109685262 | A8JR82;E2QD55; | A8JR82;E2QD55;A8JR81;I | Efa6 |
| 0,63643837 | A8JR89;O77237 | A8JR89;O77237 | Pli |
| -0,742694219 | X2JIM3;X2JCS0;( | X2JIM3;X2JCS0;Q9W406; | CG15894 |
| -0,077908198 | M9PH75;X2JEC4 | M9PH75;X2JEC4;A8JUZ9; | CG34417 |
| 0,118497213 | A8Y596;E8NHS9 | A8Y596;E8NHS9;A8Y5A0 | Gfat1 |
| -0,122267405 | Q9VWV6;O9735 | Q9VWV6;O97355;A9UNH | Tsf1 |
| 0,057908376 | Q8SXM3;D0Z769 | Q8SXM3;D0Z769;A9UNH | RpII215 |
| -0,376626333 | Q59E23;M9PBV1 | Q59E23;M9PBV1;C0IN41; | Hn |
| 0,496315638 | Q9VIV2;B1NLF4 | Q9VIV2;B1NLF4;B1NLF3 | swm |
| 0,015522639 | E1JJE6;B3DNA0; | E1JJE6;B3DNA0;M9PDU2 | OtopLa |
| 0,278931936 | B5RIR1;Q24046 | B5RIR1;Q24046 | nrv1 |
| 0,206033071 | B5RIU6;Q8T390 | B5RIU6;Q8T390 | EndoA |
| -0,65471077 | Q8IMJ0;Q8IGI8;E | Q8IMJ0;Q8IGI8;B5RJE5;Q | CG31030 |
| -0,548693975 | B6IDQ5;M9PED6 | B6IDQ5;M9PED6;Q9VRJ9 | CG4603 |
| 0,649041494 | B6IDW0;Q7KTG4 | B6IDW0;Q7KTG4;Q9VLB | Oatp30B |
| -0,152224859 | B6V6Z8 | B6V6Z8 | - |
| -0,029253006 | B7FNJ7;Q960X8; | B7FNJ7;Q960X8;C9QP23;( | Hrs |
| -0,424825668 | B7YZR7;E1JGK5 | B7YZR7;E1JGK5;Q9V895 | Mapmodulin |
| -0,203679403 | E1JHK0;M9PG37 | E1JHK0;M9PG37;Q7KT48 | CLIP-190 |
| 0,043045044 | B7YZY9;Q9Y0H4 | B7YZY9;Q9Y0H4 | Su(dx) |
| -0,390968959 | Q9VQL7;B7Z001 | Q9VQL7;B7Z001;B6IDT3 | FASN1 |
| -0,069739024 | Q9Y1L3;Q9VPL6 | Q9Y1L3;Q9VPL6;M9NEL. | kis |
| -0,538419724 | B7Z031;Q8SXU4 | B7Z031;Q8SXU4 | pes |
| -0,249115626 | B7Z090;M9PFS2; | B7Z090;M9PFS2;M9PG70; | Psn |
| 0,289633433 | Q7KUB0;Q8IQA7 | Q7KUB0;Q8IQA7;Q9VSI6 | Idh |
| -0,197259903 | O18335;B8A3X3; | O18335;B8A3X3;Q8IGH2 | Rab11 |
| 0,219325384 | Q8IRH4;B8A420; | Q8IRH4;B8A420;I0B1P6;D | Ptp61F |
| -0,218716939 | C0MJE4;Q24572; | C0MJE4;Q24572;E1JIL4 | Caf1-55 |
| -0,055985133 | C0PTW6;X2JCI0; | C0PTW6;X2JCI0;M9PJ19;. | Fas2 |
| -0,300689061 | Q0KHQ6;C3KGL | Q0KHQ6;C3KGL9;X2JED' | Pfrx |
| 0,330814362 | Q86NM8;C4IY07 | Q86NM8;C4IY07;Q24276 | Cdc37 |
| 0,258352915 | E2QD63;Q86DS1 | E2QD63;Q86DS1;C4NYP8 | HIP-R |
| -0,700039546 | C5WLN6;Q03751 | C5WLN6;Q03751-3;Q0375 | Csp |
| 0,189610799 | Q7K549;C6SUV9 | Q7K549;C6SUV9 | mlt |
| -0,511991501 | Q9VBS7;I0E2K0; | Q9VBS7;I0E2K0;C6TP58;( | CG10550 |
| -0,362681071 | C6TP60;P48596-2 | C6TP60;P48596-2;E4NKN. | Pu |
| 0,042853673 | X2J9Z8;Q9U4H0; | X2J9Z8;Q9U4H0;Q9W5E8 | CG13366 |
| 0,046094259 | C7LA75;P11147;( | C7LA75;P11147 | Hsc70-4 |
| -0,861342112 | Q95T09;Q9VVG3 | Q95T09;Q9VVG3;Q8IQQ6 | blot |
| -0,068384171 | Q9VGZ3;Q9NFX. | Q9VGZ3;Q9NFX2;C7LA9 | Irp-1B |
| 0,174695969 | C8VV03;Q7KRW | C8VV03;Q7KRW8-3;Q7KI | CG1646 |
| -0,278261185 | C8VV14;F3YDA( | C8VV14;F3YDA0;P07764; | Ald1 |

| | | | |
|---|---|---|---|
| 0,033098857 | Q95SN6;Q9VG39 | Q95SN6;Q9VG39;C8VV48 | kar |
| -0,007669449 | Q961R8;Q9VUK8 | Q961R8;Q9VUK8;C9QP25 | GlyRS |
| 0,046915054 | M9MS72;M9MS3 | M9MS72;M9MS37;H1UUI | Pka-R1 |
| -0,276567459 | Q9VW48;Q961A8 | Q9VW48;Q961A8;Q7KUT | Papss |
| -0,483795802 | Q8IQN2;Q8MQK | Q8IQN2;Q8MQK2;Q9VUY | ClC-c |
| 0,053210576 | Q9VVC8;Q8IHE7 | Q9VVC8;Q8IHE7;C9QPB0 | CG6664 |
| -0,581723531 | C9QPG6;Q9V776 | C9QPG6;Q9V776 | Cyp317a1 |
| -0,38716507 | Q961H6;Q9VM10 | Q961H6;Q9VM10;C9QPI3 | santa-maria |
| -0,203386943 | O96607;Q9VM94 | O96607;Q9VM94;M9PB54 | homer |
| 0,290957133 | M9PG83;D0IQJ8; | M9PG83;D0IQJ8;Q03751 | Csp |
| -0,170609156 | M9PD27;D1Z373 | M9PD27;D1Z373;Q9VPH7 | eRF1 |
| -0,186454137 | X2JB48;D1Z385;( | X2JB48;D1Z385;Q26365-2 | sesB |
| 0,649697622 | M9NHC5;D2NUF | M9NHC5;D2NUF4;Q9VW | CG14207 |
| -0,17809995 | Q9XYZ9;D2NUG | Q9XYZ9;D2NUG2 | GstE12 |
| -0,262961706 | Q7JZD3;D3DMK | Q7JZD3;D3DMK3;D6W4V | Eb1 |
| -0,759730021 | Q8SX89;D3DML4 | Q8SX89;D3DML4 | kuk |
| -0,040531794 | D3DML9;D3DMM | D3DML9;D3DMM7;M9NE | nuf |
| 0,855644862 | X2JEX8;D3DMQ | X2JEX8;D3DMQ1;P06607 | Yp3 |
| -0,665859858 | D3DMV8;M9NDI | D3DMV8;M9NDR2;M9PD | BicD |
| -0,360349655 | D3DMZ9;Q9V429 | D3DMZ9;Q9V429-2;Q9V4 | Trx-2 |
| 0,203542074 | M9PC65;Q8IPL3; | M9PC65;Q8IPL3;D3PFH0; | CG34126 |
| 0,500338236 | M9ZYW3;D9PTS | M9ZYW3;D9PTS4;P40417 | rl |
| -0,302903493 | E1JGK7;Q7KRI2; | E1JGK7;Q7KRI2 | lolal |
| -0,113047282 | Q95U75;E1JGN0; | Q95U75;E1JGN0;Q9V8V8 | par-1 |
| -0,186900457 | E1JGX2;P32865 | E1JGX2;P32865 | Gprk1 |
| -0,222643534 | L0CQ56;L0CRN5 | L0CQ56;L0CRN5;L0CRY8 | Hrb27C |
| 0,953753153 | G4LU68;E1JHC4; | G4LU68;E1JHC4;E1JHC3; | Ggamma30A |
| -0,135819117 | Q5BHX9;E1JHK9 | Q5BHX9;E1JHK9 | CG10283 |
| -0,399835587 | E1JHX5;Q7K0L4 | E1JHX5;Q7K0L4 | CG7611 |
| 0,354301453 | M9NDT7;M9NG1 | M9NDT7;M9NG14;M9PI8 | Dab |
| -0,167468389 | E1JI64;M9PEG9;I | E1JI64;M9PEG9;M9PEQ0; | syd |
| -0,512857437 | E1JIE5;Q9VS49 | E1JIE5;Q9VS49 | CG43781 |
| -0,187161128 | E1JIJ5;P31409 | E1JIJ5;P31409 | Vha55 |
| -0,988871257 | E1JIK0;P48810 | E1JIK0;P48810 | Hrb87F |
| -0,228103002 | E1JIZ1;E1JIZ0;A( | E1JIZ1;E1JIZ0;A0A0B4K7 | scrib |
| 0,24929746 | E1JJ33;E1JJ35;E1 | E1JJ33;E1JJ35;E1JJ32;Q8I | cpx |
| 0,365037918 | E1JJ37;Q8IPM8-3 | E1JJ37;Q8IPM8-3 | cpx |
| -0,074445089 | E1JJ73;O97067 | E1JJ73;O97067 | CG1307 |
| 0,189229329 | Q8IRV7;M9NGK | Q8IRV7;M9NGK3;X2JDK | trol |
| -0,063085556 | X2JDR3;X2JDB3 | X2JDR3;X2JDB3;E1JJC3;2 | sdk |
| -0,09511439 | X2JCI4;E1JJD5;P | X2JCI4;E1JJD5;P13217-2;I | norpA |
| -0,034812927 | E1JJF9;E1JJF8;M | E1JJF9;E1JJF8;M9NEZ5;P | Nrg |
| 0,328491211 | E1JJN1;P35128 | E1JJN1;P35128 | ben |
| 0,980149587 | E2QC56;Q7K4Q5 | E2QC56;Q7K4Q5 | CG10417 |
| 0,082847595 | E2QCP0;P84249;I | E2QCP0;P84249;P02299;Q | His3.3A |
| -0,339252472 | E2QD65;P39018 | E2QD65;P39018 | RpS19a |
| 0,922742208 | E2QD75;Q7K4N3 | E2QD75;Q7K4N3 | Cbp80 |
| 0,018973033 | Q6NR03;Q9W3H | Q6NR03;Q9W3H7;Q0KHU | sdt |
| 0,047887166 | E6EK15;Q9V4C7 | E6EK15;Q9V4C7;Q59DQ0 | PMCA |
| 0,540541331 | E6EK18 | E6EK18 | PMCA |

| | | | |
|---|---|---|---|
| 0,258195241 | Q9W0H2;E8NHA | Q9W0H2;E8NHA2;Q8MQ | Herc4 |
| 0,076588313 | M9MS06;F0JAG6 | M9MS06;F0JAG6;M9PGA | Actn |
| 0,036614736 | F0JAN1;Q9I7S8;A | F0JAN1;Q9I7S8 | Paics |
| -0,886797587 | F0JAN9;M9PEY9 | F0JAN9;M9PEY9;M9PC96 | RhoGAP68F |
| -0,642601013 | M9PFH0;M9PF27 | M9PFH0;M9PF27;Q95SE8 | Wbp2 |
| -0,87628301 | F3YDH0;P29844; | F3YDH0;P29844 | Hsc70-3 |
| -0,586239497 | F3YDH1;F3YDI6 | F3YDH1;F3YDI6;Q9VG58 | Hsp70Bbb |
| -0,039312363 | F6J1D0;P40423 | F6J1D0;P40423 | sqh |
| -0,295152664 | G3M3A2;Q29QY | G3M3A2;Q29QY2;Q06559 | RpS3 |
| 0,202562332 | G4LU41 | G4LU41 | pyd |
| 0,457036972 | Q95SF7;Q8IQ82;( | Q95SF7;Q8IQ82;G7H836;( | pst |
| 0,408327738 | Q9VXE9;Q494K8 | Q9VXE9;Q494K8;H0RNH | SMC3 |
| 0,239613851 | Q9VBI4;Q7KRZ3 | Q9VBI4;Q7KRZ3;H1UUJ5 | Exo84 |
| -0,426703771 | H1ZYJ5;H1ZYI2; | H1ZYJ5;H1ZYI2;H1ZYI0;I | chico |
| 0,13516744 | H5V852;M9PGC4 | H5V852;M9PGC4;M9PGV | sgg |
| -0,100053787 | Q8SWU9;H5V895 | Q8SWU9;H5V895;Q8T0U( | flw |
| -0,582407633 | H9XVL5;Q9V4A7 | H9XVL5;Q9V4A7 | PlexB |
| -0,342406591 | Q9VGH2;H9ZJL9 | Q9VGH2;H9ZJL9;Q6IDG7 | CG6959 |
| -0,182182948 | I1V4Y8;Q24368 | I1V4Y8;Q24368 | Iswi |
| -0,222053528 | I1V501;Q94517 | I1V501;Q94517 | HDAC1 |
| -0,195685069 | I1WYI4;P35220;C | I1WYI4;P35220;G7H851 | alpha-Cat |
| -0,273661296 | Q9NGX6;Q9I7M8 | Q9NGX6;Q9I7M8;Q8MSM | Ku80 |
| -0,338169098 | X2JH42;L0MQ04 | X2JH42;L0MQ04;Q05825; | ATPsynbeta |
| 0,29222552 | Q5LK13;L7EEU0 | Q5LK13;L7EEU0;O61305; | Dbp80 |
| -0,013637543 | M9MRD0;P35122 | M9MRD0;P35122;M9PE40 | Uch |
| -0,042658488 | X2JDP6;M9MRG | X2JDP6;M9MRG2;Q8T079 | Ndf |
| -0,7825044 | M9MRJ4;M9PEP8 | M9MRJ4;M9PEP8 | Msp300 |
| 0,131757736 | Q9VTH2;Q6NR26 | Q9VTH2;Q6NR26;X2J8Y9 | CG42671 |
| 0,403858185 | M9NFJ6;M9ND32 | M9NFJ6;M9ND32;Q0E8I8; | Shab |
| -0,416434606 | M9MSK4;M9MR | M9MSK4;M9MRR7;M9M | Sh3beta |
| 0,005173365 | M9NEX9;M9NCT | M9NEX9;M9NCT0;Q0E8F | CG5850 |
| 0,502180735 | M9NCV1;Q9VJD | M9NCV1;Q9VJD3 | fws |
| -0,121344248 | M9PDC7;O61639 | M9PDC7;O61639;Q8INU2; | Dap160 |
| -0,089366277 | M9ND57;Q9VS83 | M9ND57;Q9VS83;B3DNM | CG7546 |
| -0,260455449 | M9NDA6;Q8I7C3 | M9NDA6;Q8I7C3;M9NFH | Lasp |
| -0,100259145 | M9NDC3;O18388 | M9NDC3;O18388 | Fs(2)Ket |
| -0,195008596 | Q9VQV1;M9NDE | Q9VQV1;M9NDE3 | bark |
| -0,47435379 | Q9VLQ9;M9NDI3 | Q9VLQ9;M9NDI3 | Snx6 |
| 0,030864716 | M9NE66;X2J8V0 | M9NE66;X2J8V0;X2JCQ1 | nwk |
| -0,265274048 | M9NE68;P02516 | M9NE68;P02516 | Hsp23 |
| 0,145712535 | M9NEC7;M9NF1 | M9NEC7;M9NF11;Q9W2V | Hk |
| 0,014220556 | M9NEL1;Q9VXF | M9NEL1;Q9VXF1 | CanA-14F |
| 0,018256505 | M9NEV0;Q9VMJ | M9NEV0;Q9VMJ7 | lid |
| 0,151917775 | Q9VZ96;M9NF15 | Q9VZ96;M9NF15;M9PBP8 | axo |
| -0,076379776 | Q9VVA4;M9NFH | Q9VVA4;M9NFH8;M9PCU | CG9674 |
| 0,25655365 | M9NFP7;P16914- | M9NFP7;P16914-2;P16914 | elav |
| -0,296597799 | M9NG39;P41073- | M9NG39;P41073-3;P41073 | Pep |
| 0,096516927 | M9NGG5;Q9W59 | M9NGG5;Q9W596 | futsch |
| 0,379301707 | M9PB35;P20385- | M9PB35;P20385-2;P20385 | Cf2 |
| 0,232215881 | Q7KMK9;M9PB9 | Q7KMK9;M9PB90;Q9VKY | Cand1 |

| | | | |
|---|---|---|---|
| -0,087149302 | M9PBI5;P13395;M | M9PBI5;P13395;M9PDQ0; | alpha-Spec |
| -0,450122197 | Q3KN34;Q9W053 | Q3KN34;Q9W053;Q7Z1Y4 | zormin |
| 0,130382538 | M9PBK6;M9PDX | M9PBK6;M9PDX6;M9PH4 | prominin-like |
| -0,001349131 | M9PBL3;P02828 | M9PBL3;P02828 | Hsp83 |
| 0,311088562 | M9PE73;M9PEC8 | M9PE73;M9PEC8;M9PBQ | Dhc64C |
| -0,594609578 | M9PBS9;M9PEJ2 | M9PBS9;M9PEJ2;M9PHL4 | CG10103 |
| -0,903739293 | M9PBU2;P48554; | M9PBU2;P48554;M9PBH7 | Rac2 |
| 0,242225647 | M9PBU4;Q9VQG | M9PBU4;Q9VQG1 | Snx21 |
| -0,311127345 | M9PBU6;M9PEF | M9PBU6;M9PEF2;Q9VS8 | lqf |
| -0,135737101 | M9PBV6;Q9Y109 | M9PBV6;Q9Y109 | Nulp1 |
| -0,077985128 | M9PEJ8;M9PBW | M9PEJ8;M9PBW9;Q9VSL | rhea |
| 0,14455986 | Q9VQ94;M9PC99 | Q9VQ94;M9PC99 | Sec24CD |
| -0,110222499 | M9PFB2;M9PCB | M9PFB2;M9PCB7;M9PFF | tral |
| -0,311698278 | M9PCC1;O18640 | M9PCC1;O18640;M9PB67 | Rack1 |
| -0,35567983 | M9PCN6;P16554 | M9PCN6;P16554-2;P16554 | numb |
| 0,270383199 | Q8IPG9;M9PCW | Q8IPG9;M9PCW5;Q961I4; | Bsg |
| -0,155099233 | M9PD08;Q9VK8 | M9PD08;Q9VK85;P91944 | eRF3 |
| -0,715450287 | M9PD73;M9PDF | M9PD73;M9PDF6;Q9VIT9 | Tep4 |
| -0,227158229 | Q2PDR9;M9PDB | Q2PDR9;M9PDB2;Q8SXI6 | vari |
| -0,006792704 | M9PDK5;Q9V3J1 | M9PDK5;Q9V3J1-3;Q9V3 | VhaSFD |
| 0,378250122 | M9PDQ1;P15348 | M9PDQ1;P15348;Q6NR40 | Top2 |
| -0,554278692 | Q9W441;Q8MS1 | Q9W441;Q8MS10;M9PDV | Tsp5D |
| 0,388825734 | Q9Y151;Q9W0A | Q9Y151;Q9W0A1;M9PDW | drpr |
| 0,507256826 | M9PDX8;Q86NM | M9PDX8;Q86NM2 | nrv3 |
| -0,121957143 | M9PEE1;P91638; | M9PEE1;P91638;Q9VS62; | smid |
| -0,16852506 | M9PEG1 | M9PEG1 | CG44195 |
| -0,025996526 | M9PEL1;Q07327 | M9PEL1;Q07327 | Rop |
| 0,006570816 | M9PEU5;Q9VTE | M9PEU5;Q9VTE9 | APP-BP1 |
| -0,153631846 | M9PF16;M9PHG | M9PF16;M9PHG4 | beta-Spec |
| -0,114568075 | M9PF24;P46824;I | M9PF24;P46824;M9PFG7 | Klc |
| -0,156533559 | Q9VTK9;M9PI17 | Q9VTK9;M9PI17;Q8IQF8; | CG6084 |
| -0,000485738 | M9PF91;P61851 | M9PF91;P61851 | Sod1 |
| 0,593068441 | M9PID3;M9PG82 | M9PID3;M9PG82;M9PG2 | HIPP1 |
| -0,075815837 | M9PG40;Q8SYG | M9PG40;Q8SYG3;M9PDD | CG40045 |
| -0,115266164 | M9PG62;P25161; | M9PG62;P25161 | Rpn3 |
| 0,955953598 | M9PG76;P19889 | M9PG76;P19889 | RpLP0 |
| -0,46675237 | M9PGF7;O61307 | M9PGF7;O61307;O61307- | Ten-m |
| -0,423542658 | M9PGG4;P18824 | M9PGG4;P18824-2;P1882 | arm |
| 0,239465078 | M9PGT6;Q9W2U | M9PGT6;Q9W2U4 | PPP4R2r |
| -0,80777359 | M9PGW0;P27716 | M9PGW0;P27716 | ogre |
| 0,108888626 | O16125;M9PHL6 | O16125;M9PHL6;Q9VYV8 | bif |
| -0,644091288 | M9PHN8;X2JF40 | M9PHN8;X2JF40;Q27580 | Ahcy |
| -0,211243947 | Q6NR17;Q9VTY | Q6NR17;Q9VTY8;M9PI41 | sti |
| -0,15754954 | M9PI82;P91891 | M9PI82;P91891 | Mo25 |
| -0,437970479 | N0A138;Q9VET1 | N0A138;Q9VET1;Q9Y121 | CG10345 |
| -0,361392975 | N0A733;Q9V4S8 | N0A733;Q9V4S8 | CSN7 |
| -0,305820465 | O16043 | O16043 | Df31 |
| -0,257261912 | O18332;Q8MRA4 | O18332 | Rab1 |
| 0,241219203 | O18333;Q9U5D5 | O18333;Q9U5D5 | Rab2 |
| -0,331577301 | O18338;M9NEE8 | O18338;M9NEE8;D5AEN5 | Rab8 |

| | | | |
|---|---|---|---|
| 0,485621134 | O18353;Q7KVG9 | O18353;Q7KVG9;O18356 | Chi |
| 0,300231934 | O18413;Q9VA54 | O18413 | Rpt6 |
| -0,265182495 | Q7KK54;O44253 | Q7KK54;O44253 | Sema1b |
| 0,107253393 | X2JAB9;Q24584; | X2JAB9;Q24584;O46037 | Vinc |
| 0,40015475 | O46068;Q29R09;( | O46068;Q29R09;Q86M59;( | CG2924 |
| -0,19802475 | O46106 | O46106 | noi |
| -0,215243022 | O61491;O61491-2 | O61491;O61491-2 | Flo1 |
| -0,017606099 | O61604;Q8IQZ8;( | O61604;Q8IQZ8;Q8IQZ7;I | Fim |
| 0,128325144 | O61650;Q9VPN5; | O61650;Q9VPN5;Q867V1; | Stip1 |
| 0,152121226 | O61732;Q9W3E2 | O61732;Q9W3E2 | PIP82 |
| -0,593204498 | O62530 | O62530 | AP-2mu |
| -0,26889801 | Q8SX88;Q9VRV8 | Q8SX88;Q9VRV8;O76331 | Tnpo |
| 0,255429586 | O76742 | O76742 | Rab7 |
| -0,6434714 | O76912 | O76912 | crm |
| -0,14379247 | X2JG90;O76932 | X2JG90;O76932 | Pp4-19C |
| -0,278430303 | Q9VCV4;Q9NFX | Q9VCV4;Q9NFX3;O76934 | Irp-1A |
| -0,022830963 | Q5BIC0;O77062 | Q5BIC0;O77062 | Eaat1 |
| -0,28442955 | O77266;Q960D4;( | O77266;Q960D4;Q7K7B5 | CG4199 |
| -0,070626577 | Q9W0B8;O77285 | Q9W0B8;O77285 | alphaCOP |
| -0,065750758 | O77410;Q8MR88 | O77410;Q8MR88 | eIF3e |
| -0,33767128 | Q9VDF4;O96046 | Q9VDF4;O96046 | Cortactin |
| 0,226852417 | Q94883;Q86NK7; | Q94883;Q86NK7;O96083 | Dref |
| -0,493022919 | O96671;Q9VU53; | O96671;Q9VU53;Q0E8F0 | caps |
| -0,276131948 | O96692;Q95SU2;. | O96692;Q95SU2;A0A0B4I | Rap2l |
| -0,288178762 | O96739 | O96739 | - |
| 0,17099762 | O96967;Q9VKE2 | O96967;Q9VKE2 | Crys |
| 0,600256602 | Q7JXZ2;O97121 | Q7JXZ2;O97121 | Eip55E |
| -0,170001984 | O97477 | O97477 | Inos |
| -0,14078776 | P02283 | P02283 | His2B |
| 0,040933609 | X2JGG6;P02517 | X2JGG6;P02517 | Hsp26 |
| -0,280585607 | X2JC82;P02518 | X2JC82;P02518 | Hsp27 |
| 0,725573222 | X2JD55;Q29QE3; | X2JD55;Q29QE3;P02843;X | Yp1 |
| -0,390599569 | P05205 | P05205 | Su(var)205 |
| -0,154006958 | X2JCP8;P10987;F | X2JCP8;P10987;P02572 | Act5C |
| -0,017054876 | P12982;C0MKX3 | P12982;C0MKX3;B4F4X2; | Pp1-87B |
| -0,205640793 | P13496 | P13496 | DCTN1-p150 |
| 0,084082921 | P13677 | P13677 | inaC |
| 0,099591573 | P14199 | P14199 | ref(2)P |
| -0,263017654 | P16378-2;P16378; | P16378-2;P16378;P20353 | Galphao |
| 0,177883148 | P17210;Q6NNT8 | P17210 | Khc |
| -0,074314753 | P17336 | P17336 | Cat |
| -0,606520335 | P17917 | P17917 | PCNA |
| 0,079725901 | P18106;A0A0B4K | P18106;A0A0B4KGK6;A0. | FER |
| -0,198484421 | P21187 | P21187 | pAbp |
| -0,076382955 | P22465 | P22465 | AnxB10 |
| -0,387967428 | P22465-2;D5AEP( | P22465-2 | AnxB10 |
| 0,387374878 | P22769 | P22769 | Prosalpha4 |
| -0,283908208 | Q8IMF5;P23226-3 | Q8IMF5;P23226-3;P23226; | Map205 |
| 0,406358083 | P23625 | P23625 | Galphaq |

| | | | |
|---|---|---|---|
| 0,462556203 | P29829 | P29829 | Gbeta76C |
| -0,303776423 | P35381 | P35381 | blw |
| 0,384061813 | P36975;P36975-3 | P36975;P36975-3 | Snap25 |
| 0,30203565 | P40421 | P40421 | rdgC |
| -0,255697886 | P40797 | P40797 | pnut |
| 0,353918711 | X2JFR6;P41374 | X2JFR6;P41374 | eIF2alpha |
| 0,2754275 | P42325 | P42325 | Nca |
| 0,626515706 | P45437 | P45437 | betaCOP |
| -0,809963226 | P45594 | P45594 | tsr |
| -0,312939326 | S5M3P7;P45843 | S5M3P7;P45843 | st |
| 0,439539591 | P45888-2;P45888; | P45888-2;P45888;F6J178 | Arp2 |
| -0,435523351 | P45889 | P45889 | Arp1 |
| 0,324104309 | P49021-6;P49021- | P49021-6;P49021-7;P49021 | tim |
| 0,092371623 | P49415-2 | P49415-2 | Sdc |
| -0,390525818 | Q7KSG0;P50245; | Q7KSG0;P50245;Q9VZX9 | AhcyL2 |
| -0,008546829 | P50887 | P50887 | RpL22 |
| -0,042528788 | P52029 | P52029 | Pgi |
| 0,467206955 | Q541C3;P52485 | Q541C3;P52485 | Ubc2 |
| -0,087341944 | X2JAX3;P52486;( | X2JAX3;P52486;Q6NN75 | Ubc4 |
| 0,193518957 | P53997 | P53997 | Set |
| -0,284725189 | X2JGP4;P54399;I | X2JGP4;P54399;P54399-2 | Pdi |
| 0,456731796 | X2J6D4;P84029 | X2J6D4;P84029 | Cyt-c-p |
| -0,124637604 | Q4AB57;P84051 | Q4AB57;P84051 | His2A:CG33859 |
| 0,027130763 | Q00963;X4YME1 | Q00963 | beta-Spec |
| -0,11949412 | Q02645-2;A0A0B | Q02645-2;A0A0B4K849;A | hts |
| 0,063698451 | X2JFR1;Q8IR16;( | X2JFR1;Q8IR16;Q04047-2 | nonA |
| 0,163228989 | Q04134;Q6GKZ8 | Q04134;Q6GKZ8 | Doc\gag |
| -0,260462443 | X2JCF8;Q08180 | X2JCF8;Q08180 | rst |
| 0,130930583 | X2JE07;Q9W3R9 | X2JE07;Q9W3R9;Q8MYV | CG1677 |
| -0,062123617 | Q9XZE4;Q0E8E8 | Q9XZE4;Q0E8E8 | Mpcp2 |
| -0,077051163 | Q9VSY2;Q0E8G6 | Q9VSY2;Q0E8G6 | Cpsf5 |
| 0,061003367 | Q0E8Q7;A2VEL6 | Q0E8Q7;A2VEL6;E1JHH7 | Eato |
| 0,111516953 | Q961R5;Q0E980 | Q961R5;Q0E980 | CG8547 |
| -0,532334646 | Q0KHS6;X2JKC1 | Q0KHS6;X2JKC1;Q0KHS | Lsd-2 |
| -0,334643682 | Q0KI81;Q4QQ46; | Q0KI81;Q4QQ46 | CG42327 |
| -0,860048294 | Q24241;Q0KIE7;( | Q24241;Q0KIE7 | Ank |
| 0,567719777 | Q9XZL6;Q8IMC2 | Q9XZL6;Q8IMC2;Q9Y1J0 | Eph |
| -0,191837947 | Q1EC00;Q9VPM | Q1EC00;Q9VPM7;M9PAY | CG17078 |
| 0,279298782 | Q1RL12;Q24048- | Q1RL12;Q24048-2;A4V0B | nrv2 |
| -0,22155571 | Q23983 | Q23983 | alphaSnap |
| -0,608872096 | Q9W1G7;Q24150 | Q9W1G7;Q24150 | Nap1 |
| -0,228894552 | Q24185 | Q24185 | hook |
| -0,039708455 | X2JCY4;X2JGF5; | X2JCY4;X2JGF5;Q24212 | stnB |
| -0,106877645 | Q24253;Q960F2 | Q24253;Q960F2 | AP-1-2beta |
| -0,1669089 | Q24262 | Q24262 | flea/polyprotein |
| 0,508199692 | Q24307 | Q24307 | Diap2 |
| 0,639306386 | X2JCV2;Q24388 | X2JCV2;Q24388 | Lsp2 |
| -0,07172521 | Q24478 | Q24478 | Cp190 |
| -0,153879166 | Q8SYR3;Q24491 | Q8SYR3;Q24491 | Rsf1 |
| 0,025089264 | Q24492 | Q24492 | RpA-70 |

| | | | |
|---|---|---|---|
| -0,837848028 | Q27268;M9PC90; | Q27268;M9PC90 | Hel25E |
| -0,34485054 | Q9VZU9;Q6NLL: | Q9VZU9;Q6NLL3;Q29R24 | CG11537 |
| 0,050725937 | Q8T0C2;Q2PDW | Q8T0C2;Q2PDW1;Q9VPT | CG3662 |
| -0,048408508 | Q2XYF2;Q7JW0: | Q2XYF2;Q7JW03 | UbcE2H |
| 0,18588384 | Q32KD2 | Q32KD2 | egg |
| -0,544658025 | Q9W388;Q86B59 | Q9W388;Q86B59;Q7KVR | AP-1gamma |
| 0,12847964 | Q4V498;Q9VR48 | Q4V498;Q9VR48;Q4V4B8 | hoe2 |
| -0,07883962 | X2J9M9;Q4V5C6 | X2J9M9;Q4V5C6;X2JDZ6 | CG31729 |
| -0,450011571 | Q4V5X3;Q9VW7 | Q4V5X3;Q9VW73 | CG14184 |
| -0,222056707 | Q59DP8 | Q59DP8 | PMCA |
| -0,670853297 | Q59DP9 | Q59DP9 | PMCA |
| 0,631352107 | Q7JRF1;Q7JY00; | Q7JRF1;Q7JY00;Q5BI25;C | Syx6 |
| 0,045700709 | Q5BI50;Q961E0 | Q5BI50;Q961E0 | Cul4 |
| -0,446463267 | Q9V4C6;Q5BII0 | Q9V4C6;Q5BII0 | Zip102B |
| -0,038962046 | Q9VYT3;Q5U158 | Q9VYT3;Q5U158;Q8SZT9 | Nrd1 |
| 0,371039708 | Q9VUR4;Q6AW( | Q9VUR4;Q6AWG5;Q6NN | CG7656 |
| -0,156036377 | Q6AWH1;Q7PLG | Q6AWH1;Q7PLG1 | l(3)80Fg |
| 0,267458598 | Q7KVL7;Q9W27 | Q7KVL7;Q9W277;Q6AWI | Vps35 |
| -0,131154378 | Q6IDE2 | Q6IDE2 | CG3907 |
| -0,361489614 | Q6IDG3;S5MHR4 | Q6IDG3;S5MHR4;Q9XTM | Sec10 |
| 0,694883982 | Q6NL41;A8Y541 | Q6NL41;A8Y541 | lovit |
| -0,018929799 | Q8IML4;Q6NLA: | Q8IML4;Q6NLA3 | Vha100-1 |
| -0,036200205 | Q8IQH0;Q6NN26 | Q8IQH0;Q6NN26;Q94887 | Nrx-IV |
| -0,177216848 | Q9V4E7;Q6NND | Q9V4E7;Q6NND1;Q8IMC | Gat |
| 0,006989797 | Q6NNV7;Q5LJT( | Q6NNV7;Q5LJT0;Q6IDG8 | CG40485 |
| -0,225992203 | Q86G43;Q868A8; | Q86G43;Q868A8;Q867W7 | Nup98-96 |
| -0,231740316 | Q6NP91 | Q6NP91 | Tmem63 |
| 0,14020284 | Q9VL06;Q6NR00 | Q9VL06;Q6NR00 | Ufd4 |
| 0,052302043 | Q7JNE1 | Q7JNE1 | prod |
| -0,707700729 | Q7JQH9;Q9V3K9 | Q7JQH9;Q9V3K9 | whd |
| 0,104188919 | Q7JQI1 | Q7JQI1 | DUBAI |
| 0,255196253 | Q7JR34 | Q7JR34 | CG33144 |
| -0,239039103 | Q7JR91;Q95T37; | Q7JR91;Q95T37;Q7JRM9 | Nmda1 |
| 0,571266174 | Q8MQS0;Q7JRK | Q8MQS0;Q7JRK8;Q9VIG | tadr |
| 0,226907094 | Q7JVK6;Q6TMH | Q7JVK6 | trsn |
| -0,054075877 | Q7K010 | Q7K010 | Tsp42Ef |
| -0,151594798 | Q7K0S5 | Q7K0S5 | Gint3 |
| -0,098995209 | Q9VRD4;Q7K0S | Q9VRD4;Q7K0S8 | CG1532 |
| 0,320656459 | Q7K0X9 | Q7K0X9 | Syx7 |
| -0,012156169 | Q7K1H0 | Q7K1H0 | CG32066 |
| -0,361733119 | Q7K1L4 | Q7K1L4 | CG8468 |
| -0,578586578 | Q7K1U0 | Q7K1U0 | Arc1 |
| 0,012848536 | Q7K2B1 | Q7K2B1 | CG7222 |
| -0,346345901 | Q7K2E1 | Q7K2E1 | CG8839 |
| -0,207303365 | Q7K2G1 | Q7K2G1 | Rpn13 |
| 0,453054428 | Q9W0Z5;Q7K3E( | Q9W0Z5;Q7K3E0;Q95TB | Atf-2 |
| -0,088605881 | Q7K3J0 | Q7K3J0 | CCT8 |
| 0,122938792 | Q7K3Z3 | Q7K3Z3 | p47 |
| 0,119414647 | Q7K486 | Q7K486 | CG5721 |
| 0,560266495 | Q7K738 | Q7K738 | Ubc10 |

| | | | |
|---|---|---|---|
| -0,537733078 | Q9Y148;Q7KJA9 | Q9Y148;Q7KJA9 | sxc |
| 0,009684245 | Q7KJN8;A1Z9J0; | Q7KJN8;A1Z9J0;E1JH64;A | shot |
| -0,127077738 | Q7KUA4;Q7KJV | Q7KUA4;Q7KJV6;Q9VSD | Uba2 |
| -0,029643377 | Q7KK51;Q9W1Q | Q7KK51;Q9W1Q6;Q8MLI | CG3530 |
| 0,598903656 | Q7KMP8 | Q7KMP8 | Rpn9 |
| -0,179954529 | Q7KMQ0;M9PC1 | Q7KMQ0 | Rpt1 |
| -0,264035543 | Q9VRP3;Q7KMR | Q9VRP3;Q7KMR7 | Txl |
| 0,041855494 | Q7KN62;A0A0B4 | Q7KN62;A0A0B4LFZ4;D0 | TER94 |
| -0,066210429 | Q7KND8;Q95S25 | Q7KND8;Q95S25 | Mad1 |
| -0,377578735 | Q7KRR5;Q7KRR | Q7KRR5;Q7KRR5-2 | Pngl |
| -0,218205134 | Q7KRT4;Q9VA0 | Q7KRT4;Q9VA06 | stops |
| -0,453460693 | Q8MZ56;Q7KSA | Q8MZ56;Q7KSA4;Q9I7I6; | CG5191 |
| 0,104998906 | Q7YU32;Q7KT7 | Q7YU32;Q7KT70;Q86LF3 | Gli |
| 0,054942449 | Q7KTB3;Q8IP79; | Q7KTB3;Q8IP79;Q8IP80;C | Pect |
| 0,418207804 | Q7KTG2;Q8MRJ | Q7KTG2 | Apoltp |
| 0,822200775 | Q7KTW5 | Q7KTW5 | CG9391 |
| -0,282628377 | Q95TI4;Q7KTX7 | Q95TI4;Q7KTX7 | park |
| 0,21030426 | Q9VT60;Q7KUD | Q9VT60;Q7KUD4;Q29R2 | iPLA2-VIA |
| -0,200218201 | Q8T9D5;Q7KV2 | Q8T9D5;Q7KV27;Q9VYD | CG1640 |
| -0,118290583 | Q7KV34;Q9VYW | Q7KV34;Q9VYW8 | CG1561 |
| -0,247938792 | Q7KVP6;Q9W2I | Q7KVP6;Q9W2I0;B7YZM | CG42672 |
| -0,53045845 | Q7KVX5;Q9W4N | Q7KVX5;Q9W4N8 | Vap33 |
| -0,981904348 | Q7KY07 | Q7KY07 | Rab32 |
| -0,127964656 | Q7YU24;F6JB17; | Q7YU24;F6JB17 | Marf |
| -0,368760427 | Q9VRW4;Q7YU | Q9VRW4;Q7YU35 | Best2 |
| -0,31603241 | Q9VUX0;Q7YU5 | Q9VUX0;Q7YU53;M9PFN | CG5830 |
| 0,114106496 | X2J9B0;Q9VL24; | X2J9B0;Q9VL24;Q7YU59 | Npc1a |
| -0,1135451 | Q7YU85;A0A0B4 | Q7YU85;A0A0B4KI32;Q9 | cindr |
| 0,574736913 | Q867Z4-2;D3DM | Q867Z4-2;D3DMG1;P4228 | lola |
| 0,525176366 | Q868Z9-2;Q868Z | Q868Z9-2;Q868Z9-6;Q868 | Ppn |
| -0,335692088 | Q9VLL3;Q86BM | Q9VLL3;Q86BM5;Q9U7E | Akap200 |
| -0,17913119 | Q8T9D1;Q86BS3 | Q8T9D1;Q86BS3;Q5MAI1 | Chro |
| 0,1121521 | Q86DT8;Q86DT7 | Q86DT8;Q86DT7;Q86DT6 | STUB1 |
| -0,349711736 | Q86NN7;Q9VY9 | Q86NN7;Q9VY98;A9YHB | CG9941 |
| -0,074549357 | Q9VF74;Q86NV7 | Q9VF74;Q86NV7;Q95TY5 | CG34404 |
| -0,062116623 | Q9VR90;Q86P91; | Q9VR90;Q86P91 | Syx16 |
| -0,956481298 | Q86PM1;Q9VWI | Q86PM1;Q9VWI6 | kek5 |
| 0,354554494 | Q8I079 | Q8I079 | CG32640 |
| 0,133501689 | Q8IGE9;Q9VN21 | Q8IGE9;Q9VN21 | lost |
| -0,877806981 | Q9NFP0;Q9W28 | Q9NFP0;Q9W289;Q8IGK8 | GM130 |
| -0,545219421 | Q8T6I0;Q8IGN0; | Q8T6I0;Q8IGN0;Q8T8W3 | Past1 |
| -0,809011459 | Q8IPJ6;Q9Y0Z0; | Q8IPJ6;Q9Y0Z0;Q8IGT8 | mmy |
| 0,427939097 | Q8IMV6;Q8IGV1 | Q8IMV6;Q8IGV1;Q9VC3 | Saf-B |
| 0,114227931 | Q8IMG1;Q8IGY5 | Q8IMG1;Q8IGY5;Q9V9U | CG1910 |
| -0,46972847 | Q9VQE0;Q8IHG | Q9VQE0;Q8IHG0;Q8T9L0 | Drp1 |
| 0,849098206 | Q9V9R4;Q8IMF4 | Q9V9R4;Q8IMF4;Q8IMF3 | CG2201 |
| -0,134688059 | Q8IMI7;Q9VA91 | Q8IMI7;Q9VA91;Q3KN36 | RpS7 |
| 0,005053838 | Q9VHI8;Q8INP8; | Q9VHI8;Q8INP8;Q8INP9; | CG11980 |
| 0,752909978 | Q8IPC3;Q8SYF0; | Q8IPC3;Q8SYF0;Q8IPC5 | CYLD |
| -0,433748881 | Q8IPF5;Q9U7E6; | Q8IPF5;Q9U7E6 | Akap200 |

| | | | |
|---|---|---|---|
| -0,048782349 | Q9VMH3;Q9GQI | Q9VMH3;Q9GQR3;Q8IPK | stai |
| -0,566488902 | Q8IQD7;Q8IQD8; | Q8IQD7;Q8IQD8;Q9VT88 | scramb1 |
| 0,259307226 | Q8IQM5;Q960Q5 | Q8IQM5;Q960Q5 | CG7656 |
| -0,249866486 | Q9VXP1;Q8IR26 | Q9VXP1;Q8IR26 | CG32581 |
| -0,231772105 | Q8IRJ0;Q9W0R3; | Q8IRJ0;Q9W0R3;Q86PA5; | Ptpmeg |
| -0,126095454 | Q8MMD3;Q8MM | Q8MMD3;Q8MMD2;Q9W | Eps-15 |
| -0,245969772 | Q9VGB6;Q8MR4 | Q9VGB6;Q8MR44 | Pglym87 |
| 0,082865397 | X2JAF1;Q8MSS1 | X2JAF1;Q8MSS1 | lva |
| -0,203550339 | Q8MST1;Q9VB2 | Q8MST1;Q9VB21 | CG5639 |
| -0,425591787 | Q9VAX7;Q8MST | Q9VAX7;Q8MST5 | betaTub97EF |
| 0,476509094 | Q9VKC8;Q8MT2 | Q9VKC8;Q8MT29 | Tom70 |
| -0,125504176 | Q8SWR8;A0A0B | Q8SWR8;A0A0B4LH64;Q | Atx2 |
| -0,240176519 | Q8SWS2;Q9VYM | Q8SWS2;Q9VYM7;X2JEY | CG42258 |
| -0,172363917 | Q8SWU2;Q9VW | Q8SWU2;Q9VW13 | CG9330 |
| 0,235209147 | Q8SX35;L0MLM | Q8SX35;L0MLM5 | CG32850 |
| 0,15594546 | Q8SY33-2;Q8SY | Q8SY33-2;Q8SY33-3;Q8S | gw |
| -0,583914439 | Q9VS39;Q8SYN | Q9VS39;Q8SYN3 | CG14830 |
| 0,003844579 | Q8SYN6;Q9VSC | Q8SYN6;Q9VSC5 | CG8209 |
| 0,377142588 | Q9W3Q0;Q8SYS | Q9W3Q0;Q8SYS5;F6J915 | Rab39 |
| 0,137780507 | Q8SZU6;Q8IRG7 | Q8SZU6;Q8IRG7;M9MRY | CG12024 |
| -0,158706665 | Q8T0G8 | Q8T0G8 | Rab32 |
| 0,107290904 | Q8T0Q4 | Q8T0Q4 | shrb |
| 0,549893061 | Q9V3D9;Q8T3P8 | Q9V3D9;Q8T3P8 | Srp54k |
| -0,402272542 | Q8T4G5;Q8IQQ9 | Q8T4G5;Q8IQQ9 | CG6512 |
| -0,242949168 | Q8T6B9;A4V193; | Q8T6B9;A4V193;Q8T6B9- | hfp |
| -0,792164485 | Q9VSR8;T1W04 | Q9VSR8;T1W043;Q8T8U | CG5653 |
| 0,157606761 | Q8T8V5 | Q8T8V5 | La |
| -0,047470729 | Q8T9C5;Q9I7N0; | Q8T9C5;Q9I7N0;Q7KTD0 | MRP |
| 0,102877299 | Q8T9G8;T2FFB7; | Q8T9G8;T2FFB7;Q9VLM | AlaRS |
| -0,288245519 | Q94544;Q7JNK2 | Q94544 | sktl |
| 0,091721853 | Q95RR5;Q7K3H | Q95RR5;Q7K3H0;Q7JNX | Pi3K68D |
| 0,231421789 | Q9VEP6;Q95SP9 | Q9VEP6;Q95SP9 | AdSL |
| -0,155242284 | Q95T12;D8FT42; | Q95T12 | fwe |
| -0,079298019 | Q9VXI3;Q95TL8 | Q9VXI3;Q95TL8 | CG9911 |
| 0,056933721 | Q960E2;Q9VN44 | Q960E2;Q9VN44;Q86NQ0 | Karybeta3 |
| -0,227302551 | Q9VEH0;Q960Y | Q9VEH0;Q960Y8;Q95R41 | alt |
| -0,303155263 | Q967S7 | Q967S7 | HMS-Beagle\gag |
| 0,363572439 | Q99140 | Q99140 | kst |
| -0,274684906 | Q9VPY2;Q9GUB | Q9VPY2;Q9GUB1 | Plap |
| 0,238716761 | Q9I7K6 | Q9I7K6 | CG8223 |
| 0,205256144 | Q9I7V7;Q9I7V6; | Q9I7V7;Q9I7V6;Q8IHG2; | MESK2 |
| 0,738865534 | Q9NBD7 | Q9NBD7 | chb |
| -0,84196345 | Q9NGV4;Q9VM | Q9NGV4;Q9VM55;E1JHA | uif |
| 0,283946355 | Q9VFT4;Q9NH7 | Q9VFT4;Q9NH72 | rin |
| -0,432203293 | Q9VXV6;Q9NHD | Q9VXV6;Q9NHD7 | Scamp |
| -0,393395742 | Q9TVM2;Q8IH7 | Q9TVM2;Q8IH79 | emb |
| 0,14178594 | Q9U5W6 | Q9U5W6 | msps |
| 0,007904053 | Q9U6R9;C3KGI9 | Q9U6R9 | gammaSnap1 |
| 0,145868937 | Q9V3H2 | Q9V3H2 | Rpn11 |
| 0,055223465 | Q9V3H5;Q9Y0G | Q9V3H5;Q9Y0G0 | Cdk5alpha |

| | | | |
|---|---|---|---|
| -0,074480057 | Q9V3I2 | Q9V3I2 | Rab5 |
| 0,031478246 | Q9V3P3 | Q9V3P3 | REG |
| -0,601929347 | Q9V3V9 | Q9V3V9 | EndoGI |
| -0,217760086 | Q9V3W7 | Q9V3W7 | SF2 |
| -0,089200338 | Q9V3W9;Q9XZE | Q9V3W9;Q9XZE0;Q8IMB | Rad23 |
| -0,391344706 | Q9V3Y2 | Q9V3Y2 | CIAPIN1 |
| -0,185764949 | Q9V3Z4 | Q9V3Z4 | Rpn5 |
| -0,208262761 | Q9V426 | Q9V426 | vig |
| -0,470274607 | Q9V431 | Q9V431 | cass |
| -0,260210673 | Q9V434 | Q9V434 | AsnRS |
| 0,243317286 | Q9V4C8-6;Q9V4 | Q9V4C8-6;Q9V4C8;Q9V4 | Hcf |
| -0,72527949 | Q9VAC4 | Q9VAC4 | Nph |
| -0,177853902 | Q9VAD1;E4NKH | Q9VAD1;E4NKH3;E4NKH | CG7896 |
| 0,200926463 | Q9VAN0 | Q9VAN0 | CG11899 |
| -0,133230845 | Q9VAX8;Q86NS | Q9VAX8;Q86NS3 | CG4849 |
| -0,013142268 | Q9VB05 | Q9VB05 | ALiX |
| -0,206981023 | Q9VB23 | Q9VB23 | wdb |
| 0,940271378 | Q9VBF0;A0A0B4 | Q9VBF0;A0A0B4KH14 | CG5447 |
| -0,292200724 | Q9VBP9-2;Q9VB | Q9VBP9-2;Q9VBP9 | Npl4 |
| -0,166868846 | Q9VBQ5 | Q9VBQ5 | CG5112 |
| -0,107507706 | Q9VBX8 | Q9VBX8 | CG33095 |
| 0,178265889 | Q9VBY7;Q8IMT | Q9VBY7;Q8IMT8;Q6NL8 | veli |
| 0,052072525 | Q9VC31;Q4V4N | Q9VC31;Q4V4N9 | RabX4 |
| -0,883885066 | Q9VCF1 | Q9VCF1 | CG13603 |
| 0,084770838 | Q9VCT9 | Q9VCT9 | Usp12-46 |
| 0,189268748 | Q9VCZ2 | Q9VCZ2 | CG5346 |
| -0,221694311 | Q9VD30 | Q9VD30 | rdhB |
| 0,53347079 | Q9VD68 | Q9VD68 | CG6656 |
| -0,374340057 | Q9VE75;Q9VKF | Q9VE75 | Vha100-2 |
| 0,15942955 | Q9VEC8 | Q9VEC8 | Prx5 |
| 0,704423904 | Q9VF20 | Q9VF20 | Hmt-1 |
| -0,199752172 | Q9VFF0 | Q9VFF0 | UQCR-C1 |
| -0,359067917 | Q9VFP0 | Q9VFP0 | CG3061 |
| 0,028740565 | Q9VFP6;O76455 | Q9VFP6;O76455 | Ipp |
| -0,077817917 | Q9VFV9 | Q9VFV9 | Droj2 |
| 0,149030685 | Q9VG51 | Q9VG51 | Snx3 |
| -0,18172582 | Q9VGH5;Q8INJ6 | Q9VGH5;Q8INJ6;Q8IG99 | glo |
| 0,375424067 | Q9VGM1 | Q9VGM1 | Mrp4 |
| 0,044715246 | Q9VGR1;Q3ZAL | Q9VGR1;Q3ZAL1 | GCC88 |
| -0,07479922 | Q9VHC7;A0A0B | Q9VHC7;A0A0B4K6Z1 | rump |
| -0,372028351 | Q9VHC8 | Q9VHC8 | CG8149 |
| 0,534255981 | Q9VHI7 | Q9VHI7 | Iru |
| -0,335833232 | Q9VHL2 | Q9VHL2 | CCT7 |
| -0,093150457 | Q9VHN7;Q7KSU | Q9VHN7;Q7KSU6 | CG8036 |
| 0,189478556 | Q9VHX2 | Q9VHX2 | CG3223 |
| 0,025314331 | Q9VHX4 | Q9VHX4 | CG2767 |
| -0,272642771 | Q9VI20;Q3KN42 | Q9VI20;Q3KN42 | gzl |
| 0,142952601 | Q9VJ75 | Q9VJ75 | CG10413 |
| 0,093788783 | Q9VJD4 | Q9VJD4 | Sgt |
| -0,122402827 | Q9VK28;C0PTX | Q9VK28;C0PTX5;Q95S21 | CG16974 |

| | | | |
|---|---|---|---|
| -0,697779338 | Q9VK59 | Q9VK59 | CG5787 |
| -0,315273285 | Q9VK69 | Q9VK69 | CCT4 |
| 0,083281835 | Q9VKI8 | Q9VKI8 | CG6287 |
| -0,008488337 | Q9VKW1 | Q9VKW1 | Wdfy2 |
| -0,343263626 | Q9VKZ8 | Q9VKZ8 | Usp14 |
| -0,205095927 | Q9VL00 | Q9VL00 | CG4968 |
| -0,285203934 | Q9VL16 | Q9VL16 | CG5676 |
| -0,317925771 | Q9VL18;Q9VL18 | Q9VL18;Q9VL18-2 | eEF1delta |
| -0,292352041 | X2J5E8;Q9VL78 | X2J5E8;Q9VL78 | Fkbp59 |
| 0,11223793 | Q9VLB7;Q24349 | Q9VLB7;Q24349 | Gdi |
| -0,892922719 | Q9VM11 | Q9VM11 | CG5973 |
| 0,06948789 | Q9VM35 | Q9VM35 | CG4502 |
| -0,067118327 | X2J786;Q9VMA7 | X2J786;Q9VMA7 | Tango1 |
| 0,250584284 | X2J8X6;Q9VMC9 | X2J8X6;Q9VMC9 | KFase |
| -0,470399857 | Q9VMI5 | Q9VMI5 | CG9135 |
| -0,289637884 | Q9VMQ9 | Q9VMQ9 | cl |
| -0,017791112 | Q9VN88 | Q9VN88 | Vps37B |
| -0,160638809 | Q9VN91;D2NUK | Q9VN91 | rtp |
| 0,064383825 | Q9VNE2 | Q9VNE2 | kra |
| 0,046127319 | Q9VNI8 | Q9VNI8 | Hpr1 |
| 0,338064194 | Q9VNZ8 | Q9VNZ8 | Als2 |
| -0,244277318 | Q9VP57 | Q9VP57 | pzg |
| 0,030005137 | Q9VQF7 | Q9VQF7 | Bacc |
| -0,040554047 | Q9VQI5 | Q9VQI5 | CG31694 |
| -0,355360667 | Q9VQL1;Q95TC0 | Q9VQL1;Q95TC0 | SerRS |
| 0,018438975 | Q9VR03 | Q9VR03 | CG15435 |
| -0,270755132 | Q9VR42 | Q9VR42 | CG3008 |
| 0,60575676 | Q9VR62 | Q9VR62 | CG17598 |
| 0,124997457 | Q9VRF3 | Q9VRF3 | CG15445 |
| -0,365127563 | Q9VRJ5 | Q9VRJ5 | CHMP2B |
| 0,173878988 | Q9VRL0 | Q9VRL0 | Cyt-c1 |
| 0,30424881 | Q9VRL8 | Q9VRL8 | blanks |
| -0,0253582 | X2JGB3;Q9VRP9 | X2JGB3;Q9VRP9 | Bre1 |
| -0,320108414 | Q9VSF2 | Q9VSF2 | MED24 |
| 0,651720047 | Q9VSL3;Q9VSL6 | Q9VSL3 | se |
| 0,183923721 | Q9VSX7 | Q9VSX7 | CG4080 |
| -0,171217601 | Q9VSY6 | Q9VSY6 | aay |
| -0,060455958 | Q9VSZ1;M9PF36 | Q9VSZ1;M9PF36;M9PEX2 | CG3529 |
| 0,415696462 | Q9VTH1 | Q9VTH1 | Ube3a |
| 0,136500676 | Q9VTK7;Q95S49 | Q9VTK7;Q95S49;E1JIE7 | Duba |
| -0,689578374 | Q9VTU4 | Q9VTU4 | eIF3l |
| -0,054256439 | Q9VTY2;Q8SXE8 | Q9VTY2;Q8SXE8 | CG10638 |
| 0,409269969 | Q9VTY6 | Q9VTY6 | vih |
| 0,25232633 | X2JCD3;Q9VU13 | X2JCD3;Q9VU13;E1NZE1 | CG42709 |
| -0,702584585 | Q9VU45 | Q9VU45 | Syx13 |
| -0,02259318 | Q9VU76;M9PFE5 | Q9VU76;M9PFE5;M9PFI2 | cmb |
| -0,156763713 | Q9VUH6;Q960G3 | Q9VUH6;Q960G3 | mop |
| -0,28524526 | Q9VUQ5;Q9VUQ | Q9VUQ5;Q9VUQ5-2;M9P | AGO2 |
| -0,249742508 | Q9VUX2;M9PI73 | Q9VUX2;M9PI73 | mib1 |

| | | | |
|---|---|---|---|
| -0,092315038 | Q9VVA7;Q8MQZ | Q9VVA7;Q8MQZ0 | TSG101 |
| 0,030478795 | Q9VW54 | Q9VW54 | Rpn1 |
| 0,0221138 | Q9VW59 | Q9VW59 | RhoGDI |
| 0,202218374 | Q9VW70;Q8SYR | Q9VW70;Q8SYR0 | CG7668 |
| -0,219485601 | Q9VW76 | Q9VW76 | CG14186 |
| 0,457823435 | Q9VWD9 | Q9VWD9 | Ubqn |
| -0,772594452 | Q9VWH4-2;Q9V | Q9VWH4-2;Q9VWH4 | Idh3a |
| 0,59916242 | Q9VWJ3 | Q9VWJ3 | CG32536 |
| 0,330722809 | X2JL49;Q9VWN | X2JL49;Q9VWN5 | CG7332 |
| 0,438116074 | X2JFP6;X2JL40;( | X2JFP6;X2JL40;Q9VWS3; | Rip11 |
| 0,319567362 | Q9VWV4;Q7K57 | Q9VWV4;Q7K571;Q9VW | CG32549 |
| -0,842435837 | Q9VX17 | Q9VX17 | CG6398 |
| 0,39215215 | Q9VXE8;Q9N9Z | Q9VXE8;Q9N9Z5 | Ubc7 |
| 0,210780462 | Q9VXJ7 | Q9VXJ7 | CG42353 |
| -0,351856232 | Q9VXM4 | Q9VXM4 | MSBP |
| -0,586604436 | Q9VXQ2;X2JDY | Q9VXQ2;X2JDY8;Q86NY | Graf |
| -0,230798721 | Q9VYG2 | Q9VYG2 | Bap60 |
| 0,067683538 | X2JEX7;Q9VYQ | X2JEX7;Q9VYQ8 | Usp7 |
| -0,938831965 | Q9VZI1;M9PE30 | Q9VZI1;M9PE30 | Chd64 |
| -0,232215246 | Q9VZS6 | Q9VZS6 | nSMase |
| -0,163276037 | Q9VZU7 | Q9VZU7 | Usp5 |
| -0,321706772 | Q9VZW1;Q8SZE | Q9VZW1;Q8SZE1 | scramb2 |
| 0,084868113 | Q9W073 | Q9W073 | CG12025 |
| -0,510450363 | Q9W0A7;Q4V3I2 | Q9W0A7;Q4V3I2;Q4V3M | RabX5 |
| -0,080002467 | Q9W0B3;Q29QY | Q9W0B3;Q29QY8 | CG7967 |
| 0,284442902 | Q9W0L7;M9PDK | Q9W0L7;M9PDK3 | Usp10 |
| 0,145650228 | Q9W1B0 | Q9W1B0 | gek |
| -0,00047493 | Q9W254;O44432 | Q9W254;O44432 | qkr58E-2 |
| -0,386535009 | Q9W2M0 | Q9W2M0 | Rbpn-5 |
| -0,305871964 | Q9W2R3 | Q9W2R3 | sktl |
| 0,599716822 | Q9W2S2;Q9VEG | Q9W2S2;Q9VEG5;Q7K1J | Atg8a |
| -0,17706871 | Q9W2V9 | Q9W2V9 | l(1)G0289 |
| -0,137491862 | Q9W329;Q95RK | Q9W329;Q95RK8 | Gga |
| -0,14723587 | Q9W350;Q9NHY | Q9W350;Q9NHY0 | c11.1 |
| -0,009466807 | Q9W369;D7RHE | Q9W369;D7RHE8 | t |
| -0,240905762 | Q9W392 | Q9W392 | CCT2 |
| 0,223169327 | Q9W3T9 | Q9W3T9 | CG3040 |
| 0,083726883 | Q9W414;Q8SZ19 | Q9W414;Q8SZ19;C1C3F8; | Rpt4 |
| -0,186683019 | Q9W4A6 | Q9W4A6 | OtopLa |
| 0,056781133 | X2JAH8;Q9W4E | X2JAH8;Q9W4E2;B7Z0W | rg |
| -0,526561101 | Q9W4P5 | Q9W4P5 | VhaAC39-1 |
| 0,115378698 | Q9W4X7 | Q9W4X7 | eIF3g1 |
| 0,554772695 | Q9W503;Q8SZ53 | Q9W503;Q8SZ53 | CG3091 |
| 0,021408081 | Q9XTL2 | Q9XTL2 | Stam |
| -0,190788905 | Q9XZ11;Q86NY( | Q9XZ11 | CG7139 |
| -0,13195165 | Q9XZ58;A0A0B4 | Q9XZ58;A0A0B4KHM2 | CSN5 |
| -0,19271787 | Q9XZ61 | Q9XZ61 | Uch-L5 |
| 0,871749878 | Q9XZ68 | Q9XZ68 | betaggt-II |
| 0,799763362 | Q9XZU1 | Q9XZU1 | Cse1 |

| | | | |
|---|---|---|---|
| -0,264623006 | Q9Y114 | Q9Y114 | CG8042 |
| 0,144685109 | R9PY16;H9BVD5 | R9PY16;H9BVD5;H9BVD | CG11700 |
| -0,088188171 | X2JDP4;X2JBS2; | X2JDP4;X2JBS2;Q8MR12 | inaE |

| CG number |
| --- |
| CG1618 |
| CG2118 |
| CG5659 |
| CG12403 |
| CG4629 |
| - |
| CG1519 |
| CG42639 |
| CG9668 |
| CG6424 |
| CG32486 |
| CG8053 |
| CG8048 |
| CG6957 |
| CG32687 |
| CG15117 |
| CG4178 |
| - |
| CG5192 |
| CG6776 |
| - |
| CG5685 |

| CG number |
| --- |
| CG1817 |
| CG12529 |
| - |
| CG6899 |
| CG9990 |
| CG10118 |
| CG15860 |
| CG2727 |
| CG3164 |
| CG33970 |
| CG16857 |
| CG7675 |
| CG12814 |
| CG2254 |
| CG13077 |
| CG1599 |
| CG1461 |
| CG17759 |
| CG17941 |
| CG4322 |
| CG1886 |
| CG33672 |

CG12787
CG13887
CG9761
CG31092
CG4494
CG8735
CG1151
CG13833
CG30445
CG31689
CG1105
CG6870
CG31705
CG33129
CG6461
CG45263
CG31100
CG2140
CG3131
CG8428
CG8385
CG4122
CG10106
CG8306
CG6126
CG10842
CG1943
CG11527
CG4122
CG42279
CG43140

| CG number |
|---|
| CG11198 |
| - |
| CG1516 |
| CG5271 |
| CG1516 |
| - |
| CG3937 |
| CG42574 |
| CG43427 |
| CG5685 |
| CG5099 |
| CG1945 |
| CG6315 |
| CG3682 |
| CG17883 |
| CG11140 |

CG2791
CG8732
CG8274
CG7904
CG1090
CG33094
CG9753
CG1744
CG45110
CG17838
CG10254
CG30021
CG3991
CG7563
CG3082
CG5820
CG2835
CG8399
CG42310
CG9484
CG33101
CG17632
CG2210
CG42595
CG1555
CG8024
CG1648
CG7220
CG18076
CG30069
CG5065
CG9277
CG13425
CG2969
CG12021
CG17520
CG9745
CG32858
CG2259
CG1799
CG11840
CG11328
CG5803
CG5962
CG42611
CG3312
CG10236
CG9528
CG44128
CG10737

CG14206
CG8285
CG34125
CG3861
CG31196
CG8280
CG32544
CG10279
CG10701
CG1009
CG9075
CG8892
CG1743
CG14039
CG17248
CG32626
CG43079
CG3504
CG1411
CG10076
CG18507
CG18495
CG8222
CG46149
CG3159
CG17654
CG33113
CG4590
CG11111
CG16916
CG34120
CG17352
CG14767
CG1633
CG12210
CG5670
CG11064
CG17291
CG42768
CG9704
CG14622
CG6944
CG17927
CG5711
CG3201
CG9086
CG42253
CG10188
CG3059
CG4357

CG3127
CG32138
CG12676
CG5125
CG6647
CG17723
CG8593
CG6302
CG7619
CG8893
CG10072
CG11958
CG17060
CG7070
CG2677
CG5436
CG4944
CG4550
CG2512
CG4898
CG12348
CG5125
CG7123
CG1560
CG2238
CG5670
CG3322
CG9327
CG7875
CG10045
CG3139
CG3725
CG9012
CG7558
CG16784
CG18345
CG15693
CG4260
CG2151
CG2679
CG42667
CG42670
CG9735
CG4609
CG10578
CG1112
CG3722
CG1877
CG15793
CG10596

CG5333
CG8983
CG13213
CG32451
CG31009
CG4167
CG11155
CG1882
CG12839
CG3821
CG18817
CG15118
CG15667
CG5170
CG4899
CG18410
CG17839
CG44252
CG17484
CG12013
CG10126
CG6783
CG32137
CG10237
CG32082
CG5594
CG8552
CG1121
CG12065
CG1443
CG1782
CG13920
CG3413
CG9317
CG2849
-
CG15792
CG12781
CG33653
CG10047
CG10370
CG4599
CG11949
CG1512
CG7946
CG31038
CG10244
CG11089
CG17121
CG6969

CG5685
CG34157
CG7702
CG5823
CG18740
CG4800
CG9474
CG9448
CG9444
CG7415
CG9244
CG10602
CG15151
CG9934
CG6181
CG5355
CG16708
CG2929
CG17660
CG1753
CG5064
CG10960
CG6603
CG7739
CG42279
CG14226
CG6617
CG8188
CG34325
CG4420
CG9968
CG8231
CG15211
CG9139
CG3842
CG16721
CG9634
CG15437
CG5803
CG15792
CG8295
CG8440
CG2244
CG10225
CG4719
CG8180
CG8400
CG12769
CG3401
CG8392

CG8085
CG8416
CG2827
CG5009
CG6891
CG15279
CG14792
CG11567
CG9339
CG8937
CG10837
CG1956
CG42677
CG32697
CG32397
CG5654
CG42278
CG10776
CG10289
CG10079
CG5748
CG10119
CG4937
CG5938
CG10939
CG2666
CG6092
CG5520
CG11857
CG6977
CG9248
CG7082
CG3036
CG12202
CG3001
CG7207
CG2049
CG18375
CG1782
CG1107
CG12489
CG18290
CG11242
CG44193
CG7363
CG42388
CG6392
CG3573
CG5373
CG10328

CG5634
CG9641
CG9242
CG8310
CG43140
CG6523
CG3756
CG3618
CG10642
CG6418
CG32177
CG31732
CG3481
CG8709
CG8068
CG9765
CG31136
CG11901
CG42321
CG6203
CG6963
CG12001
CG6051
CG1539
CG17800
CG8715
CG16901
CG8529
CG11804
CG8566
CG9218
CG5174
CG2050
CG4634
CG8325
CG11856
CG42344
CG4001
CG8487
CG6383
CG10149
CG42236
CG16910
CG44154
CG17870
CG7843
CG10897
CG3454
CG8604
CG30092

CG44162
CG3615
CG30015
CG42724
CG2520
CG3416
CG8787
CG6303
CG6622
CG3379
CG6908
CG7467
CG5206
CG7583
CG5695
CG5499
CG5730
CG12878
CG6588
CG11888
CG13645
CG17838
CG7425
CG1528
CG44010
CG5555
CG7917
CG12114
CG1407
CG34407
CG12131
CG10228
CG8889
CG8472
CG13349
CG34399
CG6702
CG8739
CG9450
CG17921
CG10497
CG5594
CG14670
CG1088
CG8884
CG5085
CG1448
CG6668
CG5289
CG34371

CG9351
CG18250
CG10851
CG7340
CG32555
CG3724
CG12276
CG10206
CG9764
CG34138
CG7897
CG2064
CG12159
CG8057
CG9033
CG17509
CG8439
CG3845
CG17019
CG8443
CG8912
CG6665
CG30122
CG8201
CG34342
CG6835
CG8938
CG4527
CG4280
CG7254
CG43756
CG3762
CG18069
CG8977
CG5374
CG9983
CG1873
CG1453
CG32593
CG18102
CG18000
-
CG15118
CG32000
CG10811
CG2759
CG11533
CG9916
CG4212

CG12008
CG14998
CG15015
CG42734
CG32031
CG8104
CG17816
CG43140
CG31158
CG5212
CG15894
CG34417
CG12449
CG6186
CG1554
CG7399
CG10084
CG42492
CG9258
CG14296
CG31030
CG4603
CG3811
-
CG2903
CG5784
CG5020
CG4244
CG3523
CG3696
CG7228
CG18803
CG7176
CG5771
CG9181
CG4236
CG3665
CG3400
CG12019
CG2947
CG6395
CG12214
CG10550
CG9441
CG13366
CG4264
CG3897
CG6342
CG1646
CG6058

CG12286
CG6778
CG42341
CG8363
CG5284
CG6664
CG17453
CG12789
CG11324
CG6395
CG5605
CG16944
CG14207
CG16936
CG3265
CG5175
CG33991
CG11129
CG6605
CG31884
CG34126
CG12559
CG5738
CG8201
CG40129
CG10377
CG3694
CG10283
CG7611
CG9695
CG8110
CG43781
CG17369
CG12749
CG43398
CG32490
CG32490
CG1307
CG33950
CG5227
CG3620
CG1634
CG18319
CG10417
CG5825
CG4464
CG7035
CG32717
CG42314
CG42314

CG9153
CG4376
CG3989
CG6811
CG11009
CG4147
CG5834
CG3595
CG6779
CG43140
CG8588
CG9802
CG6095
CG5686
CG2621
CG2096
CG17245
CG6959
CG8625
CG7471
CG17947
CG18801
CG11154
CG17023
CG4265
CG4747
CG42768
CG42671
CG43128
CG8582
CG5850
CG6549
CG1099
CG7546
CG3849
CG2637
CG3921
CG8282
CG43479
CG4463
CG43388
CG9819
CG9088
CG43225
CG9674
CG4262
CG6143
CG34387
CG11924
CG5366

CG1977
CG33484
CG7740
CG1242
CG7507
CG10103
CG8556
CG3077
CG8532
CG7927
CG6831
CG10882
CG10686
CG7111
CG3779
CG31605
CG6382
CG10363
CG9326
CG17332
CG10223
CG4690
CG2086
CG8663
CG8571
CG44195
CG15811
CG7828
CG5870
CG5433
CG6084
CG11793
CG3680
CG40045
CG42641
CG7490
CG5723
CG11579
CG2890
CG3039
CG1822
CG11654
CG10522
CG4083
CG10345
CG2038
CG2207
CG3320
CG3269
CG8287

CG3924
CG1489
CG6446
CG3299
CG2924
CG2925
CG8200
CG8649
CG2720
CG11219
CG7057
CG7398
CG5915
CG2714
CG32505
CG4900
CG3747
CG4199
CG7961
CG9677
CG3637
CG5838
CG11282
CG3204
-
CG16963
CG5345
CG11143
-
CG4183
CG4466
CG2985
CG8409
CG4027
CG5650
CG9206
CG6518
CG10360
CG2204
CG7765
CG6871
CG9193
CG8874
CG5119
CG9579
CG9579
CG3422
CG1483
CG17759

CG8770
CG3612
CG40452
CG44746
CG8705
CG9946
CG7641
CG6223
CG4254
CG4314
CG9901
CG6174
CG3234
CG10497
CG8956
CG7434
CG8251
CG6720
CG8284
CG4299
CG6988
CG17903
CG33859
CG5870
CG43443
CG4211
-
CG4125
CG1677
CG4994
CG3689
CG31731
CG8547
CG9057
CG42327
CG1651
CG1511
CG17078
CG9261
CG6625
CG5330
CG10653
CG12473
CG12532
-
CG8293
CG6806
CG6384
CG5655
CG9633

CG7269
CG11537
CG3662
CG2257
CG12196
CG9113
CG15624
CG31729
CG14184
CG42314
CG42314
CG7736
CG8711
CG2177
CG2025
CG7656
CG40178
CG5625
CG3907
CG6159
CG45782
CG1709
CG6827
CG1732
CG40485
CG10198
CG11210
CG5604
CG18608
CG12891
CG8830
CG33144
CG3798
CG9264
CG11761
CG12845
CG5469
CG1532
CG5081
CG32066
CG8468
CG12505
CG7222
CG8839
CG13349
CG44246
CG8258
CG11139
CG5721
CG5788

CG10392
CG18076
CG7528
CG3530
CG10230
CG1341
CG5495
CG2331
CG2072
CG7865
CG31006
CG5191
CG3903
CG5547
CG15828
CG9391
CG10523
CG6718
CG1640
CG1561
CG42672
CG5014
CG8024
CG3869
CG10173
CG5830
CG5722
CG31012
CG12052
CG33103
CG13388
CG10712
CG5203
CG9941
CG34404
CG1467
CG12199
CG32640
CG14648
CG11061
CG6148
CG9535
CG6995
CG1910
CG3210
CG2201
CG1883
CG11980
CG5603
CG13388

CG31641
CG32056
CG7656
CG32581
CG1228
CG16932
CG17645
CG6450
CG5639
CG4869
CG6756
CG5166
CG42258
CG9330
CG32850
CG31992
CG14830
CG8209
CG12156
CG12024
CG8024
CG8055
CG4659
CG6512
CG12085
CG5653
CG10922
CG6214
CG13391
CG9985
CG11621
CG3590
CG6151
CG9911
CG1059
CG18212
-
CG12008
CG5105
CG8223
CG15669
CG32435
CG9138
CG9412
CG9195
CG13387
CG5000
CG3988
CG18174
CG5387

CG3664
CG1591
CG4930
CG6987
CG1836
CG4180
CG1100
CG4170
CG6582
CG10687
CG1710
CG7911
CG7896
CG11899
CG4849
CG12876
CG5643
CG5447
CG4673
CG5112
CG33095
CG7662
CG31118
CG13603
CG7023
CG5346
CG7077
CG6656
CG18617
CG7217
CG4225
CG3731
CG3061
CG3028
CG8863
CG6359
CG6946
CG14709
CG10703
CG9373
CG8149
CG11982
CG8351
CG8036
CG3223
CG2767
CG10277
CG10413
CG5094
CG16974

CG5787
CG5525
CG6287
CG5168
CG5384
CG4968
CG5676
CG4912
CG4535
CG4422
CG5973
CG4502
CG11098
CG9542
CG9135
CG11024
CG1115
CG10233
CG2922
CG2031
CG7158
CG7752
CG9894
CG31694
CG17259
CG15435
CG3008
CG17598
CG15445
CG4618
CG4769
CG10630
CG10542
CG7999
CG6781
CG4080
CG3705
CG3529
CG6190
CG6091
CG5642
CG10638
CG10682
CG42709
CG11278
CG10732
CG9311
CG7439
CG5841

CG9712
CG7762
CG7823
CG7668
CG14186
CG14224
CG12233
CG32536
CG7332
CG6606
CG32549
CG6398
CG4443
CG42353
CG9066
CG8948
CG4303
CG1490
CG14996
CG12034
CG12082
CG1893
CG12025
CG7980
CG7967
CG32479
CG4012
CG5821
CG4030
CG9985
CG32672
CG2221
CG3002
CG12132
CG12120
CG7033
CG3040
CG3455
CG42492
CG44835
CG2934
CG8636
CG3091
CG6521
CG7139
CG14884
CG3431
CG18627
CG13281

CG8042
CG11700
CG33174

| Description |
|---|
| comatose |
| Methylcrotonoyl-CoA carboxylase 1 |
| ariadne 1 |
| Vacuolar H[+] ATPase 68kD subunit 1 |
| - |
| mdg3 element ORF |
| Proteasome alpha7 subunit |
| Prophenoloxidase 1 |
| Rhodopsin 4 |
| - |
| - |
| eukaryotic translation initiation factor 1A |
| Vacuolar H[+] ATPase 44kD subunit |
| Oscillin |
| - |
| - |
| Larval serum protein 1 beta |
| gypsy element gag |
| Rhodopsin 6 |
| Glutathione S transferase O3 |
| Prophenoloxidase 1 (fragment) |
| Na/Ca-exchange protein |

| Description |
|---|
| Protein tyrosine phosphatase 10D |
| Zwischenferment |
| gag-int-pol |
| Protein tyrosine phosphatase 4E |
| snustorr |
| pale |
| painless |
| epithelial membrane protein |
| - |
| - |
| borderless |
| - |
| - |
| Lipid droplet subset dehydrogenase 1 |
| - |
| Vesicle-associated membrane protein 7 |
| - |
| G protein alpha q subunit |
| dachsous |
| moody |
| ATP7 |
| - |

hoepel1
-
Neprilysin 2
Lipophorin receptor 2
smt3
Lunapark
Osiris 6
-
Tyrosine decarboxylase 1
-
-
-
-
-
gamma-glutamyl transpeptidase
-
-
Cytochrome b5
Dual oxidase
spinster
ADP ribosylation factor at 79F
silver
Tetraspanin 42Ee
-
-
Cytochrome P450-4p1
-
Tiggrin
silver
Nedd4
polychaetoid

| Description |
|---|
| Acetyl-CoA carboxylase |
| Avidin |
| Pyruvate carboxylase |
| Ubiqutin-40S ribosomal protein S27A |
| Pyruvate carboxylase |
| biotin-tagged ubiquitin |
| cheerio |
| circadian trip |
| smallish |
| Na/Ca-exchange protein |
| musashi |
| fat facets |
| female lethal d |
| Phosphatidylinositol 4-phosphate 5-kinase at 59B |
| - |
| Aldehyde dehydrogenase type III |

CD98 heavy chain
Acyl-CoA synthetase long-chain
Megator
punt
-
Syndapin
Adenosine receptor
Chaoptin
tau
Syncrip
-
menage a trois
tripeptidyl-peptidase II
Calpain-A
lethal (2) k09913
Gp150
G protein alpha s subunit
-
prominin
hyperplastic discs
N-ethylmaleimide-sensitive factor 2
brown
abnormal wing discs
unextended
cinnabar
Rab32
-
-
short stop
-
-
beta-Tubulin at 56D
Heterogeneous nuclear ribonucleoprotein K
ABC transporter expressed in trachea
Patj
casein kinase IIalpha
D1 chromosomal protein
singed
Glutamate-cysteine ligase catalytic subunit
raspberry
Signal peptide peptidase
Na[+]/H[+] hydrogen exchanger 3
Fasciclin 3
Arrestin 2
Megalin
RNA-binding protein 4F
Laminin A
real-time
Src oncogene at 42A
-

Ribosomal protein S10b
bride of sevenless
-
knockdown
14-3-3epsilon
eukaryotic translation elongation factor 1 alpha 1
-
Rm62
Moesin
Puromycin sensitive aminopeptidase
eukaryotic translation initiation factor 4A
-
Glutamine synthetase 2
quick-to-court
neuronal Synaptobrevin
AMP deaminase
neuromusculin
inactivation no afterpotential D
Collapsin Response Mediator Protein
spire
-
Proteasome alpha1 subunit
PDGF- and VEGF-receptor related
Fatty acid transport protein 1
Excitatory amino acid transporter 2
Enolase
Reticulon-like1
Innexin 2
retinal degeneration B
Regulatory particle triple-A ATPase 3
-
CUB and LDLa domain
-
thioredoxin peroxidase 1
Synaptobrevin
Na pump alpha subunit
apolipophorin
Protein phosphatase 2A at 29B
Muscle-specific protein 300 kDa
Neurotactin
Dishevelled Associated Activator of Morphogenesis
Lamin
Myosin heavy chain
Arrestin 1
Myosin light chain cytoplasmic
Ubr1 ubiquitin ligase
Na[+]-driven anion exchanger 1
-
NTPase
sodium chloride cotransporter 69

Phosphoglycerate kinase
Formin-like
echinoid
neither inactivation nor afterpotential C
porin
Zinc transporter 63C
quemao
Prefoldin 2
Regulatory particle non-ATPase 10
Glyceraldehyde 3 phosphate dehydrogenase 2
sugarless
Calnexin 99A
Rab10
Pyruvate kinase
eukaryotic translation initiation factor 2B subunit beta
Heat shock protein 68
ciboulot
neither inactivation nor afterpotential E
alpha-Tubulin at 84D
Tropomyosin 1
Shaker
neither inactivation nor afterpotential C
LanB1
myospheroid
eukaryotic translation elongation factor 2
Na pump alpha subunit
Laminin B2
Proteasome alpha3 subunit
transient receptor potential
Glutathione S transferase D1
Synaptotagmin 1
Sarco/endoplasmic reticulum Ca(2+)-ATPase
Clathrin heavy chain
Actin-related protein 3
purple
transient receptor potential-like
Ribosomal protein S20
Adaptor Protein complex 2, alpha subunit
Thioredoxin reductase-1
goliath
retinal degeneration A
pasilla
Tryptophanyl-tRNA synthetase
failed axon connections
DnaJ-like-1
alpha-Esterase-7
shotgun
Cullin 1
Downstream of raf1
Msr-110

toys are us
Endoplasmic reticulum p60
F-box and leucine-rich repeat protein 6
Secretory Pathway Calcium atpase
Cadherin 99C
Heat shock gene 67Ba
-
pummelig
Tetraspanin 42En
Aspartyl-tRNA synthetase
Tetraspanin 42Ea
-
Smad anchor for receptor activation
Dodeca-satellite-binding protein 1
Photoreceptor dehydrogenase
Uracil-DNA degrading factor
-
Fatty acid transport protein 2
Adherens junction protein p120
PHGPx
-
fatty acid binding protein
-
-
Insulin receptor substrate 53 kDa
kazachoc
Phosphatidic Acid Phospholipase A1
alpha-Esterase-8
-
waterproof
Ubiquitin activating enzyme 1
-
windpipe
Carcinine transporter
Ras-like protein A
HMS-Beagle element pol
zipper
nahoda
Calcium-dependent secretion activator
Synaptotagmin 4
Regulatory particle triple-A ATPase 5
Tetratricopeptide repeat protein 2
coracle
Cullin 2
-
-
Cadherin 96Ca
-
-
cardinal

Na/Ca-exchange protein
Dystrophin
-
-
moira
Translationally controlled tumor protein
Synaptosomal-associated protein 24kDa
trabid
-
Dipeptidyl aminopeptidase III
Mitochondrial aconitase 1
-
reduced ocelli
-
Ge-1
-
Ceramide kinase
Pi4KIIalpha
-
Cystathionine beta-synthase
Signal recognition particle protein 68
-
Hsc70Cb
-
Nedd4
domeless
-
-
-
rings lost
Annexin B11
Chaperonin containing TCP1 subunit 6
-
Rabaptin-5-associated exchange factor for Rab5
-
-
gone early
modifier of rpr and grim, ubiquitously expressed
Fasciclin 3
zipper
Myelodysplasia/myeloid leukemia factor
Lissencephaly-1
Metastasis associated 1-like
Ran binding protein 3
tankyrase
-
caspar
-
beta-Tubulin at 60D
Proteasome beta1 subunit

Related to the N terminus of tre oncogene
Rho1
Transaldolase
-
-
-
stubarista
Cytochrome P450 reductase
skywalker
Heat shock protein cognate 1
eukaryotic translation initiation factor 4B
Rap1 GTPase
wing blister
Protein tyrosine phosphatase Meg2
javelin
ypsilon schachtel
cornetto
wishful thinking
fiery mountain
Epidermal growth factor receptor
Heat shock factor
Lamin C
Rho GTPase activating protein at 15B
-
-
krotzkopf verkehrt
Dak1
Glycoprotein 93
-
Cadherin 87A
Phospholipase D family member 3
papi
-
N(alpha)-acetyltransferase 15/16
Hexokinase A
ceramide transfer protein
Protein kinase N
Ankyrin-repeat, SH3-domain, and Proline-rich-region containing Protein
Ubiquitin activating enzyme 1
auxilin
defense repressor 1
Actin 87E
tubulin-binding cofactor B
Chondrocyte-derived ezrin-like domain containing protein
world cup
Nostrin
CENP-meta
Oculocerebrorenal syndrome of Lowe
Phosphotidylinositol 3 kinase 59F
nonA-like

distracted
-
burgundy
Vacuolar H[+] ATPase 36kD subunit 3
polychaetoid
-
-
-
Kinesin-like protein at 64D
-
Nedd4 family interacting protein
yuri gagarin
Alcohol dehydrogenase
Lipin
Suppressor of variegation 2-10
transforming acidic coiled-coil protein
Syntaxin 1A
eukaryotic translation elongation factor 1 gamma
ATPase 8A
Fmr1
gilgamesh
spartin
-
tropomodulin
Down syndrome cell adhesion molecule 1
lingerer
squid
Dystrobrevin
ced-6
uncoordinated-104
smooth
-
modulo
Nucleosome remodeling factor - 38kD
Tetratricopeptide repeat domain 7
Nucleoporin 358kD
bruchpilot
Phosphofructokinase
gartenzwerg
crumbs
Regulatory particle non-ATPase 6
Ran-binding protein M
kenny
klaroid
14-3-3zeta
Arsenic resistance protein 2
toutatis
Histidine decarboxylase
Amphiphysin
jitterbug

Stretchin-Mlck
Autophagy-related 9
-
-
like-AP180
Regulatory particle non-ATPase 8
Additional sex combs
BIR repeat containing ubiquitin-conjugating enzyme
Protein C kinase 53E
Histone H4 replacement
-
osa
bonus
C-terminal Binding Protein
jaguar
Histone H2A variant
Annexin B9
barentsz
Fasciclin 1
Regulatory particle non-ATPase 2
Nicotinamide mononucleotide adenylyltransferase
Syncrip
effete
Coat Protein (coatomer) gamma
kokopelli
-
Nucleoplasmin
spn-F
-
Not1
eukaryotic translation initiation factor 3 subunit j
Pcf11 cleavage and polyadenylation factor subunit
Metallophosphoesterase
Calmodulin
Regulatory particle non-ATPase 13
NADPH oxidase
Calbindin 53E
stambha A
tudor
HMG protein Z
Syndecan
kazachoc
Holocarboxylase synthetase
Vacuolar H[+]-ATPase 26kD subunit
Synapse-associated protein 47kD
Sirtuin 2
Innexin 3
atlastin
Regulatory particle triple-A ATPase 2
sidestep V

falafel
Dystroglycan
B52
granny smith
Rho GTPase activating protein p190
Phosphogluconate dehydrogenase
Activator of SUMO 1
nop5
yurt
-
Glycoprotein 210 kDa
-
-
alicorn
Tetraspanin 47F
precocious dissociation of sisters 5
Chaperonin containing TCP1 subunit 5
NAT1
-
clueless
P-element somatic inhibitor
-
-
par-1
-
Glutathione synthetase 1
Glutathione S transferase S1
Sterile20-like kinase
croquemort
Glycogen phosphorylase
Slowpoke binding protein
Vacuolar H[+] ATPase 68 kDa subunit 2
Calcium/calmodulin-dependent protein kinase II
Chaperonin containing TCP1 subunit 3
Chaperonin containing TCP1 subunit 1
Heterogeneous nuclear ribonucleoprotein at 98DE
eukaryotic translation elongation factor 1 alpha 2
Kinesin-like protein at 10A
Flotillin 2
shibire
short wing
gypsy element gag
-
anne boleyn
eukaryotic translation initiation factor 4G1
white
Asator
Cyclophilin 1
Rab14

karst
ensconsin
Cdc42-interacting protein 4
Ankyrin 2
Arginine kinase
nudE
-
polychaetoid
Exchange factor for Arf 6
Pellino
-
-
Glutamine:fructose-6-phosphate aminotransferase 1
Transferrin 1
RNA polymerase II 215kD subunit
Henna
second mitotic wave missing
Otopetrin-like a
nervana 1
Endophilin A
-
-
Organic anion transporting polypeptide 30B
Polyprotein
Hepatocyte growth factor regulated tyrosine kinase substrate
Mapmodulin
Cytoplasmic linker protein 190
Suppressor of deltex
Fatty acid synthase 1
kismet
peste
Presenilin
Isocitrate dehydrogenase
Rab11
Protein tyrosine phosphatase 61F
Chromatin assembly factor 1, p55 subunit
Fasciclin 2
6-phosphofructo-2-kinase
Cdc37
Hsc/Hsp70-interacting protein related
Cysteine string protein
mulet
-
Punch
-
Heat shock protein cognate 4
bloated tubules
Iron regulatory protein 1B
-
Aldolase 1

karmoisin
Glycyl-tRNA synthetase
Protein kinase, cAMP-dependent, regulatory subunit type 1
PAPS synthetase
Chloride channel-c
-
Cyp317a1
scavenger receptor acting in neural tissue and majority of rhodopsin is absent
homer
Cysteine string protein
eukaryotic translation release factor 1
stress-sensitive B
-
Glutathione S transferase E12
Eb1
kugelkern
nuclear fallout
Yolk protein 3
Bicaudal D
thioredoxin-2
-
rolled
lola like
par-1
G protein-coupled receptor kinase 1
Heterogeneous nuclear ribonucleoprotein at 27C
G protein subunit gamma at 30A
-
-
Disabled
sunday driver
-
Vacuolar H[+]-ATPase 55kD subunit
Heterogeneous nuclear ribonucleoprotein at 87F
scribble
complexin
complexin
-
terribly reduced optic lobes
sidekick
no receptor potential A
Neuroglian
bendless
-
Histone H3.3A
Ribosomal protein S19a
cap binding protein 80
stardust
plasma membrane calcium ATPase
plasma membrane calcium ATPase

HECT and RLD domain containing E3 ubiquitin ligase 4
alpha actinin
PAICS bifunctional enzyme
Rho GTPase activating protein at 68F
WW domain binding protein 2
Heat shock 70-kDa protein cognate 3
Hsp70Bbb
spaghetti squash
Ribosomal protein S3
polychaetoid
pastrel
Structural maintenance of chromosomes 3
Exocyst 84
chico
shaggy
flapwing
Plexin B
-
Imitation SWI
Histone deacetylase 1
alpha Catenin
Ku80
ATP synthase, beta subunit
Dead box protein 80
Ubiquitin carboxy-terminal hydrolase
Nucleosome-destabilizing factor
Muscle-specific protein 300 kDa
-
Shaker cognate b
Sh3beta
-
four way stop
Dynamin associated protein 160
-
Lasp
Female sterile (2) Ketel
bark beetle
Sorting nexin 6
nervous wreck
Heat shock protein 23
Hyperkinetic
Calcineurin A at 14F
little imaginal discs
axotactin
-
embryonic lethal abnormal vision
Protein on ecdysone puffs
futsch
Chorion factor 2
Cullin-associated and neddylation-dissociated 1

alpha Spectrin
zormin
prominin-like
Heat shock protein 83
Dynein heavy chain 64C
-
Rac2
Sorting nexin 21
liquid facets
Nuclear localized protein 1
rhea
Secretory 24CD
trailer hitch
Receptor of activated protein kinase C 1
numb
Basigin
eukaryotic translation release factor 3
Thioester-containing protein 4
varicose
Vacuolar H[+]-ATPase SFD subunit
Topoisomerase 2
Tetraspanin 5D
draper
nervana 3
smallminded
-
Ras opposite
beta-Amyloid precursor protein binding protein 1
beta Spectrin
Kinesin light chain
-
Superoxide dismutase 1
HP1 and insulator partner protein 1
-
Regulatory particle non-ATPase 3
Ribosomal protein LP0
Tenascin major
armadillo
Protein phosphatase 4 regulatory subunit 2-related protein
optic ganglion reduced
bifocal
Adenosylhomocysteinase
sticky
Mo25
-
COP9 signalosome subunit 7
Decondensation factor 31
Rab1
Rab2
Rab8

Chip
Regulatory particle triple-A ATPase 6
Semaphorin 1b
Vinculin
-
noisette
Flotillin 1
Fimbrin
Stress induced phosphoprotein 1
PIP82
Adaptor Protein complex 2, mu subunit
Transportin
Rab7
cramped
Protein phosphatase 19C
Iron regulatory protein 1A
Excitatory amino acid transporter 1
-
Coat Protein (coatomer) alpha
eukaryotic translation initiation factor 3 subunit e
Cortactin
DNA replication-related element factor
capricious
Ras-associated protein 2-like
Idefix element gag
Crystallin
Ecdysone-induced protein 55E
myo-inositol-1-phosphate synthase
Histone H2B
Heat shock protein 26
Heat shock protein 27
Yolk protein 1
Suppressor of variegation 205
Actin 5C
Protein phosphatase 1 at 87B
Dynactin 1, p150 subunit
inactivation no afterpotential C
refractory to sigma P
G protein alpha o subunit
Kinesin heavy chain
Catalase
Proliferating cell nuclear antigen
FER tyrosine kinase
poly(A) binding protein
Annexin B10
Annexin B10
Proteasome alpha4 subunit
Microtubule-associated protein 205
G protein alpha q subunit

G protein beta-subunit 76C
bellwether
Synaptosomal-associated protein 25kDa
retinal degeneration C
peanut
eukaryotic translation initiation factor 2 subunit alpha
Neurocalcin
Coat Protein (coatomer) β
twinstar
scarlet
Actin-related protein 2
Actin-related protein 1
timeless
Syndecan
Adenosylhomocysteinase like 2
Ribosomal protein L22
Phosphoglucose isomerase
Ubiquitin conjugating enzyme 2
Ubiquitin conjugating enzyme 4
Set
Protein disulfide isomerase
Cytochrome c proximal
His2A:CG33859
beta Spectrin
hu li tai shao
no on or off transient A
Doc element gag
roughest
-
Mitochondrial phosphate carrier protein 2
Cleavage and polyadenylation specific factor 5
Engulfment ABC Transporter in the ovary
-
Lipid storage droplet-2
-
Ankyrin
Eph receptor tyrosine kinase
-
nervana 2
alpha Soluble NSF attachment protein
Nucleosome assembly protein 1
hook
stoned B
Adaptor Protein complex 1/2, beta subunit
flea element polyprotein
Death-associated inhibitor of apoptosis 2
Larval serum protein 2
Centrosomal protein 190kD
Repressor splicing factor 1
Replication Protein A 70

Helicase at 25E
-
-
Ubiquitin conjugating enzyme E2H
eggless
Adaptor Protein complex 1, gamma subunit
hoepel2
-
-
plasma membrane calcium ATPase
plasma membrane calcium ATPase
Syntaxin 6
Cullin 4
Zinc/iron regulated transporter-related protein 102B
Nardilysin
-
lethal (3) 80Fg
Vacuolar protein sorting 35
-
Secretory 10
loss of visual transmission
Vacuolar H[+] ATPase 100kD subunit 1
Neurexin IV
GABA transporter
-
Nucleoporin 98-96kD
Transmembrane protein 63
Ubiquitin fusion-degradation 4-like
proliferation disrupter
withered
Deubiquitinating apoptotic inhibitor
-
N-methyl-D-aspartate receptor-associated protein
torn and diminished rhabdomeres
translin
Tetraspanin 42Ef
GDI interacting protein 3
-
Syntaxin 7
-
-
Activity-regulated cytoskeleton associated protein 1
-
-
Regulatory particle non-ATPase 13
Activating transcription factor-2
Chaperonin containing TCP1 subunit 8
p47
-
Ubiquitin conjugating enzyme 10

super sex combs
short stop
Ubiquitin-like activating enzyme 2
-
Regulatory particle non-ATPase 9
Regulatory particle triple-A ATPase 1
Thioredoxin-like
-
Mitotic arrest-deficient 1
PNGase-like
slow termination of phototransduction
-
Gliotactin
Phosphoethanolamine cytidylyltransferase
Apolipoprotein lipid transfer particle
-
parkin
calcium-independent phospholipase A2 VIA
-
-
-
VAMP-associated protein 33kDa
Rab32
Mitochondrial assembly regulatory factor
Bestrophin 2
-
Niemann-Pick type C-1a
CIN85 and CD2AP related
longitudinals lacking
Papilin
A kinase anchor protein 200
Chromator
STIP1 homology and U-box containing protein 1
-
-
Syntaxin 16
kekkon 5
-
lost
Golgi matrix protein 130 kD
Putative Achaete Scute Target 1
mummy
Scaffold attachment factor B
-
Dynamin related protein 1
-
Ribosomal protein S7
-
Cylindromatosis
A kinase anchor protein 200

stathmin
scramblase 1
-
-
Protein tyrosine phosphatase Meg
Epidermal growth factor receptor pathway substrate clone 15
Phosphoglyceromutase 87
lava lamp
-
beta-Tubulin at 97EF
Translocase of outer membrane 70
Ataxin-2
-
-
-
gawky
-
-
Rab39
-
Rab32
shrub
Signal recognition particle protein 54k
-
half pint
-
La autoantigen-like
Multidrug-Resistance like Protein 1
Alanyl-tRNA synthetase
skittles
Phosphotidylinositol 3 kinase 68D
Adenylosuccinate Lyase
flower
-
Karyopherin beta 3
aluminum tubes
HMS-Beagle element gag
karst
Phospholipase A2 activator protein
-
Misexpression suppressor of KSR 2
chromosome bows
uninflatable
rasputin
Scamp
embargoed
mini spindles
gamma Soluble NSF attachment protein 1
Regulatory particle non-ATPase 11
Cdk5 activator-like protein

Rab5
Proteasome regulator gamma
Endonuclease G inhibitor
Splicing factor 2
Rad23
Cytokine induced apoptosis inhibitor 1
Regulatory particle non-ATPase 5
vasa intronic gene
cassowary
Asparaginyl-tRNA synthetase
Host cell factor
Nucleophosmin
-
-
-
ALG-2 interacting protein X
widerborst
-
Nuclear protein localization 4
-
-
veli
RabX4
-
Ubiquitin-specific protease 12/46
-
retinol dehydrogenase B
-
Vacuolar H[+] ATPase 100kD subunit 2
Peroxiredoxin 5
Heavy metal tolerance factor 1
Ubiquinol-cytochrome c reductase core protein 1
-
Inositol polyphosphate 1-phosphatase
DnaJ-like-2
Sorting nexin 3
glorund
Multidrug resistance protein 4
GRIP and coiled-coil domain containing 88 kDa
rumpelstiltskin
-
Iruka
Chaperonin containing TCP1 subunit 7
-
-
-
godzilla E3 ubiquitin protein ligase
-
small glutamine-rich tetratricopeptide containing protein
-

-
Chaperonin containing TCP1 subunit 4
-
WD repeat and FYVE domain containing 2
Ubiquitin specific protease 14
-
-
eukaryotic translation elongation factor 1 delta
FK506-binding protein 59kD
GDP dissociation inhibitor
-
-
Transport and Golgi organization 1
Kynurenine formamidase
-
clot
Vacuolar protein sorting 37B
retinophilin
krasavietz
Hpr1
Amyotrophic lateral sclerosis 2
putzig
Bacchus
-
Seryl-tRNA synthetase
-
-
-
-
Charged multivesicular body protein 2b
Cytochrome c1
blanks
Bre1
Mediator complex subunit 24
sepia
-
astray
-
Ubiquitin protein ligase E3A
Deubiquitinating enzyme A
eukaryotic translation initiation factor 3 subunit l
-
vihar
-
Syntaxin 13
combover
myopic
Argonaute 2
mind bomb 1

Tumor susceptibility gene 101
Regulatory particle non-ATPase 1
RhoGDI
-
-
Ubiquilin
Isocitrate dehydrogenase 3a
-
-
Rab11 interacting protein
-
-
Ubiquitin conjugating enzyme 7
-
membrane steroid binding protein
GTPase regulator associated with FAK
Brahma associated protein 60kD
Ubiquitin-specific protease 7
Chd64
neutral sphingomyelinase
Ubiquitin specific protease 5
scramblase 2
-
RabX5
-
Ubiquitin specific protease 10
genghis khan
quaking related 58E-2
Rabaptin-5
skittles
Autophagy-related 8a
lethal (1) G0289
Golgi-localized, gamma-adaptin ear containing, ARF binding protein
c11.1
tan
Chaperonin containing TCP1 subunit 2
-
Regulatory particle triple-A ATPase 4
Otopetrin-like a
rugose
Vacuolar H[+] ATPase AC39 subunit 1
eukaryotic translation initiation factor 3 subunit g1
-
Signal transducing adaptor molecule
-
COP9 signalosome subunit 5
Ubiquitin carboxy-terminal hydrolase L5
beta subunit of type II geranylgeranyl transferase
Chromosome segregation 1

-
Ubiquitin
inactivation no afterpotential E
